# Does sexual conflict contribute to the evolution of novel warning patterns?

**DOI:** 10.1101/2022.05.16.492082

**Authors:** Alexander E. Hausmann, Marília Freire, Sara A. Alfthan, Chi-Yun Kuo, Mauricio Linares, Owen McMillan, Carolina Pardo-Diaz, Camilo Salazar, Richard M. Merrill

**Author notes:** AEH and MF contributed equally to this work.

## Abstract

Why warning patterns are so diverse is an enduring evolutionary puzzle. Because predators associate particular patterns with unpleasant experiences, an individual’s predation risk should decrease as the local density of its warning pattern increases, promoting pattern monomorphism. Distasteful *Heliconius* butterflies are known for their diversity of warning patterns. Here, we explore whether interlocus sexual conflict can contribute to their diversification. Male *Heliconius* use warning patterns as mating cues, but mated females may suffer costs if this leads to harassment, favoring novel patterns. Using simulations, we show that drift alone is unlikely to cause pattern diversification, but that sexual conflict can assist such process. We also find that genetic architecture influences the evolution of male preferences, which track changes in warning pattern due to sexual selection. When male attraction imposes costs on females, this affects the speed at which novel pattern alleles increase. In two experiments, females laid fewer eggs with males present. However, although males in one experiment showed less interest in females with manipulated patterns, we found no evidence that female coloration mitigates sex-specific costs. Overall, male attraction to conspecific warning patterns may impose an unrecognized cost on *Heliconius* females, but further work is required to determine this experimentally.

## Introduction

If selection can only exploit the best of the immediately available alternative phenotypes, how can novel ecological strategies evolve in already well-adapted organisms? This has traditionally been envisaged as the problem of peak shifts across the metaphorical ‘fitness landscape’ (Wright 1931). When the environment remains stable, in order to move from one adaptive peak (*i.e.* local optimum) to another, populations must first transverse a fitness valley, inhabited by intermediate and typically maladaptive phenotypes. To overcome this problem, genetic drift is often invoked as a means by which populations may avoid these fitness valleys (Wright 1931; Coyne and Orr 2004; Mallet 2010). However, when the traits in question are under positive frequency dependent selection, an additional complication is added: as peaks are defined by the abundance of its corresponding phenotype, new ‘unexplored’ peaks only become available once already populated by a substantial number of (initially maladapted) individuals.

Aposematic warning patterns, which are commonly assumed to be under strong positive frequency dependent selection, can represent considerable fitness peaks in the adaptive landscape (Mallet et al. 1990; Lindstrom et al. 2001; Borer et al. 2010; Merrill et al. 2012; Chouteau et al. 2016; Gordon et al. 2021). Because predators learn to associate particular patterns with unpleasant experiences, an individual’s risk of predation should decrease as the local density of its warning pattern increases (Müller 1879; Sherratt 2008). This can lead to the convergence of warning patterns of different prey species sharing a habitat, a process coined ‘Müllerian mimicry’ (Müller 1879). However, although naively we might expect a single warning pattern to emerge, warning patterns are often very diverse within a community (Briolat et al. 2019).

The establishment of entirely new warning signals under positive frequency-dependent selection via predators is problematic. One possibility is that during periods of relaxed selection, drift may allow new variants to rise above a threshold density until mimicry selection takes over (Mallet and Joron 1999; Sherratt 2006; Mallet 2010). Another possibility is that a model in which predators learn to avoid unpalatable prey only after sampling a fixed number is overly simplistic. For example, if predators are neophobic and generally avoid prey with unfamiliar phenotypes, novel signaling phenotypes might be favored when they are rare (Aubier and Sherratt 2015). A third possibility is that warning patterns might be multifunctional, so that their evolution is not solely governed by purifying frequency dependent processes induced by predators (Briolat et al. 2019).

Sex-specific selection provides a mechanism by which ecological diversity may be promoted (Bonduriansky 2011). Ecological adaptations, including strategies to exploit resources within the environment or avoid predators, are typically – though not always – shared between the sexes. Viability selection is normally expected to push ecological phenotypes towards a shared optimum. Sex-specific selection on the other hand may produce different adaptive optima for the two sexes (Andersson 1994; Arnqvist and Rowe 2005). This can result, for example, from requirements imposed on females to produce offspring or the need for males to find receptive mates (Bateman 1948). The existence of sex-specific optima can also lead to sexually antagonistic selection, and rapid evolution, even in opposition to viability selection (Arnqvist and Rowe 2005). For example, males may evolve strategies that increase their likelihood of securing mates. If these strategies impose costs on females, females may in turn evolve strategies to circumvent these male tactics, leading to further selection on males and so on. If sexually antagonistic selection involves ecologically relevant traits, this might result in peak shifts across the viability fitness landscape.

Since Bates (1862) first described mimicry theory, studies of *Heliconius* butterflies have made a substantial contribution to our understanding of adaptation (Merrill et al. 2015). Distasteful *Heliconius* are well known for their bright warning patterns, which are often associated with Müllerian mimicry. These warning patterns are an important ecological adaptation in *Heliconius*, and predator-induced selection coefficients for the most common local patterns are strong (Mallet et al. 1990). Despite this, *Heliconius* butterflies exhibit a striking diversity of alternative warning patterns (Bates 1862; Merrill et al. 2015). Individual species often vary in warning pattern across their range, leading to distinct geographical color pattern types, and in some cases, such as in *H. cydno, H. numata* and *H. doris*, polymorphisms exist within single geographical populations. In addition, multiple warning patterns frequently coexist within a single geographical community. A given *Heliconius* species may then join many distinct mimicry rings according to local context, leading to the well-documented mosaic of warning patterns observed across the Neotropics (Brown Jr 1976). Spatial variation in local predator and prey communities shape a rugged adaptive landscape, crucial to the maintenance of warning signal diversity.

In addition to warning potential predators, Jocelyn Crane (1955) demonstrated that the bright warning patterns of *Heliconius* stimulate male courtship in the 1950s. Since then, numerous experiments have repeatedly shown that male *Heliconius* generally prefer ‘females’ that share their own warning pattern over that of other conspecific morphs or closely related species (e.g. Jiggins et al. 2001, 2004; Kronforst et al. 2006; Merrill et al. 2011b, 2011a, 2014, 2019; Sánchez et al. 2015; Hausmann et al. 2021). It seems likely that competition between males drives these genetically determined local preferences, as the ability to efficiently locate potential mates within a visually complex environment would be beneficial (Merrill et al. 2019). However, previously mated females may suffer fitness costs if these cues lead to harassment by males during oviposition or foraging. These costs would be augmented by the fact that, although individual *Heliconius* are long lived (up to 6 months), female re-mating is a rare event in most species (Walters et al. 2012). These female-specific costs could conceivably set the stage for interlocus sexual conflict, leading to an arms race between warning pattern and male preferences and rapid evolution when patterns are released from constraints imposed by aposematism (e.g. after a local reduction in predation). Ultimately, this could be an additional factor explaining warning pattern diversification.

To explore how costs imposed on females by male attraction to warning patterns might contribute to pattern diversification in *Heliconius*, we first implemented individual-based simulations. Across a vast parameter space, we tested i) if the presence of such female costs can favor the evolution of novel patterns as opposed to drift acting in isolation, ii) which parameters are most relevant for such dynamics to occur, and iii) how the genetic architecture of male attraction traits might affect the speed at which novel patterns increase in frequency. We then performed experiments to begin to test these ideas by disrupting warning patterns of mated *Heliconius* females with marker pens or by introgressing a novel color pattern allele from a closely related species. We subsequently tested the hypotheses that i) males interact less frequently with females with disrupted patterns, ii) females lay fewer eggs in the presence of males, and that iii) this effect is less pronounced for females with experimentally disrupted patterns.

## Model and Methods

### a) Individual-based simulations

We formalized our verbal model and then implemented this as individual-based simulations in R (R Core Team 2019). By tracking the fate of a novel warning pattern allele, these simulations allowed us to compare how frequently novel patterns might evolve due to drift alone as opposed to when patterns are additionally involved in sexual conflict. This also allowed us to explore which parameters determine how and if sexual conflict contributes to diversification.

#### Arena

Individuals live, breed and die within a rectangular arena of 96 x 4 ‘patches’ with uniform habitat, each representing ~1km^2^ of forest. This arena is divided between 80 ‘central patches’, and the remaining 304 ‘peripheral patches’ (see Fig. 1A). To reduce boundary effects, the arena is wrapped into a torus.

**Figure 1:**
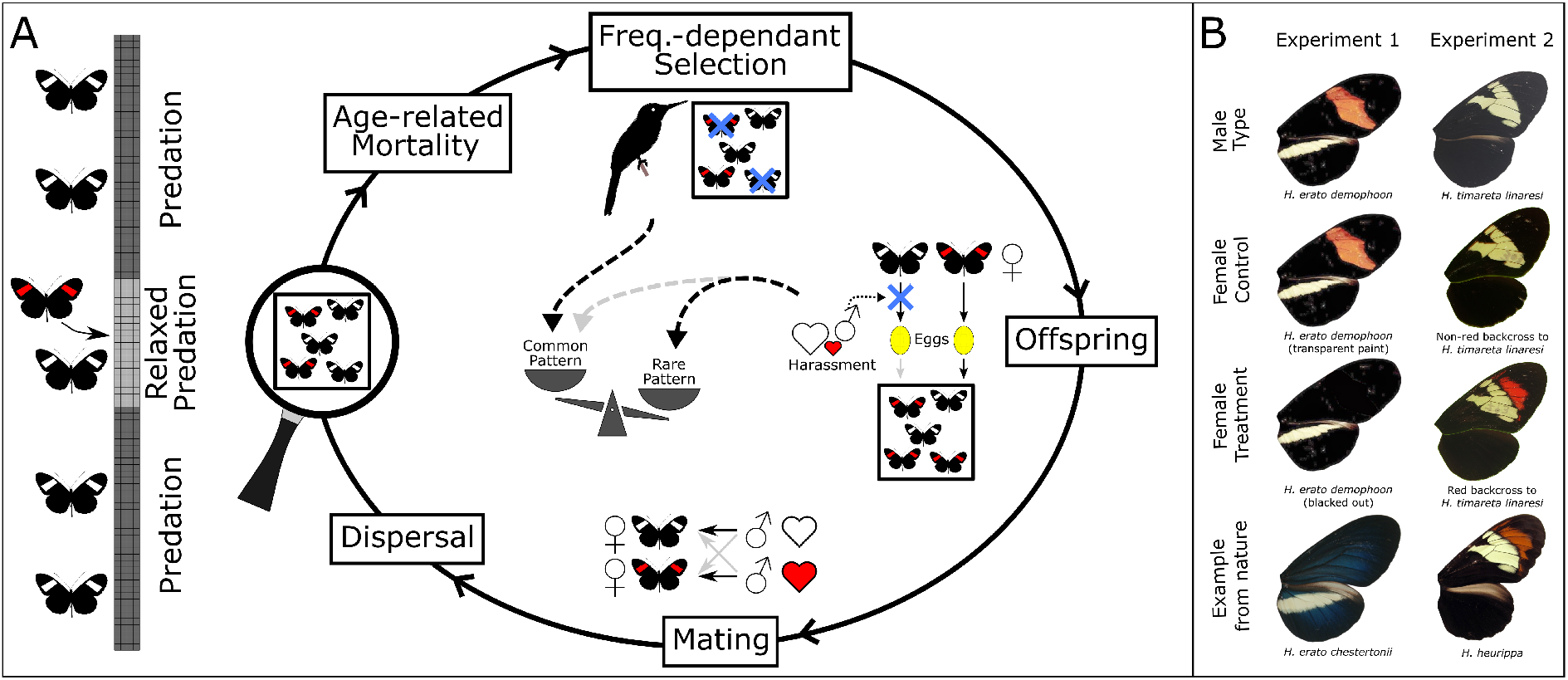
Overview of the study. A) Visualization of the individual-based model. A novel warning pattern allele is introduced into the center of the 96 x 4 patched arena, where predation is initially relaxed. The following generations consist of five phases, shown here for a single patch. Frequency dependent selection by predators always favors the most common pattern, while harassment from males imposes selection against the most common pattern (however, if female adaptions spread faster than male adaptations, this can switch). B) Pattern phenotypes of males and females (both, ‘manipulated’ and control) used in the experiments, as well as examples from nature resembling the ‘manipulated’ patterns.

#### Genetics of individual pattern and preference phenotypes

Individuals are sexual, diploid and have discrete sexes (determined by segregation of ‘sex chromosomes’). Both warning pattern and male mating preferences are genetically determined, and all loci are assumed to be autosomal and segregate independently (Merrill et al. 2015). Many years of research have established that major color pattern elements in *Heliconius* are controlled by just a few Mendelian loci (reviewed in McMillan et al. 2020). Although a handful of genes may differentiate color pattern races, here we are explicitly interested in the spread of individual color pattern alleles and so consider just a single locus. As such, individuals have a single diallelic locus determining variation in a warning pattern element (with alleles *A, a*), which is expressed in both sexes: The derived novel allele *A* is dominant over the ancestral allele *a*, reflecting strong dominance observed at *Heliconius* color pattern loci (McMillan et al. 2020). To account for mating preferences, we assume the existence of two additive quantitative characters *p_a_* and *p_n_*, controlling males’ attraction towards females of the ancestral and novel pattern, respectively. Both traits *p_a_* and *p_n_*, are scaled between 0 and 1 and are each controlled by *N* unlinked diallelic loci with equal effects (Fig. S1). Alleles have dominance relationships so that alleles increasing attraction towards the respective pattern elements are dominant over those that reduce attraction.

#### Life cycle

The life cycle consists of five stages per generation (see Fig. 1A): (1) age-related mortality; (2) frequency dependent selection of adults due to predation; (3) production of offspring; (4) mating; and (5) dispersal. Compared to a previous model from Duenez-Guzman *et al*. (2009), our model incorporates overlapping generations. This increases the biological realism of the model as *Heliconius* are long lived and breed throughout their adult life. Newly eclosed adults are assigned as age = 0 and their age increases by 1 each generation. Individuals with age > 4 are removed from the population at the beginning of each generation.

#### Predation

Predation is modelled implicitly through a patch-specific learning threshold, where predators stop eating adult butterflies of a certain pattern once the learning score *Q* is reached (following Duenez-Guzman et al. 2009). The learning process for each pattern occurs by each butterfly eaten within a patch contributing 1 to the corresponding learning score for the patch. The learning score for each patch is reset every generation and no evolution in predators is allowed. In the 80 ‘central’ patches of the arena, predation is initially relaxed for time *T.*

#### Sexual conflict and reproduction

Offspring can be produced by females either after an encounter with a male or without male encounter. We assume that during oviposition, the chance that a female encounters a male depends on the probability of encountering any particular male (*e*) and the number of males within the patch (*m*):

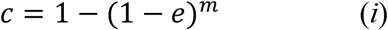

In a scenario of sexual conflict, during an encounter, a female may either be disturbed or not, and hence lay an egg or not. The probability that an encounter between a specific female and male leads to a disturbance by the male depends on the attraction *R* of this male to the female. Following Duenez-Guzman *et al.* (2009), *R* is dependent on the male’s preference trait *p* and a parameter *α*, which is positive and measures the strength of attraction to female patterns. Large values of *α* imply strong attraction to color and smaller values imply weaker attraction. The smaller *α*, the less will *R* differ between males with different preference traits (Fig. S1). *R* for a pair of male *i* with preference *p_x_* and female *j* with pattern type *x* (*x* being either *a* (ancestral) or *n* (novel)) is:

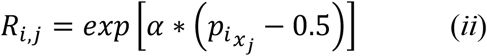

The probability that an encounter leads to a disturbance equals to *R_i,j_* divided by the maximum possible value of *R* (where *p* = 1). The total number of eggs (which directly translates to offspring), *B*, for a patch is Poisson-distributed with mean *b. b* is independent of male disturbance, and limited only by available oviposition sites and female egg laying attempts are repeated until *B* is reached. Gametes producing an egg are randomly hit by mutation. Mutations only occur at loci affecting attraction to patterns, at a constant rate of *μ* per locus per individual per generation (no double-mutations of a locus within an individual during the same generation are allowed). Maternal and paternal alleles recombine freely.

#### Mating

Mating occurs between individuals of the same patch. We assume females only mate once, and continue to lay eggs at a constant rate throughout their lives. Males are assigned to unmated females with relative probabilities equal to their attraction to patterns displayed by females (female choice is not present), which is calculated following equation (*ii*). The majority of females mate immediately after eclosion (and before dispersal); however, because each male can only mate 4 times per generation (mating is costly to *Heliconius* males) females may remain unmated if a patch by chance has >4 times more unmated females than males and then may be mated during subsequent generations. This imposes a very small cost on females with patterns that are less attractive to local males.

#### Dispersal

Finally, newly produced individuals disperse. We assume that each individual migrates to one of the eight neighboring patches or stays in its native patch with the same probability 1/9.

#### Implementation

To efficiently simulate the process of female egg laying and male harassment (by avoiding to individually simulate each egg laying attempt), we made use of the fact that the combined number of eggs placed by females of a patch was independent of male harassment. During each iteration and per patch, *R* for each possible female-male pair was computed and scaled to the maximum possible value of *R* (where *p* = 1). The mean of these values equals to the average probability of male encounters leading to disturbance, *d.* The expected proportion of eggs laid after male encounter, but without disturbance, *P*, is hence:

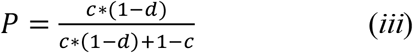

The number of eggs laid after encounter with a male was drawn from a Poisson-distribution with mean *B*P*, and number of eggs without encounter from a Poisson-distribution with mean *B*\*(1-*P*). Mated females within a patch were randomly assigned to the available eggs laid without encounter. However, unattractive females were more likely to contribute to eggs laid with encounter. The relative probability of a female being assigned to those eggs equaled to the inverse of the mean attractiveness of this female to all males in the patch (as calculated following equation (*ii*)).

We ran all simulations with equal starting conditions. The arena was initially populated with 114 individuals (57 mated females and 57 males) per patch, which was a population size similar to the equilibrium population size. Ages were distributed equally over patches and sexes within a patch and ranged from 0 to 5 (as typical for the beginning of a new generation). All individuals were initially homozygous (*aa*) for the recessive ancestral pattern and homozygous (*aa*) for the recessive ancestral allele at loci for attraction to the novel pattern (*i.e.* minimum attraction to the novel pattern in males). Also, all individuals were homozygous (*AA*) for the dominant ancestral allele at loci for attraction to the ancestral pattern (*i.e.* maximum attraction to the ancestral pattern in males). Throughout, the mutation rate *μ* was set at 10^-5.

At the beginning of each subsequent simulation run (*i.e.* at generation 1), a color pattern mutation was introduced into a randomly chosen individual of age 0 within one of the 80 central patches, where predation was initially reduced (so that it was heterozygous *Aa*). The fate of the introduced mutation was then followed until it was lost from the population, or when 2500 generations were reached. During each generation, the number of eggs laid by females within one patch, *B*, was drawn from a Poisson-distribution, with mean *b* = 20. However, for a subset of simulation runs, a bottleneck was imposed within the 80 central-most patches for the first 5 generations to increase the effects of drift, where *b* = 0.5.

#### Parameter space

We systematically varied the predator learning threshold (*Q*); the number of generations that predators were absent from the central patches of the arena (*T*); whether or not a bottleneck was imposed on the central patches in the first generations; the sex of the first mutant; the number of loci affecting each color attraction trait in males (*N*); the strength of color attraction in males (*α*); and whether sexual conflict was present (*i.e.* male harassment during egg laying) or not (reference baseline). In simulation runs where sexual conflict was present, we also varied the probability of encounter between a specific female and male (*e*). Additionally, we always noted the position in the arena where the first mutant occurred. Different settings for the parameters are shown in Table 1, together with the expected relative difference in number of offspring sired by a novel patterned female versus an ancestrally patterned female, as dependent on *α* and *e* (assuming a patch containing males typical for the first generation with minimum attraction to novel and maximum attraction to ancestral pattern). An average of 1000 iterations were run for each of the 720 parameter combinations. 576 of these parameter combinations included sexual conflict and 144 did not. Number of heterozygous and homozygous individuals at each locus in central and non-central patches were recorded during each simulated generation, and stored in a sparse matrix (Bates and Maechler 2019). Scripts to run the model are available in the supplementary R Markdown.

**Table 1:**
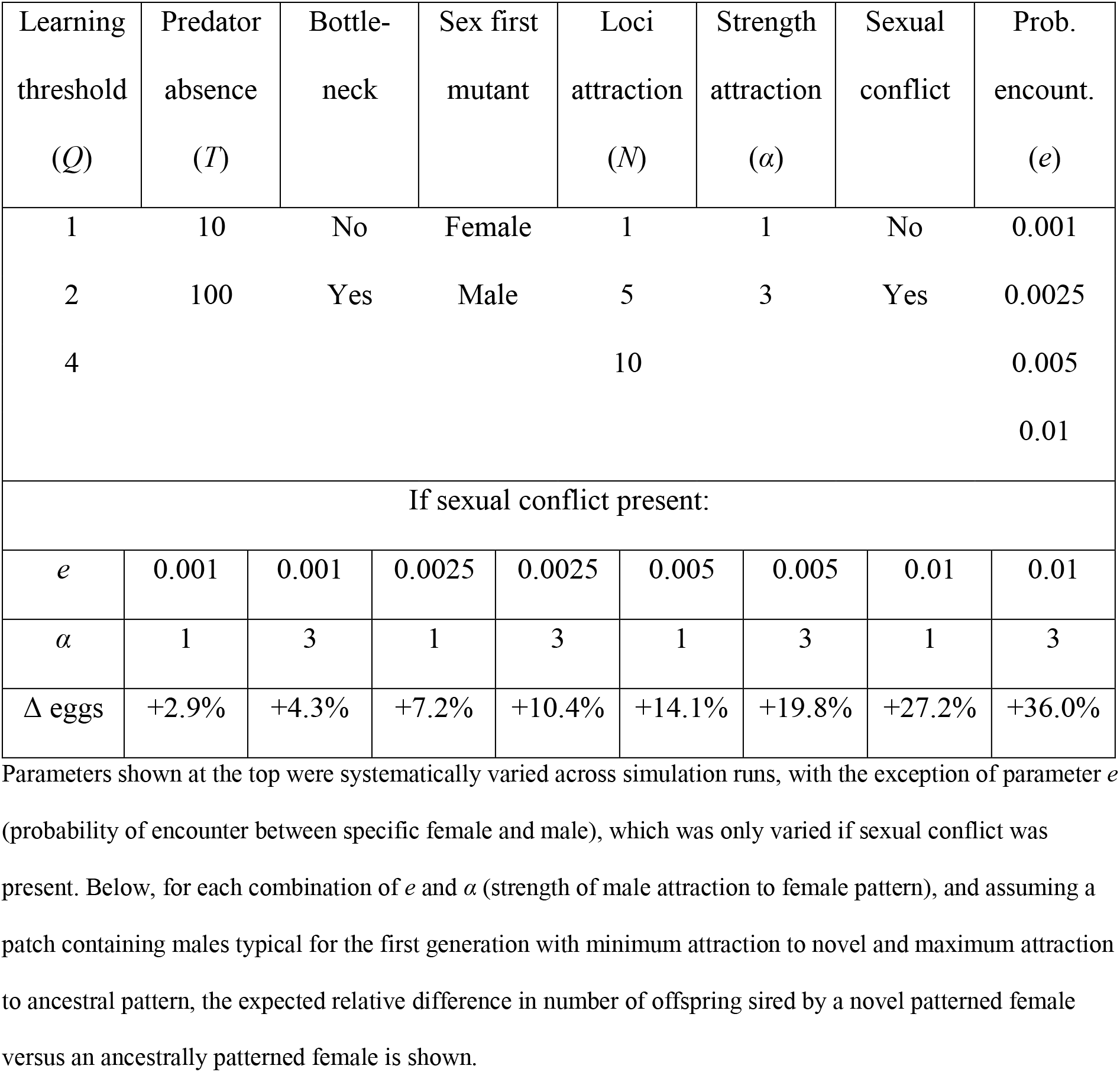
Simulation parameters and their settings.

#### Statistical analysis

All analyses were conducted in *R* (R Core Team 2019). Scripts are available in the supplementary R Markdown. Numeric covariates (*i.e. Q, T N, a, e*; see Table 1 for annotation) were transformed into ordered factors. We tested which of the simulation parameters affects the probability of the novel warning pattern allele surviving until generation 2500 (at which point simulations were always ended) using GLMs with binomial error structure (*i.e.* success = allele remains in population, always identical with going to (near) fixation; failure = allele disappears from population). To avoid complete separation in the model, fixed effects that separated the predictor variable were not included and tested with simple binomial tests (correcting p-values with the Bonferroni method). Stepwise forward-model selection based on AIC via the *MASS* package (Venables and Ripley 2002) was used to find the model that explained the data best, maximally allowing for two-way interactions. To test for the effect of genetic architecture of male attraction traits on number of generations until the novel warning pattern went to near fixation (allele frequency = 0.95; only considering runs where the novel pattern survived), we fitted a GLM with Poisson error structure with number of loci coding for each attraction trait (*N*) as fixed effect. Estimated marginal means for the different predictors were extracted using the *emmeans* package (Lenth 2019). The same package was used to perform Type III Anova to test for significance among fixed effects. Resulting p-values were corrected for multiple testing using the Bonferroni method. R^2^ values were calculated using the *dominanceanalysis* package (Bustos Navarrete and Coutinho Soares 2020).

### (b) Insectary experiments

#### Experiment 1

To investigate female-specific costs associated with male attraction to warning patterns, we first experimentally manipulated female warning patterns with marker pens. This experiment was performed between November 2014 and August 2015 in the Smithsonian Tropical Research Institute insectaries in Gamboa, Panama (9°7’24”N, 79°42’12”W). *Heliconius erato demophoon* (Fig. 1B) were collected around Gamboa and maintained in communal 2×2×2m cages (males and females separately) with ~10% sugar solution, a pollen source, and in the case of females, *Passiflora* host plants. All individuals were numbered on the ventral side of the wings. Females were assigned to a warning pattern treatment (Fig. 1B): either i) disruption of the pattern by painting over the dorsal side of the red forewing band with a black Copic™ Caio 100 marker; or one of two ‘control’ treatments, ii) painting over the dorsal side of the red forewing band with a colorless Copic™ Caio 0 marker (with same solvents as the black marker), or iii) handling but no marker. Visual equivalence of the two control treatments was confirmed by modelling based on the *H. erato* visual system (see Fig. S7 and supplementary methods). A few additional females experienced neither of the three pattern treatments and were excluded from all analyses, except where indicated.

Females were introduced individually into a 2×2×2×m experimental cage, containing a single *Passiflora biflora* host-plant (which was not re-used). Females were left to acclimatize for 48 hours and all eggs laid during this period were removed. This was followed by two 48-hour experimental periods. Three *H. e. demophoon* males were then introduced into the cage during either the first or second experimental period, so that each female experienced 48 hours with, and 48 hours without males. No two females experienced the same combination of males. Eggs were collected daily from the cages. For a subset of the females, we also recorded the time and duration of male-female interactions (hovering ‘courtship’ and chasing). Due to logistic necessities, total observation time differed between females (median = 3607.5, min = 929, max = 7207), largely relating to whether observations were carried out on one or both days in which females were housed with males. We excluded two females from subsequent analysis that did not lay eggs (though their inclusion does not quantitatively affect our results).

#### Experiment 2

To further investigate female-specific costs, but to ensure biologically realistic warning patterns, our second experiment exploited the segregation of Mendelian color pattern elements in hybrids between distinct *Heliconius* populations. This was performed between July and December 2018 in the Universidad del Rosario insectaries in La Vega, Colombia. *H. timareta linaresi* (yellow forewing bar; Fig. 1B) were collected from Guayabal (2°41’04”N, 74°53’17”W) and *H. heurippa* (yellow and red forewing bar) from Lejanías (03°34’0”N, 74°04’20.4”W), Buenavista (4°10’30”N, 73°40’41”W) and Santa María (04°53’28.2”N, 73°15’11.4”W) in Colombia. Outbred stocks were established and used to generate *H. t. linaresi* x *H. heurippa* F1 and backcrosses to *H. t. linaresi* hybrids (as described in Hausmann et al. 2021). The presence of the red forewing band is controlled by a single Mendelian locus (*optix*), and segregates in the backcross to *H. t. linaresi* so that equal numbers of individuals display a (‘novel’) yellow-red forewing band phenotype or a (‘control’) yellow forewing phenotype (Fig. 1B). Sizes of the yellow and red bands was assessed using k-means clustering (see Fig. S8 and supplementary methods). Shortly after eclosion, backcross females were mated to a *H. t. linaresi* male and then housed in a communal cage (2×4×2m) before the experiments. Experimental procedure was as in experiment 1 except that: 1) Insectary-reared individuals were used in all trials; 2) experimental cages contained several species of *Passiflora*, which could not be exchanged between trials; 3) only one male was introduced; and 4) eggs were only collected every 48 hours. As for experiment 1, total observation time differed between females (median = 3547, min = 1722, max = 3904).

#### Statistical analysis

To test for the effect of warning pattern treatment on male interest, we used GLMMs with binomial error structure. The dependent variable was the proportion of seconds with courtship, and pattern treatment was fitted as fixed effect. Female ID was fitted as random effect to correct for individual variation within treatment groups, and control for repeated measures across different days. For experiment 1, the two controls (colorless marker or handled) did not differ in their effect on male interest (*F ratio* = 0.962, *d.f.* = 1, *p* = 0.327), and were therefore combined. For experiment 2, we fitted an additional model to data from yellow-red females, where we explained proportion of seconds with courtship by size of the red band and size of the yellow band of the female (as relative to whole wing area, to correct for female size), and with female ID as a random effect. To test for the effects of male attraction to female patterns on short-term female fecundity, we used GLMMs with Poisson errors. Here the dependent variable was the number of eggs laid over a 48h period. Male presence, female pattern and their interaction were fitted as fixed effects, and female ID was fitted as random effect. Our prediction was that any differences in short term female fecundity resulting from male attraction to specific warning patterns would be observed as a significant interaction between male presence and pattern treatment. Again, for experiment 1, the two controls (colorless marker or handled) did not differ in their effect on female fecundity (*F ratio* = 1.071, *d.f.* = 1, *p* = 0.301), nor was there a significant interaction with male presence (*F ratio* = 0.993, *d.f.* = 1, *p* = 0.319), and they were therefore combined. To test for the effect of male presence on female fecundity in isolation, we fitted the same type of model structure, but only including male presence as fixed effect. For experiment 1, we additionally included females that were neither treated with a marker, nor handled (and which were excluded from all other analyses). We used the *emmeans* package (Lenth 2019) to determine the effect of the different variables (via type III Anova), and to calculate estimated marginal means (EMMs) and effect sizes (*i.e.* difference in eggs laid between experimental periods).

## Results

### (a) Simulation results

#### Sexual conflict facilitates the evolution of novel warning patterns

We ran simulations across 720 different parameter combinations (each replicated ~1000 times). The novel pattern quickly disappeared from the population in all 144000 simulation runs where only genetic drift could contribute to the diversification of warning patterns (Fig. 2A). The novel pattern allele only survived in simulations which included both, sexual conflict *and* the maximum period of relaxed predation (*i.e. T* = 100 generations; Fig. 2A). The novel allele persisted in 2865 of 288000 simulations fulfilling these two criteria (*i.e.* 1%), and in each of these it also increased in frequency to near or complete fixation (Fig. S4). By far the most important additional parameter in these simulations was the probability of a male-female encounter, which explained 64% of the variance (Fig. 2B): with the notable exception of 2 simulation runs, where a value of 0.005 was sufficient, this parameter had to be at 0.01 for the novel pattern allele to be retained. However, stepwise model selection also revealed that strength of male attraction to color (*α*), predator learning threshold (*Q*), presence of a bottleneck in the central patches during the first generations (‘bottleneck’), sex of the first mutant (‘sex’), as well as the interactions bottleneck\**α*, bottleneck\**Q*, bottleneck*sex, and *α*\**Q*, also all significantly affected the retention of the novel allele (Fig. 2B and Table S2). Overall, the most promising combination of parameter settings included sexual conflict, *Q* = 1, *T* = 100, no bottleneck, female first mutant, *N* = 10, *α* = 3 and *e* = 0.01, where the novel allele was retained in 26% of runs (Table S1). We also found that the novel allele was more likely to survive when the initial mutation occurred in one of the most central patches (*i.e.* far away from active predation, Fig. S2). Indeed, the eventual fate of the novel pattern allele seemed to be largely determined during these first generations of relaxed predation in the central patches, as its frequency at generation 100 was closely correlated with future retention or loss of the allele (Fig. S3).

**Figure 2:**
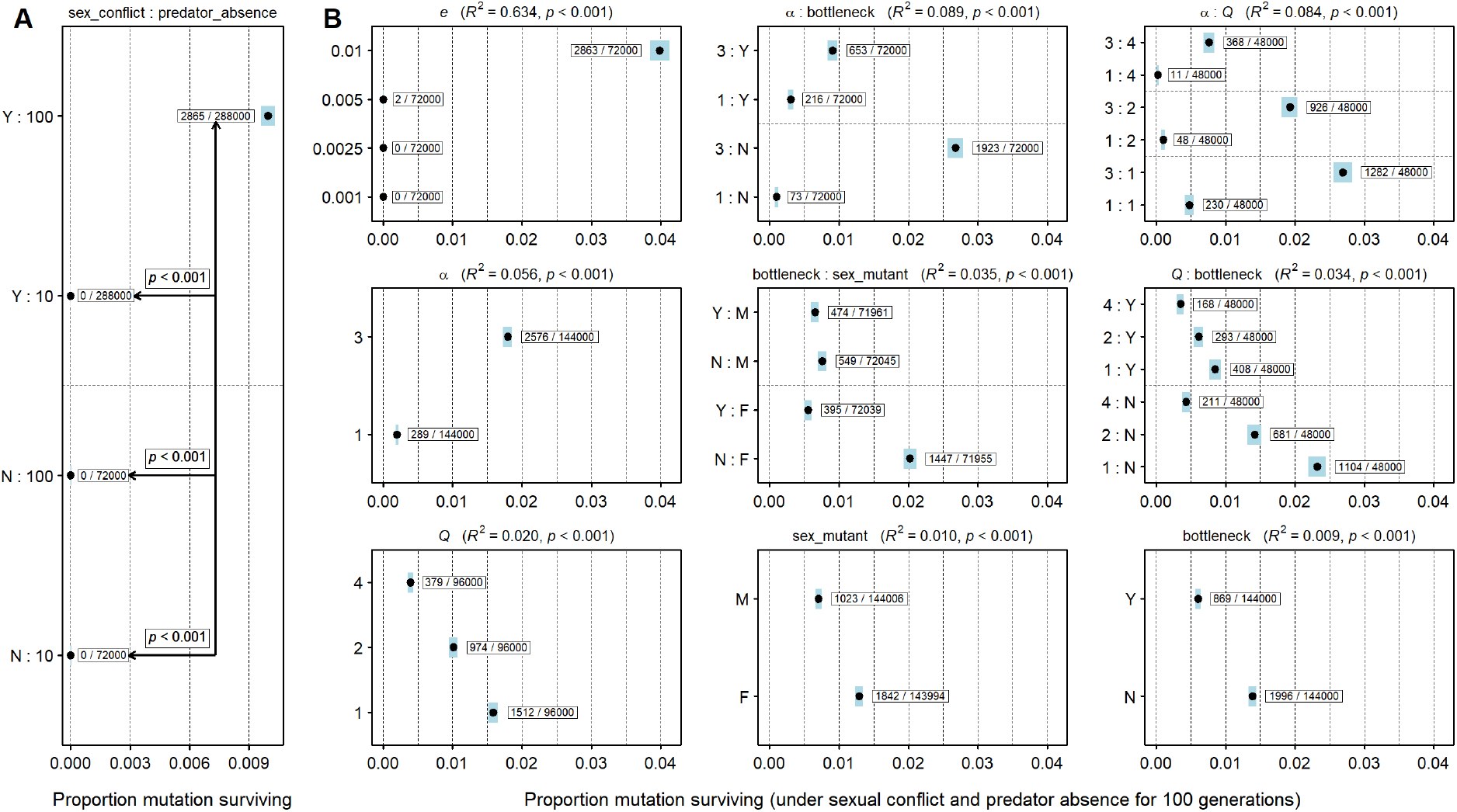
Presence of sexual conflict and duration of relaxed predation are the strongest predictors of retention of novel pattern, followed by probability of male encounter. A: Effect of sexual conflict (‘Y’ = yes, ‘N’ = no) and duration of predator absence from central patches on probability of novel warning pattern retention. Proportions and 95% binomial confidence intervals are displayed for each combination, together with p-values from binomial tests. B: Effects of simulation parameters retained during model selection (same parameter annotation as in Table 1). R^2^ value and p-value are shown above each panel. Interactions are indicated with a ‘:’ sign above the respective panel, and combinations are separated by a ‘:’ sign at the y-axis (‘Y’ = yes, ‘N’ = no, ‘M’ = male, ‘F’ = female). Estimated marginal means (circles) and their 95% confidence intervals (blue rectangles) are shown for each predictor.

#### Male attraction to colors tracks warning pattern evolution

Overall, the mean frequency of alleles causing male attraction to the novel pattern closely tracked the frequency of the novel pattern allele, but lagged behind some generations, as typically expected during chase-away selection (Fig. 3A, see also supplementary animation). The magnitude of this lag depended on the genetic architecture of male attraction traits, with more complex architectures (*i.e.* more preference loci) generating a greater discrepancy between pattern and male attraction to this pattern (Fig. S4). In contrast, the mean frequency of alleles determining attraction to the ancestral pattern changed very little (Fig. 3A; note that male attraction to the two pattern types was controlled by independent loci): Across simulation runs, the average male was at no point more attracted to the novel than to the ancestral pattern (Fig. S5). Therefore, once the novel pattern exceeded the frequency of the ancestral pattern, both mimicry selection and selection imposed by male harassment acted in the same direction for a period of time, both favoring for the novel pattern (Fig. S5).

**Figure 3:**
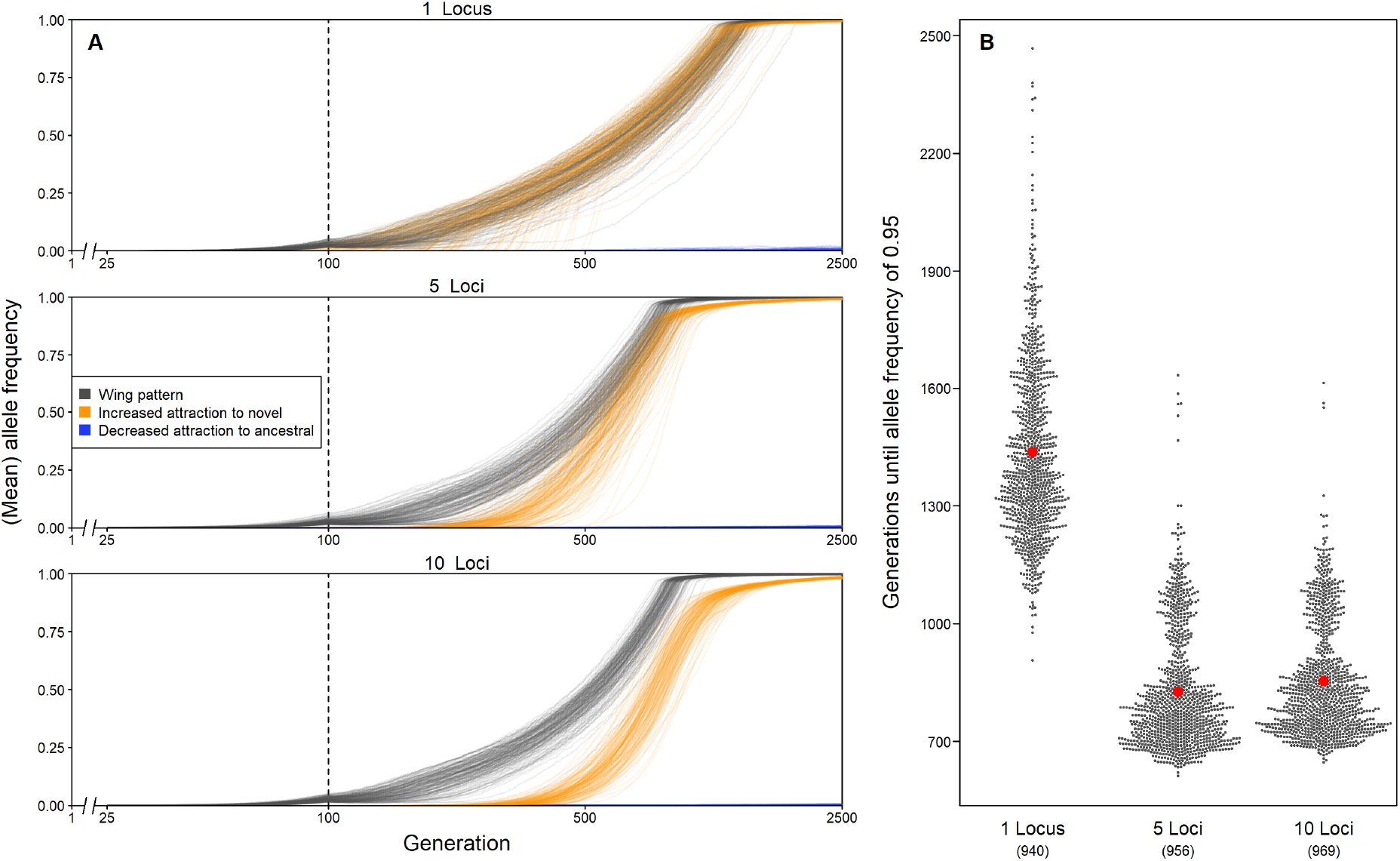
Male attraction tracks changes in warning pattern and its genetic architecture affects the speed at which novel patterns spread. A: Allele frequencies over generation time for novel alleles at the warning pattern locus (gray), loci for male attraction to the novel pattern (orange; novel alleles increase attraction) and loci for male attraction to the ancestral pattern (blue; novel alleles reduce attraction). Means were calculated whenever multiple loci were involved. Panels are split by different genetic architectures of male attraction traits (1, 5 or 10 loci). Results from only one evolutionary scenario are shown (sexual conflict present; *Q* = 2; *T* = 100; no bottleneck; sex first mutant = female; *α* = 3; *e* = 0.01) and only including those where the novel pattern persisted. x-axis is log-scaled, and frequencies at generations < 25 are not displayed (as they are invisibly small). Vertical line at 100 generations indicates end of relaxed predation. B: Speed at which the novel pattern allele reaches a frequency of 0.95, as dependent on number of loci for male attraction traits (including data across all other simulation parameters). Estimated marginal means are shown in red (CIs are invisibly narrow).

#### Genetic architecture of male attraction influences the speed at which novel pattern alleles spread in simulations with sexual conflict

Although the number of loci encoding attraction phenotypes was not retained in our model testing for survival of the novel pattern allele, it did have a strong effect on how fast the novel pattern allele’s frequency increased in the population, in cases where it persisted (Fig. 3B and S6). With just one locus determining attraction to each pattern, simulation runs required on average ~600 generations longer to reach a frequency of 0.95 of the novel pattern allele compared to simulation runs with 5 loci (z-test: *z* = 394.812, *p* < 0.001) or 10 loci (z-test: *z* = 375.969, *p* < 0.001). Surprisingly, simulation runs with 5 or with 10 loci did not differ much from each other in this respect (and the trend was even slightly reversed, z-test: *z* = −20.558, *p* < 0.001). These differences were observed across all possible combinations of the other simulation parameters (see supplementary R Markdown).

### (b) Empirical results

#### Warning patterns influence male harassment of mated *H. erato* females

In experiment 1, we carried out observations for 58 individual females, including 30 experimental butterflies with disrupted warning patterns, and 28 butterflies subjected to one of the two control treatments (14 of each). After removing outliers (five experimental and four control individuals, see Fig. S9 and supplementary methods), females with intact warning patterns were harassed more often than those with disrupted patterns (Fig. 4A, *F ratio* = 13.34, *d.f.* = 1,*p* < 0.001; inclusion of the outliers did not qualitatively affect the results: Fig. S9A, *F ratio* = 6.24, *d.f.* = 1, *p* = 0.013). In experiment 2, we carried out observations for 45 individual females, including 20 ‘experimental’ butterflies with yellow-red forewing pattern, and 25 ‘control’ butterflies with yellow forewing pattern. Overall, males were much more responsive than in experiment 1. Surprisingly, including all individuals, females with ‘experimental’ pattern were harassed more often than those with ‘control’ pattern (Fig. S9B, *F ratio* = 4.54, *d.f.* = 1, *p* = 0.033), though this trend was no longer significant when outliers (two ‘experimental’ and one ‘control’ individual) were removed (Fig. 4C, *F ratio* = 3.61, *d.f.* = 1, *p* = 0.058). However, for yellow-red females with measurements of the band sizes available (12 of the 18 from Fig. 4C), we found that male harassment decreased as the size of the red band increased (Fig. 10A, z-test for slope ≠ 0: z = 2.566, p = 0.010), whereas the size of the yellow band had no effect (Fig. 10B, z-test: z = 0.116, p = 0.908).

**Figure 4:**
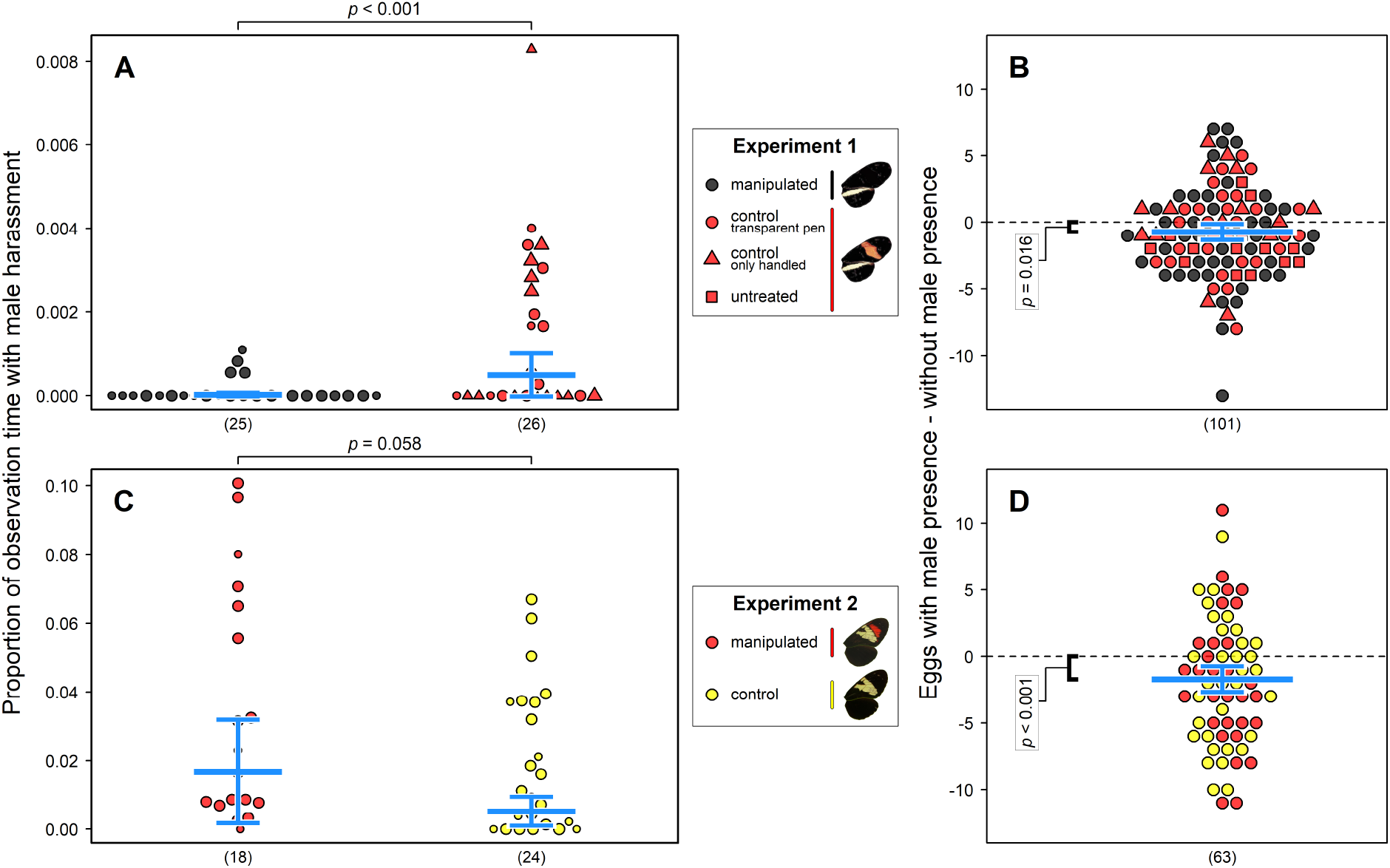
Males harass mated females with intact warning patterns more often in experiment 1. Females lay fewer eggs in the presence of males in both experiments. Panels A+B correspond to experiment 1 (*H. erato*), panels C+D to experiment 2 (*H. timareta*). Warning pattern treatment is indicated by dot color and shape. A+C: Proportion of total observation time males spent harassing females with different patterns. Area of dots is relative to total number of observed seconds. Estimated marginal means and their confidence intervals (CIs) are shown in blue. B+D: Difference between eggs laid with and without male presence. Values below 0 indicate that male presence reduces fecundity. Effect sizes and their confidence intervals (CIs) are shown in blue.

#### Male presence reduces the number of eggs laid by females

There was considerable variation in the number of eggs laid, both, between experimental periods (*i.e.* males present and males absent) for individual females, as well as between females (Fig. 4B+D). Nonetheless, in both experiments, females laid fewer eggs in the presence of males (experiment 1, including females without warning pattern treatment: *F ratio* = 5.83, *d.f.* = 1, *p* = 0.016 (Fig. 4B); experiment 2: *F ratio* = 12.49, *d.f.* = 1, *p* < 0.001 (Fig. 4D)). Over a two-day period, females in the presence of males laid ~0.73 [CI: 0.14-1.32] fewer eggs in experiment 1 (a reduction of 13%; GLMM estimate without male = 5.45, with male = 4.72) and 1.70 [CI: 0.74-2.66] fewer eggs in experiment 2 (a reduction of 19%; GLMM estimate without male = 8.86, with male = 7.15). However, we found no evidence of an interaction between the presence of males and warning pattern treatment on the number of eggs laid (experiment 1: *F ratio* = 1.110, *d.f.* = 1, *p* = 0.291; experiment 2: *F ratio* = 0.005, *d.f.* = 1, *p* = 0.943). Hatching success, measured for a subset of females in experiment 2, was also unaffected by male presence (Fig. S11).

## Discussion

The warning patterns of *Heliconius* butterflies have become a textbook example of natural selection (e.g. Barton *et al.*, 2007; Futuyama & Kirkpatrick, 2017), but the origins of their considerable diversity remain problematic. We explored whether sex-specific selection might contribute to the evolution of novel warning patterns. Using individual-based simulations, we have shown that drift alone is unlikely to account for the spread of novel warning pattern alleles. Instead our simulations suggest that harassment of previously mated females by males attracted to their bright warning patterns – in association with periods of initially relaxed predation – could facilitate the spread of novel pattern alleles. We also showed that genetic architecture of male attraction traits can influence how quickly a novel pattern may spread through the population, as it determines how fast males adapt their corresponding mating preferences. Data from insectary experiments also provide some support that sexual conflict can facilitate the spread of novel patterns: The presence of males reduced short-term female fecundity, highlighting that unwanted male attention may well favor the evolution of female ‘defensive’ traits in *Heliconius.* However, we found no evidence in our experiments that novel warning patterns mitigate female-specific costs.

In our simulations, we considered a large parameter space to investigate which variables may facilitate the spread of novel warning pattern alleles. Notably, in all 144000 simulations without sexual conflict, the novel allele was rapidly lost from the population, regardless of other parameter values. In contrast, among the remaining 576000 simulations where sexual conflict was present, the novel allele was retained in ~0.5% of runs. Despite the striking diversity of warning patterns in *Heliconius*, the spread of a novel pattern allele is presumably a rare event, so even this apparently modest increase could make a substantial difference across 12 million years (Kozak et al. 2015) of *Heliconius* diversification. Overall, our simulations suggest that drift acting in isolation may be unlikely to facilitate the evolution of novel warning patterns, and that sex-specific selection may be an important, previously unrecognized factor.

Alongside a potential role for sex-specific selection, our simulations also suggest that increased periods of relaxed predation and high probabilities of encounter between mated females and males are important factors determining the fate of novel pattern alleles. Periods of relaxed predation have frequently been invoked as a necessary prerequisite for the spread of novel warning patterns in aposematic butterflies (Mallet and Joron 1999; Mallet 2010). High population density (leading to increased probabilities of encounter) has also been shown theoretically to drive diversification through interlocus sexual conflict (Gavrilets 2000). Notably, in 2 of our simulations runs, the novel pattern persisted with a moderate encounter probability between females and males. This suggests that even at lower encounter rates, sexual conflict can contribute to the evolution of novel warnings patterns. Of course, the interpretation of our simulations must depend on how likely its parameters reflect reality. The benefits that we introduced to females with novel patterns in our simulations (during relaxed predation; see Table 1) are broadly comparable to the higher fecundity experienced by females in the absence of males in our empirical data. However, although a great deal is now known of *Heliconius* biology (Merrill et al. 2015; Jiggins 2017), making them an excellent subject for exploration with individual based simulations, it is important to note that we still know little about their predators, both in terms of population densities or learning functions (Jiggins 2017). Similarly, although *Heliconius* can exist in high densities, this – to our knowledge – has not been systematically assessed.

In evolutionary arms-races, adaptations in one party are contested by counter-adaptations in the other (Arnqvist and Rowe 2005). The genetic architecture underlying these adaptations can play a crucial role during these dynamics, as it may affect how fast one party can distance itself, or conversely catch up, to the other party. Our simulations show that male preferences rapidly track changes in female wing pattern cue, and that very simple genetic architectures speed up this process. This in turn decreases the speed at which the novel pattern allele spreads across the population, as female benefits from novel patterns are reduced when male interest in this pattern increases. However, this does not affect whether or not the novel pattern eventually reaches high frequencies in our simulations. The final fate of the novel pattern allele in our simulations seems to be largely determined in the first generations after its occurrence, during which the novel pattern is at very low frequency in the population, and hence, there is not yet strong selection for increased attraction to this pattern.

We modelled attraction of males to novel or ancestral patterns with two independent traits, controlled by different sets of loci (following Duenez-Guzman et al. 2009). Alleles that reduce attraction to the ancestral pattern did not markedly increase in frequency in any of our simulations. Males in our simulations suffered few costs from being attracted to females. This seems reasonable considering the sparsity of receptive *Heliconius* females in the wild, securing access to which possibly outweighs costs associated with attraction to the ‘wrong’ female (Estrada and Jiggins 2008). In our simulations, this meant that a reversal in harassment probability of differently colored females never occurred (*i.e.* the novel pattern was never harassed more than the ancestral pattern). If both attraction traits were controlled by the same loci (*i.e.* an increase in attraction to the novel pattern requires a decrease in attraction to the ancestral pattern), this might lead to such reversals and possibly increase the probability for color polymorphisms to occur. This would be an interesting question for further empirical and theoretical work, but is beyond the scope of the current study.

Both our experiments support the hypothesis that male presence is costly for mated females, at least in the short term, as shown by a reduction of eggs laid. Although we cannot rule out the possibility that competition between females and males over food resources, as opposed to male harassment, accounts for this reduction in laying rate, we consider this unlikely. In our experiments, butterflies were provided with multiple flowers and feeders (similar to cages housing much larger number of butterflies, where average laying rates per butterfly are within the same range), and it seems unlikely that this was a limiting resource. Although it has not previously been shown experimentally, *Heliconius* researchers frequently keep mated females separate from males to increase egg yield (independent of overall butterfly density). We therefore consider it more likely that male harassment interrupting females during egg laying, or during foraging or scouting for host plants, explains our data. In support of this view, the reduction in the eggs laid was more pronounced in experiment 2, in which we also observed much higher levels of male-female interactions (despite having only a single male with each female).

*H. erato* and *H. timareta* females respectively laid on average 13% and 19% fewer eggs when males were present. Although this may seem a relatively small reduction, female *Heliconius* only lay a few eggs per day, and if consistent across a female’s reproductive life (up to several months), this would represent a significant reduction in fitness. Notably, this is comparable to per locus estimates of selection acting on warning pattern due to predation (albeit at the lower end; e.g. per locus s = 0.13–0.40 in *H. erato* and *H. melpomene*, Mallet et al. 1990). Once again, extrapolation of our results to natural populations must be treated with some caution. In particular, the activity and density of individuals in our insectary enclosures might not reflect the situation in the wild. For example, it wild *Heliconius* males often show much higher activity than captive individuals (potentially leading to high rates of encounter between females and males). Local abundance of *Heliconius* in the wild *can* be very high, and interactions frequent, but there is considerable variation in density between sites (Merrill, pers. obs.). As such, it is unclear whether the densities in our insectary experiments reflect those in natural populations and this remains an important caveat. Nevertheless, given the large effective population sizes of many *Heliconius* species, even a relatively small fitness cost resulting from male harassment could have significant effects on the evolution of associated traits (like warning pattern).

Although our experiments reveal that male presence can reduce short-term female fecundity, they provide only limited evidence for a key prediction of our hypothesis: That novel warning patterns should mitigate costs resulting from male harassment. *H. erato* females with disrupted patterns did receive less attention from males; however, there was no effect of pattern treatment on short term female fecundity (*i.e.* number of eggs laid with males present). In our experiments with *H. timareta*, we found no evidence that novel patterns reduce male harassment (indeed, there is some evidence that males are more interested in females with the red band) or that warning pattern affects the number of eggs laid with males present.

Despite this, it is *perhaps* premature to rule out a role of sexual conflict as a factor contributing to the evolution of novel warning patterns. The increased interest of males directed towards females with intact patterns observed in our first experiment, mirrors previous experiments testing male attraction to population-specific warning patterns in *H. erato* (e.g. Muñoz et al. 2010; Finkbeiner et al. 2014; Merrill et al. 2014). Indeed, our warning pattern manipulation of *H. erato demophoon* creates a similar phenotype to *H. e. chestertonii* (see Fig. 1B), which Muñoz *et al*. (2010) have shown to be less attractive to neighboring, red-banded populations of *H. erato*. Why the differences in interactions we observed do not translate into differences in female short-term fecundity is not immediately clear. One possibility is that anti-aphrodisiac pheromones, transferred from males to females during mating (Gilbert 1976), which have also been hypothesized to reduce male harassment (Estrada et al. 2011), may mitigate any detectable costs associated with initial male interest. Another possibility is that our experiments simply lack power, especially considering the large variation in the number of eggs laid.

The results of our experiments with *H. timareta* may simply reflect the lack of strong differences in visual attraction. In particular, mate choice experiments with *H. t. linaresi* males – which were run concurrently with experiments reported here – revealed that males show only very weak preferences for the conspecific yellow pattern over a yellow-red pattern, as used in the present study (Hausmann et al. 2021). Recently, it has also become apparent that the composite color patterns of *H. heurippa* may not be the target of differences in male attraction between different species of *Heliconius* (Mavárez et al. 2021). We initially chose to study these taxa because introgressing a red band into *H. t. linaresi* recreates a *H. heurippa*-like pattern (see Fig. 1B), and is thought to reflect the evolutionary history of this putative hybrid species. We consider the initial acquisition of color pattern alleles through hybridization and introgression an especially likely scenario affecting the kinds of dynamics we describe here. A stronger effect of warning pattern in mitigating fecundity loss due to male presence may be observed if repeating our experiments with other *Heliconius* species that are known to show stronger color based mate choice, such as *H. melpomene* (Merrill et al. 2019). Further experiments across a broader range of populations would be important to robustly test a role for sexual conflict in driving warning pattern divergence in *Heliconius*.

The rise of a new variant at an ecologically relevant locus due to interlocus sexual conflict is a challenging concept, especially if it is at the same time constrained by positive frequency dependent natural selection (e.g. Sherratt 2008; Briolat et al. 2019). As the new variant will be less advantageous for one sex than the other (in our example, males with novel warning pattern suffer higher costs than females), an additional component of (intralocus) sexual conflict is introduced, where male and female adaptations at the same locus are in conflict (Schenkel et al. 2018). Such combined effects can hinder, but possibly also induce antagonistic coevolution between males and females (Pennell and Morrow 2013; Pennell et al. 2016). A role for sex-specific selection driving divergence in primarily ecological traits has been suggested previously (Bonduriansky 2011). In poison frogs, it seems that sexual selection due to female preferences for bright colors has contributed to inter-population differences in warning signals (Maan and Cummings 2009). Similarly, evidence suggests that male harassment drives phenotypic diversity in wing color in damselflies (Svensson et al. 2005). Although there seems to be little evidence that these damselfly color morphs are ‘ecologically relevant’ (*i.e.* affecting an individual’s survival due to interaction with heterospecific individuals, or the abiotic environment), diversity appears to enhance population performance more generally by reducing overall fitness costs to females from sexual conflict (Takahashi et al. 2014). Among *Papilio* butterflies, which are often female-limited Batesian mimics, non-mimetic ‘male-like’ females might exist to avoid unwanted attention of males. In *P. dardanus*, males do indeed prefer to approach mimetic over non-mimetic (male-like) females (Cook et al. 1994). Our study contributes to this body of work by explicitly testing for fitness effects resulting from sexual conflict relating to an ecological trait.

In conclusion, our theoretical results show that drift alone might be unlikely to drive diversification of warning patterns. However, once these patterns are additionally involved in mate choice, and the two sexes have different reproductive strategies (e.g. males are adapted to mate as often as they can, while females are not), sexual conflict can arise and contribute to warning pattern diversification. The speed at which novel patterns increase in frequency will then depend on the genetic architecture underlying adaptations in the sexes arising from this arms-race. Our empirical results show that females can indeed suffer fitness costs from unwanted male attention, but this was not mitigated by females displaying unusual patterns. The failure of detecting such interaction could be caused by a number of factors, the most likely of which is that color based preferences of males in the taxa we used are not strong enough to see a clear effect. Future work may repeat these experiments with taxa that are known to show stronger color based mate choice.

## Supporting information

Analysis scripts markdown

Empirical data for erato experiments

Wing JND values

Model parameter settings

Summary simulation data

Supplementary methods and results

Empirical data for timereta experiments

## Data accessibility

Supplementary methods and results, data and analysis scripts (in form of an R Markdown document) are included as electronic supplementary material.

## Acknowledgements

We are grateful to the Ministerio del Ambiente and the Autoridad Nacional de Licencias Ambientales (ANLA, permit 530 awarded to the Universidad del Rosario) for permission to collect butterflies in Panama and Colombia, respectively. We are very grateful to the Abondano-Almeida family for being a great support to AEH and MF in Colombia; Juan Sebastián Sánchez, Óscar Penagos, Isabel Leon and Rachel Crisp for assistance in the insectaries. RMM is indebted to Chris Jiggins for valuable discussions. SN was funded by a British Ecological Society Small Research Grant awarded to RMM, and RMM was additionally supported by a Junior Research Fellowship from King’s College Cambridge. AEH, MF and C-YK were funded by an Emmy Noether fellowship and research grant awarded to RMM by the Deutsche Forschungsgemeinschaft (DFG) (Grant Number: GZ: ME 4845/1-1).

## Author contributions

RMM, AEH and MF designed the research. AEH and RMM wrote the simulation model. AEH, MF, SN and RMM collected the behavioral data. AEH and MF analyzed the behavioral data and C-YK analyzed the wing spectral data. RMM, ML, WOM, CP-D, and CS and contributed experimental animals and facilities in addition to academic supervision. AEH, MF and RMM wrote the paper with contributions from all authors.

