## Supplementary material for "Does sexual conflict contribute to the evolution of novel warning patterns?": Analysis scripts markdown

Anonymous (for double-blind review)

last edited: 17 June 2021

**Authors of manuscript**: Anonymous (for double-blind review)

**Published in**: Under review (2021)

**DOI**:

#### 1 Initial explantions and setup

Load required libraries

```
# For binomial tests
suppressWarnings(suppressMessages(library(binom)))
# For retrieving estimated marginal
# means (EMMs) from models
suppressWarnings(suppressMessages(library(emmeans)))
# For stepwise AIC
suppressWarnings(suppressMessages(library(MASS)))
# For dominance analyses (R2
# calculation)
suppressWarnings(suppressMessages(library(dominanceanalysis)))
# For reading png images
suppressWarnings(suppressMessages(library(png)))
# For displaying png images
suppressWarnings(suppressMessages(library(grid)))
# For mixed-models
suppressWarnings(suppressMessages(library(lme4)))
# For beeswarm plots
suppressWarnings(suppressMessages(library(beeswarm)))
```

Call `plot_proportion_stats.R`, which contains a function that will later be used for producing the locally jittered stripchart graphs of the male attraction data.

```
github <- "https://raw.githubusercontent.com"
repos <- "/SpeciationBehaviour/visual_preference_heurippa_linaresi"
folder <- "/master/Analyses_and_Data/Skripts_called_from_Markdown"
source(paste0(github, repos, folder,
    "/Plot_Proportion_Stats.R"))
```

With this function, we calculate the selection coefficient for the novel pattern mutation in females:

```
calc_s <- function(number_indiv, chance_encounter,
    p1, p2, alpha) {
```

```
    # Probability of encountering a male
    ccc <- 1 - (1 - chance_encounter)^number_indiv
    # mean attraction of males to
    # females
    ddd <- mean(c(exp(alpha * (p1 -
        0.5))/exp(alpha * (0.5)), rep(exp(alpha *
        (p2 - 0.5))/exp(alpha * (0.5)),
        number_indiv)))
    # Proportion of eggs laid after
    # encounter
    propo <- (ccc * (1 - ddd))/(ccc *
        (1 - ddd) + (1 - ccc))
    # Expected fraction of eggs coming
    # from each female type
    eggs1 <- propo + (1/number_indiv) *
        (1 - propo)
    eggs2 <- (1/number_indiv) * (1 -
        propo)
    # Double check if it equals 1
    eggs1 + (number_indiv - 1) * eggs2
    # Maximum fitness
    max_fit <- max(c(eggs1, eggs2))
    # relative fitnesses
    w1 <- eggs1/max_fit
    w2 <- eggs2/max_fit
    # Selection coefficient
    s <- 1 - w2

    return(s)
}
```

This function will be used later to draw the allele frequency lines, or those for phenotype ratios:

```
add_lines <- function(all_mat, x, n_mat,
    cols) {
    for (mato in 1:n_mat) {
        lines((2500 - length(all_mat[[mato]][,
            x]) + 1):2500, all_mat[[mato]][,
            x], col = adjustcolor(cols[mato],
            1/10))
    }
}
```

We define a function to later add estimators and CIs to the plots.

```
add_CI <- function(EMM_obj, at = 1:2,
    match_col = "Male_Type", match_vec,
    response = "prob") {
    if (match_col == "") {
        segments(at - 0.65, EMM_obj[,
            response], at - 0.35, EMM_obj[,
            response], col = "white",
            lwd = 5.5, lend = 2)
        arrows(at - 0.5, EMM_obj$asymp.LCL,
            at - 0.5, EMM_obj$asymp.UCL,
```

```
            col = "white", lwd = 4.5,
            length = 0.15, code = 3,
            angle = 90, lend = 2)
        segments(at - 0.65, EMM_obj[,
            response], at - 0.35, EMM_obj[,
            response], col = "dodgerblue",
            lwd = 4, lend = 1)
        arrows(at - 0.5, EMM_obj$asymp.LCL,
            at - 0.5, EMM_obj$asymp.UCL,
            col = "dodgerblue", lwd = 3,
            length = 0.15, code = 3,
            angle = 90, lend = 1)
    } else {
        segments(at - 0.65, EMM_obj[,
            response][match(match_vec,
            EMM_obj[, match_col])],
            at - 0.35, EMM_obj[, response][match(match_vec,
                EMM_obj[, match_col])],
            col = "white", lwd = 5.5,
            lend = 2)
        arrows(at - 0.5, EMM_obj$asymp.LCL[match(match_vec,
            EMM_obj[, match_col])],
            at - 0.5, EMM_obj$asymp.UCL[match(match_vec,
                EMM_obj[, match_col])],
            col = "white", lwd = 4.5,
            length = 0.15, code = 3,
            angle = 90, lend = 2)
        segments(at - 0.65, EMM_obj[,
            response][match(match_vec,
            EMM_obj[, match_col])],
            at - 0.35, EMM_obj[, response][match(match_vec,
                EMM_obj[, match_col])],
            col = "dodgerblue", lwd = 4,
            lend = 1)
        arrows(at - 0.5, EMM_obj$asymp.LCL[match(match_vec,
            EMM_obj[, match_col])],
            at - 0.5, EMM_obj$asymp.UCL[match(match_vec,
                EMM_obj[, match_col])],
            col = "dodgerblue", lwd = 3,
            length = 0.15, code = 3,
            angle = 90, lend = 1)
    }
}
```

Super neat function to add a picture to a plot. Taken from here: https://stackoverflow.com/questions/27800307/adding-a-picture-to-plot-in-r

```
addImg <- function(
  obj, # an image file imported as an array (e.g. png::readPNG, jpeg::readJPEG)
  x = NULL, # mid x coordinate for image
  y = NULL, # mid y coordinate for image
  width = NULL, # width of image (in x coordinate units)
  interpolate = TRUE # Apply linear interpolation?
){
```

```
  if(is.null(x) | is.null(y) | is.null(width)){
    stop("Must provide args 'x', 'y', and 'width'")
    }
  # A vector of the form c(x1, x2, y1, y2) giving the extremes of the user coordinates
  # of the plotting region
  USR <- par()$usr
  # The current plot dimensions, (width, height), in inches
  PIN <- par()$pin
  # number of x-y pixels for the image
  DIM <- dim(obj)
  # pixel aspect ratio (y/x)
  ARp <- DIM[1]/DIM[2]
  # convert width units to inches
  WIDi <- width/(USR[2]-USR[1])*PIN[1]
  # height in inches
  HEIi <- WIDi * ARp
  # height in units
  HEIu <- HEIi/PIN[2]*(USR[4]-USR[3])
  rasterImage(image = obj,
              xleft = x-(width/2), xright = x+(width/2),
              ybottom = y-(HEIu/2), ytop = y+(HEIu/2),
              interpolate = interpolate)
}
```

### 2 Simulation code

#### 2.1 Definition of functions used for simulations

Written by Richard M. Merrill and Alexander E. Hausmann

August 2014 & June-October 2020

 &

Simulations were run in R 3.6.1

**NOTE**: R introduced a new random number generator (rng) as default with version 3.6. We use this default rng; our results can therefore only be reproduced when run with version 3.6+

**NOTE**: We had all these functions saved within one R script, called `functions.R`, which we then called in our actual simulation scripts via the `source` command (see also next section).

Functions:

##### 2.1.1 Produce indices for the genomic loci

The indices created by this function signalize the following functions where to look for certain loci in the table. What these loci mean will become clear in the explanation for the next function. This is a sole helper function, making the code below easier to follow (and a little faster)

```
loci_indices <- function(number_loci) {
    return(list(i_attr_novel = 6:(6 +
        number_loci - 1), i_attr_ances = (6 +
        number_loci):(6 + 2 * number_loci -
        1), i_original = c(5, 6:(6 +
        number_loci - 1), (6 + number_loci):(6 +
        2 * number_loci - 1)), i_copy = c(6 +
```

```
        2 * number_loci, (6 + 2 * number_loci +
        1):(6 + 3 * number_loci), (6 +
        3 * number_loci + 1):(6 + 4 *
        number_loci)), i_attr_all = c(6:(6 +
        number_loci - 1), (6 + number_loci):(6 +
        2 * number_loci - 1), (6 + 2 *
        number_loci + 1):(6 + 3 * number_loci),
        (6 + 3 * number_loci + 1):(6 +
            4 * number_loci)), i_all = c(c(5,
        6:(6 + number_loci - 1), (6 +
            number_loci):(6 + 2 * number_loci -
            1)), c(6 + 2 * number_loci,
        (6 + 2 * number_loci + 1):(6 +
            3 * number_loci), (6 + 3 *
            number_loci + 1):(6 + 4 *
            number_loci))), total_number_loci = 2 *
        number_loci + 1))
}
```

##### 2.1.2 Create starting population

Total population gets safed in a list, filled with `length(patches)` tables (one table for each patch). All tables are of type integer. One row of a table represents one individual.

In columns `[indices$i_copy]` (starting column name with "copy") we save the genotype of a male that a female mated with. These columns are used later for mutation and recombination during production of offspring. They only matter for mated female individuals. We do realize that this is taking up a lot of memory, but there seems to be no more efficient way. If a female would only produce offspring once, after mating, we could directly give a pointer to the dad during the mutation and the recombination function, without having to save the genotype of the dad. However, this is not the case in our simulations, where a female can continue laying after the dad died. Saving all genotypes from fathers in a "dads database" does not seem more inefficient.

**Explanation of the columns** (except for the loci that are copies of the mate, *i.e.* columns `[indices$i_copy]`).

- sex: 0 or 1 (male or female)
- age: varies between 0 and 5 (individuals with 5 die in the beginning of a cycle)
- patch: ID of the patch, varying between 1 and length(patches)
- mated: 0 or 1 (unmated or mated; only relevant for females)

**NOTE**: Loci (all diallelic): 'a' highlights the recessive, 'A' the dominant state

- Colour: 0 = "aa" (ancestral colour), 1 ="Aa" or "aA" (novel colour), 2 = "AA" (novel colour)
- R_dom_xxx: 0 = "aa" (no attraction to novel colour), 1 ="Aa" or "aA" (attraction to novel colour), 2 = "AA" (attraction to novel colour)
- W_dom_xxx: exact same logic

Creating the starting population:

1. Each central patch gets filled with the same number of individuals, and each non-central patch with the same number
2. We fill each patch with equally many males and females (at unequal patch size, one more female)
3. All ages (0-5) appear equally often within a patch and within the sexes (unless total patch size cannot be divided by 6, in that case lower age classes appear more often); if patch size divided by 6 does not result in an even number, ages cannot be distributed evenly over the two sexes; then there is one more male of age 0, one more female of age 1, one more male of age 2, one more female of age 3 and so on

4. Each individual within a patch gets an identifier = patch_ID
5. `locus_colour` gets set to 0 for all individuals
6. All loci for attraction to novel colour (and the copies of the mate) get set to 0
7. All loci for attraction to ancestral colour (and the copies of the mate) get set to 2

```r
beginners <- function(number_per_central,
    number_per_rest, patches, females_mated,
    number_loci) {
    # Set up empty population list
    population <- vector(mode = "list",
        length = patches)

    # Create populations per patch (a
    # sparse matrix is not faster here,
    # but actually much slower, it
    # seems)
    for (create_patch in as.integer(1:patches)) {
        # If all females are mated, give all
        # females a 1 in the mated column
        if (females_mated) {
            # Population may look different in
            # central patch or non-central
            # patch, if we decided to fill it
            # with different numbers of
            # individuals
            if (create_patch %in% central_vertical) {
                population[[create_patch]] <- cbind((rep(c(0L,
                  1), ceiling(number_per_central/2)))[1:number_per_central],
                  c(rep(0L:5L, each = floor(number_per_central/6)),
                    rep(0L, number_per_central%%6)),
                  rep(create_patch,
                    number_per_central),
                  (rep(c(1L, 0), ceiling(number_per_central/2)))[1:number_per_central],
                  matrix(rep(c(rep(0L,
                    number_loci + 1),
                    rep(2L, number_loci),
                    rep(0L, number_loci +
                      1), rep(2L, number_loci)),
                    each = number_per_central),
                    ncol = number_loci *
                      4 + 2))
            } else {
                population[[create_patch]] <- cbind((rep(c(0L,
                  1), ceiling(number_per_rest/2)))[1:number_per_rest],
                  c(rep(0L:5L, each = floor(number_per_rest/6)),
                    rep(0L, number_per_rest%%6)),
                  rep(create_patch,
                    number_per_rest),
                  (rep(c(1L, 0), ceiling(number_per_rest/2)))[1:number_per_rest],
                  matrix(rep(c(rep(0L,
                    number_loci + 1),
                    rep(2L, number_loci),
                    rep(0L, number_loci +
                      1), rep(2L, number_loci)),
```

```
                each = number_per_rest),
                ncol = number_loci *
                  4 + 2))

        }
    } else {
        # If females are not mated, give all
        # females a 0 in the mated column
        # Population may look different in
        # central patch or non-central
        # patch, if we decided to fill it
        # with different numbers of
        # individuals
        if (create_patch %in% central_vertical) {
            population[[create_patch]] <- cbind((rep(c(0L,
              1), ceiling(number_per_central/2)))[1:number_per_central],
              c(rep(0L:5L, each = floor(number_per_central/6)),
                rep(0L, number_per_central%%6)),
              rep(create_patch,
                number_per_central),
              rep(0L, number_per_central),
              matrix(rep(c(rep(0L,
                number_loci + 1),
                rep(2L, number_loci),
                rep(0L, number_loci +
                  1), rep(2L, number_loci)),
                each = number_per_central),
                ncol = number_loci *
                  4 + 2))
        } else {
            population[[create_patch]] <- cbind((rep(c(0L,
              1), ceiling(number_per_rest/2)))[1:number_per_rest],
              c(rep(0L:5L, each = floor(number_per_rest/6)),
                rep(0L, number_per_rest%%6)),
              rep(create_patch,
                number_per_rest),
              rep(0L, number_per_rest),
              matrix(rep(c(rep(0L,
                number_loci + 1),
                rep(2L, number_loci),
                rep(0L, number_loci +
                  1), rep(2L, number_loci)),
                each = number_per_rest),
                ncol = number_loci *
                  4 + 2))

        }
    }
    # Give names to columns (R for loci
    # coding for attraction to novel, W
    # for loci coding for attraction to
    # ancestral)
    colnames(population[[create_patch]]) <- c("sex",
```

```
            "age", "patch", "mated",
            "Colour", paste0("R_dom_",
                1:number_loci), paste0("W_dom_",
                1:number_loci), "copyColour",
            paste0("copyR_dom_", 1:number_loci),
            paste0("copyW_dom_", 1:number_loci))
        # Make everything integer
        mode(population[[create_patch]]) <- "integer"
    }

    return(population)
}
```

###### 2.1.3 Predation

Predators sample butterflies. Once threshold is reached, no longer attack that colour pattern

```
predation <- function(individuals, nrow_indiv) {
    # Determine predation threshold
    if ((individuals[1, 3] %in% central_vertical) &
        (n %in% birds_die:birds_return)) {
        Q <- reduced_predation
    } else {
        Q <- pred_constant
    }

    if (Q > 0) {
        # Determine novel- and
        # ancestral-coloured butterflies
        novelYN <- individuals[, 5] !=
            0L
        living_Aa_AA <- (1:nrow_indiv)[novelYN]
        living_aa <- (1:nrow_indiv)[!novelYN]
        number_living_Aa_AA <- length(living_Aa_AA)
        number_living_aa <- length(living_aa)

        # Predation by colour (maximally
        # sample Q individuals of each type)
        novel_killed <- living_Aa_AA[sample.int(number_living_Aa_AA,
            size = min(Q, number_living_Aa_AA))]
        ancestr_killed <- living_aa[sample.int(number_living_aa,
            size = min(Q, number_living_aa))]
        return(individuals[-c(novel_killed,
            ancestr_killed), , drop = F])
    } else {
        return(individuals)
    }
}
```

###### 2.1.4 Set up attraction traits

Calculate attractions based on equation in Duenez Guzman *et al.* (2009) (hybrid speciation) paper. Also calculates `max_attr` depending on `strength_of_attraction` (alpha in Duenez Guzman *et al.* (2009))

Calculate number of Aa (=1L) and AA (=2L) loci:

```
attraction_dom <- function(x) {
    # Count alleles that are not aa
    return(rowSums(x > 0L))
}
```

Attraction to a certain colour is calculated in one function and scaled to the maximum attraction

```
attraction_colour <- function(ind_table) {
    # Calculate attraction score
    # weighted by maximum attraction
    attraction <- exp(strength_of_attraction *
        ((attraction_dom(ind_table)/number_loci) -
            1/2))/max_attr
    return(attraction)
}
```

###### 2.1.5 Mating

Mating is dependent on encounter, which is random with respect to males within a patch, then: Mating is proportional to a males attraction for a females colour pattern. EXAMPLE: if a male has all loci for attraction to novel colour at the dominant state (Aa (=1L) or AA (=2L)) probability of mating with a novel-coloured female is 1. If not all loci are at this state, the probability is adjusted (and will depend on `strength_of_attraction`). E.g. `strength_of_attraction` = 4, male has 2 loci with novel allele, `number_loci` = 5; THEN: attraction to novel-coloured female = 0.09071795

```
boy_meets_girl <- function(boyz_rows,
    unmated_rows, individuals) {

    # Determine length of boys and
    # unmated girls vector
    nrow_boyz <- length(boyz_rows)
    nrow_girls <- length(unmated_rows)
    # Shuffle through the unmated girls
    girls_id <- sample.int(nrow_girls)

    # Determine maximum sample size
    # (since males can only mate
    # max_mate times, not all females
    # will get mated)
    sample_size <- min(nrow_girls, nrow_boyz *
        max_mate)

    # If all females have novel colour,
    # it is easy
    if (sum(individuals[unmated_rows,
        5] > 0L) == nrow_girls) {
        # Sample males depending on their
        # attraction to novel colour By
        # repeating the male IDs max_mate
        # times, we have an easy trick how
        # to cap at max_mate draws
        daddies <- sample(rep(1:nrow_boyz,
            max_mate), size = sample_size,
            prob = rep(attraction_colour(individuals[boyz_rows,
                indices$i_attr_novel,
```

```
            drop = F]), max_mate))

        # If all females have ancestral
        # colour, it is also easy
    } else {
        if (sum(individuals[unmated_rows,
            5] == 0L) == nrow_girls) {
            daddies <- sample(rep(1:nrow_boyz,
                max_mate), size = sample_size,
                prob = rep(attraction_colour(individuals[boyz_rows,
                  indices$i_attr_ances,
                  drop = F]), max_mate))

            # If females are differently
            # coloured:
        } else {
            # Get attraction for novel pattern
            # of each male
            attr_novel <- attraction_colour(individuals[boyz_rows,
                indices$i_attr_novel,
                drop = F])
            # Get attraction for ancetral colour
            # of each male
            attr_ancestr <- attraction_colour(individuals[boyz_rows,
                indices$i_attr_ances,
                drop = F])
            # Sample males female-by-female
            daddies <- rep(NA, sample_size)
            for (x in 1:sample_size) {
                daddies[x] <- ifelse(individuals[x,
                  5] > 0L, sample.int(nrow_boyz,
                  prob = attr_novel,
                  size = 1), sample.int(nrow_boyz,
                  prob = attr_ancestr,
                  size = 1))
            }
            # Now we check how often each male
            # appeared
            freq_daddy <- table(daddies)
            # If one (or multiple) appeared more
            # than max_mate times, repeat while
            # loop until even distribution...
            freq_max_mateplus <- freq_daddy >
                max_mate
            sum_freq_max_mateplus <- sum(freq_max_mateplus)
            while (sum_freq_max_mateplus >
                0) {
                girl_new <- c()
                # Find which girls are affected
                for (correct in 1:sum_freq_max_mateplus) {
                  matchy <- daddies ==
                    as.numeric(names(freq_daddy)[freq_max_mateplus][correct])
                  girl_new <- c(girl_new,
```

```r
                (1:sample_size)[matchy][5:sum(matchy)])
            }
            # Extract males, which still have
            # not mated max_mate times
            new_boyz <- c(as.numeric(names(freq_daddy)[freq_daddy <
              max_mate]), (1:nrow_boyz)[!(1:nrow_boyz) %in%
              names(freq_daddy)])
            l_newb <- length(new_boyz)
            # Sample from those
            for (x in girl_new) {
              # has to be so complicated since
              # new_boyz may be of length = 1 and
              # then sample() does nonsense
              daddies[x] <- ifelse(individuals[unmated_rows[girls_id][x],
                5] > 0L, new_boyz[sample.int(l_newb,
                prob = attr_novel[new_boyz],
                size = 1)], new_boyz[sample.int(l_newb,
                prob = attr_ancestr[new_boyz],
                size = 1)])
            }
            # Repeat check
            freq_daddy <- table(daddies)
            freq_max_mateplus <- freq_daddy >
              max_mate
            sum_freq_max_mateplus <- sum(freq_max_mateplus)
        }
      }
    }

    # Assign these females as mated and
    # fill male genotype in columns
    # [indices$i_copy] (important for
    # recombination later)
    individuals[unmated_rows[girls_id[1:sample_size]],
        4] <- 1L
    individuals[unmated_rows[girls_id[1:sample_size]],
        indices$i_copy] <- individuals[boyz_rows[daddies],
        indices$i_original, drop = F]

    return(individuals)
}
```

###### 2.1.6 Generating offspring

Number of new individuals is determined by patch capacity, a Poisson distribution around 20 (or 0.5 during bottleneck). Can be thought of as ~ number available Passiflora. Females within a patch are chosen at random to lay an egg. 'Laying' depends on `chance_encounter` with a male and then the males' attraction for the females colour pattern determines whether she does or doesn't lay (similar to mating).

Mutation can occur during production of new individuals in ANY attraction allele. We assume that only one locus in an individual can be hit once by mutation

```r
Mutation <- function(individuals, nrow_indiv,
    number_eggs, baby_id) {
```

```
    # Number of mutations to be
    # distributed. Individuals
    # (number_eggs) times positions that
    # can mutate (4*number_loci) times
    # mutation rate (prob_mutation)
    numb_mut <- rpois(n = 1, lambda = number_eggs *
        4 * number_loci * prob_mutation)
    if (numb_mut > 0) {
        # Position them (sampling without
        # replacement, i.e. no
        # double-mutation)
        ind_pos <- sample.int(number_eggs *
            4 * number_loci, size = numb_mut)
        # Check for homozygotes in the loci
        # responsible for attraction to
        # colours (including the gamete
        # copies of the father!)
        classy <- individuals[baby_id,
            indices$i_attr_all, drop = F][ind_pos] !=
            1L
        # All homozygotes become
        # heterozygotes
        individuals[baby_id, indices$i_attr_all][ind_pos][classy] <- 1L
        # Heterozygotes randomly become one
        # of the types of homozygote
        individuals[baby_id, indices$i_attr_all][ind_pos][!classy] <- sample(c(0L,
            2L), size = sum(!classy),
            replace = T)
    }
    return(individuals)
}
```

Recombination of parental alleles (no linkage)

```
Recombination <- function(individuals,
    nrow_indiv, number_eggs, baby_id) {
    # Selection of gamete in
    # heterozygous loci
    hetero <- individuals[baby_id, indices$i_all] ==
        1L
    individuals[baby_id, indices$i_all][hetero] <- sample(c(0L,
        2L), sum(hetero), replace = T)
    # Recombination. Easy trick: We do
    # this by calculating the mean, e.g.
    # 0 and 2 gives a 1
    for (locus in 1:indices$total_number_loci) {
        individuals[baby_id, indices$i_original[locus]] <- as.integer(rowMeans(individuals[baby_id,
            c(indices$i_original[locus],
                indices$i_copy[locus]),
            drop = F]))
    }
    return(individuals)
}
```

The main function for generating offspring (here is where sexual conflict plays a role):

```r
make_babies <- function(individuals,
    nrow_indiv, mated_females, boyz_rows) {

    # Work out number of expected eggs
    # per patch

    if (individuals[1, 3] %in% central_vertical &
        n %in% 1:(bottleneck_ends -
            1) & bottleneck) {
        expected_eggs <- 0.5
    } else {
        expected_eggs <- 20L
    }

    # Sample mothers

    # If no sexual conflict or no males
    # alive, all females have the same
    # chance
    if (!sex_conflict | length(boyz_rows) ==
        0) {
        number_eggs <- rpois(1, expected_eggs)
        if (number_eggs > 0) {
            babies <- mated_females[sample.int(length(mated_females),
                size = number_eggs,
                replace = T)]
            eggs_laid <- T
        } else {
            eggs_laid <- F
        }
    } else {
        # If there is sexual conflict, but
        # all females look the same, all
        # females again have the same chance
        nrow_mated_females <- length(mated_females)
        if (sum(individuals[mated_females,
            5] > 0L) == nrow_mated_females |
            sum(individuals[mated_females,
                5] == 0L) == nrow_mated_females) {
            number_eggs <- rpois(1,
                expected_eggs)
            if (number_eggs > 0) {
                babies <- mated_females[sample.int(length(mated_females),
                  size = number_eggs,
                  replace = T)]
                eggs_laid <- T
            } else {
                eggs_laid <- F
            }
            # If they look differently, sexual
            # conflict starts playing a role:
        } else {
            # Calculate average male attraction
```

```
        # to novel or ancestral colour
        mean_attr_novel <- mean(attraction_colour(individuals[boyz_rows,
            indices$i_attr_novel,
            drop = F]))
        mean_attr_ancestr <- mean(attraction_colour(individuals[boyz_rows,
            indices$i_attr_ances,
            drop = F]))
        # Calculate for each female the
        # harassment probability (this is
        # the average male attraction to the
        # colour she displays)
        prob_harass <- ifelse(individuals[mated_females,
            5] > 0L, mean_attr_novel,
            mean_attr_ancestr)

        # This calculates the encounter
        # probability
        chance_encounter <- 1 -
            ((1 - prob_encounter)^length(boyz_rows))
        # This calculates the probability of
        # an egg being laid after encounter
        # in ONE single attempt It depends
        # on encounter probability and
        # average harassment probability
        P_with_encounter <- chance_encounter *
            (1 - mean(prob_harass))
        # This calculates the total
        # proportion of eggs laid after
        # encounter
        Prop_with_encounter <- P_with_encounter/(P_with_encounter +
            (1 - chance_encounter))
        if (Prop_with_encounter >
            1 | Prop_with_encounter <
            0) {
            stop("Prop_with_encounter makes no sense. Something wrong in the math.")
        }
        # Number of eggs laid after
        # encounter and without encounter
        # are Poisson distributed
        eggs_after_encounter <- rpois(1,
            expected_eggs * Prop_with_encounter)
        eggs_without_encounter <- rpois(1,
            expected_eggs * (1 -
              Prop_with_encounter))
        number_eggs <- eggs_after_encounter +
            eggs_without_encounter

        # A part of the eggs get sampled to
        # the females based on harassment
        # probability (the less likely they
        # are harassed), the other part of
        # the eggs get sampled with equal
        # probability
```

```
        nrow_mated_females <- length(mated_females)
        if (number_eggs > 0) {
            if (eggs_after_encounter >
              0) {
              harassed_mothers <- sample.int(nrow_mated_females,
                size = eggs_after_encounter,
                prob = 1 - prob_harass,
                replace = T)
            } else {
              harassed_mothers <- c()
            }
            if (eggs_without_encounter >
              0) {
              unharassed_mothers <- sample.int(nrow_mated_females,
                size = eggs_without_encounter,
                replace = T)
            } else {
              unharassed_mothers <- c()
            }
            eggs_laid <- T
            babies <- mated_females[c(harassed_mothers,
              unharassed_mothers)]
        } else {
            eggs_laid <- F
        }
    }
}

# If eggs were laid, add newborns
# and perform mutation+recombination
if (eggs_laid) {
    # Add babies to dataset
    individuals <- individuals[c(1:nrow_indiv,
        babies), , drop = F]
    baby_id <- nrow_indiv + (1:number_eggs)
    # Sample sexes and make them unmated
    # and age 0
    individuals[baby_id, 1] <- sample(c(0L,
        1L), size = number_eggs,
        replace = TRUE)
    individuals[baby_id, 2] <- 0L
    individuals[baby_id, 4] <- 0L

    # Mutation & Recombination

    individuals <- Mutation(individuals,
        nrow_indiv, number_eggs,
        baby_id)
    individuals <- Recombination(individuals,
        nrow_indiv, number_eggs,
        baby_id)
}
return(individuals)
```

```
}
```

###### 2.1.7 Dispersal

Only local dispersal possible (1 patch in each dimension). Modulo calculation in both dimensions makes arena torus-shaped

```
fly_away <- function(x) {
    length_x <- length(x)
    # Move horizontally
    x <- (x + sample(c(0, arena_height,
        -arena_height), size = length_x,
        replace = T) - 1)%%patches +
        1
    # Move vertically
    x <- as.integer(floor((x - 1)/arena_height) *
        arena_height + ((x + sample(c(0,
        1, -1), size = length_x, replace = T) -
        1)%%arena_height) + 1)
    return(x)
}
```

###### 2.1.8 LIFE CYCLE

This calls the above functions consecutively

```
life_cycle <- function(individuals,
    n) {

    # Dying of age
    individuals <- individuals[individuals[,
        2] < 5L, , drop = F]

    # If no individuals, next generation
    # is empty
    nrow_indiv <- nrow(individuals)
    if (nrow_indiv > 0) {

        # Ageing
        individuals[, 2] <- individuals[,
            2] + 1L

        # Predation
        individuals <- predation(individuals,
            nrow_indiv)

        # Only do the following steps if
        # there is individuals in the patch
        nrow_indiv <- nrow(individuals)
        if (nrow_indiv > 0) {

            # Determine males
            boyz_rows <- (1:nrow_indiv)[individuals[,
                1] == 1L]
            # Determine mated females
```

```
            mated_females <- (1:nrow_indiv)[individuals[,
                4] == 1L]
            if (length(mated_females) >
                0) {
                # Egg laying etc.
                individuals <- make_babies(individuals,
                  nrow_indiv, mated_females,
                  boyz_rows)
                # Determine males again (new borns
                # added)
                boyz_rows <- (1:nrow(individuals))[individuals[,
                  1] == 1L]
            }
            if (length(boyz_rows) >
                0) {
                # Determine unmated females
                unmated_rows <- (1:nrow(individuals))[individuals[,
                  4] == 0L & individuals[,
                  1] == 0L]
                if (length(unmated_rows) >
                  0) {
                  # Mate these females
                  individuals <- boy_meets_girl(boyz_rows,
                    unmated_rows, individuals)
                }
            }
            # Dispersal
            new_born <- individuals[,
                2] == 0L
            individuals[new_born, 3] <- fly_away(individuals[new_born,
                3])
        }
    }
    return(individuals)
}
```

#### 2.2 Running a simulation under one specific scenario

Below, we present the complete code you can copy-paste into a new R script, to run the simulations. This is not supposed to be run within RStudio, but as a command line program. Arguments can be passed via the command line to the `commandArgs` function. In our simulations, we varied the 4th command line argument (`setting`), which is the row in `parameter_settings.csv` to extract model parameters from. That table has 120 rows. Additionally, we varied the 6th command line argument (`number_loci`) between values 1, 5 and 10 (and each `setting` was run once for each `number_loci`). This gives 120x3 = 360 possible parameter combinations. Each of those was run 2000 times (5th command line argument, `simulations`). Within the simulation code, we additionally randomized between whether a male or a female first mutant occurs, giving another parameter with 2 possible values. Hence, we had a total of 360x2 = 720 parameter combinations, each simulated ~1000 times.

To efficiently do the 360 calls to this function via command line, we used Windows batch scripts (on Mac and Linux, the equivalent can be used).

On Windows, a call to the R script containing below code could look like this (where you'd have to replace all parts inside `__ __` with your own settings): `"__ENTER_R.exe_DIR_HERE__" CMD BATCH --vanilla --slave`

```
"--args __ENTER_PARAMS_SEPERATED_BY_SPACE__" __ENTER_NAME_R_SCRIPT__
```

The reason why we simulated each scenario with a separate call to R is because the longer R runs, the more unnecessary 'garbage' gets accumulated and the slower it becomes. Hence, it makes sense to close and restart R after one scenario was simulated. In total, the code takes about 1-2 months to run.

Explanations on what happens where can be found inside the next code chunk. You can also scroll down some pages to the section "Allele frequency Gif", where some of this comes up again!

Here is the code (it won't be executed in this Markdown though):

```
# Setup

# Fetch command line arguments
params <- commandArgs(trailingOnly = TRUE)
# Main directory
main_dir <- params[1]
# Folder containing Functions.R and
# parameter_settings.csv
info_dir <- params[2]
# Output folder
out_dir <- params[3]
# Which simulation scenario to run
setting <- as.integer(params[4])
# Number of simulations to be run
# (i.e. row of
# parameter_settings.csv)
simulations <- as.integer(params[5])
# Number of loci coding for one
# attraction phenotype
number_loci <- as.integer(params[6])
# Maximum number of females a male
# can mate during one generation In
# all our simulations, we set this
# to 4
max_mate <- as.integer(params[7])

# Replace all \ by /
main_dir <- gsub("\\\\", "/", main_dir)
info_dir <- gsub("\\\\", "/", info_dir)
out_dir <- gsub("\\\\", "/", out_dir)

# Make sure we have a / at the end
# of each path
main_dir <- paste0(trimws(main_dir),
    ifelse(trimws(main_dir) == "", "",
        "/"))
info_dir <- paste0(trimws(info_dir),
    ifelse(trimws(info_dir) == "", "",
        "/"))
out_dir <- paste0(trimws(out_dir), ifelse(trimws(out_dir) ==
    "", "", "/"))

# Load library for sparse matrices
if (!"Matrix" %in% rownames(installed.packages())) {
    install.packages("Matrix")
```

```r
}
suppressMessages(suppressWarnings(library(Matrix)))

# Set working directory
setwd(main_dir)

# Call simulation functions
source(paste0(info_dir, "Functions.R"))

# Load the 120 possible parameter
# combinations we are testing
parameter_table <- read.csv(paste0(info_dir,
    "parameter_settings.csv"), header = T,
    stringsAsFactors = F)


# Parameter constants

# Arena dimensions
arena_width <- 4L
arena_height <- 96L
# Number of central patches (in
# vertical direction (i.e. patches
# 39-58 when 96 in height))
arena_center <- 20L
# Generation when the birds
# disappear
birds_die <- 1L
# Number of individuals per colour
# phenotype picked out during
# relaxed predation
reduced_predation <- 0L
# Generation in which the bottleneck
# in the central patches ends (i.e.
# Passiflora plants are available
# again and egg laying goes back to
# normal)
bottleneck_ends <- 6L
# Maximum age of individuals of
# starting population
max_age <- 5L
# All females in starting population
# are mated
females_mated <- T
# Mutation rate per individual per
# locus per generation
prob_mutation <- 10^-5
# Maximum number of generations per
# simulation
generations <- 2500

# From this, deduce following other
# constants: Total arena patches
```

```
patches <- arena_width * arena_height
# Get central patches (if they
# cannot be perfectly centered,
# round up)
central_vertical <- (1:patches)[((1:patches -
    1)%%arena_height + 1) %in% ifelse(arena_height%%2 ==
    0, ceiling(0.5 * arena_height -
    0.5 * arena_center + 1), ceiling(0.5 *
    (arena_height + 1) - 0.5 * (arena_center -
    1))):ifelse(arena_height%%2 == 0,
    ceiling(0.5 * arena_height + 0.5 *
        arena_center), ceiling(0.5 *
        (arena_height + 1) + 0.5 * (arena_center -
        1)))]
# Get non-central patches
not_central <- (1:patches)[!(1:patches) %in%
    central_vertical]


# Load parameters specific to the
# scenario that should be simulated

# Sexual conflict (yes or no)
sex_conflict <- parameter_table$sex_conflict[setting]
# Number of generations no predation
# in central patches
birds_return <- parameter_table$predator_absence[setting]
# Whether we start off from a
# bottleneck population
bottleneck <- parameter_table$bottleneck[setting]
# Number of individuals per colour
# phenotype picked out by predators
pred_constant <- parameter_table$pred_constant[setting]
# Start seed of the first simulation
start_seed <- as.integer(parameter_table$start_seed[setting])

# Following parameters are only
# relevant if sex_conflict==TRUE

# alpha in Duenez-Guzman et al.
# (2009), for attraction of a male
# to certain colour A value of 1
# means that a male with all
# attraction loci for colour X at Aa
# or AA state and all attraction
# loci for colour Y at aa state will
# have a preference index for X of
# about 0.73 (where preference index
# is (associations with X)/(all
# associations)).  A value of 1
# means that the same male will have
# a preference index of about 0.95
strength_of_attraction <- parameter_table$strength_attract[setting]
```

```r
# Probability that a female
# encounters a male during egg
# laying
prob_encounter <- parameter_table$prob_encounter[setting]
# From this, calculate maximum
# attraction value
max_attr <- exp(strength_of_attraction *
    (1 - 1/2) * 1)


# Setup population

normal_pop <- beginners(number_per_central = 114,
    number_per_rest = 114, females_mated = females_mated,
    patches = patches, number_loci = number_loci)
# Remember total number of columns
# for one of the tables from the
# list
number_columns <- 6 + number_loci *
    4


# Set up data collection

# We create a very generously big
# results table with 50% of maximum
# number of possible rows
results <- sparseMatrix(i = integer(0),
    j = integer(0), x = 0, dims = c(as.integer(simulations *
        0.5 * generations), 12 + 8 *
        number_loci))

colnames(results) <- c("Scenario", "Simulation",
    "Generation", "start_seed_of_simulation",
    "sex_mutant", "mutator_patch", "N_non_central",
    "N_central", "Cnc_het", paste0("Rnc",
        1:number_loci, "_het"), paste0("Wnc",
        1:number_loci, "_het"), "Cnc_hom",
    paste0("Rnc", 1:number_loci, "_hom"),
    paste0("Wnc", 1:number_loci, "_hom"),
    "Cc_het", paste0("Rc", 1:number_loci,
        "_het"), paste0("Wc", 1:number_loci,
        "_het"), "Cc_hom", paste0("Rc",
        1:number_loci, "_hom"), paste0("Wc",
        1:number_loci, "_hom"))
# Determine the columns that are
# corresponding to counts for colour
# pattern mutants
mutant_columns <- c(9, 9 + number_loci *
    c(2, 4, 6) + 1:3)
# Get the relevant columns to
# extract information from
# simulation results loci_indices is
```

```r
# a function defined in Functions.R
indices <- loci_indices(number_loci)

# Install a counter to know the row
# where to save the results
sim_counter <- 1
# Determine steps of 100 simulations
# after which results should be
# saved.  This is for safety, in
# case your computer crashes, etc.,
# then, by setting the right seed,
# you can pick up where the
# simulations stopped before.
simu_to_save <- seq(0, ifelse((simulations%%100) ==
    0, simulations - 100, simulations),
    100)


# Simulation runs

# Repeat this as many times as the
# 'simulations' parameter says

for (sim in as.integer(1:simulations)) {

    # Give info about the new simulation
    print(paste0("Setting: ", setting,
        " -- Simulation: ", sim))
    print(Sys.time())

    # Set seed
    set.seed(start_seed)

    # Make new individuals and introduce
    # mutation at colour pattern locus

    # Get starting population
    individuals <- normal_pop
    # Place a mutation at the colour
    # locus to turn one new-born female
    # from a central patch into a
    # novel-coloured butterfly
    mutator_patch <- sample(central_vertical,
        size = 1)
    mutator_indiv <- sample.int(114,
        size = 1)
    individuals[[mutator_patch]][mutator_indiv,
        5] <- 1L
    # Remember sex of mutant
    sex_mutator <- unname(individuals[[mutator_patch]][mutator_indiv,
        1])
    # Make that individual age 0
    # (new-born in previous generation)
```

```r
individuals[[mutator_patch]][mutator_indiv,
    2] <- 0L

# Get number of individuals
total_individuals <- sum(lengths(individuals))/number_columns

# Run through generations

for (n in as.integer(1:generations)) {
    print(paste0("--", n, "--",
        total_individuals))

    # Run current generation
    for (unito in 1:patches) {
        individuals[[unito]] <- life_cycle(individuals[[unito]],
            n = unito)
    }

    # Equally fast alternative:
    # individuals <- lapply(individuals,
    # life_cycle)

    # Here is some nice code to check
    # which steps in the code take the
    # longest (via profiling):
    # library(proftools)
    # Rprof('prof.out')
    # lapply(individuals, life_cycle)
    # Rprof(NULL)
    # print(summaryRprof('prof.out'))

    # Change list position of migrators
    # (make them 'officially' migrate);
    # this has to happen in a separate
    # step, such that the migrators only
    # appear in their new patch in the
    # next loop
    for (unito in 1:patches) {
        curr_n <- nrow(individuals[[unito]])
        if (curr_n > 0) {
            changers <- (1:curr_n)[individuals[[unito]][,
              3] != unito]
            if (length(changers) >
              0) {
              for (move_around in changers) {
                individuals[[individuals[[unito]][move_around,
                  3]]] <- rbind(individuals[[individuals[[unito]][move_around,
                  3]]], individuals[[unito]][move_around,
                  , drop = F])
                # Or do:
                # individuals[[individuals[[unito]][move_around,3]]]<-
                # do.call(rbind,list(individuals[[individuals[[unito]][move_around,3]]],
                # individuals[[unito]][move_around,,drop=F]))
```

```
            }
            individuals[[unito]] <- individuals[[unito]][-changers,
              , drop = F]
          }
      }
  }


  # Collect data

  # Count total number of individuals,
  # as well as central-square
  # individuals
  total_individuals <- as.integer(sum(lengths(individuals))/number_columns)
  sum_is_central <- as.integer(sum(lengths(individuals[central_vertical]))/number_columns)
  # Additionally to saving overall
  # info (setting, simulation number,
  # generation, seed, sex of first
  # mutant, patch where first mutant
  # occured, number of total
  # individuals, number of individuals
  # in central squares), we also count
  # number of homozygotes and
  # heterozygotes for each locus
  # (colour or attraction), separately
  # for central and non-central
  # squares
  results[sim_counter, ] <- c(setting,
      sim, n, start_seed, sex_mutator,
      mutator_patch, total_individuals -
          sum_is_central, sum_is_central,
      as.integer(Reduce(`+`, lapply(individuals[not_central],
          function(x) colSums(x[,
            indices$i_original,
            drop = F] == 1L)))),
      as.integer(Reduce(`+`, lapply(individuals[not_central],
          function(x) colSums(x[,
            indices$i_original,
            drop = F] == 2L)))),
      as.integer(Reduce(`+`, lapply(individuals[central_vertical],
          function(x) colSums(x[,
            indices$i_original,
            drop = F] == 1L)))),
      as.integer(Reduce(`+`, lapply(individuals[central_vertical],
          function(x) colSums(x[,
            indices$i_original,
            drop = F] == 2L)))))

  # Increase the simulation counter by
  # 1
  sim_counter <- sim_counter +
      1
```

```
        # Break loop if mutation died out
        if (sum(results[sim_counter -
            1, mutant_columns]) == 0)
            break
    }

    # Increase seed by 1
    start_seed <- start_seed + 1L

    # Remove R garbage created in this
    # simulation (not sure this actually
    # helps, as it also eats up some
    # time and is only partly
    # successful...)
    gc()

    # Save results (as sparse matrix) if
    # current simulation is a multiple
    # of 100
    if (sim %in% simu_to_save) {
        writeMM(results, file = paste0(out_dir,
            "loci_", number_loci, "_setting_",
            setting, "_mate_", max_mate,
            "_simulations_", simulations,
            ".txt"))
    }
}

# Save results (as sparse matrix) at
# very end
writeMM(results, file = paste0(out_dir,
    "loci_", number_loci, "_setting_",
    setting, "_mate_", max_mate, "_simulations_",
    simulations, ".txt"))
```

#### 2.3 Combining results into tables

Finally, we get a folder with 360 individual small files, which we now combine into three larger tables (one for each set of loci coding for male attraction - those cannot be all in one table, as we need different amounts of columns in the tables). Here is the code (it won't be executed in this Markdown though):

```
# Use the 'matrix' package to read
# in previously saved files
library(matrix)

for (ll in c(1, 5, 10)) {

    # Get list of files matching pattern
    # with current number of loci
    listo <- list.files("YOUR_OUTPUT_DIR_FROM_BEFORE",
        pattern = paste0("loci_", ll,
            "_"), full.names = T)

    # Order this list by the setting ID
```

```r
# in the name of the file (ranging
# between 1 and 120)
ids <- unname(sapply(listo, function(x) as.numeric(strsplit(x,
    "_")[[1]][length(strsplit(x,
    "_")[[1]]) - 4])))
listo <- listo[order(ids)]

# Read in the first file (using
# readMM from the 'matrix' package)
resultsn <- readMM(listo[1])
# Remove rows of all 0s (appended at
# the end of the table)
resultsn <- resultsn[resultsn[,
    1] != 0, ]
# Convert to data frame
resultsn <- as.data.frame(as.matrix(resultsn))
# Give column names (some names
# depend on how many loci for male
# attraction were simulated)
namis <- c("Scenario", "Simulation",
    "Generation", "start_seed_of_simulation",
    "sex_mutant", "mutator_patch",
    "N_non_central", "N_central",
    "Cnc_het", paste0("Rnc", 1:ll,
        "_het"), paste0("Wnc", 1:ll,
        "_het"), "Cnc_hom", paste0("Rnc",
        1:ll, "_hom"), paste0("Wnc",
        1:ll, "_hom"), "Cc_het",
    paste0("Rc", 1:ll, "_het"),
    paste0("Wc", 1:ll, "_het"),
    "Cc_hom", paste0("Rc", 1:ll,
        "_hom"), paste0("Wc", 1:ll,
        "_hom"))
names(resultsn) <- namis

# Now, we repeat the exact same for
# all other 119 files, and we append
# them to the previous data frame
for (iii in 2:length(listo)) {
    resultsn_sub <- readMM(listo[iii])
    resultsn_sub <- resultsn_sub[resultsn_sub[,
        1] != 0, ]
    resultsn_sub <- as.data.frame(as.matrix(resultsn_sub))
    namis <- c("Scenario", "Simulation",
        "Generation", "start_seed_of_simulation",
        "sex_mutant", "mutator_patch",
        "N_non_central", "N_central",
        "Cnc_het", paste0("Rnc",
            1:ll, "_het"), paste0("Wnc",
            1:ll, "_het"), "Cnc_hom",
        paste0("Rnc", 1:ll, "_hom"),
        paste0("Wnc", 1:ll, "_hom"),
        "Cc_het", paste0("Rc", 1:ll,
```

```
                "_het"), paste0("Wc",
                1:ll, "_het"), "Cc_hom",
            paste0("Rc", 1:ll, "_hom"),
            paste0("Wc", 1:ll, "_hom"))
        names(resultsn_sub) <- namis
        # Bind it to previous data frame
        resultsn <- rbind(resultsn,
            resultsn_sub)
    }

    # Write the combined table to a csv
    # file
    write.csv(resultsn, paste0("simulation_",
        ll, ifelse(ll == 1, "locus",
            "loci"), ".csv"), row.names = F,
        quote = F)

    # Finally, if you want, you can
    # delete all the small files that we
    # now combined with something like:
    # file.remove(listo)
}
```

### 3 Simulation results

#### 3.1 Data read-in and manipulation

Read in parameter settings for simulations

```
param <- read.csv("Data/Simulations/parameter_settings.csv",
    header = T, stringsAsFactors = F)
# For some aesthetics later, we
# change TRUE and FALSE to 0 and 1
param$sex_conflict <- as.numeric(param$sex_conflict)
param$bottleneck <- as.numeric(param$bottleneck)
```

The columns indicate:

- **sex_conflict** = Whether sexual conflict was absent (0) or present (1) in the current scenario
- **bottleneck** = Whether a bottleneck was absent (0) or present (1) in the current scenario
- **prob_encounter** = Parameter $e$ in the manuscript. Probability a female encounters one particular male individual of a patch during egg laying. Can take values of 0.001, 0.0025, 0.005 or 0.01
- **predator_absence** = Parameter $T$ in the manuscript. Number of generations that predators are absent from central patches
- **strength_attract** = Parameter $/alpha$ in the manuscript. Strength of attraction based on colour pattern in males. Can take values of 1 or 3. The higher the value, the stronger the colour-based attraction
- **pred_constant** = Parameter $Q$ in the manuscript. Learning threshold of predators, *i.e.* number of butterflies of a certain colour pattern that birds eat during one generation
- **start_seed** = First seed of first Simulation under the current scenario

Read in data from 1 attraction locus towards each colour, data from 5 loci, and data from 10 loci

```
l1 <- read.csv("Data/Simulations/simulation_1locus.csv",
    header = T, stringsAsFactors = F)
```

```r
l5 <- read.csv("Data/Simulations/simulation_5loci.csv",
    header = T, stringsAsFactors = F)
l10 <- read.csv("Data/Simulations/simulation_10loci.csv",
    header = T, stringsAsFactors = F)
```

The columns indicate:

- **Scenario** = 'ID' of evolutionary scenario, *i.e.* row in `param` table corresponding to the relevant combination of parameter settings
- **Simulation** = ID of simulation run. Ranges from 1 to 2000 for each Scenario
- **Generation** = ID of current number a life cycle was iterated. Starts at 1 and can go up to 2500, if novel colour allele persists in population
- **start_seed_of_simulation** = The seed that was set at the beginning of the current `Simulation`
- **sex_mutant** = Whether the first mutant placed in the arena was a female (=0) or a male (=1)
- **mutator_patch** = The patch in the arena where the first mutant was placed. The arena had a total of 384 patches, with 96 'rows' and 4 'columns'. Their IDs were organized descending by column, e.g. the first column had IDs 1 to 96, the second 97 to 192, and so on. 80 of these patches were considered central and placed in exactly the 'center' of the 'columns' (meaning e.g. at IDs 39 to 58 in the first column, at IDs 135 to 154 in the second column, and so on)
- **N_non_central** = Total number of individuals in non-central patches at the end of the current `Generation`
- **N_central** = Total number of individuals in central patches at the end of the current `Generation`
- **Cnc_het** = Number of individuals in non-central patches at the end of the current `Generation` that are heterozygous at the colour pattern locus

**NOTE: 'XX' in the following denotes the locus ID of one of the two sets of attraction loci. E.g. `R1` is the first locus coding for attraction to the novel warning pattern.**

- **RncXX_het** = Number of individuals in non-central patches at the end of the current `Generation` that are heterozygous for the attraction locus `XX` for attraction to the novel warning pattern
- **WncXX_het** = Number of individuals in non-central patches at the end of the current `Generation` that are heterozygous for the attraction locus `XX` for attraction to the ancestral warning pattern
- **Cnc_hom** = Number of individuals in non-central patches at the end of the current `Generation` that are homozygous at the colour pattern locus
- **RncXX_hom** = Number of individuals in non-central patches at the end of the current `Generation` that are homozygous for the attraction locus `XX` for attraction to the novel warning pattern
- **WncXX_hom** = Number of individuals in non-central patches at the end of the current `Generation` that are homozygous for the attraction locus `XX` for attraction to the ancestral warning pattern
- **Cc_het** = Number of individuals in central patches at the end of the current `Generation` that are heterozygous at the colour pattern locus
- **RcXX_het** = Number of individuals in central patches at the end of the current `Generation` that are heterozygous for the attraction locus `XX` for attraction to the novel warning pattern
- **WcXX_het** = Number of individuals in central patches at the end of the current `Generation` that are heterozygous for the attraction locus `XX` for attraction to the ancestral warning pattern
- **Cc_hom** = Number of individuals in central patches at the end of the current `Generation` that are homozygous at the colour pattern locus
- **RcXX_hom** = Number of individuals in central patches at the end of the current `Generation` that are homozygous for the attraction locus `XX` for attraction to the novel warning pattern
- **WcXX_hom** = Number of individuals in central patches at the end of the current `Generation` that are homozygous for the attraction locus `XX` for attraction to the ancestral warning pattern

For each parameter setting, count number of successes (*i.e.* mutation survived until generation 2500). This does NOT include sex as a factor yet, which we will add as another parameter below. The new table contains 9 columns. The first 7 showing the setting for 7 of the relevant parameters, the other two showing number of 'successes' (*i.e.* mutation survives until generation 2500) and number of 'tries', *i.e.* how many simulations were run.

```
outcome <- data.frame(rbind(param[,
    1:6], param[, 1:6], param[, 1:6]),
    number_loci = as.integer(c(rep(c(1,
        5, 10), each = nrow(param)))),
    number_successes = c(sapply(1:nrow(param),
        function(x) sum(l1$Scenario ==
            x & l1$Generation == 2500)),
        sapply(1:nrow(param), function(x) sum(l5$Scenario ==
            x & l5$Generation == 2500)),
        sapply(1:nrow(param), function(x) sum(l10$Scenario ==
            x & l10$Generation == 2500))),
    number_tries = rep(2000, 3), stringsAsFactors = F)
```

Now we add as an additional parameter the sex of the first mutant. For this, we calculate for each parameter combination how many of the ‘successes’ and how many of the ‘tries’ came from male first mutants.

```
male_successes <- c()
male_first_mutant <- c()
for (i in 1:nrow(outcome)) {
    # Get current parameter setting (not
    # including the number of loci
    # setting)
    current_param <- ((i - 1)%%nrow(param)) +
        1
    # Get number of loci setting
    current_loci <- ((i - 1)%/%nrow(param)) +
        1
    # First, find which rows match the
    # current scenario and have a male
    # as first mutant
    scenario_mutant_check <- get(c("l1",
        "l5", "l10")[current_loci])$Scenario ==
        current_param & get(c("l1",
        "l5", "l10")[current_loci])$sex_mutant ==
        1
    # Count number of 'successes' (=2500
    # generations reached) where first
    # mutant was male in the respective
    # dataset
    male_successes <- c(male_successes,
        sum(get(c("l1", "l5", "l10")[current_loci])$Generation[scenario_mutant_check] ==
            2500))
    # Count number of 'trials'
    # (Generation = 1 = starting
    # generation) where first mutant was
    # male in the respective dataset
    male_first_mutant <- c(male_first_mutant,
        sum(get(c("l1", "l5", "l10")[current_loci])$Generation ==
            1 & scenario_mutant_check))
}
```

We create a new table by duplicating the previous table, in each adjusting number of successes and number of tries to either male or female first mutants.

```
# First, create the table for male
# first mutants
outcome_males <- outcome
outcome_males$sex_first_mutant <- "M"
outcome_males <- outcome_males[, c(1:7,
    10, 8:9)]
outcome_males$number_successes <- male_successes
outcome_males$number_tries <- male_first_mutant

# Same for females. Successes and
# failures are always Total minus
# male first mutants
outcome_females <- outcome
outcome_females$sex_first_mutant <- "F"
outcome_females <- outcome_females[,
    c(1:7, 10, 8:9)]
outcome_females$number_successes <- outcome$number_successes -
    male_successes
outcome_females$number_tries <- outcome$number_tries -
    male_first_mutant

# Bind together
outcome <- rbind(outcome_males, outcome_females)
```

Some aesthetics and save output as table.

```
# Order by value/letter
outcome <- outcome[order(outcome$sex_conflict,
    outcome$bottleneck, outcome$prob_encounter,
    outcome$predator_absence, outcome$strength_attract,
    outcome$pred_constant, outcome$number_loci,
    outcome$sex_first_mutant), ]

# Change 0 and 1 to N (no) and Y
# (yes)
outcome$sex_conflict <- ifelse(outcome$sex_conflict ==
    1, "Y", "N")
outcome$bottleneck <- ifelse(outcome$bottleneck ==
    1, "Y", "N")

# Save table
write.table(outcome, "Data/Simulations/summary_simulations.txt",
    row.names = F, quote = F)
```

#### 3.2 Which simulation parameters influence whether the novel pattern persists?

##### 3.2.1 Simple statistics

We make the categorical columns factors.

```
outcome$sex_conflict <- factor(outcome$sex_conflict)
outcome$bottleneck <- factor(outcome$bottleneck)
outcome$sex_first_mutant <- factor(outcome$sex_first_mutant)
```

Count how many of the **scenarios** (a certain parameter combination) led to survival of the novel pattern in at least one simulation run, *i.e.* as long as the pattern survived in one simulation run of a scenario, the scenario gets counted.

```
setting_success <- sapply(1:8, function(x) sapply(as.character(unique(outcome[,
    x])), function(y) sum(as.character(outcome[,
    x]) == y & outcome$number_successes >
    0), USE.NAMES = TRUE))
names(setting_success) <- names(outcome)[1:8]
print(setting_success)
```

```
## $sex_conflict
##  N  Y
##  0 61
##
## $bottleneck
##  N  Y
## 24 37
##
## $prob_encounter
##  0.001 0.0025  0.005   0.01
##      0      0      2     59
##
## $predator_absence
##  10 100
##   0  61
##
## $strength_attract
##  1  3
## 23 38
##
## $pred_constant
##  1  2  4
## 26 18 17
##
## $number_loci
##  1  5 10
## 20 21 20
##
## $sex_first_mutant
##  F  M
## 29 32
```

We repeat this, but this time we count the number of simulation run within these scenarios, where the allele survived.

```
number_success <- sapply(1:8, function(x) sapply(as.character(unique(outcome[,
    x])), function(y) sum(outcome$number_successes[as.character(outcome[,
    x]) == y]), USE.NAMES = TRUE))
names(number_success) <- names(outcome)[1:8]
print(number_success)
```

```
## $sex_conflict
##    N    Y
##    0 2865
##
## $bottleneck
```

```
##    N    Y
## 1996  869
##
## $prob_encounter
##  0.001 0.0025  0.005   0.01
##      0      0      2   2863
##
## $predator_absence
##   10  100
##    0 2865
##
## $strength_attract
##    1    3
##  289 2576
##
## $pred_constant
##    1    2    4
## 1512  974  379
##
## $number_loci
##   1   5  10
## 940 956 969
##
## $sex_first_mutant
##    F    M
## 1842 1023
```

Check which scenario had the highest proportion of success:

```r
outcome[which.max(outcome$number_successes/outcome$number_tries),
    ]
```

… and the exact proportion of successes:

```r
format(round(max(outcome$number_successes/outcome$number_tries),
    3), nsmall = 3)
```

```
## [1] "0.260"
```

Find the proportion of successes with sexual conflict present

```r
format(round(sum(outcome$number_successes[outcome$sex_conflict ==
    "Y"])/sum(outcome$number_tries[outcome$sex_conflict ==
    "Y"]), 3), nsmall = 3)
```

```
## [1] "0.005"
```

##### 3.2.2 Statistical tests

The overview of number of successes across different parameter settings before showed that we will run into some problems if we now just start doing a normal logistic regression and include all 8 fixed effects. Specifically, there is perfect separation in 3 variables, namely: sex_conflict, predator_absence and prob_encounter. Sexual conflict always has to be present and predators always have to return after 100 generations for the novel mutation to survive until generation 2500. Also, at least a prob_encounter value of 0.005 is necessary (and even at 0.005, we still have quasi-perfect separation).

Now, there is several possibilities to deal with this. * 1) Use packages such as brglm2, glmnet or logistf to fit GLMs including all 8 predictors, but accounting for the perfect separation. * 2) Go Bayesian and use the right kind of priors. * 3) Run a model without including sex_conflict and predator_absence, and dealing in some way with prob_encounter that still allows running the GLM (without the separation issues).

For further discussion, see also: https://stats.stackexchange.com/questions/11109/how-to-deal-with-perfect-separation-in-logistic-regression

We tried 1) and 2), which both gave better models than the usual glm, but still, confidence intervals for the estimators for all fixed effects were gigantic. Both even resulted in huge confidence intervals when discarding sex_conflict and predator_absence, but keeping prob_encounter as numeric covariate. Hence, we decided to use approach 3).

###### 3.2.2.1 Binomial tests

Before all that, we will use a simple binomial test to check how significant the effect of sex_conflict and predator_absence is. First, we produce a simple overview table showing the proportion each setting resulted in a survival until generation 2500.

```
conflict_return <- data.frame(sex_conflict = c("N",
    "N", "Y", "Y"), predator_absence = c(10,
    100, 10, 100), sum_survived = c(sum(outcome$number_successes[outcome$sex_conflict ==
    "N" & outcome$predator_absence ==
    10]), sum(outcome$number_successes[outcome$sex_conflict ==
    "N" & outcome$predator_absence ==
    100]), sum(outcome$number_successes[outcome$sex_conflict ==
    "Y" & outcome$predator_absence ==
    10]), sum(outcome$number_successes[outcome$sex_conflict ==
    "Y" & outcome$predator_absence ==
    100])), sum_tries = c(sum(outcome$number_tries[outcome$sex_conflict ==
    "N" & outcome$predator_absence ==
    10]), sum(outcome$number_tries[outcome$sex_conflict ==
    "N" & outcome$predator_absence ==
    100]), sum(outcome$number_tries[outcome$sex_conflict ==
    "Y" & outcome$predator_absence ==
    10]), sum(outcome$number_tries[outcome$sex_conflict ==
    "Y" & outcome$predator_absence ==
    100])))
conflict_return <- cbind(conflict_return,
    binom.confint(conflict_return$sum_survived,
        conflict_return$sum_tries, method = "exact")[,
        4:6])
print(data.frame(conflict_return[, 1:4],
    round(conflict_return[, 5:7], 4)))
```

```
##   sex_conflict predator_absence sum_survived sum_tries   mean  lower  upper
## 1            N               10            0     72000 0.0000 0.0000 0.0001
## 2            N              100            0     72000 0.0000 0.0000 0.0001
## 3            Y               10            0    288000 0.0000 0.0000 0.0000
## 4            Y              100         2865    288000 0.0099 0.0096 0.0103
```

Now, we test each combination other than conflict present + return after 100 versus this setting. We also correct for multiple testing with the Bonferroni method. We can see that the proportion under conflict present + return after 100 is significantly higher than in all other combinations (where the proportion is always exactly 0).

```
tests <- data.frame(test = c("0_10 vs 1_100",
    "0_100 vs 1_100", "1_10 vs 1_100"),
    p.value = c(prop.test(x = conflict_return$sum_survived[c(1,
        4)], n = conflict_return$sum_tries[c(1,
```

```
        4)])$p.value, prop.test(x = conflict_return$sum_survived[c(2,
        4)], n = conflict_return$sum_tries[c(2,
        4)])$p.value, prop.test(x = conflict_return$sum_survived[c(3,
        4)], n = conflict_return$sum_tries[c(3,
        4)])$p.value))
tests$p.value <- p.adjust(tests$p.value,
    method = "bonferroni")
tests$p.value <- format(round(tests$p.value,
    3), nsmall = 3)
print(tests)
```

```
##             test p.value
## 1  0_10 vs 1_100   0.000
## 2 0_100 vs 1_100   0.000
## 3  1_10 vs 1_100   0.000
```

**3.2.2.2 GLMs** In the GLMs, we use two 'tricks' to bypass the problem of complete separation:

First, we discard the data from cases without sexual conflict or/and only 10 generations of predator-free phase.

```
glm_data <- outcome[outcome$sex_conflict ==
    "Y" & outcome$predator_absence ==
    100, ]
```

Second, as we had such few different values for all numeric predictors, we decided to make them ordered factors (this also helps bypassing the problem with quasi-complete separation in probability of encounter).

```
glm_data$prob_encounter <- as.ordered(glm_data$prob_encounter)
glm_data$strength_attract <- as.ordered(glm_data$strength_attract)
glm_data$pred_constant <- as.ordered(glm_data$pred_constant)
glm_data$number_loci <- as.ordered(glm_data$number_loci)
```

Change the contrasts used for dummy coding (remember the old setting to reset later)

```
old_contr <- options("contrasts")
options(contrasts = c("contr.sum", "contr.poly"))
```

Create an intercept-only logistic regression. Then use forward stepwise AIC calculation to find which factors/covariates play a role explaining survival of the novel pattern allele. Note that we allow a maximum of two-way interactions, to not overcomplicate the model. We suppress warnings from the stepAIC function, which repeatedly warns the following "glm.fit: fitted probabilities numerically 0 or 1 occurred". This is because of some settings never leading to successes, which is not ideal for the model. However, the model still converges.

```
glm1 <- glm(cbind(number_successes,
    number_tries - number_successes) ~
    1, data = glm_data, family = "binomial")

step_glm <- suppressWarnings(stepAIC(glm1,
    scope = list(upper = ~(bottleneck +
        prob_encounter + strength_attract +
        pred_constant + number_loci +
        sex_first_mutant)^2, lower = ~1),
    direction = "forward", trace = FALSE))
```

Test for significance. Correct p-values with the Bonferroni method. All effects are highly significant.

```
p_values <- joint_tests(step_glm)
p_values$p.value <- p.adjust(p_values$p.value,
    method = "bonferroni")
print(p_values)
```

```
##  model term                       df1 df2 F.ratio p.value
##  prob_encounter                     3 Inf  36.570 <.0001
##  strength_attract                   1 Inf 569.593 <.0001
##  pred_constant                      2 Inf 160.863 <.0001
##  bottleneck                         1 Inf  31.493 <.0001
##  sex_first_mutant                   1 Inf 118.329 <.0001
##  strength_attract:pred_constant     2 Inf  56.039 <.0001
##  strength_attract:bottleneck        1 Inf 304.385 <.0001
##  pred_constant:bottleneck           2 Inf  58.521 <.0001
##  bottleneck:sex_first_mutant        1 Inf 220.306 <.0001
```

Now we calculate the R^2. We suppress again the usual warning about fitted probabilities.

```
dominance_fact <- suppressWarnings(averageContribution(dominanceAnalysis(step_glm),
    fit.functions = "r2.m")$r2.m)
print_r2 <- data.frame(fixed_effect = names(dominance_fact),
    R2 = round(unname(dominance_fact),
        3))
print(print_r2[order(print_r2$R2, decreasing = T),
    ])
```

```
##                      fixed_effect    R2
## 1                  prob_encounter 0.634
## 6    strength_attract:bottleneck 0.089
## 8 strength_attract:pred_constant 0.084
## 2                strength_attract 0.056
## 7    bottleneck:sex_first_mutant 0.035
## 9       pred_constant:bottleneck 0.034
## 3                   pred_constant 0.020
## 5                sex_first_mutant 0.010
## 4                      bottleneck 0.009
```

Now we calculated the estimated marginal means (EMMs) for all levels of each of the fixed effects. This is very complicated code, which basically does something very simple: It just calls the `emmeans` function and then just re-formats the output. Finally, for each fixed effect, we add the number of successes and trials, as well as p-value and the R^2 value. Note that `emmeans` returns the warning of 'NOTE: Results may be misleading due to involvement in interactions', which is very normal. It's just warning to not look at the EMMs of single terms, if those are also involved in interactions.

```
emms <- list()
# Fixed effect names
t_name <- levels(p_values$`model term`)[p_values$`model term`]
for (i in attr(step_glm$terms, "term.labels")) {
    # First we differentiate between
    # interaction and no-interaction
    # terms. First the terms without
    # interaction:
    if (!grepl(":", i)) {
        # Calculate EMMs (we have to solve
        # this with eval-parse, as we want
        # to call the term 'i', so we first
```

```
        # have to construct the call to
        # emmeans as a string)
        emcall <- paste0("summary(emmeans(step_glm,pairwise~",
            i, ",transform='response'))")
        emms[[length(emms) + 1]] <- eval(parse(text = emcall))
        # list length:
        l <- length(emms)
        # Now we have to differentiate
        # between factors and non-factors
        # (although, almost all terms should
        # be factors, but emmeans might
        # change them to numeric in its
        # output table). First if we have a
        # factor as fixed
        if (is.factor(emms[[l]][[1]][,
            1])) {
            # Count number successes
            emms[[l]][[1]]$successes <- sapply(levels(emms[[l]][[1]][,
                1])[emms[[l]][[1]][,
                1]], function(x) sum(glm_data$number_successes[glm_data[,
                names(emms[[l]][[1]])[1]] ==
                x]))
            # Count number tries
            emms[[l]][[1]]$tries <- sapply(levels(emms[[l]][[1]][,
                1])[emms[[l]][[1]][,
                1]], function(x) sum(glm_data$number_tries[glm_data[,
                names(emms[[l]][[1]])[1]] ==
                x]))
            # Almost same if fixed effect is not
            # a factor, just a bit easier
        } else {
            emms[[l]][[1]]$successes <- sapply(emms[[l]][[1]][,
                1], function(x) sum(glm_data$number_successes[glm_data[,
                names(emms[[l]][[1]])[1]] ==
                x]))
            emms[[l]][[1]]$tries <- sapply(emms[[l]][[1]][,
                1], function(x) sum(glm_data$number_tries[glm_data[,
                names(emms[[l]][[1]])[1]] ==
                x]))
        }
        # Now come all cases where the term
        # is an interaction We have to find
        # all possible combinations, and
        # thus the table formatting is a bit
        # more complex. But follows
        # basically the same rules as above,
        # so won't be explained in detail.
    } else {
        # Split by ':' sign (return two
        # strings, as we have max. two-way
        # interactions)
        splitter <- strsplit(i, ":")[[1]]
        # Calculate EMMs (we have to solve
```

```r
# this with eval-parse, as we want
# to call the two terms from
# 'splitter', so we first have to
# construct the call to emmeans as a
# string)
emcall <- paste0("summary(emmeans(step_glm,pairwise~",
    splitter[1], "|", splitter[2],
    ",transform='response'))")
emms[[length(emms) + 1]] <- eval(parse(text = emcall))
# list length:
l <- length(emms)

if (is.factor(emms[[l]][[1]][,
    1])) {
    if (is.factor(emms[[l]][[1]][,
        2])) {
        emms[[l]][[1]]$successes <- sapply(1:nrow(emms[[l]][[1]]),
          function(x) sum(glm_data$number_successes[glm_data[,
            names(emms[[l]][[1]])[1]] ==
            levels(emms[[l]][[1]][,
              1])[emms[[l]][[1]][x,
              1]] & glm_data[,
            names(emms[[l]][[1]])[2]] ==
            levels(emms[[l]][[1]][,
              2])[emms[[l]][[1]][x,
              2]]]))
        emms[[l]][[1]]$tries <- sapply(1:nrow(emms[[l]][[1]]),
          function(x) sum(glm_data$number_tries[glm_data[,
            names(emms[[l]][[1]])[1]] ==
            levels(emms[[l]][[1]][,
              1])[emms[[l]][[1]][x,
              1]] & glm_data[,
            names(emms[[l]][[1]])[2]] ==
            levels(emms[[l]][[1]][,
              2])[emms[[l]][[1]][x,
              2]]]))
    } else {
        emms[[l]][[1]]$successes <- sapply(1:nrow(emms[[l]][[1]]),
          function(x) sum(glm_data$number_successes[glm_data[,
            names(emms[[l]][[1]])[1]] ==
            levels(emms[[l]][[1]][,
              1])[emms[[l]][[1]][x,
              1]] & glm_data[,
            names(emms[[l]][[1]])[2]] ==
            emms[[l]][[1]][x,
              2]]))
        emms[[l]][[1]]$tries <- sapply(1:nrow(emms[[l]][[1]]),
          function(x) sum(glm_data$number_tries[glm_data[,
            names(emms[[l]][[1]])[1]] ==
            levels(emms[[l]][[1]][,
              1])[emms[[l]][[1]][x,
              1]] & glm_data[,
            names(emms[[l]][[1]])[2]] ==
```

```
                    emms[[l]][[1]][x,
                      2]]))
            }
        } else {
            if (is.factor(emms[[l]][[1]][,
                2])) {
                emms[[l]][[1]]$successes <- sapply(1:nrow(emms[[l]][[1]]),
                  function(x) sum(glm_data$number_successes[glm_data[,
                    names(emms[[l]][[1]])[1]] ==
                    emms[[l]][[1]][x,
                      1] & glm_data[,
                    names(emms[[l]][[1]])[2]] ==
                    levels(emms[[l]][[1]][,
                      2])[emms[[l]][[1]][x,
                      2]]]))
                emms[[l]][[1]]$tries <- sapply(1:nrow(emms[[l]][[1]]),
                  function(x) sum(glm_data$number_tries[glm_data[,
                    names(emms[[l]][[1]])[1]] ==
                    emms[[l]][[1]][x,
                      1] & glm_data[,
                    names(emms[[l]][[1]])[2]] ==
                    levels(emms[[l]][[1]][,
                      2])[emms[[l]][[1]][x,
                      2]]]))
            } else {
                emms[[l]][[1]]$successes <- sapply(1:nrow(emms[[l]][[1]]),
                  function(x) sum(glm_data$number_successes[glm_data[,
                    names(emms[[l]][[1]])[1]] ==
                    emms[[l]][[1]][x,
                      1] & glm_data[,
                    names(emms[[l]][[1]])[2]] ==
                    emms[[l]][[1]][x,
                      2]]))
                emms[[l]][[1]]$tries <- sapply(1:nrow(emms[[l]][[1]]),
                  function(x) sum(glm_data$number_tries[glm_data[,
                    names(emms[[l]][[1]])[1]] ==
                    emms[[l]][[1]][x,
                      1] & glm_data[,
                    names(emms[[l]][[1]])[2]] ==
                    emms[[l]][[1]][x,
                      2]]))
            }
        }
    }
    # Add R2 and p-value
    emms[[l]][[3]] <- unname(dominance_fact[names(dominance_fact) ==
        i])
    emms[[l]][[4]] <- ifelse(p_values$p.value[t_name ==
        i] < 0.001, "< 0.001", paste0("= ",
        as.character(unname(format(round(p_values$p.value[t_name ==
            i], 3), nsmall = 3)))))
    names(emms[[l]])[3:4] <- c("r2.m",
        "p.value")
```

```
}
```

```
## NOTE: Results may be misleading due to involvement in interactions
## NOTE: Results may be misleading due to involvement in interactions
## NOTE: Results may be misleading due to involvement in interactions
## NOTE: Results may be misleading due to involvement in interactions
```

```
# Give list the names following the
# terms
names(emms) <- attr(step_glm$terms,
    "term.labels")

# reorder based on R2
emms <- emms[order(sapply(1:length(emms),
    function(x) emms[[x]]$r2.m), decreasing = T)]
```

##### 3.2.3 Plots

**3.2.3.1 Effect of different parameters** Now we plot the results from the binomial tests/GLM. Comments within the following code should hopefully be sufficient to explain what's happening here.

First, we define some vectors to rename the variable names:

```
new_titles <- c("bquote(italic(e)",
    "bquote(alpha~':'~bottleneck", "bquote(alpha~':'~italic(Q)",
    "bquote(alpha", "bquote(bottleneck~':'~sex_mutant",
    "bquote(italic(Q)~':'~bottleneck",
    "bquote(italic(Q)", "bquote(sex_mutant",
    "bquote(bottleneck")
```

Now the plot:

```
png("Figures/simulations_effect_parameters.png",
    width = 2700, height = 1500, res = 300)
# Open plot grid
layout(cbind(rep(1, 3), matrix(2:(length(emms) +
    1), ncol = ceiling(length(emms)/3),
    byrow = T)), widths = c(0.8, 1,
    1, 1))
par(oma = c(1.4, 0, 0, 0))
par(mar = c(1.8, 3.1, 1.3, 1))

# Binomial tests for the effect of
# sexual conflict and return of
# predators
plot(1, 1, type = "n", xlab = "", ylab = "",
    xaxt = "n", yaxt = "n", xlim = c(min(conflict_return$lower),
        max(conflict_return$upper)),
    ylim = c(0.5, nrow(conflict_return) +
        0.5), yaxs = "i")
# Vertical lines highlighting x-axis
# values
abline(v = seq(0, 1, 0.003), lty = "dashed",
    lwd = 0.5)

# Confidence intervals
```

```r
segments(conflict_return$lower, 1:nrow(conflict_return),
    conflict_return$upper, 1:nrow(conflict_return),
    lend = 3, lwd = 12, col = "lightblue")
# Estimators
points(conflict_return$mean, 1:nrow(conflict_return),
    pch = 20, cex = 1.3)

# Text for successes / tries
text_to_add <- paste0(conflict_return$sum_survived,
    " / ", conflict_return$sum_tries)
widths <- strwidth(text_to_add, cex = 0.6)
heights <- strheight(text_to_add, cex = 0.6)

# Add number of cases either left or
# right
left_right <- ifelse(conflict_return$upper <=
    (par("usr")[2] - 1.1 * widths -
        0.05 * diff(par("usr")[1:2])),
    1, -1)
add_at <- ifelse(left_right == 1, conflict_return$upper +
    0.03 * diff(par("usr")[1:2]), conflict_return$lower -
    0.03 * diff(par("usr")[1:2]))

# Add successes / tries to graph,
# with a rectangle around it
rect(add_at, (1:nrow(conflict_return)) -
    0.7 * heights, add_at + left_right *
    1.1 * widths, (1:nrow(conflict_return)) +
    0.7 * heights, col = "white", border = "black",
    lwd = 0.3)

for (textor in 1:length(text_to_add)) {
    if (left_right[textor] == 1) {
        text(add_at[textor] + 0.05 *
            widths[textor], (1:nrow(conflict_return))[textor],
            text_to_add[textor], pos = 4,
            offset = 0, cex = 0.6)
    } else {
        text(add_at[textor] - 0.05 *
            widths[textor], (1:nrow(conflict_return))[textor],
            text_to_add[textor], pos = 2,
            offset = 0, cex = 0.6)
    }
}

# Reformat p-values (if smaller than
# <0.001)
p_vals <- ifelse(tests$p.value < 0.001,
    "bquote(italic(p)~'<'~0.001)", paste0("bquote(italic(p)~'='",
        tests$p.value, ")"))

# Width and height of text
widths_p <- sapply(1:length(p_vals),
```

```
    function(x) eval(parse(text = paste0("strwidth(",
        p_vals[x], ",cex=0.8)"))))
heights_p <- sapply(1:length(p_vals),
    function(x) eval(parse(text = paste0("strheight(",
        p_vals[x], ",cex=0.8)"))))

# Add arrows showing which
# comparison is looked at
arrows(add_at[4] + 0.5 * left_right[4] *
    1.1 * widths[4], nrow(conflict_return) -
    0.7 * heights[4], add_at[4] + 0.5 *
    left_right[4] * 1.1 * widths[4],
    1, lwd = 1.4, lend = 3, code = 1,
    length = 0.05)
arrows((add_at + left_right * 1.1 *
    widths)[-4], 1:(nrow(conflict_return) -
    1), add_at[4] + 0.5 * left_right[4] *
    1.1 * widths[4], 1:(nrow(conflict_return) -
    1), lwd = 1.4, lend = 3, code = 1,
    length = 0.05)

# Add p-values with rectangle around
rect(add_at[4] + 0.5 * left_right[4] *
    1.1 * widths[4] - 0.03 * diff(par("usr")[1:2]),
    (1:(nrow(conflict_return) - 1)) +
        0.01 * diff(par("usr")[3:4]),
    add_at[4] + 0.5 * left_right[4] *
        1.1 * widths[4] - 0.03 * diff(par("usr")[1:2]) -
        1.1 * widths_p, (1:(nrow(conflict_return) -
        1)) + 0.01 * diff(par("usr")[3:4]) +
        1.4 * heights_p, col = "white",
    border = "black", lwd = 0.3)
addtxt1 <- "text(add_at[4]+0.5*left_right[4]*1.1*widths[4]-0.03*diff(
     par('usr')[1:2])-1.05*widths_p,"
addtxt2 <- "(1:(nrow(conflict_return)-1))+0.01*diff(par('usr')[3:4])+0.7*heights_p,"
eval(parse(text = paste0(addtxt1, addtxt2,
    p_vals, ",pos = 4,offset = 0,cex=0.8)")))

# Axis titles and axes
mtext("sex_conflict : predator_absence",
    3, line = 0.2, cex = 0.54)
mtext("Proportion mutation surviving",
    1, line = 2, cex = 0.7)
axis(1, labels = F, tck = -0.02, at = seq(0,
    0.009, 0.003))
axis(1, lwd = 0, line = -0.7, cex.axis = 0.9,
    at = seq(0, 0.009, 0.003))
axis(2, at = 1:nrow(conflict_return),
    labels = F, tck = -0.02)
axis(2, at = 1:nrow(conflict_return),
    paste0(conflict_return$sex_conflict,
        " : ", conflict_return$predator_absence),
    lwd = 0, las = 2, line = -0.6, cex.axis = 0.9)
```

```
# Add divider between levels of
# sexual conflict
abline(h = 2.5, lty = "dashed", lwd = 0.5)

# Add an 'A' to this plot region
par(xpd = NA)
text(par("usr")[1] - (0.18/par("pin")[1]) *
    diff(par("usr")[1:2]), par("usr")[4] +
    (0.04/par("pin")[2]) * diff(par("usr")[3:4]),
    "A", font = 2, cex = 1.5, pos = 2,
    offset = 0)
par(xpd = F)

# GLM results for all other effects

par(mar = c(1.8, 3.1, 1.3, 0.5))

# Get the lowest value of the lower
# CIs and the highest value of the
# upper CIs (to later sclae the
# x-axes of all graphs to)
min_y <- min(sapply(1:length(emms),
    function(x) min(emms[[x]]$emmeans$asymp.LCL)))
max_y <- max(sapply(1:length(emms),
    function(x) max(emms[[x]]$emmeans$asymp.UCL)))

# Go one by one through the EMMs,
# starting with the one with the
# biggest R2 value
for (i in 1:length(emms)) {
    # Open empty plot
    plot(1, 1, type = "n", xlab = "",
        ylab = "", xaxt = "n", yaxt = "n",
        xlim = c(min_y, max_y), ylim = c(0.5,
            nrow(emms[[i]]$emmeans) +
                0.5), yaxs = "i")
    # Provide vertical lines at all
    # x-values, with 0.005 intervals
    abline(v = seq(0, 1, 0.005), lty = "dashed",
        lwd = 0.5)
    # Add CIs
    segments(emms[[i]]$emmeans$asymp.LCL,
        1:nrow(emms[[i]]$emmeans), emms[[i]]$emmeans$asymp.UCL,
        1:nrow(emms[[i]]$emmeans), lend = 3,
        lwd = 12, col = "lightblue")
    # Add EMMs
    points(emms[[i]]$emmeans$prob, 1:nrow(emms[[i]]$emmeans),
        pch = 20, cex = 1.3)

    # Text for successes / tries
    text_to_add <- paste0(emms[[i]]$emmeans$successes,
        " / ", emms[[i]]$emmeans$tries)
    widths <- strwidth(text_to_add,
```

```
    cex = 0.6)
heights <- strheight(text_to_add,
    cex = 0.6)

# Add number of cases either left or
# right
left_right <- ifelse(emms[[i]]$emmeans$asymp.UCL <=
    (par("usr")[2] - 1.1 * widths -
        0.05 * diff(par("usr")[1:2])),
    1, -1)
add_at <- ifelse(left_right == 1,
    emms[[i]]$emmeans$asymp.UCL +
        0.03 * diff(par("usr")[1:2]),
    emms[[i]]$emmeans$asymp.LCL -
        0.03 * diff(par("usr")[1:2]))

# Add successes / tries with a
# rectangle around the text
rect(add_at, (1:nrow(emms[[i]]$emmeans)) -
    0.7 * heights, add_at + left_right *
    1.1 * widths, (1:nrow(emms[[i]]$emmeans)) +
    0.7 * heights, col = "white",
    border = "black", lwd = 0.3)

for (textor in 1:length(text_to_add)) {
    if (left_right[textor] == 1) {
        text(add_at[textor] + 0.05 *
            widths[textor], (1:nrow(emms[[i]]$emmeans))[textor],
            text_to_add[textor],
            pos = 4, offset = 0,
            cex = 0.6)
    } else {
        text(add_at[textor] - 0.05 *
            widths[textor], (1:nrow(emms[[i]]$emmeans))[textor],
            text_to_add[textor],
            pos = 2, offset = 0,
            cex = 0.6)
    }
}

# Above each graph, add the name of
# the fixed effect, the R2 and the
# p-value
par(xpd = NA)
btxt1 <- " ~ '  ('*italic(R)^2 ~ '=' ~ .(format(round(emms[[i]]$r2.m,3),"
btxt2 <- "nsmall=3))*', '*italic(p) ~ .(emms[[i]]$p.value) *')'"
btxt3 <- "),3,line=0.1,cex=0.54)"
eval(parse(text = paste0("mtext(",
    new_titles[i], btxt1, btxt2,
    btxt3)))
par(xpd = F)
# Add axes
axis(1, labels = F, tck = -0.02)
```

```r
    axis(1, lwd = 0, line = -0.7, cex.axis = 0.9)
    axis(2, at = 1:nrow(emms[[i]]$emmeans),
        labels = F, tck = -0.02)
    # If interaction term is present,
    # the y-axis will look differently
    if (grepl(":", names(emms)[[i]])) {
        axis(2, at = 1:nrow(emms[[i]]$emmeans),
            paste0(levels(emms[[i]]$emmeans[,
                1])[emms[[i]]$emmeans[,
                1]], " : ", levels(emms[[i]]$emmeans[,
                2])[emms[[i]]$emmeans[,
                2]]), lwd = 0, las = 2,
            line = -0.6, cex.axis = 0.9)
        uni1 <- length(unique(emms[[i]]$emmeans[,
            1]))
        uni2 <- length(unique(emms[[i]]$emmeans[,
            2]))
        abline(h = seq(uni1, (uni1 *
            uni2) - uni1, uni1) + 0.5,
            lty = "dashed", lwd = 0.5)
    } else {
        axis(2, at = 1:nrow(emms[[i]]$emmeans),
            levels(emms[[i]]$emmeans[,
                1])[emms[[i]]$emmeans[,
                1]], lwd = 0, las = 2,
            line = -0.6, cex.axis = 0.9)
    }

    # Add x-axis and the letter 'B'
    if (i == 8) {
        # Splitting this so it doesn't
        # exceed line in Markdown
        txt1 <- "Proportion mutation surviving"
        txt2 <- " (under sexual conflict and predator absence for 100 generations)"
        mtext(paste0(txt1, txt2), 1,
            line = 2, cex = 0.7)
    }
    if (i == 1) {
        par(xpd = T)
        text(par("usr")[1] - (0.18/par("pin")[1]) *
            diff(par("usr")[1:2]), par("usr")[4] +
            (0.04/par("pin")[2]) * diff(par("usr")[3:4]),
            "B", font = 2, cex = 1.5,
            pos = 2, offset = 0)
        par(xpd = F)
    }
}
invisible(dev.off())
```

Show png we just made:

```r
fig_disp <- readPNG("Figures/simulations_effect_parameters.png")
grid.raster(fig_disp)
```

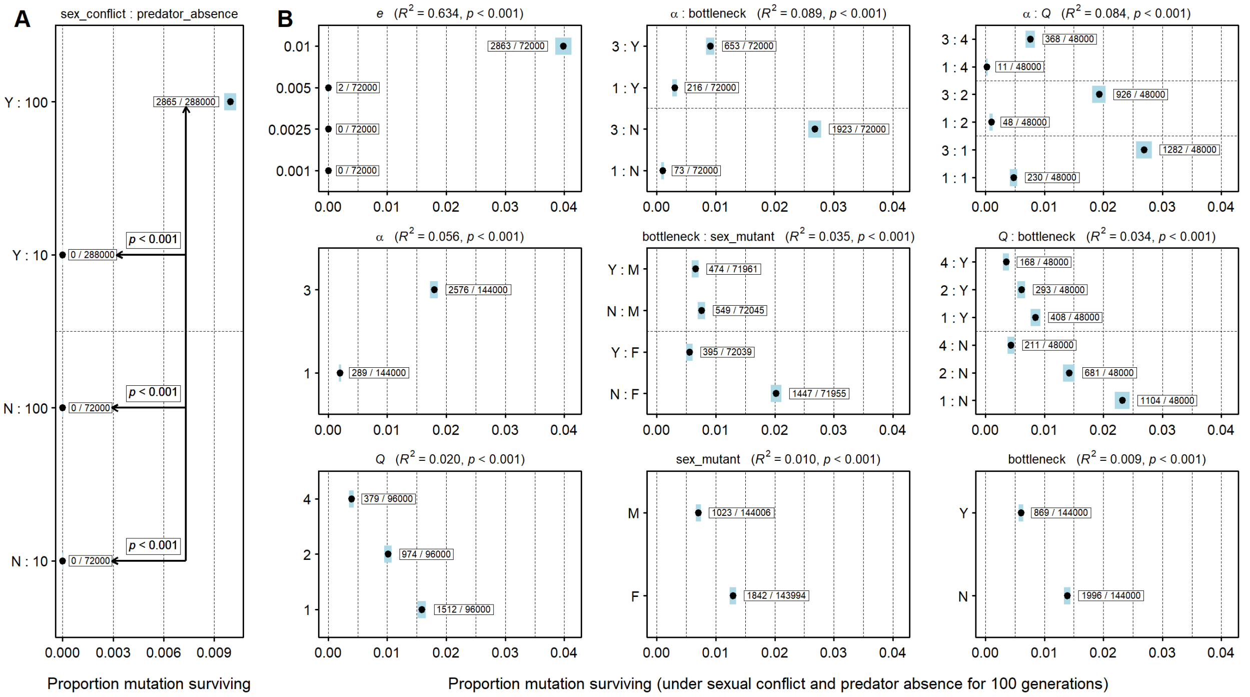


**3.2.3.2 'Heatmap table' for all parameter combinations** First, we want to calculate the selection coefficients, dependent on the parameters */alpha* (strength of male attraction) and *e* (probability of encountering a single male). Note that these calculations are done for the starting population in the patch the first mutant occurs, *i.e.* where predators are absent and preference phenotypes ancestral; the calculation is done under the assumption that the first mutant is a female (if it's a male, it should have no effect). By 'selection coefficient', we mean the proportional increase in offspring that the mutant generates. E.g. mutant lays 12 eggs, non-mutant lays 10 eggs, then the selection coefficient is 0.2 (20% more eggs). Note, the coefficient can only be positive here, as predators are absent at this moment and there is no associated cost.

The starting population in all our simulations is always created as follows:

```
number_per_patch <- 114
starting_pop <- data.frame(sex = (rep(c(0L,
    1), ceiling(number_per_patch/2)))[1:number_per_patch],
    age = c(rep(0L:5L, each = floor(number_per_patch/6)),
        rep(0L, number_per_patch%%6)))
```

This is the number of males actually alive during the first generation. In the starting population, a few individuals will be age = 5, which die immediately in the beginning of the cycle. Therefore, not all 114 individuals will make it until the part of the cycnle where offspring is being produced (and male harassment takes place). The number might be 1 higher in some simulation runs, as the mutant is chosen at random from the 114 individuals and then immediately becomes age 0. This shouldn't matter much and just affect the selection coefficient very slightly.

```
(sum_males <- sum(starting_pop$sex ==
    1 & starting_pop$age < 5))
```

```
## [1] 47
```

These are the possible values of /*alpha* and *e*:

```
values_e <- c(0.001, 0.0025, 0.005,
    0.01)
values_a <- c(1, 3)
```

Calculate selection coefficients for all *e*, first looking at /*alpha*=1, then /*alpha*=3. Display as nicely formatted table, finally save as vector (for plotting later).

```
sel_coef <- sapply(values_e, function(x) sapply(values_a,
    function(y) calc_s(chance_encounter = x,
        alpha = y, number_indiv = sum_males,
        p1 = 0, p2 = 1)))
colnames(sel_coef) <- paste0("e = ",
    values_e)
rownames(sel_coef) <- paste0("a = ",
    values_a)
(sel_coef <- format(round(sel_coef,
    3), nsmall = 3))
```

```
##       e = 0.001 e = 0.0025 e = 0.005 e = 0.01
## a = 1 "0.029"   "0.072"    "0.141"   "0.272"
## a = 3 "0.043"   "0.104"    "0.198"   "0.360"
```

```
sel_coef <- as.vector(sel_coef)
```

Now, we define the order in which the different confidence intervals should appear in the 'heatmap table'. Note that here we only include simulation scenarios where `sex_conflict=="Y"` and `predator_absence==100`. In total, those are 288 scenarios.

```
table_order <- as.vector(sapply(c("F",
    "M"), function(c1) sapply(c(1, 5,
    10), function(c2) sapply(c("Y",
    "N"), function(c3) sapply(c(1, 2,
    4), function(c4) sapply(c(0.001,
    0.0025, 0.005, 0.01), function(c5) sapply(c(1,
    3), function(c6) (1:nrow(outcome))[outcome$sex_conflict ==
    "Y" & outcome$predator_absence ==
    100 & outcome$sex_first_mutant ==
    c1 & outcome$number_loci == c2 &
    outcome$bottleneck == c3 & outcome$pred_constant ==
    c4 & outcome$prob_encounter == c5 &
    outcome$strength_attract == c6])))))))
```

For all scenarios, we calculate a binomial confidence interval (for display, we then format it by transforming it into % and rounding it).

```
bincoef <- as.data.frame(t(matrix(unlist(sapply(table_order,
    function(x) binom.confint(outcome$number_successes[x],
        outcome$number_tries[x], methods = "exact")[4:6])),
    nrow = 3, ncol = length(table_order))))
bincoef_pretty <- paste0(round(bincoef[,
    1] * 100, 1), "%", " [", round(bincoef[,
    2] * 100, 1), "-", round(bincoef[,
    3] * 100, 1), "%]")
```

Now we plot the actual 'heatmap table':

```r
png("Figures/simulations_heatmap.png",
    width = 4000, height = 6000, res = 300)

par(mar = rep(0.15, 4))
plot(1, 1, type = "n", xlab = "", ylab = "",
    xaxt = "n", yaxt = "n", xaxs = "i",
    yaxs = "i", xlim = c(1, 11), ylim = c(39,
        0), bty = "n")

# Draw heat colours (the more red,
# the more the allele survived
# (maximum 75% intensity))
for (i in 1:length(bincoef_pretty)) {
    rect(xleft = 2 + (i - 1)%%8 + 1,
        ybottom = 4 + (i - 1)%/%8, xright = 3 +
            (i - 1)%%8 + 1, ytop = 3 +
            (i - 1)%/%8, border = NA,
        col = adjustcolor("red", (bincoef[i,
            1]/max(bincoef[, 1])) *
            0.75))
}

# Now draw the grid
par(xpd = T)
segments(c(3:11), 0, c(3:11), 39)
segments(3, c(0:1, 3:39), 11, c(0:1,
    3:39))
segments(3, 2, 11, 2, lwd = 2, lty = "dashed")
segments(3, c(3, 39), 11, c(3, 39),
    lwd = 3)
segments(c(1, 1.5, 2, 2.5), 3, c(1,
    1.5, 2, 2.5), 39)
segments(2.5, 3:39, 3, 3:39)
segments(2, seq(3, 39, 3), 11, seq(3,
    39, 3), lwd = 1.8)
segments(1.5, seq(3, 39, 6), 11, seq(3,
    39, 6), lwd = 2.2)
segments(1, seq(3, 39, 18), 11, seq(3,
    39, 18), lwd = 2.2)
segments(c(3, 11), 3, c(3, 11), 39,
    lwd = 3)
par(xpd = F)

# Add the proportions and CIs
for (i in 1:length(bincoef_pretty)) {
    if (bincoef[i, 1] > 0) {
        text(2.5 + (i - 1)%%8 + 1, 3.5 +
            (i - 1)%/%8, bincoef_pretty[i],
            cex = 0.8)
    } else {
        text(2.5 + (i - 1)%%8 + 1, 3.5 +
            (i - 1)%/%8, "-", cex = 1)
    }
```

```r
}

# Add the different parameter values
# on top
text(c(3, 4) + 0.5, 0.5, substitute(paste(italic("e"),
    " = 0.001")))
text(c(5, 6) + 0.5, 0.5, substitute(paste(italic("e"),
    " = 0.0025")))
text(c(7, 8) + 0.5, 0.5, substitute(paste(italic("e"),
    " = 0.005")))
text(c(9, 10) + 0.5, 0.5, substitute(paste(italic("e"),
    " = 0.01")))
text(seq(3, 9, 2) + 0.5, 1.5, expression(paste(italic(alpha),
    " = 1")))
text(seq(4, 10, 2) + 0.5, 1.5, expression(paste(italic(alpha),
    " = 3")))
# Splitting for Markdown aesthetics
txt_s <- "substitute(paste(italic(Delta),' = ',"
invisible(sapply(1:length(sel_coef),
    function(x) text(x + 2.5, 2.5, eval(parse(text = paste0(txt_s,
        sel_coef[x], "))"))))))

# Add the parameter values on the
# left
text(2.68, 3:38 + 0.5, substitute(paste(italic("Q"))),
    srt = 90)
text(2.82, 3:38 + 0.5, paste0("= ",
    rep(c(1, 2, 4), 12)), srt = 90)
text(2.25, seq(4, 37, 3) + 0.5, c("Bottleneck",
    "No Bottleneck"), srt = 90)
text(1.75, seq(6, 36, 6), paste0("Loci = ",
    c(1, 5, 10)), srt = 90)
text(1.25, c(12, 30), c("First Mutant = Female",
    "First Mutant = Male"), srt = 90)

invisible(dev.off())
```

Show png we just made:

```r
fig_disp <- readPNG("Figures/simulations_heatmap.png")
grid.raster(fig_disp)
```

| | | | | $e$ = 0.001 | $e$ = 0.001 | $e$ = 0.0025 | $e$ = 0.0025 | $e$ = 0.005 | $e$ = 0.005 | $e$ = 0.01 | $e$ = 0.01 |
|---|---|---|---|---|---|---|---|---|---|---|---|
| | | | | α = 1 | α = 3 | α = 1 | α = 3 | α = 1 | α = 3 | α = 1 | α = 3 |
| | | | | Δ = 0.029 | Δ = 0.043 | Δ = 0.072 | Δ = 0.104 | Δ = 0.141 | Δ = 0.198 | Δ = 0.272 | Δ = 0.36 |
| First Mutant = Female | Loci = 1 | Bottleneck | Q = 1 | - | - | - | - | - | - | 2.4% [1.5-3.5%] | 3.8% [2.7-5.2%] |
| | | | Q = 2 | - | - | - | - | - | - | 0.9% [0.4-1.7%] | 3.7% [2.6-5%] |
| | | | Q = 4 | - | - | - | - | - | - | - | 2.4% [1.5-3.6%] |
| | | No Bottleneck | Q = 1 | - | - | - | - | - | - | 2% [1.2-3.1%] | 23.9% [21.3-26.7%] |
| | | | Q = 2 | - | - | - | - | - | - | - | 16.2% [14-18.6%] |
| | | | Q = 4 | - | - | - | - | - | - | - | 6% [4.6-7.6%] |
| | Loci = 5 | Bottleneck | Q = 1 | - | - | - | - | - | - | 2.3% [1.5-3.4%] | 3.5% [2.5-4.9%] |
| | | | Q = 2 | - | - | - | - | - | - | 1.2% [0.6-2.1%] | 3.9% [2.8-5.3%] |
| | | | Q = 4 | - | - | - | - | - | - | 0.2% [0-0.7%] | 2.3% [1.5-3.4%] |
| | | No Bottleneck | Q = 1 | - | - | - | - | - | - | 1.2% [0.6-2.1%] | 23.9% [21.3-26.7%] |
| | | | Q = 2 | - | - | - | - | - | - | - | 18.9% [16.5-21.4%] |
| | | | Q = 4 | - | - | - | - | - | - | - | 5.1% [3.8-6.6%] |
| | Loci = 10 | Bottleneck | Q = 1 | - | - | - | - | - | - | 1.9% [1.1-2.9%] | 4.2% [3.1-5.7%] |
| | | | Q = 2 | - | - | - | - | - | - | 0.4% [0.1-1%] | 3.8% [2.7-5.2%] |
| | | | Q = 4 | - | - | - | - | - | - | 0.2% [0-0.7%] | 2.7% [1.8-3.9%] |
| | | No Bottleneck | Q = 1 | - | - | - | - | - | - | 2.3% [1.5-3.4%] | 26% [23.3-28.9%] |
| | | | Q = 2 | - | - | - | - | - | - | - | 15.8% [13.6-18.2%] |
| | | | Q = 4 | - | - | - | - | - | - | - | 4.3% [3.1-5.7%] |
| First Mutant = Male | Loci = 1 | Bottleneck | Q = 1 | - | - | - | - | - | 0.1% [0-0.6%] | 2.9% [2-4.2%] | 4.6% [3.4-6%] |
| | | | Q = 2 | - | - | - | - | - | - | 1.2% [0.6-2.1%] | 3.4% [2.4-4.7%] |
| | | | Q = 4 | - | - | - | - | - | - | 0.2% [0-0.7%] | 3% [2-4.2%] |
| | | No Bottleneck | Q = 1 | - | - | - | - | - | - | 0.9% [0.4-1.7%] | 9.8% [8-11.8%] |
| | | | Q = 2 | - | - | - | - | - | - | - | 4.9% [3.7-6.5%] |
| | | | Q = 4 | - | - | - | - | - | - | - | 1.8% [1.1-2.9%] |
| | Loci = 5 | Bottleneck | Q = 1 | - | - | - | - | - | 0.1% [0-0.6%] | 2.7% [1.8-3.9%] | 4.3% [3.1-5.7%] |
| | | | Q = 2 | - | - | - | - | - | - | 0.4% [0.1-1%] | 4.9% [3.7-6.4%] |
| | | | Q = 4 | - | - | - | - | - | - | 0.4% [0.1-1%] | 2.1% [1.3-3.2%] |
| | | No Bottleneck | Q = 1 | - | - | - | - | - | - | 0.4% [0.1-1%] | 9.8% [8-11.8%] |
| | | | Q = 2 | - | - | - | - | - | - | - | 5.9% [4.5-7.6%] |
| | | | Q = 4 | - | - | - | - | - | - | - | 2.3% [1.4-3.4%] |
| | Loci = 10 | Bottleneck | Q = 1 | - | - | - | - | - | - | 3.5% [2.5-4.9%] | 4.5% [3.3-5.9%] |
| | | | Q = 2 | - | - | - | - | - | - | 0.7% [0.3-1.4%] | 4.8% [3.6-6.3%] |
| | | | Q = 4 | - | - | - | - | - | - | 0.1% [0-0.6%] | 3.2% [2.2-4.5%] |
| | | No Bottleneck | Q = 1 | - | - | - | - | - | - | 0.5% [0.2-1.2%] | 10.6% [8.8-12.6%] |
| | | | Q = 2 | - | - | - | - | - | - | - | 6.1% [4.7-7.8%] |
| | | | Q = 4 | - | - | - | - | - | - | - | 1.4% [0.8-2.4%] |

###### 3.2.3.3 Distribution of last generations and geographic position of first mutants

Extract the last generations

```
last_gen_l1 <- c(sapply(1:(nrow(l1) -
    1), function(x) ifelse(l1$Simulation[x +
    1] != l1$Simulation[x], x, NA)),
    nrow(l1))
last_gen_l1 <- last_gen_l1[!is.na(last_gen_l1)]
last_gen_l5 <- c(sapply(1:(nrow(l5) -
    1), function(x) ifelse(l5$Simulation[x +
    1] != l5$Simulation[x], x, NA)),
    nrow(l5))
last_gen_l5 <- last_gen_l5[!is.na(last_gen_l5)]
last_gen_l10 <- c(sapply(1:(nrow(l10) -
    1), function(x) ifelse(l10$Simulation[x +
    1] != l10$Simulation[x], x, NA)),
    nrow(l10))
last_gen_l10 <- last_gen_l10[!is.na(last_gen_l10)]
```

Plot distribution of last generations

```
png("Figures/simulations_distribution_last_gen.png",
    width = 2000, height = 1500, res = 300)
par(mar = c(4.1, 4, 3.5, 1))
suppressWarnings(plot(hist(c(l1$Generation[last_gen_l1],
    l5$Generation[last_gen_l5], l10$Generation[last_gen_l10]),
    breaks = seq(0, 2500, 1), plot = F)$count,
```

```
    log = "xy", lend = 2, type = "h",
    lwd = 3, xlab = "log Last Generation",
    ylab = "log Frequency", xaxt = "n"))
axis(1, c(1, 5, 10, 50, 100, 500, 1000,
    2500))
# Add important events
axis(3, at = c(6, 11, 101, 2500), las = 2,
    c("1st dies", "Pred. 1", "Pred. 2",
        "Max"), line = -0.7, lwd = 0)
invisible(dev.off())
```

Show png we just made:

```
fig_disp <- readPNG("Figures/simulations_distribution_last_gen.png")
grid.raster(fig_disp)
```

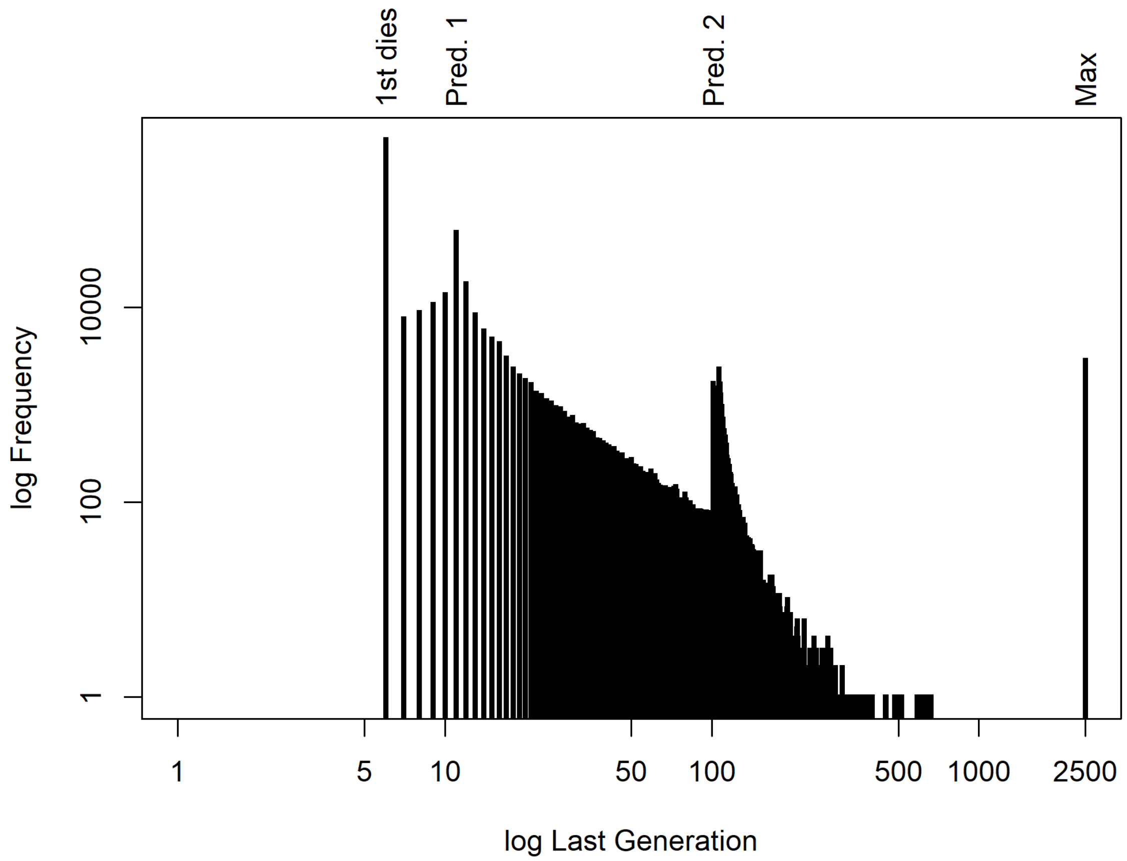


subset l1, l5 and l10 to only those rows where 2500 was reached

```
l1_2500 <- l1[l1$Generation == 2500,
    ]
l5_2500 <- l5[l5$Generation == 2500,
    ]
l10_2500 <- l10[l10$Generation == 2500,
    ]
```

Now we check in which areas of the grid the mutation happened among the cases where the mutation survived 2500 generations. First, calculate for all central patches how many times the first mutant occurred in them.

```
freq_per_patch <- sapply(c(39:58, 135:154,
    231:250, 327:346), function(x) sum(c(l1_2500$mutator_patch,
    l5_2500$mutator_patch, l10_2500$mutator_patch) ==
    x))
max_freq <- max(freq_per_patch)
```

First, we plot this as a 'map', where rectangles with darker colour indicate that first mutants placed in that region are more likely to lead to fixation of the novel allele.

```
png("Figures/simulations_first_mutant_location.png",
    width = 2000, height = 1500, res = 300)
par(mar = c(0, 0, 0, 3))
plot(1, 1, type = "n", xaxt = "n", yaxt = "n",
    xlab = "", ylab = "", xaxs = "i",
    yaxs = "i", xlim = c(0, 4), ylim = c(-5,
        25), bty = "n")
abline(h = 0:20)
segments(0:4, 0, 0:4, 20)

# Add the rectangles and colour by
# frequency of a mutation in that
# patch leading to fixation
invisible(sapply(1:80, function(x) rect(xleft = (0:3)[((x -
    1)%/%20) + 1 == (1:4)], xright = (1:4)[((x -
    1)%/%20) + 1 == (1:4)], ybottom = (19:0)[((x -
    1)%%20) + 1 == (1:20)], ytop = (20:1)[((x -
    1)%%20) + 1 == (1:20)], col = adjustcolor("black",
    freq_per_patch[x]/max_freq))))

text(2, 22, "Predation")
text(2, -2, "Predation")
arrows(2, c(23, -3), 2, c(24.5, -4.5),
    length = 0.1)
par(xpd = NA)
segments(4, c(0, 20), 4.2, c(9.5, 10.5))
text(4.3, 10, "Relaxed Predation", srt = 270)
par(xpd = F)

invisible(dev.off())
```

Show png we just made:

```
fig_disp <- readPNG("Figures/simulations_first_mutant_location.png")
grid.raster(fig_disp)
```

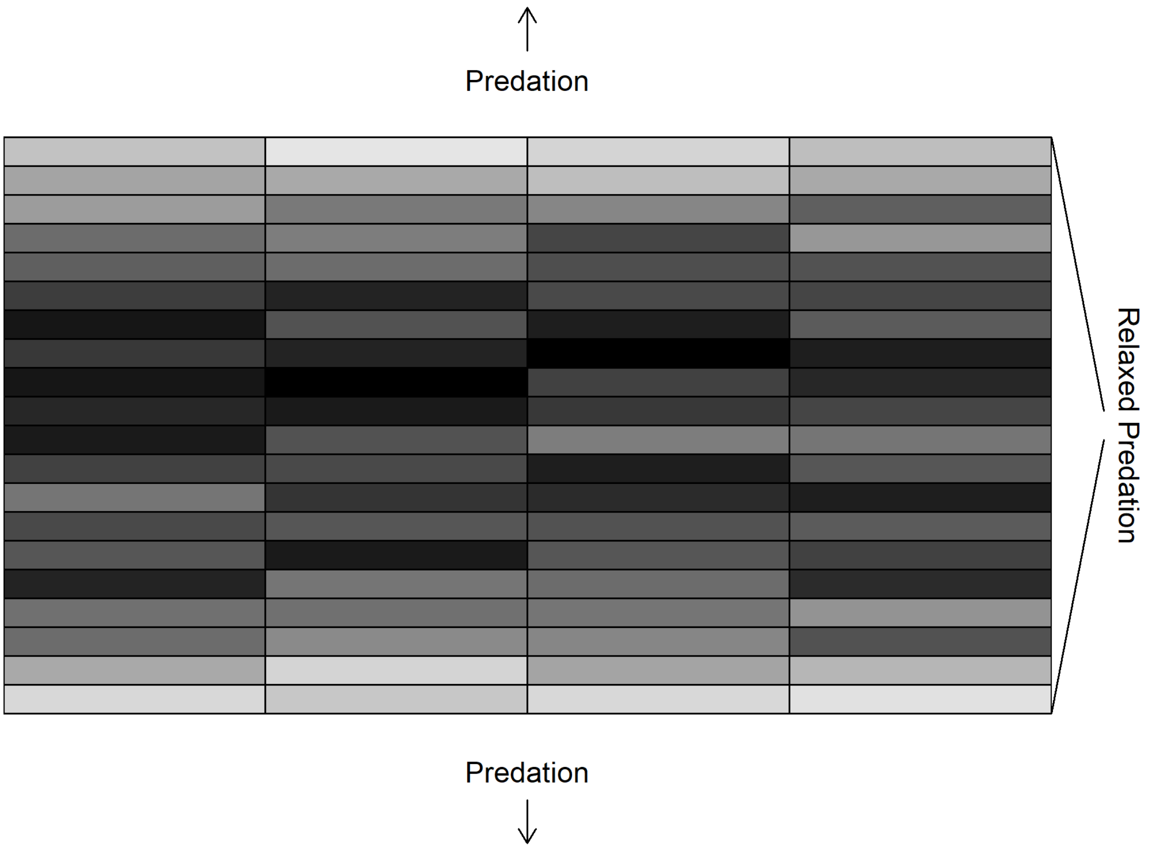


As an alternative solution, we plot this as a histogram:

```
png("Figures/simulations_location_first_mutant_histogram.png",
    width = 2000, height = 1500, res = 300)
par(mar = c(3.5, 4, 0.5, 1))
# Get counts for each region (looks
# complicated, but we basically just
# have to single out those patch IDs
# that are central)
hist(ifelse(c(l1_2500$mutator_patch,
    l5_2500$mutator_patch, l10_2500$mutator_patch) %in%
    39:58, c(l1_2500$mutator_patch,
    l5_2500$mutator_patch, l10_2500$mutator_patch) -
    38, ifelse(c(l1_2500$mutator_patch,
    l5_2500$mutator_patch, l10_2500$mutator_patch) %in%
    135:154, c(l1_2500$mutator_patch,
    l5_2500$mutator_patch, l10_2500$mutator_patch) -
    134, ifelse(c(l1_2500$mutator_patch,
    l5_2500$mutator_patch, l10_2500$mutator_patch) %in%
    231:250, c(l1_2500$mutator_patch,
    l5_2500$mutator_patch, l10_2500$mutator_patch) -
    230, c(l1_2500$mutator_patch, l5_2500$mutator_patch,
    l10_2500$mutator_patch) - 326))),
    main = "", xlab = "", ylab = "",
    xaxt = "n", breaks = seq(0, 20,
        length.out = 10))
# Indicate predation and
```

```
# non-predation regime
par(xpd = NA)
text(c(0, 20), par("usr")[3] - 0.15 *
    diff(par("usr")[3:4]), "Predation")
arrows(c(1, 19), par("usr")[3] - 0.11 *
    diff(par("usr")[3:4]), c(-1, 21),
    par("usr")[3] - 0.11 * diff(par("usr")[3:4]),
    length = 0.1)
par(xpd = F)

mtext("Simulations where Mutation Persisted",
    2, line = 3)

axis(1, at = c(0, 10, 20), c("Northern",
    "Center of Relaxed", "Southern"),
    lwd = 0, line = -1.5)
axis(1, at = c(0, 10, 20), c("End",
    "Predation Zone", "End"), lwd = 0,
    line = -0.7)
invisible(dev.off())
```

Show png we just made:

```
fig_disp <- readPNG("Figures/simulations_location_first_mutant_histogram.png")
grid.raster(fig_disp)
```

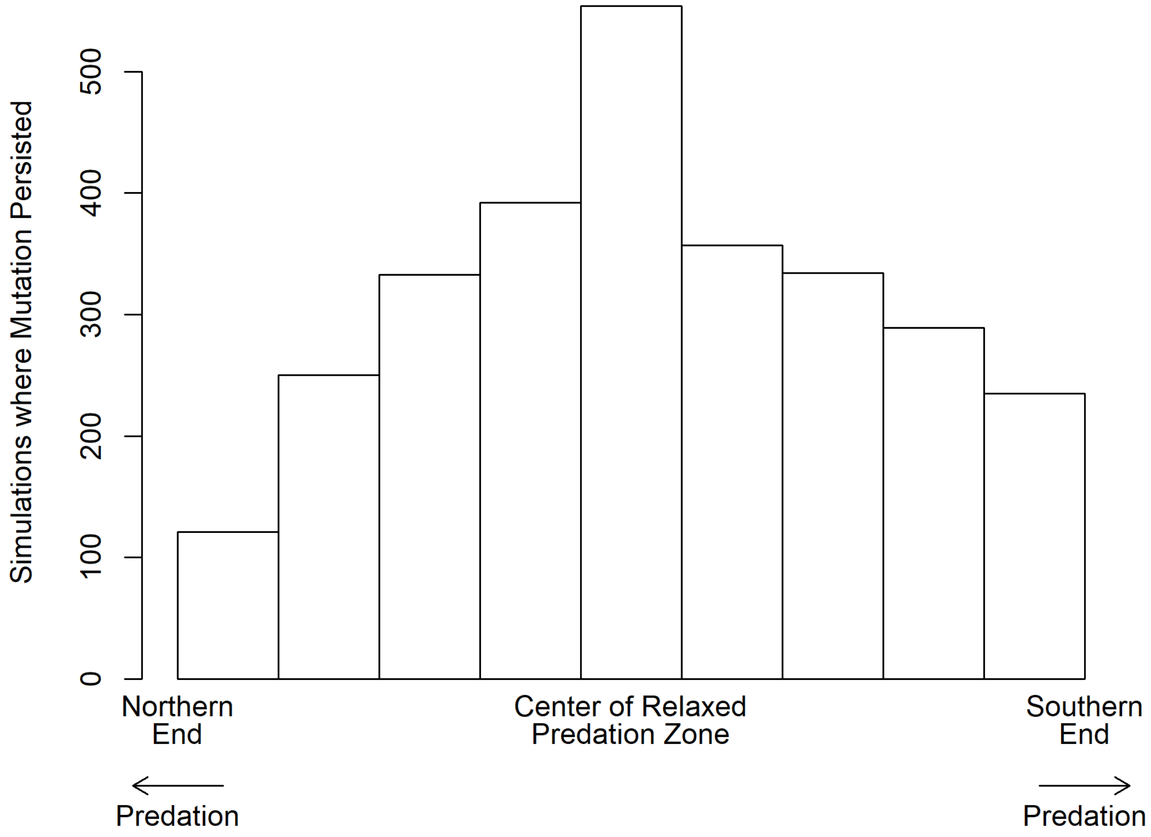


The mutation is most likely to persist if it occurs in a patch far from the predation-rich areas!

#### 3.3 How does the genetic architecture of attraction traits affect allele frequency dynamics?

In this section, we look at allele frequency dynamics: in general, but also with a particular emphasis on what role the genetic architecture of male attraction traits plays.

##### 3.3.1 Plots

**3.3.1.1 Number of loci and strength of attraction determine attraction phenotypes** First of all, we want to get a sense of how males with different attraction genotypes are attracted to colours as compared to a male with minimum attraction.

Calculate minimum and maximum attraction that can be reached, given different values for the strength of attraction.

```
strength_of_attraction1 <- 3
max_attr1 <- exp(strength_of_attraction1 *
    (1 - 1/2) * 1)
min_attr1 <- exp(strength_of_attraction1 *
    (0 - 1/2) * 1)

strength_of_attraction2 <- 1
max_attr2 <- exp(strength_of_attraction2 *
    (1 - 1/2) * 1)
min_attr2 <- exp(strength_of_attraction2 *
    (0 - 1/2) * 1)
```

Plot the relative attraction values (all relative to male with minimum attraction under given "strength of attraction" parameter). To bring it all to the same scale, we scale everything by `min_attr1` and `max_attr1`. If we don't do this, not all lines start in the same position, but those for "strength of attraction" = 1 would start higher up, as `min_attr2` > `min_attr1`. This might seem odd at first, but one has to keep in mind that these attraction values always have to be seen as relative to other attraction values under the same "strength of attraction" parameter. `max_attr1/min_attr1` equals to 20.1, while `max_attr2/min_attr2` only equals to 2.7.

```
png("Figures/simulations_attraction_loci.png",
    width = 2000, height = 1500, res = 300)
# Open empty plot and scale y-axis
# for minimum and maximum under
# strength of attraction = 3.
# logarithmize the y-axis (as we
# want to look at x-fold changes)
par(mar = c(3, 3.5, 0.5, 0.5))
plot(1, 1, type = "n", xaxt = "n", yaxt = "n",
    xlab = "", ylab = "", xlim = c(0,
        10), ylim = c(min_attr1/max_attr1,
        1), xaxs = "i", yaxs = "i",
    log = "y")
# Put thin lines at different
# x-values
abline(v = 1:10, lwd = 0.2)

# Define three colours
colores <- c(1, 2, 5)

# Add relative attraction (relative
```

```
# to maximum) for strength of
# attraction = 3 for the three
# different genetic architectures
for (i in 1:3) {
    n <- c(1, 5, 10)[i]
    atr_lv <- exp(strength_of_attraction1 *
        (((0:n)/n) - 1/2) * 1)/max_attr1
    lines(0:n, atr_lv, col = colores[i],
        lwd = 2.5)
}

# Add relative attraction (relative
# to maximum) for strength of
# attraction = 1 for the three
# different genetic architectures.
# Note that we scale this to the
# ratio of minimum by maximum
# between the two different sets of
# min and max values
for (i in 1:3) {
    n <- c(1, 5, 10)[i]
    atr_lv <- (exp(strength_of_attraction2 *
        (((0:n)/n) - 1/2) * 1))/max_attr2 *
        ((min_attr1/max_attr1)/(min_attr2/max_attr2))
    lines(0:n, atr_lv, col = colores[i],
        lwd = 2.5, lty = "dashed")
}

# Provide two legends (one for
# strenth of attraction, the other
# for genetic architecture)
legend(x = 7.63, y = 10^(par("usr")[3] +
    0.159 * (-par("usr")[3])), lwd = 2.5,
    lty = c("dashed", "solid"), c("1",
        "3"), title = expression(paste(italic(alpha))),
    cex = 0.85)
legend("bottomright", lwd = 2.5, col = colores,
    as.character(c(1, 5, 10)), title = "Loci",
    cex = 0.85)

# Axes and titles
axis(1, at = 0:10, lwd = 0, line = -0.9,
    cex.axis = 0.7)
mtext("Alleles causing attraction",
    1, line = 1.8, cex = 0.9)
# Values of x-fold changes
x <- c(1, 2, 5, 10, 20)
axis(2, at = (min_attr1/max_attr1) *
    x, paste0(x, "x"), lwd = 0, line = -0.5,
    cex.axis = 1, las = 2)
axis(2, at = (min_attr1/max_attr1) *
    x, labels = F, tck = -0.015)
mtext("Probability relative to minimum attraction",
```

```
    2, line = 2.5, cex = 0.9)

box(lwd = 1)
dev.off()
```

```
## pdf
##   2
```

Show png we just made:

```
fig_disp <- readPNG("Figures/simulations_attraction_loci.png")
grid.raster(fig_disp)
```

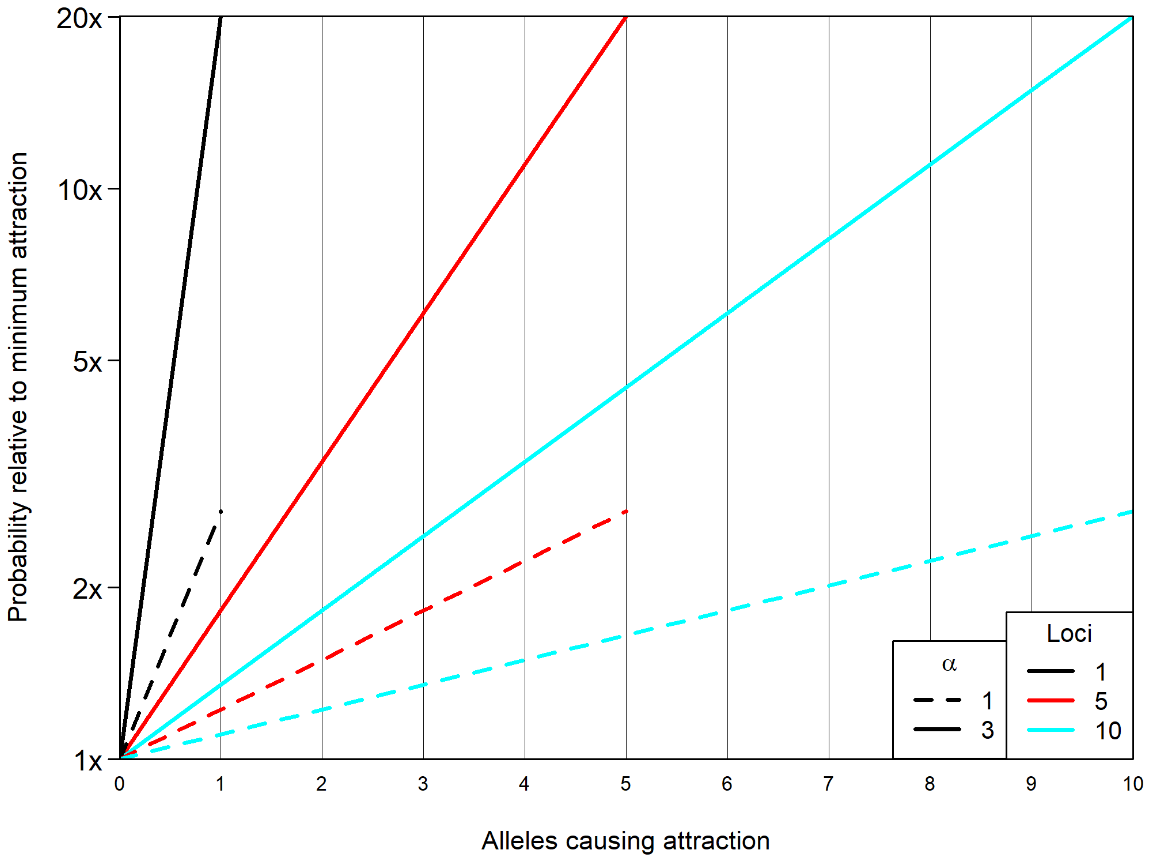


**3.3.1.2 Histograms of allele frequency dynamics** Calculate allele frequency at all loci coding either for wing pattern or for attraction to either colour for all last generations, as well as only for generation 2500 where the novel pattern survived.

Next, we calculate the maximum allele frequencies for the same two datasets.

Finally, we find the generations where an allele frequency of 0.95 for novel allele was reached at the colour pattern locus by those simulations, where frequencies reached 2500 (all simulations where the novel allele persisted reached 0.95).

We create frequency histograms for each set of loci (colour, preference for novel, preference for ancestral), for all types of datasets. For the last-generation dataset, as well as the maximum-frequency dataset, we create a set of histograms with 50 BINs. For the loci for attraction to the ancestral pattern, we additionally create a histogram with 1000 BINs for the last-generation dataset. We also create such high-resolution histograms for the colour pattern locus in the maximum-frequency dataset.

```r
# Histogram breaks:
number_breaks <- 51
breaks <- seq(0, 1, length.out = number_breaks)
number_breaks_zoom <- 1001
breaks_zoom <- seq(0, 1, length.out = number_breaks_zoom)
# Loci we check:
loci <- c(1, 5, 10)
# To ease coding:
z <- "last_gen_l"
# Empty list to save the indices
# where allele frequency of 0.95 was
# reached the first time
freq_exceed_095 <- list()
for (lo in loci) {

    # Frequencies at last generations:

    # For all simulation runs:
    freq_all <- as.data.frame(cbind((get(paste0("l",
        lo))$Cc_het[get(paste0(z, lo))] +
        get(paste0("l", lo))$Cnc_het[get(paste0(z,
            lo))] + 2 * get(paste0("l",
        lo))$Cc_hom[get(paste0(z, lo))] +
        2 * get(paste0("l", lo))$Cnc_hom[get(paste0(z,
            lo))])/(2 * get(paste0("l",
        lo))$N_central[get(paste0(z,
        lo))] + 2 * get(paste0("l",
        lo))$N_non_central[get(paste0(z,
        lo))]), sapply(1:lo, function(x) (get(paste0("l",
        lo))[get(paste0(z, lo)), paste0("Rc",
        x, "_het")] + get(paste0("l",
        lo))[get(paste0(z, lo)), paste0("Rnc",
        x, "_het")] + 2 * get(paste0("l",
        lo))[get(paste0(z, lo)), paste0("Rc",
        x, "_hom")] + 2 * get(paste0("l",
        lo))[get(paste0(z, lo)), paste0("Rnc",
        x, "_hom")])/(2 * get(paste0("l",
        lo))$N_central[get(paste0(z,
        lo))] + 2 * get(paste0("l",
        lo))$N_non_central[get(paste0(z,
        lo))])), sapply(1:lo, function(x) (get(paste0("l",
        lo))[get(paste0(z, lo)), paste0("Wc",
        x, "_het")] + get(paste0("l",
        lo))[get(paste0(z, lo)), paste0("Wnc",
        x, "_het")] + 2 * get(paste0("l",
        lo))[get(paste0(z, lo)), paste0("Wc",
        x, "_hom")] + 2 * get(paste0("l",
        lo))[get(paste0(z, lo)), paste0("Wnc",
        x, "_hom")])/(2 * get(paste0("l",
        lo))$N_central[get(paste0(z,
        lo))] + 2 * get(paste0("l",
        lo))$N_non_central[get(paste0(z,
        lo))]))))
```

```
eval(parse(text = paste0("names(freq_all)",
    "<-c('colour',paste0('l',1:lo),paste0('ancestral_pref_l',1:lo))")))
# Repeat for only those simulations
# where novel allele survived
freq_fix <- as.data.frame(cbind((get(paste0("l",
    lo, "_2500"))$Cc_het + get(paste0("l",
    lo, "_2500"))$Cnc_het + 2 *
    get(paste0("l", lo, "_2500"))$Cc_hom +
    2 * get(paste0("l", lo, "_2500"))$Cnc_hom)/(2 *
    get(paste0("l", lo, "_2500"))$N_central +
    2 * get(paste0("l", lo, "_2500"))$N_non_central),
    sapply(1:lo, function(x) (get(paste0("l",
        lo, "_2500"))[, paste0("Rc",
        x, "_het")] + get(paste0("l",
        lo, "_2500"))[, paste0("Rnc",
        x, "_het")] + 2 * get(paste0("l",
        lo, "_2500"))[, paste0("Rc",
        x, "_hom")] + 2 * get(paste0("l",
        lo, "_2500"))[, paste0("Rnc",
        x, "_hom")])/(2 * get(paste0("l",
        lo, "_2500"))$N_central +
        2 * get(paste0("l", lo,
            "_2500"))$N_non_central)),
    sapply(1:lo, function(x) (get(paste0("l",
        lo, "_2500"))[, paste0("Wc",
        x, "_het")] + get(paste0("l",
        lo, "_2500"))[, paste0("Wnc",
        x, "_het")] + 2 * get(paste0("l",
        lo, "_2500"))[, paste0("Wc",
        x, "_hom")] + 2 * get(paste0("l",
        lo, "_2500"))[, paste0("Wnc",
        x, "_hom")])/(2 * get(paste0("l",
        lo, "_2500"))$N_central +
        2 * get(paste0("l", lo,
            "_2500"))$N_non_central))))
eval(parse(text = paste0("names(freq_fix)",
    "<-c('colour',paste0('l',1:lo),paste0('ancestral_pref_l',1:lo))")))
# Create a set of histograms for
# each set of loci in these tables
# First for all simulations
assign(paste0("all_last_colour",
    lo), hist(freq_all$colour, breaks = breaks,
    plot = F))
assign(paste0("all_last_pref", lo),
    hist(unlist(freq_all[, 2:(1 +
        lo)]), breaks = breaks,
        plot = F))
assign(paste0("all_last_ancestral_pref",
    lo), hist(unlist(freq_all[,
    (2 + lo):(1 + 2 * lo)]), breaks = breaks,
    plot = F))
# calculate average frequencies
# (divide by total cases and (for
```

```
# cases of more than one preference
# locus) by total number of
# preference loci)
eval(parse(text = paste0("all_last_colour",
    lo, "$counts<-all_last_colour",
    lo, "$counts/nrow(freq_all)")))
eval(parse(text = paste0("all_last_pref",
    lo, "$counts<-all_last_pref",
    lo, "$counts/(nrow(freq_all)*",
    lo, ")")))
eval(parse(text = paste0("all_last_ancestral_pref",
    lo, "$counts<-all_last_ancestral_pref",
    lo, "$counts/(nrow(freq_all)*",
    lo, ")")))

# Now repeat histograms for only
# those simulations where generation
# 2500 was reached
assign(paste0("colour", lo), hist(freq_fix$colour,
    breaks = breaks, plot = F))
assign(paste0("pref", lo), hist(unlist(freq_fix[,
    2:(1 + lo)]), breaks = breaks,
    plot = F))
assign(paste0("ancestral_pref",
    lo), hist(unlist(freq_fix[,
    (2 + lo):(1 + 2 * lo)]), breaks = breaks,
    plot = F))
# calculate average frequencies
# (divide by total cases and (for
# cases of more than one preference
# locus) by total number of
# preference loci)
eval(parse(text = paste0("colour",
    lo, "$counts<-colour", lo, "$counts/nrow(freq_all)")))
eval(parse(text = paste0("pref",
    lo, "$counts<-pref", lo, "$counts/(nrow(freq_all)*",
    lo, ")")))
eval(parse(text = paste0("ancestral_pref",
    lo, "$counts<-ancestral_pref",
    lo, "$counts/(nrow(freq_all)*",
    lo, ")")))

# Maximum frequencies & first
# generation >= 0.95:

# Extract the generation where
# maximum freqency was reached per
# simulation. Also extract the
# generation where frequency
# exceeded 0.95 for the first time
# (in case it did) Number of
# simulations:
simul_par <- 2000
```

```
# Empty index vector
index_max_freq <- rep(NA, nrow(param) *
    simul_par)
index_095 <- rep(NA, nrow(param) *
    simul_par)
# Allele frequency at all
# generations
allele_freq_novel <- (get(paste0("l",
    lo))$Cc_het + get(paste0("l",
    lo))$Cnc_het + 2 * get(paste0("l",
    lo))$Cc_hom + 2 * get(paste0("l",
    lo))$Cnc_hom)/(2 * get(paste0("l",
    lo))$N_central + 2 * get(paste0("l",
    lo))$N_non_central)
nrow_l <- nrow(get(paste0("l", lo)))
# Walk through all parameter
# combinations
for (i in 1:nrow(param)) {
    # These indices match:
    where_i <- (1:nrow_l)[get(paste0("l",
        lo))$Scenario == i]
    # Now find start and end index of
    # each simulation. We make use of
    # the tables being ordered:
    starters <- c(where_i[1], where_i[-1][get(paste0("l",
        lo))$Simulation[where_i[-1]] >
        get(paste0("l", lo))$Simulation[where_i[-length(where_i)]]])
    enders <- c(starters[-1] - 1,
        where_i[length(where_i)])
    # Find which index has the maximum
    index_max_freq[((i - 1) * simul_par +
        1):(i * simul_par)] <- sapply(1:simul_par,
        function(s) {
            z <- (starters[s]:enders[s])
            z[max.col(t(allele_freq_novel[z]),
              "last")]
        })
    # Find which index is the first to
    # reach 0.95 (if not reached, an NA
    # will be returned)
    index_095[((i - 1) * simul_par +
        1):(i * simul_par)] <- sapply(1:simul_par,
        function(s) {
            z <- (starters[s]:enders[s])
            z[allele_freq_novel[z] >=
              0.95][1]
        })
}
# Remove the NAs
freq_exceed_095[[length(freq_exceed_095) +
    1]] <- index_095[!is.na(index_095)]

# Create a new table, extracting
```

```
# only those generations with
# maximum allele frequency
max_freq_t <- get(paste0("l", lo))[index_max_freq,
    ]

# Add new 'survivor' column
max_freq_t$survivor <- FALSE
# We walk through the different
# scenarios and check if a certain
# simulation run can be found in the
# table of survivors
for (i in 1:nrow(param)) {
    sim_with_success <- get(paste0("l",
        lo, "_2500"))$Simulation[get(paste0("l",
        lo, "_2500"))$Scenario ==
        i]
    max_freq_t$survivor[max_freq_t$Scenario ==
        i & max_freq_t$Simulation %in%
        sim_with_success] <- TRUE
}
# Add allele frequency (already
# calculated above)
max_freq_t$allele_freq_col <- allele_freq_novel[index_max_freq]

# Calculate allele frequencies when
# pattern is at maximum frequency.
# For all simulation runs:
freq_all_max <- as.data.frame(cbind((max_freq_t$Cc_het +
    max_freq_t$Cnc_het + 2 * max_freq_t$Cc_hom +
    2 * max_freq_t$Cnc_hom)/(2 *
    max_freq_t$N_central + 2 * max_freq_t$N_non_central),
    sapply(1:lo, function(x) (max_freq_t[,
        paste0("Rc", x, "_het")] +
        max_freq_t[, paste0("Rnc",
            x, "_het")] + 2 * max_freq_t[,
        paste0("Rc", x, "_hom")] +
        2 * max_freq_t[, paste0("Rnc",
            x, "_hom")])/(2 * max_freq_t$N_central +
        2 * max_freq_t$N_non_central)),
    sapply(1:lo, function(x) (max_freq_t[,
        paste0("Wc", x, "_het")] +
        max_freq_t[, paste0("Wnc",
            x, "_het")] + 2 * max_freq_t[,
        paste0("Wc", x, "_hom")] +
        2 * max_freq_t[, paste0("Wnc",
            x, "_hom")])/(2 * max_freq_t$N_central +
        2 * max_freq_t$N_non_central))))
eval(parse(text = paste0("names(freq_all_max)",
    "<-c('colour',paste0('l',1:lo),paste0('ancestral_pref_l',1:lo))")))
# Extract only those simulations
# where novel allele survived
freq_fix_max <- freq_all_max[max_freq_t$survivor,
    ]
```

```
# As before, we create a set of
# histograms for each set of loci in
# these tables First for all
# simulations
assign(paste0("all_max_colour",
    lo), hist(freq_all_max$colour,
    breaks = breaks, plot = F))
assign(paste0("all_max_pref", lo),
    hist(unlist(freq_all_max[, 2:(1 +
        lo)]), breaks = breaks,
        plot = F))
assign(paste0("all_max_ancestral_pref",
    lo), hist(unlist(freq_all_max[,
    (2 + lo):(1 + 2 * lo)]), breaks = breaks,
    plot = F))
# calculate average frequencies
# (divide by total cases and (for
# cases of more than one preference
# locus) by total number of
# preference loci)
eval(parse(text = paste0("all_max_colour",
    lo, "$counts<-all_max_colour",
    lo, "$counts/nrow(freq_all_max)")))
eval(parse(text = paste0("all_max_pref",
    lo, "$counts<-all_max_pref",
    lo, "$counts/(nrow(freq_all_max)*",
    lo, ")")))
eval(parse(text = paste0("all_max_ancestral_pref",
    lo, "$counts<-all_max_ancestral_pref",
    lo, "$counts/(nrow(freq_all_max)*",
    lo, ")")))

# Now repeat histograms for only
# those simulations where generation
# 2500 was reached
assign(paste0("all_max_surv_colour",
    lo), hist(freq_fix_max$colour,
    breaks = breaks, plot = F))
assign(paste0("all_max_surv_pref",
    lo), hist(unlist(freq_fix_max[,
    2:(1 + lo)]), breaks = breaks,
    plot = F))
assign(paste0("all_max_surv_ancestral_pref",
    lo), hist(unlist(freq_fix_max[,
    (2 + lo):(1 + 2 * lo)]), breaks = breaks,
    plot = F))
# calculate average frequencies
# (divide by total cases and (for
# cases of more than one preference
# locus) by total number of
# preference loci)
eval(parse(text = paste0("all_max_surv_colour",
    lo, "$counts<-all_max_surv_colour",
```

```
        lo, "$counts/nrow(freq_all_max)")))
    eval(parse(text = paste0("all_max_surv_pref",
        lo, "$counts<-all_max_surv_pref",
        lo, "$counts/(nrow(freq_all_max)*",
        lo, ")")))
    eval(parse(text = paste0("all_max_surv_ancestral_pref",
        lo, "$counts<-all_max_surv_ancestral_pref",
        lo, "$counts/(nrow(freq_all_max)*",
        lo, ")")))


    # Make histograms with 1000 BINs

    # Zoom on colour pattern allele
    # frequency at maximum allele freq
    assign(paste0("all_max_colour_zoom",
        lo), hist(freq_all_max$colour,
        breaks = breaks_zoom, plot = F))
    # Zoom on ancestral pref allele
    # frequency at last generation
    assign(paste0("all_last_ancestral_pref_zoom",
        lo), hist(unlist(freq_all[,
        (2 + lo):(1 + 2 * lo)]), breaks = breaks_zoom,
        plot = F))
    eval(parse(text = paste0("all_max_colour_zoom",
        lo, "$counts<-all_max_colour_zoom",
        lo, "$counts/nrow(freq_all_max)")))
    eval(parse(text = paste0("all_last_ancestral_pref_zoom",
        lo, "$counts<-all_last_ancestral_pref_zoom",
        lo, "$counts/(nrow(freq_all)*",
        lo, ")")))

    # Same again for only those
    # simulations, where novel colour
    # survives
    assign(paste0("all_max_surv_colour_zoom",
        lo), hist(freq_fix_max$colour,
        breaks = breaks_zoom, plot = F))
    assign(paste0("all_last_surv_ancestral_pref_zoom",
        lo), hist(unlist(freq_fix[,
        (2 + lo):(1 + 2 * lo)]), breaks = breaks_zoom,
        plot = F))
    eval(parse(text = paste0("all_max_surv_colour_zoom",
        lo, "$counts<-all_max_surv_colour_zoom",
        lo, "$counts/nrow(freq_all_max)")))
    eval(parse(text = paste0("all_last_surv_ancestral_pref_zoom",
        lo, "$counts<-all_last_surv_ancestral_pref_zoom",
        lo, "$counts/(nrow(freq_all)*",
        lo, ")")))

}
```

First, we plot the allele frequencies at the last generation, colour coding whether a simulation resulted in the novel allele to survive or not.

Summarise all frequencies in one vector (only for the big dataset, *i.e.* all generations).

```
all_last_all_freqs <- c(all_last_colour1$counts,
    all_last_colour5$counts, all_last_colour10$counts,
    all_last_pref1$counts, all_last_pref5$counts,
    all_last_pref10$counts, all_last_ancestral_pref1$counts,
    all_last_ancestral_pref5$counts,
    all_last_ancestral_pref10$counts)
# Also calculate the range of the
# frequencies The 0.5 makes sure
# that e.g. that the bar for a
# proportion resulting from 1 count
# is half as big as of a proportion
# resulting from 2 counts
all_last_y_range <- c(0.5 * min(all_last_all_freqs[all_last_all_freqs >
    0]), 1)
```

Let's plot this:

```
png("Figures/simulations_average_allele_freq_survivors_all_last.png",
    width = 2800, height = 2200, res = 300)
layout(matrix(1:9, ncol = 3))
par(mar = c(2, 2, 0.5, 0.5))
par(oma = c(2, 3.5, 2, 0))

# Title for each row of plots
y_titl <- c("Colour locus", "Preference (novel)",
    "Preference (ancestral)")
# Name of each of the 3 histogram
# categories
ht <- c("colour", "pref", "ancestral_pref")
# Go through three datasets
for (lo in loci) {
    # Loop over the 3 different
    # categories of loci
    for (subpl in 1:3) {
        # Open plot
        plot(1, 1, type = "n", bty = "n",
            log = "y", xlab = "", ylab = "",
            main = "", xlim = c(0, 1),
            ylim = all_last_y_range,
            yaxs = "i")
        # If the right moment, add some
        # titles
        if (subpl == 1) {
            mtext(paste0(lo, " preference loci"),
                3, line = 1.1, font = 2)
            mtext(paste0(format(round(100 *
                (nrow(get(paste0("l",
                  lo, "_2500")))/(nrow(param) *
                  2000)), 3), nsmall = 3),
                "% simulations survival"),
                3, line = -0.5)
        }
        if (lo == 1) {
```

```
            mtext(y_titl[subpl], 2,
                line = 4.3, font = 2)
        }
        # Add histograms manually, with
        # y-axis being log-scaled (all
        # simulations in black, simulations
        # that reached 2500 in red)
        rect(col = "gray20", border = NA,
            get(paste0("all_last_",
                ht[subpl], lo))$breaks[1:(number_breaks -
                1)], rep(all_last_y_range[1],
                number_breaks), get(paste0("all_last_",
                ht[subpl], lo))$breaks[2:number_breaks],
            get(paste0("all_last_",
                ht[subpl], lo))$counts)
        rect(col = "red", border = NA,
            get(paste0(ht[subpl], lo))$breaks[1:(number_breaks -
                1)], rep(all_last_y_range[1],
                number_breaks), get(paste0(ht[subpl],
                lo))$breaks[2:number_breaks],
            get(paste0(ht[subpl], lo))$counts)
    }
}

# Add axis titles
mtext("log-Proportion", 2, outer = T,
    line = 0.4)
mtext("(Mean) allele Frequency", 1,
    outer = T, line = 0.4)

invisible(dev.off())
```

Show png we just made:

```
fig_disp <- readPNG("Figures/simulations_average_allele_freq_survivors_all_last.png")
grid.raster(fig_disp)
```

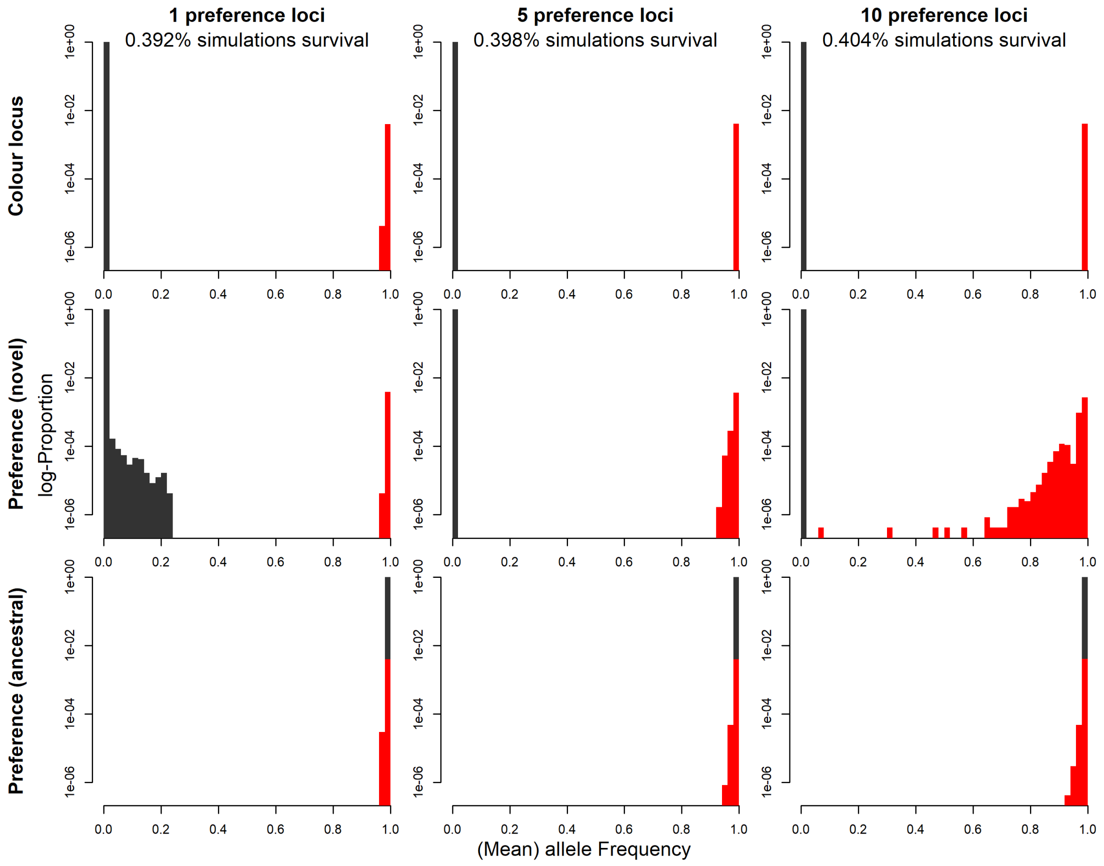


We repeat the exact same, this time for the maximum allele frequency of the novel pattern that was reached:

```
# Summarise all frequencies in one
# vector (only for the big dataset,
# i.e. all generations)
all_max_all_freqs <- c(all_max_colour1$counts,
    all_max_colour5$counts, all_max_colour10$counts,
    all_max_pref1$counts, all_max_pref5$counts,
    all_max_pref10$counts, all_max_ancestral_pref1$counts,
    all_max_ancestral_pref5$counts,
    all_max_ancestral_pref10$counts)

all_max_y_range <- c(0.5 * min(all_max_all_freqs[all_max_all_freqs >
    0]), 1)

png("Figures/simulations_average_allele_freq_survivors_all_max.png",
    width = 2800, height = 2200, res = 300)
layout(matrix(1:9, ncol = 3))
par(mar = c(2, 2, 0.5, 0.5))
par(oma = c(2, 3.5, 2, 0))

for (lo in loci) {
    for (subpl in 1:3) {
        plot(1, 1, type = "n", bty = "n",
            log = "y", xlab = "", ylab = "",
            main = "", xlim = c(0, 1),
            ylim = all_max_y_range,
```

```
            yaxs = "i")
        if (subpl == 1) {
            mtext(paste0(lo, " preference loci"),
                3, line = 1.1, font = 2)
            mtext(paste0(format(round(100 *
                (nrow(get(paste0("l",
                  lo, "_2500")))/(nrow(param) *
                  2000)), 3), nsmall = 3),
                "% simulations survival"),
                3, line = -0.5)
        }
        if (lo == 1) {
            mtext(y_titl[subpl], 2,
                line = 4.3, font = 2)
        }
        rect(col = "gray20", border = NA,
            get(paste0("all_max_", ht[subpl],
                lo))$breaks[1:(number_breaks -
                1)], rep(all_max_y_range[1],
                number_breaks), get(paste0("all_max_",
                ht[subpl], lo))$breaks[2:number_breaks],
            get(paste0("all_max_", ht[subpl],
                lo))$counts)
        rect(col = "red", border = NA,
            get(paste0("all_max_surv_",
                ht[subpl], lo))$breaks[1:(number_breaks -
                1)], rep(all_max_y_range[1],
                number_breaks), get(paste0("all_max_surv_",
                ht[subpl], lo))$breaks[2:number_breaks],
            get(paste0("all_max_surv_",
                ht[subpl], lo))$counts)
    }
}

mtext("log-Proportion", 2, outer = T,
    line = 0.4)
mtext("(Mean) allele Frequency", 1,
    outer = T, line = 0.4)

invisible(dev.off())
```

Show png we just made:

```
fig_disp <- readPNG("Figures/simulations_average_allele_freq_survivors_all_max.png")
grid.raster(fig_disp)
```

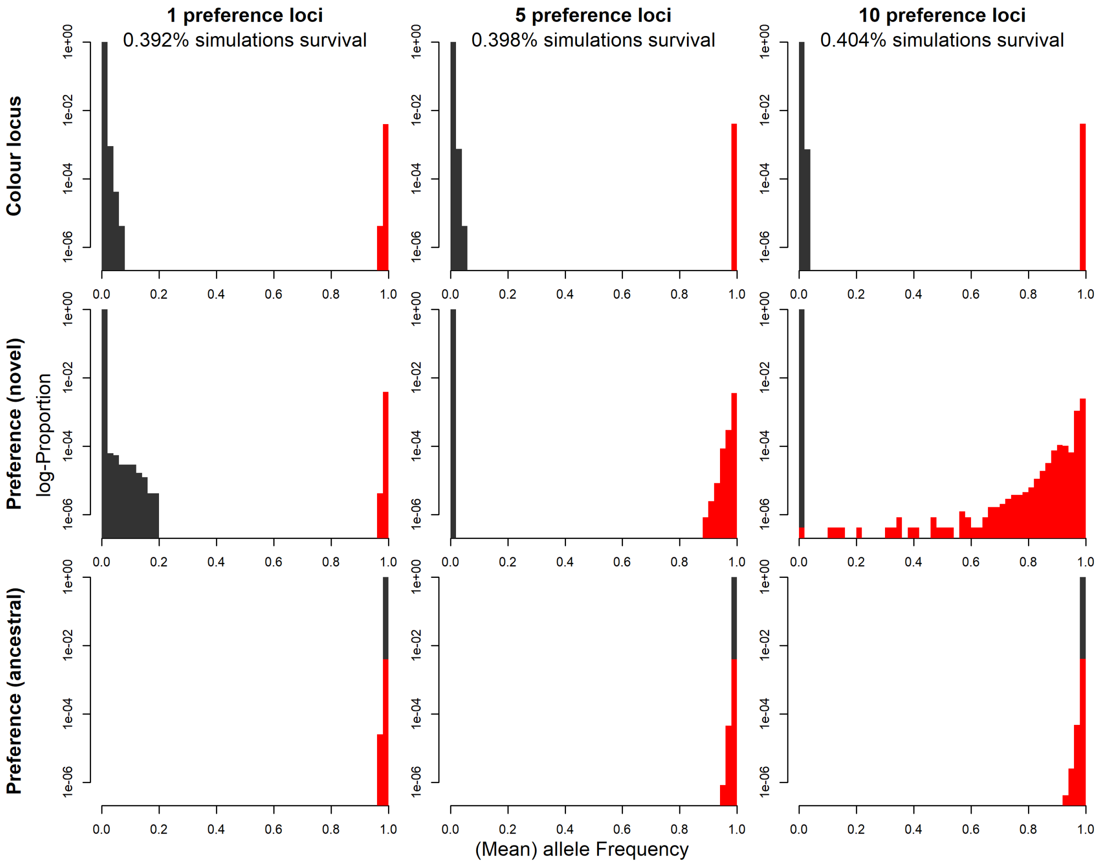


To get a better idea of these histograms for the colour pattern locus, as well as for loci coding for preference to the ancestral type, we create two 'zoomed in' histograms (following the same logic as above).

For colour pattern (here we look at the maximum-frequency dataset):

```
# Summarise all frequencies in one
# vector (only for the big dataset,
# i.e. all generations)
all_max_all_freqs <- c(all_max_colour_zoom1$counts,
    all_max_colour_zoom5$counts, all_max_colour_zoom10$counts)

# The 0.5 makes sure that e.g. that
# the bar for a proportion resulting
# from 1 count is half as big as of
# a proportion resulting from 2
# counts
all_max_y_range <- c(0.5 * min(all_max_all_freqs[all_max_all_freqs >
    0]), 1)

png("Figures/simulations_colour_allele_freq_survivors_all_max_zoom.png",
    width = 2800, height = 2200, res = 300)
layout(matrix(1:6, ncol = 2, byrow = T))
par(mar = c(2, 2, 0.5, 0.5))
par(oma = c(2, 3.5, 0, 0))

for (lo in loci) {
    for (subpl in 1:2) {
```

```
        plot(1, 1, type = "n", bty = "n",
            log = "y", xlab = "", ylab = "",
            main = "", xlim = list(c(0,
                0.066), c(0.934, 1))[[subpl]],
            ylim = all_max_y_range,
            yaxs = "i", xaxt = "n")
        axis(1, at = 0:1, labels = F)
        axis(1, at = list(seq(0, 0.06,
            0.01), seq(0.94, 1, 0.01))[[subpl]])
        if (subpl == 1) {
            mtext(paste0(lo, " preference loci"),
                2, line = 4.3)
        }
        rect(col = "gray20", border = NA,
            get(paste0("all_max_colour_zoom",
                lo))$breaks[1:(number_breaks_zoom -
                1)], rep(all_max_y_range[1],
                number_breaks_zoom),
            get(paste0("all_max_colour_zoom",
                lo))$breaks[2:number_breaks_zoom],
            get(paste0("all_max_colour_zoom",
                lo))$counts)
        rect(col = "red", border = NA,
            get(paste0("all_max_surv_colour_zoom",
                lo))$breaks[1:(number_breaks_zoom -
                1)], rep(all_max_y_range[1],
                number_breaks_zoom),
            get(paste0("all_max_surv_colour_zoom",
                lo))$breaks[2:number_breaks_zoom],
            get(paste0("all_max_surv_colour_zoom",
                lo))$counts)
    }
}

mtext("log-Proportion (only colour locus)",
    2, outer = T, line = 0.4)
mtext("Allele Frequency", 1, outer = T,
    line = 0.4)

invisible(dev.off())
```

Show png we just made:

```
fig_disp <- readPNG("Figures/simulations_colour_allele_freq_survivors_all_max_zoom.png")
grid.raster(fig_disp)
```

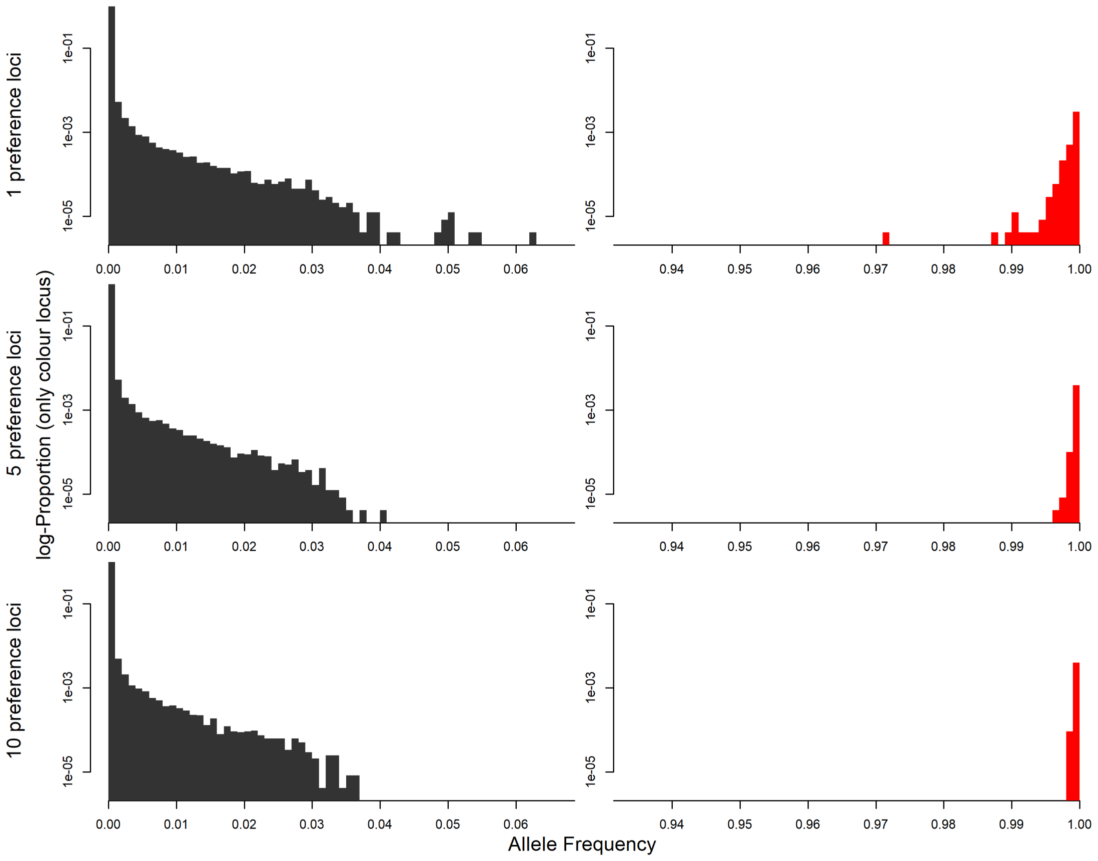


For preference for ancestral pattern (here we look at the last-generation dataset):

```
# Summarise all frequencies in one
# vector (only for the big dataset,
# i.e. all generations)
all_last_all_freqs <- c(all_last_ancestral_pref_zoom1$counts,
    all_last_ancestral_pref_zoom5$counts,
    all_last_ancestral_pref_zoom10$counts)

# The 0.5 makes sure that e.g. that
# the bar for a proportion resulting
# from 1 count is half as big as of
# a proportion resulting from 2
# counts
all_last_y_range <- c(0.5 * min(all_last_all_freqs[all_last_all_freqs >
    0]), 1)

png("Figures/simulations_ancestral_pref_allele_freq_survivors_all_last_zoom.png",
    width = 1500, height = 2200, res = 300)
layout(matrix(1:3, ncol = 1, byrow = T))
par(mar = c(2, 2, 0.5, 0.2))
par(oma = c(2, 4, 1, 0))

for (lo in loci) {
    plot(1, 1, type = "n", bty = "n",
        log = "y", xlab = "", ylab = "",
        main = "", xlim = c(0.934, 1),
```

```
        ylim = all_last_y_range, yaxs = "i",
        xaxt = "n")
    axis(1, at = 0:1, labels = F)
    axis(1, at = seq(0.94, 1, 0.01))
    mtext(paste0(lo, " preference loci"),
        2, line = 4.8)
    rect(col = "gray20", border = NA,
        get(paste0("all_last_ancestral_pref_zoom",
            lo))$breaks[1:(number_breaks_zoom -
            1)], rep(all_last_y_range[1],
            number_breaks_zoom), get(paste0("all_last_ancestral_pref_zoom",
            lo))$breaks[2:number_breaks_zoom],
        get(paste0("all_last_ancestral_pref_zoom",
            lo))$counts)
    rect(col = "red", border = NA, get(paste0("all_last_surv_ancestral_pref_zoom",
        lo))$breaks[1:(number_breaks_zoom -
        1)], rep(all_last_y_range[1],
        number_breaks_zoom), get(paste0("all_last_surv_ancestral_pref_zoom",
        lo))$breaks[2:number_breaks_zoom],
        get(paste0("all_last_surv_ancestral_pref_zoom",
            lo))$counts)
}

mtext("log-Proportion (only loci for preference to ancestral)",
    2, outer = T, line = 0.8)
mtext("Mean allele Frequency", 1, outer = T,
    line = 0.4)

invisible(dev.off())
```

Show png we just made:

```
fig_disp <- readPNG("Figures/simulations_ancestral_pref_allele_freq_survivors_all_last_zoom.png")
grid.raster(fig_disp)
```

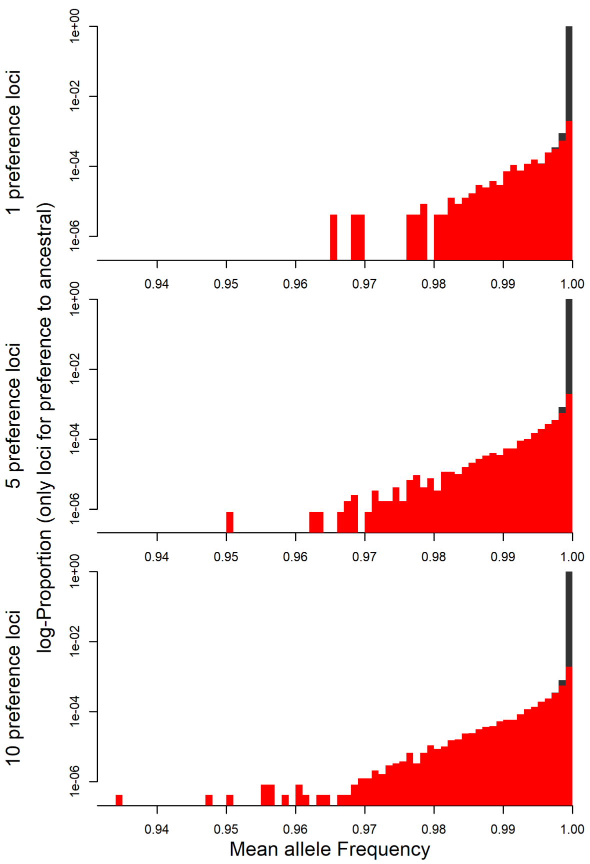


**3.3.1.3 Line plots of allele frequency dynamics** Finally, we visualize the full allele frequency dynamics for all scenarios where the novel pattern survived until generation 2500.

Get all scenarios where the novel allele survived (note that here we don't differentiate between male and female first mutant; this will follow below)

```
all_scen <- sort(unique(c(l1_2500$Scenario,
    l5_2500$Scenario, l10_2500$Scenario)))
# Reorder by order in graph:
all_scen <- all_scen[order(param$pred_constant[all_scen],
    param$prob_encounter[all_scen],
    param$strength_attract[all_scen],
    param$bottleneck[all_scen])]
```

To calculate the total number of plots we need, we check for each sex of first mutants how many times the novel pattern survived (across all datasets, *i.e.* irrespective of number of loci):

```
all_scen_mf <- sapply(all_scen, function(x) c(sum(c(l1_2500$sex_mutant[l1_2500$Scenario ==
    x] == 0, l5_2500$sex_mutant[l5_2500$Scenario ==
    x] == 0, l10_2500$sex_mutant[l10_2500$Scenario ==
    x] == 0)), sum(c(l1_2500$sex_mutant[l1_2500$Scenario ==
    x] == 1, l5_2500$sex_mutant[l5_2500$Scenario ==
    x] == 1, l10_2500$sex_mutant[l10_2500$Scenario ==
    x] == 1))))
```

Then, we take the number of non-zero cells from all_scen_mf, and this times 3, as we split by number of loci. Additionally, we have to add one 'column' of plots to display the parameter settings for each scenario.

```
number_of_plots <- sum(all_scen_mf >
    0) * 4
```

Now, we plot this:

```
first_row <- T
png("Figures/simulations_allele_freq_scenarios.png",
    width = 2000, height = 6500, res = 300)

layout(matrix(1:number_of_plots, ncol = 4,
    byrow = T))
par(oma = c(3, 0, 1.55, 2))

# Go through all scenarios
for (sceni in 1:length(all_scen)) {
    scenario <- all_scen[sceni]
    # For each scenario, go through the
    # two sexes
    for (sex_mutant in 0:1) {
        # If the scenario and this sex of
        # the first mutant ever led to
        # survival of the novel allele:
        if (all_scen_mf[sex_mutant +
            1, sceni] > 0) {
            # empty plot for parameter settings:
            par(mar = c(0.5, 0.3, 0.5,
                1.8))
            # Empty plot
            plot(1, 1, type = "n", xaxt = "n",
                yaxt = "n", xlab = "",
                ylab = "", bty = "n",
                ylim = c(0.1, 0.9),
                xlim = c(0, 2), xaxs = "i")
            # Add parameter settings to plot
            texti <- param[all_scen[sceni],
                ]
            text(1.3, 0.86, "Sexual conflict:   ",
                pos = 2, offset = 0,
                cex = 0.95)
            text(1.3, 0.86, c("no",
                "yes")[texti$sex_conflict +
                1], pos = 4, offset = 0,
                cex = 0.95)
            text(1.3, 0.74, "Predators return:   ",
                pos = 2, offset = 0,
                cex = 0.95)
            text(1.3, 0.74, texti$predator_absence,
                pos = 4, offset = 0,
                cex = 0.95)
            text(1.3, 0.62, "Learn. threshold:   ",
                pos = 2, offset = 0,
                cex = 0.95)
            text(1.3, 0.62, texti$pred_constant,
                pos = 4, offset = 0,
```

```
        cex = 0.95)
    text(1.3, 0.5, "Prob. encounter:   ",
        pos = 2, offset = 0,
        cex = 0.95)
    text(1.3, 0.5, texti$prob_encounter,
        pos = 4, offset = 0,
        cex = 0.95)
    text(1.3, 0.38, "Strength attr.:   ",
        pos = 2, offset = 0,
        cex = 0.95)
    text(1.3, 0.38, texti$strength_attract,
        pos = 4, offset = 0,
        cex = 0.95)
    text(1.3, 0.26, "Bottleneck:   ",
        pos = 2, offset = 0,
        cex = 0.95)
    text(1.3, 0.26, c("no",
        "yes")[texti$bottleneck +
        1], pos = 4, offset = 0,
        cex = 0.95)
    text(1.3, 0.14, "Sex first mutant:   ",
        pos = 2, offset = 0,
        cex = 0.95)
    text(1.3, 0.14, c("Female",
        "Male")[sex_mutant +
        1], pos = 4, offset = 0,
        cex = 0.95)

    # Now, each row of the plotting
    # scheme receives one plot for each
    # set of loci
    for (lo in loci) {

        # Extract results for current
        # scenario
        scenario_simulation <- get(paste0("l",
          lo, "_2500"))[get(paste0("l",
          lo, "_2500"))$Scenario ==
          scenario & get(paste0("l",
          lo, "_2500"))$sex_mutant ==
          sex_mutant, c("Scenario",
          "Simulation")]
        # Get column names for preference
        # loci
        novel_attr_c_hom <- paste0("Rc",
          1:lo, "_hom")
        novel_attr_nc_hom <- paste0("Rnc",
          1:lo, "_hom")
        novel_attr_c_het <- paste0("Rc",
          1:lo, "_het")
        novel_attr_nc_het <- paste0("Rnc",
          1:lo, "_het")
        ances_attr_c_hom <- paste0("Wc",
```

```
  1:lo, "_hom")
ances_attr_nc_hom <- paste0("Wnc",
  1:lo, "_hom")
ances_attr_c_het <- paste0("Wc",
  1:lo, "_het")
ances_attr_nc_het <- paste0("Wnc",
  1:lo, "_het")

# We logarithmize the x-axis
par(mar = c(0.5, 0.3,
  0.5, 0.3))
plot(c(1, 2500), c(0,
  1), log = "x", type = "n",
  xaxt = "n", yaxt = "n",
  xlab = "", ylab = "",
  xaxs = "i", yaxs = "i")
abline(v = 100, lty = "dashed",
  lwd = 0.5)
# If it's the first 'row' of plots,
# add info on the number of loci
if (first_row) {
  if (lo == 10) {
    first_row <- F
  }
  par(xpd = NA)
  mtext(paste(lo, " Loci"),
    3, line = 0.5)
  par(xpd = F)
}
# If this scenario led to at least
# one run where the mutation
# survived:
if (nrow(scenario_simulation) >
  0) {

  # Run through single simulations,
  # calculate allele frequency for the
  # colour locus and average allele
  # frequency at the attraction loci
  # (first for novel, then for
  # ancestral pattern)
  all_freq_patt <- matrix(ncol = nrow(scenario_simulation),
    nrow = 2500)
  all_freq_attr_nov <- matrix(ncol = nrow(scenario_simulation),
    nrow = 2500)
  all_freq_attr_anc <- matrix(ncol = nrow(scenario_simulation),
    nrow = 2500)
  for (setto in 1:nrow(scenario_simulation)) {
    matchers <- (1:nrow(get(paste0("l",
      lo))))[get(paste0("l",
      lo))$Scenario ==
      scenario_simulation$Scenario[setto] &
      get(paste0("l",
```

```
    lo))$Simulation ==
    scenario_simulation$Simulation[setto]]
all_freq_patt[,
  setto] <- (2 *
  get(paste0("l",
    lo))$Cc_hom[matchers] +
  2 * get(paste0("l",
    lo))$Cnc_hom[matchers] +
  get(paste0("l",
    lo))$Cc_het[matchers] +
  get(paste0("l",
    lo))$Cnc_het[matchers])/(2 *
  get(paste0("l",
    lo))$N_non_central[matchers] +
  2 * get(paste0("l",
    lo))$N_central[matchers])
all_freq_attr_nov[,
  setto] <- (2 *
  rowSums(get(paste0("l",
    lo))[matchers,
    novel_attr_c_hom,
    drop = F]) +
  2 * rowSums(get(paste0("l",
    lo))[matchers,
    novel_attr_nc_hom,
    drop = F]) +
  rowSums(get(paste0("l",
    lo))[matchers,
    novel_attr_c_het,
    drop = F]) +
  rowSums(get(paste0("l",
    lo))[matchers,
    novel_attr_nc_het,
    drop = F]))/(2 *
  lo * get(paste0("l",
  lo))$N_non_central[matchers] +
  2 * lo * get(paste0("l",
    lo))$N_central[matchers])
all_freq_attr_anc[,
  setto] <- (2 *
  rowSums(get(paste0("l",
    lo))[matchers,
    ances_attr_c_hom,
    drop = F]) +
  2 * rowSums(get(paste0("l",
    lo))[matchers,
    ances_attr_nc_hom,
    drop = F]) +
  rowSums(get(paste0("l",
    lo))[matchers,
    ances_attr_c_het,
    drop = F]) +
  rowSums(get(paste0("l",
```

```
          lo))[matchers,
          ances_attr_nc_het,
          drop = F]))/(2 *
        lo * get(paste0("l",
        lo))$N_non_central[matchers] +
        2 * lo * get(paste0("l",
          lo))$N_central[matchers])
    }
    # Add as partly transparent lines to
    # graph Note that we plot
    # 1-frequency of alleles that code
    # for increased attraction to the
    # ancestral pattern
    invisible(sapply(1:nrow(scenario_simulation),
      function(x) add_lines(all_mat = list(all_freq_patt,
        all_freq_attr_nov,
        1 - all_freq_attr_anc),
        x = x, n_mat = 3,
        cols = c("gray30",
          rgb(255/255,
            149/255, 8/255),
          rgb(28/255,
            57/255, 252/255)))))
  }
  # Add y-axis
  if (lo == 1) {
    axis(2, at = seq(0,
      1, 0.25), labels = F,
      tck = -0.015)
    axis(2, at = seq(0,
      1, 0.25), lwd = 0,
      line = -0.6, las = 2)
  }
  if (lo == 10) {
    axis(4, at = seq(0,
      1, 0.25), labels = F,
      tck = -0.015)
    axis(4, at = seq(0,
      1, 0.25), lwd = 0,
      line = -0.6, las = 2)
  }
  # Add x-axis
  if (sceni == length(all_scen) &
    sex_mutant == 1) {
    axis(1, at = c(5,
      20, 100, 500, 2500),
      labels = F, tck = -0.015)
    axis(1, at = c(5,
      20, 100, 500, 2500),
      lwd = 0, line = -0.8)
    if (lo == 5) {
      mtext("Generation",
        1, line = 1.8)
```

```
                    }
                }
            }
        }
    }
}
invisible(dev.off())
```

Show png we just made:

```
fig_disp <- readPNG("Figures/simulations_allele_freq_scenarios.png")
grid.raster(fig_disp)
```

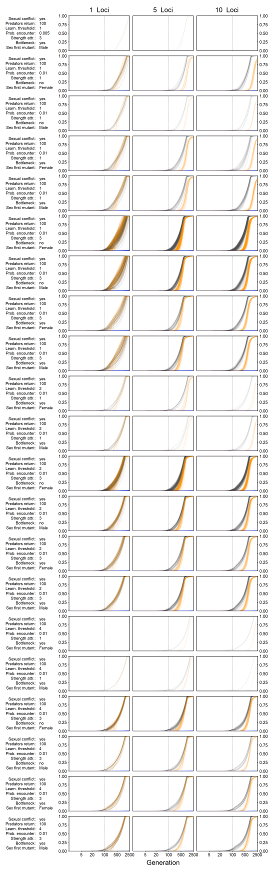


**3.3.1.4 Line plots of 'Phenotype Ratios'** In this section, we transcribe information from the above plot into phenotype ratios, where we'll show A) how is the ratio of novel over ancestral pattern and B) what is the ratio of overall male attraction score to novel over ancestral pattern.

First, we calculate these ratios:

```
# Empty lists to fill
ratio_pattern_all <- list()
ratio_attraction_all <- list()
yrange_pattern_all <- list()
yrange_attraction_all <- list()
mult_f_all <- list()
first_gen_pattern_1_crossed_all <- list()
first_gen_attr_approach_1_all <- list()
```

```
# Calculate phenotype ratios
for (sceni in 1:length(all_scen)) {
    # Extract current scenario
    scenario <- all_scen[sceni]

    # For each scenario, go through the
    # two sexes
    for (sex_mutant in 0:1) {
        # If the scenario and this sex of
        # the first mutant ever led to
        # survival of the novel allele:
        if (all_scen_mf[sex_mutant +
            1, sceni] > 0) {

            # Each row of the plotting scheme
            # receives one plot for each set of
            # loci
            for (lo in loci) {

                # Extract results for current
                # scenario
                scenario_simulation <- get(paste0("l",
                  lo, "_2500"))[get(paste0("l",
                  lo, "_2500"))$Scenario ==
                  scenario & get(paste0("l",
                  lo, "_2500"))$sex_mutant ==
                  sex_mutant, c("Scenario",
                  "Simulation")]
                # Get column names for preference
                # loci
                novel_attr_c_hom <- paste0("Rc",
                  1:lo, "_hom")
                novel_attr_nc_hom <- paste0("Rnc",
                  1:lo, "_hom")
                novel_attr_c_het <- paste0("Rc",
                  1:lo, "_het")
                novel_attr_nc_het <- paste0("Rnc",
                  1:lo, "_het")
                ances_attr_c_hom <- paste0("Wc",
                  1:lo, "_hom")
                ances_attr_nc_hom <- paste0("Wnc",
                  1:lo, "_hom")
                ances_attr_c_het <- paste0("Wc",
                  1:lo, "_het")
                ances_attr_nc_het <- paste0("Wnc",
                  1:lo, "_het")

                # If this scenario led to at least
                # one run where the mutation
                # survived:
                if (nrow(scenario_simulation) >
                  0) {
```

```r
# Calculate the phenotype ratios
ratio_pattern <- matrix(ncol = nrow(scenario_simulation),
  nrow = 2500)
ratio_attraction <- matrix(ncol = nrow(scenario_simulation),
  nrow = 2500)
for (setto in 1:nrow(scenario_simulation)) {
  # Get matching rows
  matchers <- (1:nrow(get(paste0("l",
    lo))))[get(paste0("l",
    lo))$Scenario ==
    scenario_simulation$Scenario[setto] &
    get(paste0("l",
      lo))$Simulation ==
      scenario_simulation$Simulation[setto]]
  # Pattern ratio
  ratio_pattern[,
    setto] <- (get(paste0("l",
    lo))$Cnc_hom[matchers] +
    get(paste0("l",
      lo))$Cc_hom[matchers] +
    get(paste0("l",
      lo))$Cnc_het[matchers] +
    get(paste0("l",
      lo))$Cc_het[matchers])/((get(paste0("l",
    lo))$N_non_central[matchers] +
    get(paste0("l",
      lo))$N_central[matchers]) -
    (get(paste0("l",
      lo))$Cnc_hom[matchers] +
      get(paste0("l",
        lo))$Cc_hom[matchers] +
      get(paste0("l",
        lo))$Cnc_het[matchers] +
      get(paste0("l",
        lo))$Cc_het[matchers]))
  # Total number of individuals
  number_indiv <- get(paste0("l",
    lo))$N_non_central[matchers] +
    get(paste0("l",
      lo))$N_central[matchers]
  # Attraction ratios As novel
  # mutations are dominant,
  # heterozygotes are equal to
  # homzygotes here.
  novel_attr_m <- sapply(1:lo,
    function(x) rowSums(get(paste0("l",
      lo))[matchers,
      c(novel_attr_nc_het[x],
        novel_attr_nc_hom[x],
        novel_attr_c_het[x],
        novel_attr_c_hom[x])])/number_indiv)
  ancestr_attr_m <- sapply(1:lo,
    function(x) rowSums(get(paste0("l",
```

```r
        lo))[matchers,
        c(ances_attr_nc_het[x],
          ances_attr_nc_hom[x],
          ances_attr_c_het[x],
          ances_attr_c_hom[x])])/number_indiv)
    # We use the same formula as in the
    # simulations to calculate an
    # attraction score
    attraction_novel <- exp(param$strength_attract[scenario] *
      ((rowSums(novel_attr_m)/lo) -
        1/2))/exp(param$strength_attract[scenario] *
      (1/2))
    attraction_ancestr <- exp(param$strength_attract[scenario] *
      ((rowSums(ancestr_attr_m)/lo) -
        1/2))/exp(param$strength_attract[scenario] *
      (1/2))
    ratio_attraction[,
      setto] <- (attraction_novel/attraction_ancestr)
  }

  # Get ranges of values
  yvals_pattern <- c(as.vector(log(ratio_pattern)))
  yrange_pattern <- max(abs(yvals_pattern[!is.infinite(yvals_pattern)]))
  yvals_attraction <- c(as.vector(log(ratio_attraction)))
  # Split this into two lines for
  # Markdown aesthetics
  infcheck <- !is.infinite(yvals_attraction)
  yrange_attraction <- max(abs(yvals_attraction[infcheck]))
  # The ratio between pattern and
  # attraction will be used later to
  # scale one to the other (as we want
  # both ratios in one plot, but their
  # scaling is very different)
  mult_f <- yrange_pattern/yrange_attraction

  # Calculate when certain events took
  # place, which we'll use later as a
  # background in the plot.  First
  # generation the pattern ratio
  # crosses 1:1
  idp <- (1:nrow(ratio_pattern))
  first_gen_pattern_1_crossed <- idp[rowMeans(ratio_pattern) >
    1][1]
  # First generation the attraction
  # ratio approaches 1 (i.e. 0.99)
  ida <- (1:nrow(ratio_attraction))
  first_gen_attr_approach_1 <- ida[rowMeans(ratio_attraction) >
    0.99][1]

  # Store results
  ratio_pattern_all[[length(ratio_pattern_all) +
    1]] <- ratio_pattern
  ratio_attraction_all[[length(ratio_attraction_all) +
```

```
                  1]] <- ratio_attraction
                yrange_pattern_all[[length(yrange_pattern_all) +
                  1]] <- yrange_pattern
                yrange_attraction_all[[length(yrange_attraction_all) +
                  1]] <- yrange_attraction
                mult_f_all[[length(mult_f_all) +
                  1]] <- mult_f
                first_gen_pattern_1_crossed_all[[length(first_gen_pattern_1_crossed_all) +
                  1]] <- first_gen_pattern_1_crossed
                first_gen_attr_approach_1_all[[length(first_gen_attr_approach_1_all) +
                  1]] <- first_gen_attr_approach_1
              } else {
                # If no simulation run resulted in
                # survival, enter NA
                ratio_pattern_all[[length(ratio_pattern_all) +
                  1]] <- NA
                ratio_attraction_all[[length(ratio_attraction_all) +
                  1]] <- NA
                yrange_pattern_all[[length(yrange_pattern_all) +
                  1]] <- NA
                yrange_attraction_all[[length(yrange_attraction_all) +
                  1]] <- NA
                mult_f_all[[length(mult_f_all) +
                  1]] <- NA
                first_gen_pattern_1_crossed_all[[length(first_gen_pattern_1_crossed_all) +
                  1]] <- NA
                first_gen_attr_approach_1_all[[length(first_gen_attr_approach_1_all) +
                  1]] <- NA
              }
          }
      }
    }
}
```

Get most extreme y-value ranges from all scenarios

```
yranges_all <- c(-max(unlist(yrange_pattern_all),
    unlist(yrange_attraction_all), na.rm = T),
    max(unlist(yrange_pattern_all),
        unlist(yrange_attraction_all),
        na.rm = T))
```

For the plot, we follow a very similar logic as in the previous one. Important differences will be pointed out.

```
first_row <- T
png("Figures/simulations_phenotype_rat_scenarios.png",
    width = 2000, height = 6500, res = 300)

layout(matrix(1:number_of_plots, ncol = 4,
    byrow = T))
par(oma = c(3, 0, 1.55, 2))

curr_count <- 0
for (sceni in 1:length(all_scen)) {
    scenario <- all_scen[sceni]
```

```
    for (sex_mutant in 0:1) {
        if (all_scen_mf[sex_mutant +
            1, sceni] > 0) {
            par(mar = c(0.5, 0.3, 0.5,
                1.8))
            plot(1, 1, type = "n", xaxt = "n",
                yaxt = "n", xlab = "",
                ylab = "", bty = "n",
                ylim = c(0.1, 0.9),
                xlim = c(0, 2), xaxs = "i")
            texti <- param[all_scen[sceni],
                ]
            text(1.3, 0.86, "Sexual conflict:   ",
                pos = 2, offset = 0,
                cex = 0.95)
            text(1.3, 0.86, c("no",
                "yes")[texti$sex_conflict +
                1], pos = 4, offset = 0,
                cex = 0.95)
            text(1.3, 0.74, "Predators return:   ",
                pos = 2, offset = 0,
                cex = 0.95)
            text(1.3, 0.74, texti$predator_absence,
                pos = 4, offset = 0,
                cex = 0.95)
            text(1.3, 0.62, "Learn. threshold:   ",
                pos = 2, offset = 0,
                cex = 0.95)
            text(1.3, 0.62, texti$pred_constant,
                pos = 4, offset = 0,
                cex = 0.95)
            text(1.3, 0.5, "Prob. encounter:   ",
                pos = 2, offset = 0,
                cex = 0.95)
            text(1.3, 0.5, texti$prob_encounter,
                pos = 4, offset = 0,
                cex = 0.95)
            text(1.3, 0.38, "Strength attr.:   ",
                pos = 2, offset = 0,
                cex = 0.95)
            text(1.3, 0.38, texti$strength_attract,
                pos = 4, offset = 0,
                cex = 0.95)
            text(1.3, 0.26, "Bottleneck:   ",
                pos = 2, offset = 0,
                cex = 0.95)
            text(1.3, 0.26, c("no",
                "yes")[texti$bottleneck +
                1], pos = 4, offset = 0,
                cex = 0.95)
            text(1.3, 0.14, "Sex first mutant:   ",
                pos = 2, offset = 0,
                cex = 0.95)
```

```r
text(1.3, 0.14, c("Female",
    "Male")[sex_mutant +
    1], pos = 4, offset = 0,
    cex = 0.95)

for (lo in loci) {

    scenario_simulation <- get(paste0("l",
      lo, "_2500"))[get(paste0("l",
      lo, "_2500"))$Scenario ==
      scenario & get(paste0("l",
      lo, "_2500"))$sex_mutant ==
      sex_mutant, c("Scenario",
      "Simulation")]

    curr_count <- curr_count +
      1
    # We logarithmize the x- and y-axis
    par(mar = c(0.5, 0.3,
      0.5, 0.3))
    plot(1, 1, type = "n",
      ylim = yranges_all,
      xlim = c(1, 2500),
      log = "x", xlab = "",
      ylab = "", xaxt = "n",
      yaxt = "n", xaxs = "i",
      yaxs = "i")
    # Add horizontal lines for ratios of
    # 1:1
    abline(h = 0, lty = "dashed",
      lwd = 0.75)

    if (nrow(scenario_simulation) >
      0) {
      # Highlight the four phases by
      # drawing 4 polygons, separated by 3
      # lines. Those lines (=boundaries of
      # polygons) are at: 100 (last
      # generation of relaxed predation),
      # first_gen_pattern_1_crossed (first
      # generation where average ratio for
      # pattern corsses 1:1) and
      # first_gen_attr_approach_1 (first
      # generation where average ratio for
      # attraction is bigger than 0.99).
      polygon(c(10^-10,
        rep(100, 2), 10^-10),
        rep(c(-10^10, 10^10),
          each = 2), border = NA,
        col = adjustcolor("goldenrod",
          0.08))
      polygon(c(100, rep(first_gen_pattern_1_crossed_all[[curr_count]],
        2), 100), rep(c(-10^10,
```

```
      10^10), each = 2),
      border = NA, col = adjustcolor("goldenrod",
        0.16))
    if (is.na(first_gen_attr_approach_1_all[[curr_count]])) {
      polygon(c(first_gen_pattern_1_crossed_all[[curr_count]],
        rep(10^10, 2),
        first_gen_pattern_1_crossed_all[[curr_count]]),
        rep(c(-10^10,
          10^10), each = 2),
        border = NA, col = adjustcolor("goldenrod",
          0.24))
    } else {
      polygon(c(first_gen_pattern_1_crossed_all[[curr_count]],
        rep(first_gen_attr_approach_1_all[[curr_count]],
          2), first_gen_pattern_1_crossed_all[[curr_count]]),
        rep(c(-10^10,
          10^10), each = 2),
        border = NA, col = adjustcolor("goldenrod",
          0.24))
      polygon(c(first_gen_attr_approach_1_all[[curr_count]],
        rep(10^10, 2),
        first_gen_attr_approach_1_all[[curr_count]]),
        rep(c(-10^10,
          10^10), each = 2),
        border = NA, col = adjustcolor("goldenrod",
          0.32))
      abline(v = first_gen_attr_approach_1_all[[curr_count]],
        lty = "dashed",
        lwd = 0.5)
    }
    abline(v = c(100,
      first_gen_pattern_1_crossed_all[[curr_count]]),
      lty = "dashed",
      lwd = 0.5)
  }
  if (first_row) {
    if (lo == 10) {
      first_row <- F
    }
    par(xpd = NA)
    mtext(paste(lo, " Loci"),
      3, line = 0.5)
    par(xpd = F)
  }
  if (nrow(scenario_simulation) >
    0) {

    # Add phenotype ratios as lines
    # (note that all are transformed to
    # the logarithm). In gray are the
    # pattern ratios, in green the
    # attraction ratios. Note that the
    # attraction ratios are multiplied
```

```
    # by the minimum ratio of pattern
    # divided by attraction ratio
    invisible(sapply(1:nrow(scenario_simulation),
      function(x) add_lines(all_mat = list(log(ratio_pattern_all[[curr_count]]),
        min(unlist(mult_f_all),
          na.rm = T) *
          log(ratio_attraction_all[[curr_count]])),
        x = x, n_mat = 2,
        cols = c("gray30",
          rgb(23/255,
            179/255, 152/255)))))
  }
  # Add y-axis as ratios
  if (lo == 1) {
    axis(2, at = log(c(10^-4.5,
      10^-2, 1, 10^2,
      10^4.5)), labels = F,
      tck = -0.015, col = NA,
      col.ticks = "gray30")
    axis(2, at = log(c(10^-4.5,
      10^-2, 1, 10^2,
      10^4.5)), parse(text = c(paste0("10^",
      c("-4.5", "-2")),
      "1", paste0("10^",
        c("2", "4.5")))),
      lwd = 0, line = -0.7,
      las = 2, col.axis = "gray30",
      cex.axis = 0.85)
  }
  if (lo == 10) {
    # Right y-axis: First calculate the
    # position of those labels we want
    # to show (scaling to left y-axis)
    label_pos <- log(c(1/20,
      1/5, 1, 5/1, 20/1)) *
      min(unlist(mult_f_all),
        na.rm = T)
    axis(4, at = label_pos,
      labels = F, tck = -0.015,
      col = NA, col.ticks = rgb(23/255,
        179/255, 152/255))
    axis(4, at = label_pos,
      c("1/20", "1/5",
        "1", "5", "20"),
      lwd = 0, line = -0.7,
      las = 2, col.axis = rgb(23/255,
        179/255, 152/255),
      cex.axis = 0.85)

  }

  if (sceni == length(all_scen) &
    sex_mutant == 1) {
```

```
         axis(1, at = c(5,
          20, 100, 500, 2500),
          labels = F, tck = -0.015)
         axis(1, at = c(5,
          20, 100, 500, 2500),
          lwd = 0, line = -0.8)
         if (lo == 5) {
          mtext("Generation",
           1, line = 1.8)
         }
        }
      }
    }
  }
}
invisible(dev.off())
```

Show png we just made:

```
fig_disp <- readPNG("Figures/simulations_phenotype_rat_scenarios.png")
grid.raster(fig_disp)
```

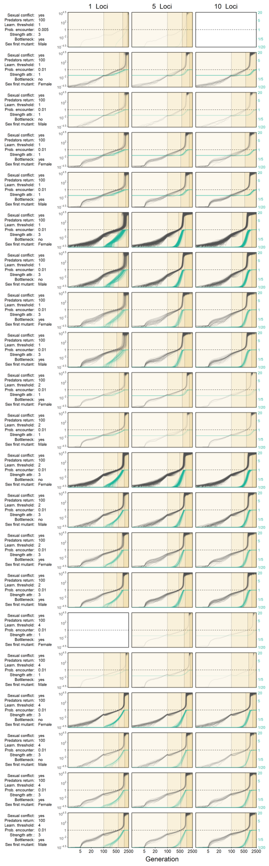


**3.3.1.5 Allele frequency Gif** We'll produce an animation video for one simulation run from a specific scenario (sexual conflict present, predators absent from central patches for 100 generations, no bottleneck, 10 loci coding for attraction phenotypes probability of encounter = 0.01, strength of attraction = 3, predator learning threshold = 2).

```
scenario <- (1:nrow(param))[param$sex_conflict ==
    1 & param$bottleneck == 0 & param$prob_encounter ==
    0.01 & param$predator_absence ==
    100 & param$strength_attract ==
    3 & param$pred_constant == 2]
```

First, general setup for simulations (explanations are almost the same as in the beginning of the Markdown, where we showed how to run the simulations).

```
# Arena dimensions
arena_width <- 4L
arena_height <- 96L
# Number of central patches (in
# vertical direction (i.e. patches
# 39-58 when 96 in height))
arena_center <- 20L
# Generation when the birds
# disappear
birds_die <- 1L
# Number of individuals per colour
# phenotype picked out during
# relaxed predation
reduced_predation <- 0L
# Generation in which the bottleneck
# in the central patches ends (i.e.
# Passiflora plants are available
# again and egg laying goes back to
# normal)
bottleneck_ends <- 6L
# Maximum age of individuals of
# starting population
max_age <- 5L
# All females in starting population
# are mated
females_mated <- T
# Mutation rate per individual per
# locus per generation
prob_mutation <- 10^-5
# Maximum number of females a male
# can mate during one generation
max_mate <- 4
# Maximum number of generations per
# simulation
generations <- 2500
```

We search for simulation runs with same starting seed, in which the first mutant is placed in the exact same spot and all other starting conditions are equal.

```
seed_equal <- table(c(l1_2500$start_seed_of_simulation,
    l5_2500$start_seed_of_simulation,
    l10_2500$start_seed_of_simulation))[table(c(l1_2500$start_seed_of_simulation,
    l5_2500$start_seed_of_simulation,
    l10_2500$start_seed_of_simulation)) ==
    3]
```

Get those from scenario 39 (the one we selected above)

```
scen39 <- as.numeric(names(seed_equal)[as.numeric(names(seed_equal)) >
    389999 & as.numeric(names(seed_equal)) <
    4e+05])
# Check if the sex and placement of
# the first mutants is indeed
# idential:
(sex_patch <- data.frame(seed = scen39,
    sex_l1 = sapply(scen39, function(x) l1_2500$sex_mutant[l1_2500$start_seed_of_simulation ==
        x]), sex_l5 = sapply(scen39,
        function(x) l5_2500$sex_mutant[l5_2500$start_seed_of_simulation ==
            x]), sex_l10 = sapply(scen39,
        function(x) l10_2500$sex_mutant[l10_2500$start_seed_of_simulation ==
            x]), patch_l1 = sapply(scen39,
        function(x) l1_2500$mutator_patch[l1_2500$start_seed_of_simulation ==
            x]), patch_l5 = sapply(scen39,
        function(x) l5_2500$mutator_patch[l5_2500$start_seed_of_simulation ==
            x]), patch_l10 = sapply(scen39,
        function(x) l10_2500$mutator_patch[l10_2500$start_seed_of_simulation ==
            x])))
```

```
# Yes! Now take the first row with a
# female first mutant
seed_for_simul <- sex_patch$seed[rowSums(sex_patch[,
    2:4]) == 0][1]
```

Set parameters specific to scenario:

```
# Whether we start off from a
# bottleneck population
bottleneck <- param$bottleneck[scenario]
# Sexual conflict (yes or no)
sex_conflict <- param$sex_conflict[scenario]
# Probability that a female
# encounters a male during egg
# laying Only relevant if
# sex_conflict==TRUE
prob_encounter <- param$prob_encounter[scenario]
# Number generations no predation
birds_return <- param$predator_absence[scenario]
# alpha in Duenez-Guzman et al.
# (2009), for attraction of a male
# to certain colour A value of 1
# means that a male with all
# attraction loci for colour X at Aa
# or AA state and all attraction
# loci for colour Y at aa state will
# have a preference index for X of
# about 0.73 (where preference index
# is (associations with X)/(all
# associations)).  A value of 1
# means that the same male will have
# a preference index of about 0.95
strength_of_attraction <- param$strength_attract[scenario]
# From this, calculate maximum
```

```r
# attraction value
max_attr <- exp(strength_of_attraction *
    (1 - 1/2) * 1)
# Number of individuals per colour
# phenotype picked out by predators
pred_constant <- param$pred_constant[scenario]
```

From this, deduce following other constants:

```r
# Total arena patches
patches <- arena_width * arena_height
# Get central patches (if they
# cannot be perfectly centered,
# round up)
central_vertical <- (1:patches)[((1:patches -
    1)%%arena_height + 1) %in% ifelse(arena_height%%2 ==
    0, ceiling(0.5 * arena_height -
    0.5 * arena_center + 1), ceiling(0.5 *
    (arena_height + 1) - 0.5 * (arena_center -
    1))):ifelse(arena_height%%2 == 0,
    ceiling(0.5 * arena_height + 0.5 *
        arena_center), ceiling(0.5 *
        (arena_height + 1) + 0.5 * (arena_center -
        1)))]
# Get non-central patches
not_central <- (1:patches)[!(1:patches) %in%
    central_vertical]
```

Create lists to store the allele frequencies from the different generations. Each list will contain three matrices (one for each set of loci). One matrix will have 384 columns (one for each patch) and 2500 rows (one for each generation). Also, we store a list to save the position of the first mutant.

```r
results_per_loci_colour <- list()
results_per_loci_pref_novel <- list()
results_per_loci_pref_ances <- list()
all_mut_patches <- list()
```

Now come the actual simulations. Iterate through genetic architectures.

```r
for (number_loci in loci) {

    # Setup population

    # Remember total number of columns
    # for one of the tables from the
    # list
    number_columns <- 6 + number_loci *
        4

    # Indices for function (loci_indices
    # is a function defined in
    # Functions.R)
    indices <- loci_indices(number_loci)

    # Results matrices (for allele
    # frequencies)
```

```r
freq_matrix_colour <- matrix(ncol = patches,
    nrow = 0)
freq_matrix_pref_novel <- matrix(ncol = patches,
    nrow = 0)
freq_matrix_pref_ances <- matrix(ncol = patches,
    nrow = 0)

# Set start seed
set.seed(seed_for_simul)

# Make new individuals and introduce
# mutation at colour pattern locus

# Get starting population
individuals <- beginners(number_per_central = 114,
    number_per_rest = 114, females_mated = females_mated,
    patches = patches, number_loci = number_loci)
# Place a mutation at the colour
# locus to turn one new-born female
# from a central patch into a
# novel-coloured butterfly
mutator_patch <- sample(central_vertical,
    size = 1)
all_mut_patches[[length(all_mut_patches) +
    1]] <- mutator_patch
mutator_indiv <- sample.int(114,
    size = 1)
individuals[[mutator_patch]][mutator_indiv,
    5] <- 1L
# Make that individual age 0
# (new-born in previous generation)
individuals[[mutator_patch]][mutator_indiv,
    2] <- 0L

# Run through generations
for (n in as.integer(1:generations)) {

    # Run simulation
    for (unito in 1:patches) {
        individuals[[unito]] <- life_cycle(individuals[[unito]],
            n = unito)
    }

    # Change list position of migrators
    # (make them 'officially' migrate);
    # this has to happen in a separate
    # step, such that the migrators only
    # appear in their new patch in the
    # next loop
    for (unito in 1:patches) {
        curr_n <- nrow(individuals[[unito]])
        if (curr_n > 0) {
            changers <- (1:curr_n)[individuals[[unito]][,
```

```
            3] != unito]
          if (length(changers) >
            0) {
            for (move_around in changers) {
              individuals[[individuals[[unito]][move_around,
                3]]] <- rbind(individuals[[individuals[[unito]][move_around,
                3]]], individuals[[unito]][move_around,
                , drop = F])
            }
            individuals[[unito]] <- individuals[[unito]][-changers,
              , drop = F]
          }
      }
  }

  # Collect data

  # Total number of individuals
  total_individuals <- lengths(individuals)/number_columns
  # This counts number of
  # heterozygotes and homozygotes
  # separately for each patch, and at
  # each locus position
  freqs_hetero <- lapply(individuals,
      function(x) colSums(x[,
          indices$i_original,
          drop = F] == 1L))
  freqs_homo <- lapply(individuals,
      function(x) colSums(x[,
          indices$i_original,
          drop = F] == 2L))
  # Append frequencies to matrix
  freq_matrix_colour <- rbind(freq_matrix_colour,
      (as.integer(sapply(1:length(freqs_homo),
          function(x) freqs_hetero[[x]][1])) +
          2 * as.integer(sapply(1:length(freqs_homo),
            function(x) freqs_homo[[x]][1])))/(2 *
          total_individuals))
  freq_matrix_pref_novel <- rbind(freq_matrix_pref_novel,
      (as.integer(sapply(1:length(freqs_homo),
          function(x) mean(freqs_hetero[[x]][2:(number_loci +
            1)]))) + 2 * as.integer(sapply(1:length(freqs_homo),
          function(x) mean(freqs_homo[[x]][2:(number_loci +
            1)]))))/(2 * total_individuals))
  freq_matrix_pref_ances <- rbind(freq_matrix_pref_ances,
      1 - ((as.integer(sapply(1:length(freqs_homo),
          function(x) mean(freqs_hetero[[x]][(number_loci +
            2):(2 * number_loci +
            1)]))) + 2 * as.integer(sapply(1:length(freqs_homo),
          function(x) mean(freqs_homo[[x]][(number_loci +
            2):(2 * number_loci +
            1)]))))/(2 * total_individuals)))
```

```
        # If novel mutation dies out, stop
        # simulations (never the case in
        # this example, as we know these
        # simulations will reach generation
        # 2500)
        if (sum(freq_matrix_colour[nrow(freq_matrix_colour),
            ]) == 0)
            break
    }

    # Append results to list
    results_per_loci_colour[[length(results_per_loci_colour) +
        1]] <- freq_matrix_colour
    results_per_loci_pref_novel[[length(results_per_loci_pref_novel) +
        1]] <- freq_matrix_pref_novel
    results_per_loci_pref_ances[[length(results_per_loci_pref_ances) +
        1]] <- freq_matrix_pref_ances
}
```

Make the movie!

Construct a matrix with x and y coordinates of the patches

```
patchmat <- cbind(rep(1:4, each = 96),
    rep(96:1, 4))
```

Define range for colour palette

```
col_rang <- 0:1
# Define colour ramp
paletto <- colorRampPalette(c("blue",
    "red"))
# Sample 1000 colours
bluered <- paletto(1000)
```

Go through all 2500 generations

```
for (cycler in 1:2500) {
    # Create a new png, with a layout to
    # contain 9 plots
    png(paste0("movie/img", ifelse(cycler <
        10, "000", ifelse(cycler < 100,
        "00", ifelse(cycler < 1000,
            "0", ""))), cycler, ".png"),
        width = 1500, height = 2500,
        res = 300)
    layout(matrix(1:9, nrow = 3))
    par(oma = c(0, 1.5, 3, 0))
    # This identifier describes at which
    # of three lists we look
    for (loci_set in 1:3) {
        # This identifier described within
        # each list which matrix we consider
        for (type_loci in 1:3) {
            # Get current matrix
            curro <- get(paste0("results_per_loci_",
```

```r
    c("colour", "pref_novel",
      "pref_ances")[type_loci]))[[loci_set]]
# Open empty plot fit to the arena
# dimensions
par(mar = c(0.25, 0.25,
    0.25, 0.25))
plot(1, 1, type = "n", xaxt = "n",
    yaxt = "n", xlab = "",
    ylab = "", xaxs = "i",
    yaxs = "i", xlim = c(0.5,
      4.5), ylim = c(0.5,
      96.5))
# Assign a colour to each
# polygon/patch -> blue - frequency
# = 0; -> red - frequency = 1
col_dots <- bluered[sapply(1:patches,
    function(x) which.min(abs(seq(col_rang[1],
      col_rang[2], (1/999)) -
      curro[cycler, x])))]
# Plot the separate patches
invisible(sapply(1:patches,
    function(x) polygon(c(patchmat[x,
      1] - 0.5, patchmat[x,
      1] + 0.5, patchmat[x,
      1] + 0.5, patchmat[x,
      1] - 0.5), c(patchmat[x,
      2] - 0.5, patchmat[x,
      2] - 0.5, patchmat[x,
      2] + 0.5, patchmat[x,
      2] + 0.5), border = NA,
      col = col_dots[x])))
# Denote the 'central' region where
# predation is initially relaxed
abline(h = c(38.5, 58.5),
    col = "white")
# Highlight in yellow where first
# mutant occurred
polygon(c(patchmat[all_mut_patches[[loci_set]],
    1] - 0.5, patchmat[all_mut_patches[[loci_set]],
    1] + 0.5, patchmat[all_mut_patches[[loci_set]],
    1] + 0.5, patchmat[all_mut_patches[[loci_set]],
    1] - 0.5), c(patchmat[all_mut_patches[[loci_set]],
    2] - 0.5, patchmat[all_mut_patches[[loci_set]],
    2] - 0.5, patchmat[all_mut_patches[[loci_set]],
    2] + 0.5, patchmat[all_mut_patches[[loci_set]],
    2] + 0.5), border = "yellow",
    lwd = 0.7, col = NA)
# Add titles
if (type_loci == 1) {
    mtext(c("1 Loci", "5 Loci",
      "10 Loci")[loci_set],
      3, line = 0.3)
}
```

```
            if (loci_set == 1) {
                mtext(c("Colour", "Pref. novel",
                  "Pref. ancestral")[type_loci],
                  2, line = 0.3)
                if (type_loci == 1) {
                  mtext(paste0("Gen.: ",
                    ifelse(cycler <
                      10, "000", ifelse(cycler <
                      100, "00", ifelse(cycler <
                      1000, "0", ""))),
                    cycler), 3, line = 2)
                }
            }
            # Add info about important events
            if (loci_set == 3) {
                if (type_loci == 1) {
                  par(xpd = NA)
                  text(par("usr")[2],
                    par("usr")[4] +
                      0.15 * diff(par("usr")[3:4]),
                    paste0(ifelse(cycler >
                      5, "First Mutant died",
                      "")), cex = 0.85,
                    pos = 2, offset = 0)
                  text(par("usr")[2],
                    par("usr")[4] +
                      0.105 * diff(par("usr")[3:4]),
                    paste0(ifelse(cycler >
                      100, "Predators returned",
                      "")), cex = 0.85,
                    pos = 2, offset = 0)
                  par(xpd = F)
                }
            }
        }
    }
    # mtext('Freq. 'Red'
    # Allels',3,line=-1,outer=T)
    invisible(dev.off())
}
```

Create the movie:

```
# This requires a valid build of
# ffmpeg, set as a system path Cut
# code into pieces for Markdown
# formatting
txt1 <- "ffmpeg -r 25 -i movie/img%04d.png -c:v libx264 -vf fps=25"
txt2 <- " -pix_fmt yuv420p movie/allele_freq_over_time_example.mp4"
system(paste0(txt1, txt2))
# Delete all pngs
invisible(file.remove(list.files("movie/",
    pattern = ".png", full.names = T)))
```

###### 3.3.1.6 The importance of allele frequencies at generation 100

A decisive moment seems to occur at generation 100 (return of predators to the central patches of the arena), at which the frequency dynamics seem to change a lot. We want to investigate whether the allele frequency (or the phenotype frequency) for the novel pattern at generation 100 is a good predictor for simulation outcome.

We calculate the allele and the phenotype frequency of the novel pattern at generation 100, for the three different datasets:

```
all_freq_c_100_l1 <- (l1$Cc_het[l1$Generation ==
    100] + 2 * l1$Cc_hom[l1$Generation ==
    100])/(2 * l1$N_central[l1$Generation ==
    100])
all_freq_c_100_l5 <- (l5$Cc_het[l5$Generation ==
    100] + 2 * l5$Cc_hom[l5$Generation ==
    100])/(2 * l5$N_central[l5$Generation ==
    100])
all_freq_c_100_l10 <- (l10$Cc_het[l10$Generation ==
    100] + 2 * l10$Cc_hom[l10$Generation ==
    100])/(2 * l10$N_central[l10$Generation ==
    100])

all_freq_c_pheno_100_l1 <- (l1$Cc_het[l1$Generation ==
    100] + l1$Cc_hom[l1$Generation ==
    100])/(l1$N_central[l1$Generation ==
    100])
all_freq_c_pheno_100_l5 <- (l5$Cc_het[l5$Generation ==
    100] + l5$Cc_hom[l5$Generation ==
    100])/(l5$N_central[l5$Generation ==
    100])
all_freq_c_pheno_100_l10 <- (l10$Cc_het[l10$Generation ==
    100] + l10$Cc_hom[l10$Generation ==
    100])/(l10$N_central[l10$Generation ==
    100])
```

We plot these. As red dots, we plot those simulation runs, where the novel pattern persisted. First, determine which dot is black and which is red:

```
colo100_l1 <- ifelse(paste0(l1$Scenario[l1$Generation ==
    100], "_", l1$Simulation[l1$Generation ==
    100]) %in% paste0(l1_2500$Scenario,
    "_", l1_2500$Simulation), "red",
    ifelse(l1$Scenario[l1$Generation ==
        100] %in% (1:nrow(param))[param$sex_conflict ==
        1], "black", "grey"))
colo100_l5 <- ifelse(paste0(l5$Scenario[l5$Generation ==
    100], "_", l5$Simulation[l5$Generation ==
    100]) %in% paste0(l5_2500$Scenario,
    "_", l5_2500$Simulation), "red",
    ifelse(l5$Scenario[l5$Generation ==
        100] %in% (1:nrow(param))[param$sex_conflict ==
        1], "black", "grey"))
colo100_l10 <- ifelse(paste0(l10$Scenario[l10$Generation ==
    100], "_", l10$Simulation[l10$Generation ==
    100]) %in% paste0(l10_2500$Scenario,
    "_", l10_2500$Simulation), "red",
    ifelse(l10$Scenario[l10$Generation ==
```

```
        100] %in% (1:nrow(param))[param$sex_conflict ==
        1], "black", "grey"))
```

Now, we plot this:

```
png("Figures/simulations_freq_after_100.png",
    width = 3500, height = 1500, res = 300)
layout(matrix(1:2, ncol = 2))

# First, we look at the allele
# frequency
par(mar = c(0.4, 3.5, 0.4, 0.6))
plot(1, 1, type = "n", xaxt = "n", yaxt = "n",
    xlab = "", ylab = "", xlim = c(0.5,
        1.5), ylim = range(c(all_freq_c_100_l1,
        all_freq_c_100_l5, all_freq_c_100_l10)))
text(par("usr")[1] + (0.3/par("pin")[1]) *
    diff(par("usr")[1:2]), par("usr")[4] -
    (0.23/par("pin")[2]) * diff(par("usr")[3:4]),
    "A", font = 2, cex = 1.5, pos = 2,
    offset = 0)
# Beeswarm
beeswarm(c(all_freq_c_100_l1, all_freq_c_100_l5,
    all_freq_c_100_l10), at = 1, add = T,
    cex = 0.045, xaxt = "n", yaxt = "n",
    pwcol = c(colo100_l1, colo100_l5,
        colo100_l10))
# And the corresponding boxplot on
# top of it in blue
boxplot(c(all_freq_c_100_l1, all_freq_c_100_l5,
    all_freq_c_100_l10), at = 1, add = T,
    outline = F, xaxt = "n", yaxt = "n",
    border = "dodgerblue2", col = NA)
# Axes and titles
axis(2, labels = F, tck = -0.015)
axis(2, las = 2, line = -0.5, lwd = 0)
mtext("Frequency of novel colour pattern allele",
    2, line = 2.7, cex = 0.9)
mtext("in central patches at generation 100",
    2, line = 1.9, cex = 0.9)

# We repeat the same for phenotype
# frequency
plot(1, 1, type = "n", xaxt = "n", yaxt = "n",
    xlab = "", ylab = "", xlim = c(0.5,
        1.5), ylim = range(c(all_freq_c_pheno_100_l1,
        all_freq_c_pheno_100_l5, all_freq_c_pheno_100_l10)))
text(par("usr")[1] + (0.3/par("pin")[1]) *
    diff(par("usr")[1:2]), par("usr")[4] -
    (0.23/par("pin")[2]) * diff(par("usr")[3:4]),
    "B", font = 2, cex = 1.5, pos = 2,
    offset = 0)
beeswarm(c(all_freq_c_pheno_100_l1,
    all_freq_c_pheno_100_l5, all_freq_c_pheno_100_l10),
```

```
    at = 1, add = T, cex = 0.045, xaxt = "n",
    yaxt = "n", pwcol = c(colo100_l1,
        colo100_l5, colo100_l10))
boxplot(c(all_freq_c_pheno_100_l1, all_freq_c_pheno_100_l5,
    all_freq_c_pheno_100_l10), at = 1,
    add = T, outline = F, xaxt = "n",
    yaxt = "n", border = "dodgerblue2",
    col = NA)
axis(2, labels = F, tck = -0.015)
axis(2, las = 2, line = -0.5, lwd = 0)
mtext("Frequency of novel colour pattern phenotype",
    2, line = 2.7, cex = 0.9)
mtext("in central patches at generation 100",
    2, line = 1.9, cex = 0.9)

invisible(dev.off())
```

Show png we just made:

```
fig_disp <- readPNG("Figures/simulations_freq_after_100.png")
grid.raster(fig_disp)
```

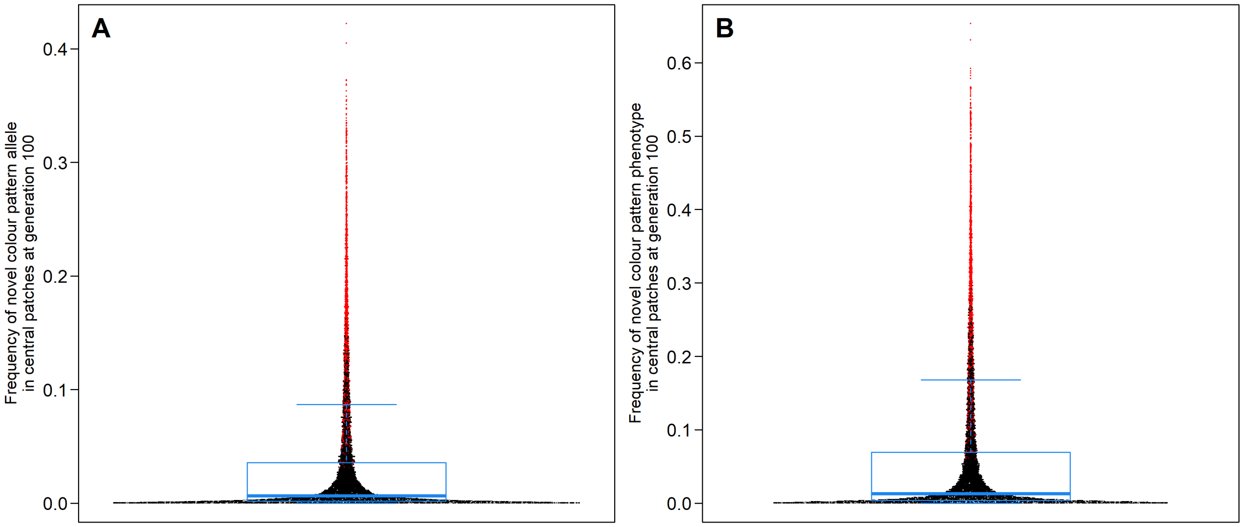


##### 3.3.2 Specifically testing for genetic architecture of attraction traits

Double check if indeed all simulations that reached generation 2500 also had an allele-freq>=0.95:

```
print(lengths(freq_exceed_095))
```

```
## [1] 940 956 969
```

```
print(c(nrow(unique(l1_2500[, c("Scenario",
    "Simulation")])), nrow(unique(l5_2500[,
    c("Scenario", "Simulation")])),
    nrow(unique(l10_2500[, c("Scenario",
        "Simulation")]))))
```

```
## [1] 940 956 969
```

To perform a test, we need to have a number as response variable, which describes the allele frequency dynamics. We decided to extract the generation value at which 0.95 was reached:

```
generations <- sapply(1:3, function(x) get(paste0("l",
    loci[x]))$Generation[freq_exceed_095[[x]]])
```

Get the respective simulation scenario (out of the list of the 120 scenarios)

```
scenarios <- unlist(sapply(1:3, function(x) get(paste0("l",
    loci[x]))$Scenario[freq_exceed_095[[x]]]))
```

Combine this in a table, adding a column on number of loci, and sex of the first mutant

```
glm_data_gen <- data.frame(param[scenarios,
    1:6], number_loci = unlist(sapply(1:3,
    function(x) rep(loci[x], lengths(generations)[x]))),
    sex_first_mutant = unlist(sapply(1:3,
        function(x) get(paste0("l",
            loci[x]))$sex_mutant[freq_exceed_095[[x]]])),
    generation = unlist(generations),
    stringsAsFactors = F)
```

Make the predictors (ordered) factors (like before)

```
glm_data_gen$bottleneck <- factor(ifelse(glm_data_gen$bottleneck ==
    1, "Y", "N"))
glm_data_gen$sex_first_mutant <- factor(ifelse(glm_data_gen$sex_first_mutant ==
    1, "M", "F"))
glm_data_gen$strength_attract <- as.ordered(glm_data_gen$strength_attract)
glm_data_gen$pred_constant <- as.ordered(glm_data_gen$pred_constant)
glm_data_gen$number_loci <- as.ordered(glm_data_gen$number_loci)
```

###### 3.3.2.1 Testing for the effect of genetic architecture of attraction traits alone (+ plot)

Analyse the effect of number of loci alone on generation

```
generation_glm_only_loci <- glm(generation ~
    number_loci, family = "poisson",
    data = glm_data_gen)
```

Get EMMs and contrasts

```
emms_gen_loci <- summary(emmeans(generation_glm_only_loci,
    pairwise ~ number_loci, transform = "response",
    adjust = "none"))
emms_gen_loci$contrasts$p.value <- p.adjust(emms_gen_loci$contrasts$p.value,
    method = "bonferroni")
```

```
print(emms_gen_loci)
```

```
## $emmeans
##  number_loci   rate     SE  df asymp.LCL asymp.UCL
##  1           1436.6 1.2363 Inf    1434.2    1439.1
##  5            826.0 0.9295 Inf     824.2     827.8
##  10           853.1 0.9383 Inf     851.3     855.0
##
## Confidence level used: 0.95
##
## $contrasts
##  contrast estimate    SE  df z.ratio p.value
##  1 - 5      610.66 1.547 Inf 394.812 <.0001
##  1 - 10     583.51 1.552 Inf 375.969 <.0001
##  5 - 10     -27.15 1.321 Inf -20.558 <.0001
```

We later plot this effect of number of loci on generations until 0.95, together with allele frequency curves (as done before) for one scenario.

First, we calculate allele frequencies for a specific scenario (sexual conflict present, predators absent from central patches for 100 generations, no bottleneck, probability of encounter = 0.01, strength of attraction = 3, predator learning threshold = 2, sex of first mutant = female), across different number of loci coding for attraction phenotypes (1, 5 or 10). This is the same settings we used to make the Gif.

```
scenario <- (1:nrow(param))[param$sex_conflict ==
    1 & param$bottleneck == 0 & param$prob_encounter ==
    0.01 & param$predator_absence ==
    100 & param$strength_attract ==
    3 & param$pred_constant == 2]
sex_mutant <- 0

# Empty lists:
all_freq_patt_all <- list()
all_freq_attr_nov_all <- list()
all_freq_attr_anc_all <- list()

# Iterate through number of loci
for (lo in c(1, 5, 10)) {
    # Extract results for this scenario
    scenario_simulation <- get(paste0("l",
        lo, "_2500"))[get(paste0("l",
        lo, "_2500"))$Scenario == scenario &
        get(paste0("l", lo, "_2500"))$sex_mutant ==
            0, c("Scenario", "Simulation")]

    # Get column names for preference
    # loci
    novel_attr_c_hom <- paste0("Rc",
        1:lo, "_hom")
    novel_attr_nc_hom <- paste0("Rnc",
        1:lo, "_hom")
    novel_attr_c_het <- paste0("Rc",
        1:lo, "_het")
    novel_attr_nc_het <- paste0("Rnc",
        1:lo, "_het")
```

```
ances_attr_c_hom <- paste0("Wc",
    1:lo, "_hom")
ances_attr_nc_hom <- paste0("Wnc",
    1:lo, "_hom")
ances_attr_c_het <- paste0("Wc",
    1:lo, "_het")
ances_attr_nc_het <- paste0("Wnc",
    1:lo, "_het")

# Run through single simulations,
# calculate allele frequency for the
# dominant allele at the colour
# locus and average allele frequency
# for the dominant alleles at at the
# attraction loci (first for novel,
# then for ancestral pattern)
all_freq_patt <- matrix(ncol = nrow(scenario_simulation),
    nrow = 2500)
all_freq_attr_nov <- matrix(ncol = nrow(scenario_simulation),
    nrow = 2500)
all_freq_attr_anc <- matrix(ncol = nrow(scenario_simulation),
    nrow = 2500)
for (setto in 1:nrow(scenario_simulation)) {
    matchers <- (1:nrow(get(paste0("l",
        lo))))[get(paste0("l", lo))$Scenario ==
        scenario_simulation$Scenario[setto] &
        get(paste0("l", lo))$Simulation ==
            scenario_simulation$Simulation[setto]]
    all_freq_patt[, setto] <- (2 *
        get(paste0("l", lo))$Cc_hom[matchers] +
        2 * get(paste0("l", lo))$Cnc_hom[matchers] +
        get(paste0("l", lo))$Cc_het[matchers] +
        get(paste0("l", lo))$Cnc_het[matchers])/(2 *
        get(paste0("l", lo))$N_non_central[matchers] +
        2 * get(paste0("l", lo))$N_central[matchers])
    all_freq_attr_nov[, setto] <- (2 *
        rowSums(get(paste0("l",
            lo))[matchers, novel_attr_c_hom,
            drop = F]) + 2 * rowSums(get(paste0("l",
        lo))[matchers, novel_attr_nc_hom,
        drop = F]) + rowSums(get(paste0("l",
        lo))[matchers, novel_attr_c_het,
        drop = F]) + rowSums(get(paste0("l",
        lo))[matchers, novel_attr_nc_het,
        drop = F]))/(2 * lo * get(paste0("l",
        lo))$N_non_central[matchers] +
        2 * lo * get(paste0("l",
            lo))$N_central[matchers])
    all_freq_attr_anc[, setto] <- (2 *
        rowSums(get(paste0("l",
            lo))[matchers, ances_attr_c_hom,
            drop = F]) + 2 * rowSums(get(paste0("l",
        lo))[matchers, ances_attr_nc_hom,
```

```
            drop = F]) + rowSums(get(paste0("l",
            lo))[matchers, ances_attr_c_het,
            drop = F]) + rowSums(get(paste0("l",
            lo))[matchers, ances_attr_nc_het,
            drop = F]))/(2 * lo * get(paste0("l",
            lo))$N_non_central[matchers] +
            2 * lo * get(paste0("l",
                lo))$N_central[matchers])
    }
    all_freq_patt_all[[length(all_freq_patt_all) +
        1]] <- all_freq_patt
    all_freq_attr_nov_all[[length(all_freq_attr_nov_all) +
        1]] <- all_freq_attr_nov
    all_freq_attr_anc_all[[length(all_freq_attr_anc_all) +
        1]] <- all_freq_attr_anc
}
```

Now, we plot this:

```
png("Figures/simulations_allele_freq_generations_number_loci.png",
    width = 3300, height = 2000, res = 300)
# Open plot layout
layout(matrix(c(1:3, rep(4, 3)), ncol = 2),
    widths = c(1.4, 1))
par(oma = c(1.8, 4, 0.05, 1))

# This defines the first generation
# to be displayed
start_at_gen <- 25
# This is the actual beginning of
# the x-axis. However, we'll
# 'pretend' as if the x-axis starts
# at 1
min_xaxs <- 20

# We add one plot for each set of
# loci
for (lo in 1:3) {

    par(mar = c(1.5, 0.3, 1.2, 0.3))
    # We logarithmize the x-axis and
    # make it start at 20
    plot(1, 1, xlim = c(min_xaxs, 2500),
        ylim = 0:1, type = "n", xaxt = "n",
        yaxt = "n", xlab = "", ylab = "",
        xaxs = "i", yaxs = "i", log = "x")
    # Add a vertical line where
    # predators return
    abline(v = 100, lty = "dashed",
        lwd = 1)
    # If we're in the second plot, add
    # legend
    if (loci[lo] == 5) {
        legend("left", pch = 15, pt.cex = 1.7,
```

```r
        col = c("gray30", rgb(255/255,
            149/255, 8/255), rgb(28/255,
            57/255, 252/255)), c("Wing pattern",
            "Increased attraction to novel",
            "Decreased attraction to ancestral"),
        cex = 1.05)
    mtext("(Mean) allele frequency",
        2, line = 3)
}
# Add info on number of loci above
# each plot
par(xpd = NA)
text(10^(mean(par("usr")[1:2])),
    1.05, paste(loci[lo], ifelse(loci[lo] ==
        1, " Locus", " Loci")),
    cex = 1.3)
par(xpd = F)

# Add frequencies as partly
# transparent lines to graph Note
# that we plot 1 minus the frequency
# of alleles that code for increased
# attraction to the ancestral
# pattern
invisible(sapply(1:nrow(scenario_simulation),
    function(x) add_lines(all_mat = list(all_freq_patt_all[[lo]][start_at_gen:2500,
        ], all_freq_attr_nov_all[[lo]][start_at_gen:2500,
        ], 1 - all_freq_attr_anc_all[[lo]][start_at_gen:2500,
        ]), x = x, n_mat = 3, cols = c("gray30",
        rgb(255/255, 149/255, 8/255),
        rgb(28/255, 57/255, 252/255)))))

# Add y-axis
axis(2, at = seq(0, 1, 0.25), labels = F,
    tck = -0.015)
axis(2, at = seq(0, 1, 0.25), lwd = 0,
    line = -0.6, las = 2)
# Add x-axis Note that we plot the
# number '1' at the position of
# min_xaxs
axis(1, at = c(min_xaxs, start_at_gen,
    100, 500, 2500), labels = F,
    tck = -0.015)
axis(1, at = c(min_xaxs, start_at_gen,
    100, 500, 2500), c(1, start_at_gen,
    100, 500, 2500), lwd = 0, line = -0.8)
# Now, 'cut out' a part of the
# x-axis between 20 and 25
two_pts <- log(c(min_xaxs, start_at_gen))
cut_pts_left <- exp(mean(two_pts) -
    c(0.18, 0.08) * diff(two_pts))
cut_pts_right <- exp(mean(two_pts) +
    c(0.08, 0.18) * diff(two_pts))
```

```
    par(xpd = NA)
    polygon(c(cut_pts_left[1], cut_pts_right[1],
        cut_pts_right[2], cut_pts_left[2]),
        rep(c(-0.035, 0.035), each = 2),
        col = "white", border = NA)
    segments(c(cut_pts_left[1], cut_pts_right[1]),
        rep(-0.035, 2), c(cut_pts_left[2],
            cut_pts_right[2]), rep(0.035,
            2))
    par(xpd = F)
    # Add x-axis if we're in the last
    # plot
    if (loci[lo] == 10) {
        mtext("Generation", 1, line = 1.8)
    }
}


# Plot effect of number of loci on
# fixation dynamics
par(mar = c(1.5, 6.6, 1.2, 0.45))
plot(1, 1, type = "n", xaxs = "i", xlab = "",
    ylab = "", xaxt = "n", yaxt = "n",
    xlim = c(0.5, 3.5), ylim = range(glm_data_gen$generation))
# Add data as beeswarms
beeswarm(generation ~ number_loci, at = 1:3,
    add = T, cex = 0.45, pch = 20, data = glm_data_gen,
    col = "gray30")
# Add EMMs and their CIs (CIs won't
# be visible, as they're so small)
points(1:3, emms_gen_loci$emmeans$rate,
    col = "red", pch = 20, cex = 2.2)
segments(1:3, emms_gen_loci$emmeans$asymp.LCL,
    1:3, emms_gen_loci$emmeans$asymp.UCL,
    col = "red", lwd = 2, lend = 3)
# Axes, labels, etc.
axis(2, at = seq(700, 2500, 300), lwd = 0,
    line = -0.5, las = 2)
axis(2, at = seq(700, 2500, 300), labels = F,
    tck = -0.015)
axis(1, at = 1:3, paste0(c(1, 5, 10),
    c(" Locus", " Loci", " Loci")),
    lwd = 0, line = -0.1, cex.axis = 1.3)
axis(1, at = 1:3, paste0("(", unname(table(glm_data_gen$number_loci)),
    ")"), lwd = 0, line = 1.1)
mtext("Generations until allele frequency of 0.95",
    2, line = 3.4)
# Add panel ID
text(par("usr")[1] + 0.08 * diff(par("usr")[1:2]),
    par("usr")[3] + 0.97 * diff(par("usr")[3:4]),
    "B", font = 2, cex = 1.5, pos = 2,
    offset = 0)
# We add the 'A' for the frequency
```

```
# plots following the dimensions of
# this current plot.
par(xpd = NA)
text(par("usr")[1] - 1.96 * diff(par("usr")[1:2]),
    par("usr")[3] + 0.97 * diff(par("usr")[3:4]),
    "A", font = 2, cex = 1.5, pos = 2,
    offset = 0)
par(xpd = F)

invisible(dev.off())
```

Show png we just made:

```
fig_disp <- readPNG("Figures/simulations_allele_freq_generations_number_loci.png")
grid.raster(fig_disp)
```

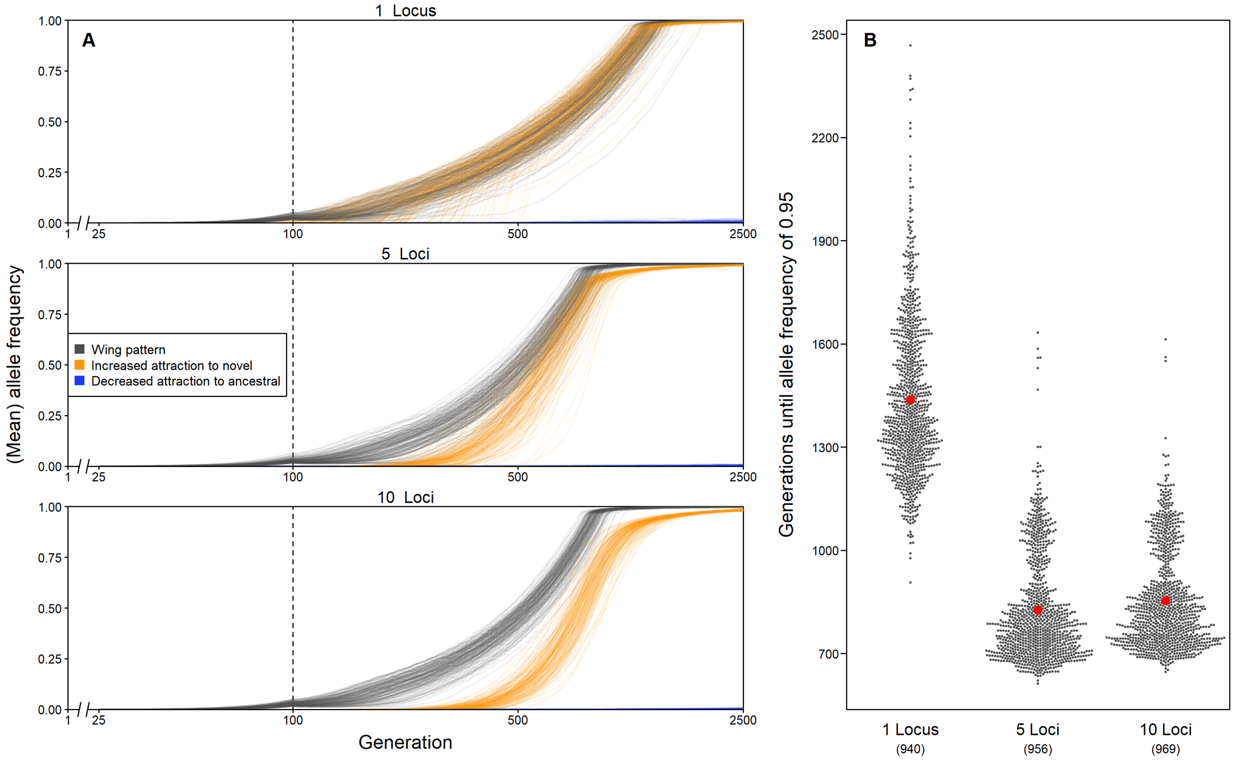


###### 3.3.2.2 Interactions between genetic architecture of attraction traits and other parameters

Check if this strong effect of number of loci might be biased by the fact that the occurence of fixation might correlate with certain scenarios, which then are actually driving this effect.

- A) Check if certain scenarios are more commonly represented among simulations that went to fixation in certain loci sets rather than others.

Get unique scenarios

```
uni_scen <- unique(glm_data_gen[, c(3,
    5, 8, 2, 6)])
# Order them and remove irritating
```

```r
# row names
uni_scen <- uni_scen[order(uni_scen$prob_encounter,
    uni_scen$strength_attract, uni_scen$sex_first_mutant,
    uni_scen$bottleneck, uni_scen$pred_constant),
    ]
rownames(uni_scen) <- NULL
```

Use the opportunity to give a scenario ID to each scenario (excluding number of loci) in the GLM table (used for later).

```r
glm_data_gen$ID <- as.character(sapply(1:nrow(glm_data_gen),
    function(x) (1:nrow(uni_scen))[uni_scen$prob_encounter ==
        glm_data_gen$prob_encounter[x] &
        uni_scen$strength_attract ==
            glm_data_gen$strength_attract[x] &
        uni_scen$sex_first_mutant ==
            glm_data_gen$sex_first_mutant[x] &
        uni_scen$bottleneck == glm_data_gen$bottleneck[x] &
        uni_scen$pred_constant == glm_data_gen$pred_constant[x]]))
# Add leading 0s
glm_data_gen$ID <- ifelse(nchar(glm_data_gen$ID) ==
    1, paste0("0", glm_data_gen$ID),
    glm_data_gen$ID)
```

Summarize how many times each scenario is found in the table for the three sets of loci

```r
uni_scen$perc_1_loci <- sapply(1:nrow(uni_scen),
    function(x) sum(glm_data_gen$number_loci ==
        "1" & glm_data_gen$prob_encounter ==
        uni_scen$prob_encounter[x] &
        glm_data_gen$strength_attract ==
            uni_scen$strength_attract[x] &
        glm_data_gen$sex_first_mutant ==
            uni_scen$sex_first_mutant[x] &
        glm_data_gen$bottleneck == uni_scen$bottleneck[x] &
        glm_data_gen$pred_constant ==
            uni_scen$pred_constant[x]))
uni_scen$perc_5_loci <- sapply(1:nrow(uni_scen),
    function(x) sum(glm_data_gen$number_loci ==
        "5" & glm_data_gen$prob_encounter ==
        uni_scen$prob_encounter[x] &
        glm_data_gen$strength_attract ==
            uni_scen$strength_attract[x] &
        glm_data_gen$sex_first_mutant ==
            uni_scen$sex_first_mutant[x] &
        glm_data_gen$bottleneck == uni_scen$bottleneck[x] &
        glm_data_gen$pred_constant ==
            uni_scen$pred_constant[x]))
uni_scen$perc_10_loci <- sapply(1:nrow(uni_scen),
    function(x) sum(glm_data_gen$number_loci ==
        "10" & glm_data_gen$prob_encounter ==
        uni_scen$prob_encounter[x] &
        glm_data_gen$strength_attract ==
            uni_scen$strength_attract[x] &
        glm_data_gen$sex_first_mutant ==
```

```
            uni_scen$sex_first_mutant[x] &
        glm_data_gen$bottleneck == uni_scen$bottleneck[x] &
        glm_data_gen$pred_constant ==
            uni_scen$pred_constant[x]))
```

Calculate binomial confidence intervals and round

```
confint_1 <- round(100 * binom.confint(uni_scen$perc_1_loci,
    sum(uni_scen$perc_1_loci), method = "exact")[,
    4:6], 1)
confint_5 <- round(100 * binom.confint(uni_scen$perc_5_loci,
    sum(uni_scen$perc_5_loci), method = "exact")[,
    4:6], 1)
confint_10 <- round(100 * binom.confint(uni_scen$perc_10_loci,
    sum(uni_scen$perc_10_loci), method = "exact")[,
    4:6], 1)
```

Add as columns to table and shorten column names

```
uni_scen$perc_1_loci <- paste0(confint_1[,
    1], " [", confint_1[, 2], "-", confint_1[,
    3], "]%")
uni_scen$perc_5_loci <- paste0(confint_5[,
    1], " [", confint_5[, 2], "-", confint_5[,
    3], "]%")
uni_scen$perc_10_loci <- paste0(confint_10[,
    1], " [", confint_10[, 2], "-",
    confint_10[, 3], "]%")
# Make column names shorter:
# e=prob-encounter,
# a=strength_attraction,
# sex=sex_first_mutant,
# bottle=bottleneck, Q=pred_constant
names(uni_scen)[1:5] <- c("e", "a",
    "sex", "bottle", "Q")
print(uni_scen)
```

```
##        e a sex bottle Q       perc_1_loci       perc_5_loci      perc_10_loci
## 1  0.005 3   M      Y 1      0.1 [0-0.6]%      0.1 [0-0.6]%        0 [0-0.4]%
## 2  0.010 1   F      N 1    2.1 [1.3-3.3]%    1.3 [0.7-2.2]%    2.4 [1.5-3.5]%
## 3  0.010 1   F      Y 1    2.6 [1.6-3.8]%    2.4 [1.5-3.6]%        2 [1.2-3]%
## 4  0.010 1   F      Y 2      1 [0.4-1.8]%    1.3 [0.7-2.2]%    0.4 [0.1-1.1]%
## 5  0.010 1   F      Y 4        0 [0-0.4]%      0.2 [0-0.8]%      0.2 [0-0.7]%
## 6  0.010 1   M      N 1      1 [0.4-1.8]%    0.4 [0.1-1.1]%    0.5 [0.2-1.2]%
## 7  0.010 1   M      Y 1    3.1 [2.1-4.4]%    2.8 [1.9-4.1]%      3.6 [2.5-5]%
## 8  0.010 1   M      Y 2    1.3 [0.7-2.2]%    0.4 [0.1-1.1]%    0.7 [0.3-1.5]%
## 9  0.010 1   M      Y 4      0.2 [0-0.8]%    0.4 [0.1-1.1]%      0.1 [0-0.6]%
## 10 0.010 3   F      N 1 24.9 [22.2-27.8]% 24.5 [21.8-27.3]% 26.3 [23.6-29.2]%
## 11 0.010 3   F      N 2   17.3 [15-19.9]% 19.9 [17.4-22.5]% 16.4 [14.1-18.9]%
## 12 0.010 3   F      N 4      6.5 [5-8.3]%    5.4 [4.1-7.1]%      4.5 [3.3-6]%
## 13 0.010 3   F      Y 1      4 [2.9-5.5]%    3.7 [2.6-5.1]%    4.3 [3.1-5.8]%
## 14 0.010 3   F      Y 2    3.8 [2.7-5.3]%      4 [2.8-5.4]%    3.8 [2.7-5.2]%
## 15 0.010 3   F      Y 4    2.6 [1.6-3.8]%    2.4 [1.5-3.6]%      2.8 [1.8-4]%
## 16 0.010 3   M      N 1   10.6 [8.7-12.8]%  10.5 [8.6-12.6]%   11.1 [9.2-13.3]%
## 17 0.010 3   M      N 2    5.2 [3.9-6.8]%    6.2 [4.7-7.9]%      6.3 [4.8-8]%
```

```
## 18 0.010 3   M      N 4      1.9 [1.1-3]%    2.3 [1.4-3.5]%    1.4 [0.8-2.4]%
## 19 0.010 3   M      Y 1    4.9 [3.6-6.5]%      4.5 [3.3-6]%    4.6 [3.4-6.2]%
## 20 0.010 3   M      Y 2    3.7 [2.6-5.1]%    5.2 [3.9-6.8]%    5.1 [3.8-6.6]%
## 21 0.010 3   M      Y 4    3.2 [2.2-4.5]%    2.2 [1.4-3.3]%    3.3 [2.3-4.6]%
```

Seems not to be the case!

- B) Use a GLM to check for the effect of number of loci across simulation scenarios.

Run GLM with interactions and plot it using `emmeans`.

```
generation_glm_only_loci_scenario <- glm(generation ~
    number_loci * ID, family = "poisson",
    data = glm_data_gen)
png("Figures/simulations_generations_effect_loci_scenarios.png",
    width = 1500, height = 3500, res = 300)
plot(emmeans(generation_glm_only_loci_scenario,
    ~number_loci | ID, transform = "response"),
    xlab = "Generations until allele frequency of 0.95 was reached",
    ylab = "Number of Loci")
```

```
## Warning: Removed 2 rows containing missing values (geom_segment).
```

```
## Warning: Removed 2 rows containing missing values (geom_point).
```

```
invisible(dev.off())
```

Show png we just made:

```
fig_disp <- readPNG("Figures/simulations_generations_effect_loci_scenarios.png")
grid.raster(fig_disp)
```

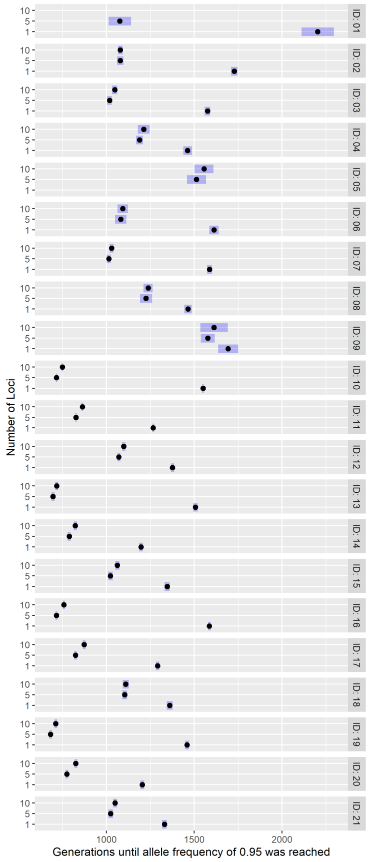


Also, this seems not to be the case!

- C) Let's anyway also check for interactions between other parameters and number of loci:

Include number_loci as well as all possible interactions it might be involved in. Don't include prob_encounter, as there is only two levels, one of which only has two datapoints

```
generation_glm <- glm(generation ~ number_loci +
    bottleneck + strength_attract +
    pred_constant + sex_first_mutant +
    number_loci:bottleneck + number_loci:strength_attract +
    number_loci:pred_constant + number_loci:sex_first_mutant,
    family = "poisson", data = glm_data_gen)
```

Reduce this model with stepwise AIC (this time backward)

```
step_glm_gen <- suppressWarnings(stepAIC(generation_glm,
    scope = list(lower = ~1), direction = "backward",
    trace = FALSE))
```

Get p-values and correct for multiple testing

```
p_values_gen <- joint_tests(step_glm_gen)
p_values_gen$p.value <- p.adjust(p_values_gen$p.value,
    method = "bonferroni")
print(p_values_gen)
```

```
##  model term                   df1 df2   F.ratio p.value
##  number_loci                    2 Inf 13015.927 <.0001
```

```
##   bottleneck                       1 Inf  1159.512 <.0001
##   strength_attract                 1 Inf 21230.363 <.0001
##   pred_constant                    2 Inf  8425.767 <.0001
##   sex_first_mutant                 1 Inf    13.526 0.0019
##   number_loci:strength_attract     2 Inf  3310.706 <.0001
##   number_loci:pred_constant        4 Inf  7260.693 <.0001
##   number_loci:sex_first_mutant     2 Inf     3.802 0.1786
```

Get dominance

```
dominance_fact_gen <- suppressWarnings(averageContribution(dominanceAnalysis(step_glm_gen),
    fit.functions = "r2.m")$r2.m)
print_r2_gen <- data.frame(fixed_effect = names(dominance_fact_gen),
    R2 = round(unname(dominance_fact_gen),
        3))
print(print_r2_gen[order(print_r2_gen$R2,
    decreasing = T), ])
```

```
##                    fixed_effect    R2
## 7    number_loci:pred_constant 0.267
## 6 number_loci:strength_attract 0.203
## 1                  number_loci 0.165
## 8 number_loci:sex_first_mutant 0.165
## 3             strength_attract 0.025
## 4                pred_constant 0.016
## 2                   bottleneck 0.002
## 5             sex_first_mutant 0.000
```

Three interactions seem to have a large effect. Let's visualize those:

```
plot(emmeans(step_glm_gen, ~number_loci |
    pred_constant, transform = "response"),
    xlab = "Generations until allele frequency of 0.95 was reached")
```

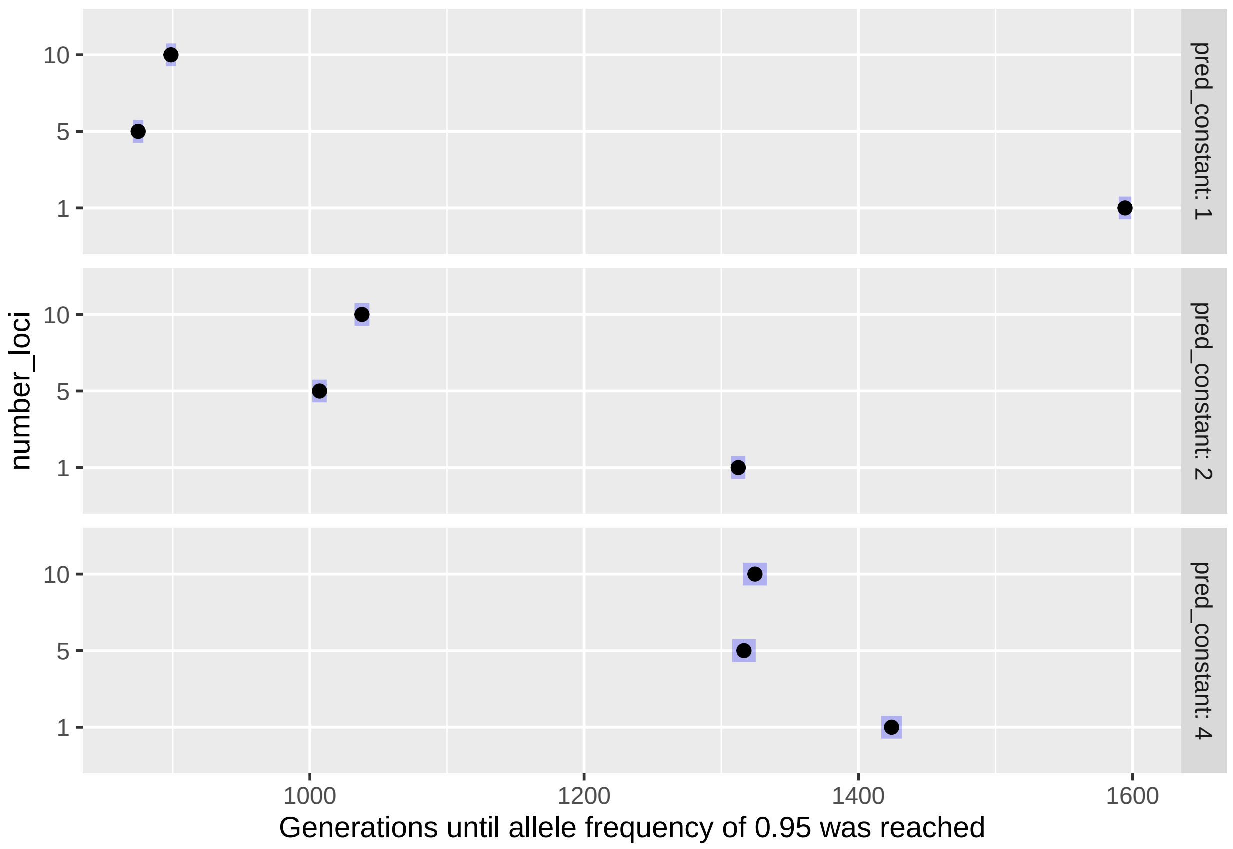


```
plot(emmeans(step_glm_gen, ~number_loci |
    strength_attract, transform = "response"),
    xlab = "Generations until allele frequency of 0.95 was reached")
```

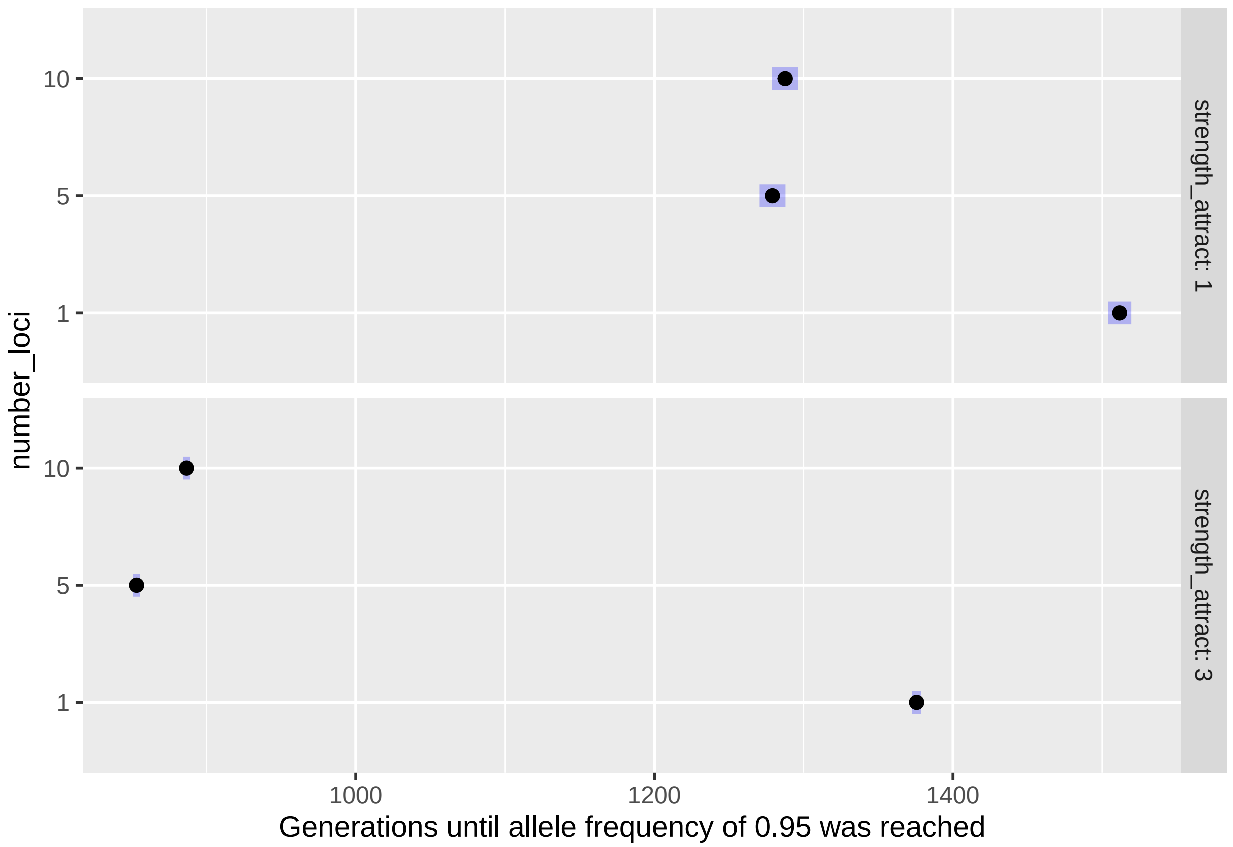


```
plot(emmeans(step_glm_gen, ~number_loci |
    sex_first_mutant, transform = "response"),
    xlab = "Generations until allele frequency of 0.95 was reached")
```

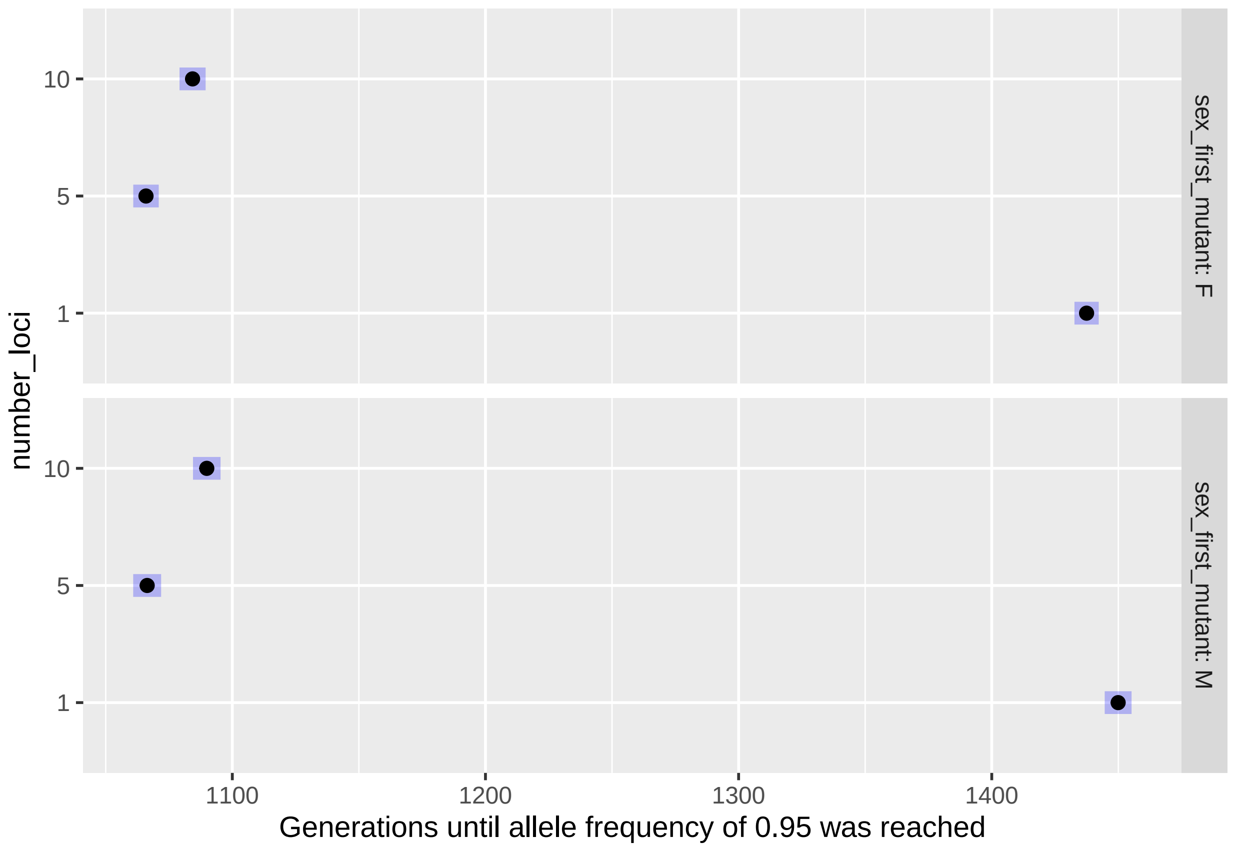


The interaction between number_loci and pred_constant is particularly intriguing! While the trend 1>5<10 is visible across predation rates, it seems to be more pronounced the smaller the predation rate is!

Let's visualize this more prettily.

Save EMMs:

```
EMM_loci_pred <- summary(emmeans(step_glm_gen,
    ~number_loci | pred_constant, transform = "response"))
```

Plot it (similar as before)

```
png("Figures/simulations_generations_effect_loci_pred.png",
    width = 2700, height = 1500, res = 300)
par(mar = c(3, 3.5, 1.3, 0.5))

# Get the lowest value of the lower
# CIs and the highest value of the
# upper CIs
min_y <- min(EMM_loci_pred$asymp.LCL)
max_y <- max(EMM_loci_pred$asymp.UCL)

# Open empty plot
plot(1, 1, type = "n", xlab = "", ylab = "",
    xaxt = "n", yaxt = "n", xlim = c(min_y,
        max_y), ylim = c(0.5, nrow(EMM_loci_pred) +
        0.5), yaxs = "i")
# Provide vertical lines at all
```

```r
# x-values
abline(v = seq(600, 1600, 200), lty = "dashed",
    lwd = 0.5)
# Add CIs
segments(EMM_loci_pred$asymp.LCL, 1:nrow(EMM_loci_pred),
    EMM_loci_pred$asymp.UCL, 1:nrow(EMM_loci_pred),
    lend = 3, lwd = 12, col = "lightblue")
# Add EMMs
points(EMM_loci_pred$rate, 1:nrow(EMM_loci_pred),
    pch = 20, cex = 1.3)

# Add the R2 and the p-value
mtext(bquote(italic(N) ~ ":" ~ italic(Q) ~
    "  (" * R^2 ~ "=" ~ .(format(round(print_r2_gen$R2[print_r2_gen$fixed_effect ==
    "number_loci:pred_constant"], 3),
    nsmall = 3)) * ", p < 0.001" * ")"),
    3, line = 0.2, cex = 0.75)
# Add axes
axis(1, labels = F, tck = -0.02)
axis(1, lwd = 0, line = -0.7, cex.axis = 0.9)
axis(2, at = 1:nrow(EMM_loci_pred),
    labels = F, tck = -0.01)
axis(2, at = 1:nrow(EMM_loci_pred),
    paste0(levels(EMM_loci_pred[, 1])[EMM_loci_pred[,
        1]], " : ", levels(EMM_loci_pred[,
        2])[EMM_loci_pred[, 2]]), lwd = 0,
    las = 2, line = -0.6, cex.axis = 0.9)
uni1 <- length(unique(EMM_loci_pred[,
    1]))
uni2 <- length(unique(EMM_loci_pred[,
    2]))
abline(h = seq(uni1, (uni1 * uni2) -
    uni1, uni1) + 0.5, lty = "dashed",
    lwd = 0.5)

mtext("Generations until allele frequency of 0.95 was reached",
    1, line = 2, cex = 0.9)
mtext("Number of loci : Predation constant",
    2, line = 2.7, cex = 0.9)

invisible(dev.off())
```

Show png we just made:

```r
fig_disp <- readPNG("Figures/simulations_generations_effect_loci_pred.png")
grid.raster(fig_disp)
```

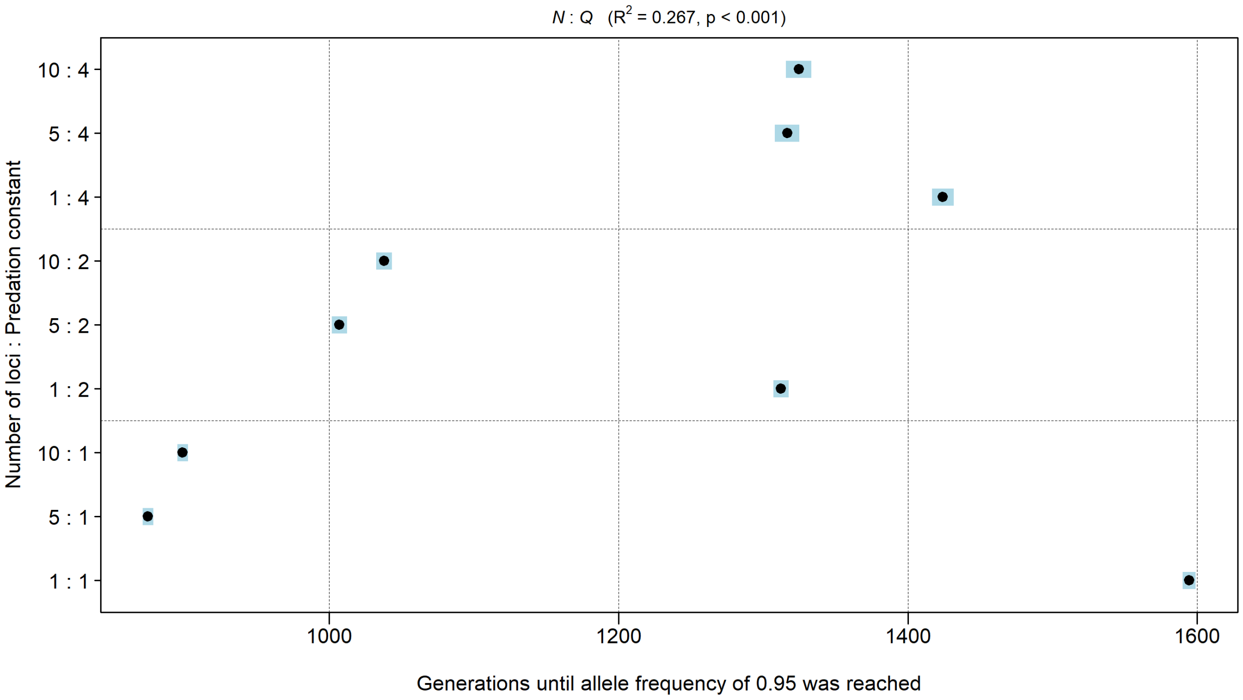


### 4 Empirical results

#### 4.1 Data read-in and manipulation

Read in JND data:

```
jnd <- read.csv("Data/Empirical/JND.csv",
    header = T, stringsAsFactors = F)
```

The columns indicate:

- **ID** = the IDs of the six erato individuals tested
- **JND_wing_black_vs_black_marker_on_wing_red** = JND between regular black from wing and blacked out red patch
- **JND_wing_red_vs_transparent_marker_on_wing_red** = JND between regular red patch and red patch painted with transparent marker

Read in data from experiment 1 (with Heliconius erato):

```
erato_data <- read.csv("Data/Empirical/erato_conflict_data.csv",
    header = T, stringsAsFactors = F)
```

The columns indicate:

- **female** = Female ID
- **treatment** = Wing pattern treatment. Four categories are available: `completely_untreated` = female was not treated or handled; `only_handled` = female was only handled, but wing pattern was untreated; `colourless_marker` = female red pattern area was treated with a transparent Copic marker;

`blacked_out` = female red pattern area was treated with a black Copic marker

- **date_enter_yyyy_mm_dd** = date female entered the experiment (beginning of the 6-day period where female is either with or without male in a cage, laying eggs)
- **habituation_day1_eggs** = number of eggs on the first day (acclimatization)
- **habituation_day2_eggs** = number of eggs on the second day (acclimatization)
- **Day_3_eggs_A** = number of eggs on the third day (experimental phase A)
- **Day_4_eggs_A** = number of eggs on the fourth day (experimental phase A)
- **Day_5_eggs_B** = number of eggs on the fifth day (experimental phase B)
- **Day_6_eggs_B** = number of eggs on the sixth day (experimental phase B)
- **Males_in_A_or_B** = Whether males were present in cage in phase A or B
- **Male_X** = three columns, indicating IDs of males present in cage
- **Eggs_with_males** = number of eggs laid with male present (e.g. if males present in phase A, then this is the sum of columns: `Day_3_eggs_A` and `Day_4_eggs_A`)
- **Eggs_without_males** = same logic, just that this is the sum of the days where males were absent
- **observation_time_1** = total number of seconds butterflies were observed on the first day that males were present
- **court_time_1** = total number of seconds with harassment behaviour (courting, chasing, mating attempts from males, etc.) on the first day that males were present
- **observation_time_2** = same as `observation_time_1`, but on the second day that males were present
- **court_time_2** = same as `court_time_1`, but on the second day that males were present
- **total_observation_time** = sum of `observation_time_1` and `observation_time_2`
- **total_court_time** = sum of `court_time_1` and `court_time_2`
- **first_court** = time in seconds until first courtship interaction occurs. Note that if no courtship interaction occurs, this is set to `NA`

Read in data from experiment 2 (with Heliconius timareta):

```
timareta_data <- read.csv("Data/Empirical/timareta_conflict_data.csv",
    header = T, stringsAsFactors = F)
```

Several column names are identical in this table as in the previous one, and these will not be described again. The columns that are 'new' indicate:

- **brood** = Hybrid brood ID (check also tables in doi.org/10.5061/dryad.pc866t1ng containing male individuals from same broods for further details)
- **wing_colour** = Wing pattern of backcross females. Two categories are available: `yellow_red` = female displays a yellow and a red band on the wing, similar to *H. heurippa*; `yellow` = female displas only a yellow band, similar to *H. timareta linaresi*
- **whole_wing_cm2** = Total area of right (mostly intact) forewing, in cm^2
- **yellow_cm2** = Total area of yellow band on right (mostly intact) forewing, in cm^2
- **red_cm2** = Total area of red band on right (mostly intact) forewing, in cm^2
- **habituation_day1_2_eggs** = number of eggs on the first and second day combined (acclimatization)
- **Day_3_4_eggs_A** = number of eggs on the third and fourth day combined (experimental phase A)
- **Day_5_6_eggs_B** = number of eggs on the fifth and sixth day combined (experimental phase B)
- **Male** = Since only one male was added in these experiments, there is only one column indicating male ID
- **Day_3_4_hatch_A** = Number of caterpillars that hatched from eggs laid in phase A (`NA` where not assessed)
- **Day_5_6_hatch_B** = Number of caterpillars that hatched from eggs laid in phase B (`NA` where not assessed)

#### 4.2 JND for manipulated *H. erato* wings

Here, we'll simply plot the JND data:

```
png("Figures/empirical_jnds.png", width = 5400,
    height = 3600, res = 600)
par(mar = c(1.5, 3.3, 0.4, 0.4))
plot(1, 1, type = "n", xaxt = "n", yaxt = "n",
    xlab = "", ylab = "", xlim = c(0.5,
        2.5), xaxs = "i", ylim = range(c(jnd$JND_wing_black_vs_black_marker_on_wing_red,
        jnd$JND_wing_red_vs_transparent_marker_on_wing_red)))
# Add beeswarms
beeswarm(jnd$JND_wing_black_vs_black_marker_on_wing_red,
    at = 1, add = T, pch = 21, cex = 2,
    spacing = 1.1)
beeswarm(jnd$JND_wing_red_vs_transparent_marker_on_wing_red,
    at = 2, add = T, pch = 21, cex = 2,
    spacing = 1.1)
# Add boxplots
boxplot(jnd$JND_wing_black_vs_black_marker_on_wing_red,
    outline = F, at = 1, add = T, col = adjustcolor("white",
        0), border = "dodgerblue2",
    yaxt = "n")
boxplot(jnd$JND_wing_red_vs_transparent_marker_on_wing_red,
    outline = F, at = 2, add = T, col = adjustcolor("white",
        0), border = "dodgerblue2",
    yaxt = "n")
# Axes and titles
axis(1, at = 1:2, labels = F, tck = -0.01)
axis(1, at = 1:2, c("'blacked-out red pattern' vs 'black pattern'",
    "'transparently-painted red pattern' vs 'red pattern'"),
    lwd = 0, line = -0.5)
axis(2, labels = F, tck = -0.01)
axis(2, las = 2, lwd = 0, line = -0.3)
mtext("Just noticeable difference (JND)",
    2, line = 2.4)

invisible(dev.off())
```

Show png we just made:

```
fig_disp <- readPNG("Figures/empirical_jnds.png")
grid.raster(fig_disp)
```

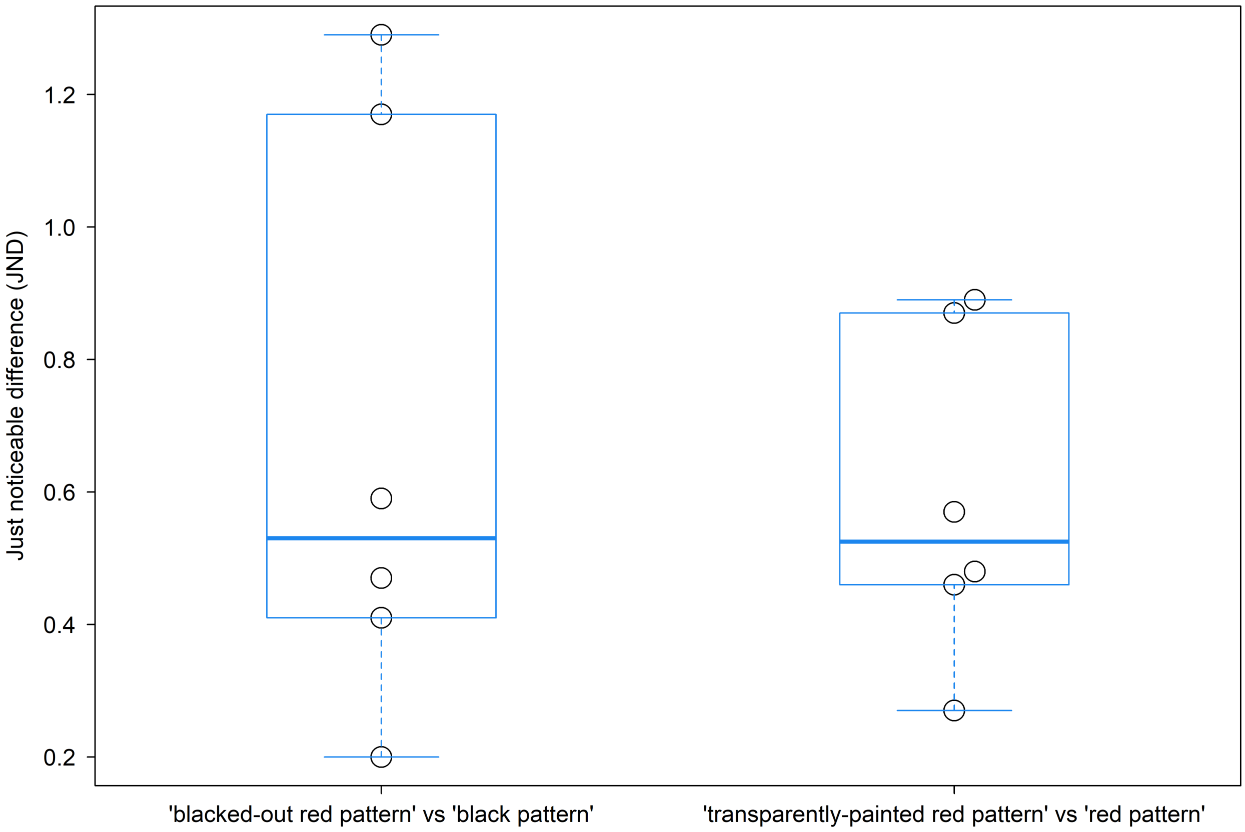


#### 4.3 Sizes of colour bands on backcross to *H. timareta* females

For a subset of females from experiment 2, we have measurements of whole wing size and sizes of the yellow and red band available. We plot the band sizes, once for absolute values and once for values corrected by total wing area.

First, we add two new columns, for relative size of the bands:

```
timareta_data$yellow_rel <- timareta_data$yellow_cm2/timareta_data$whole_wing_cm2
timareta_data$red_rel <- timareta_data$red_cm2/timareta_data$whole_wing_cm2
```

Now the plot. Note: These images will **NOT** be provided in the online supplement. If you rerun this, comment out the code where these variables appear (which, in this case, leaves you with empty plots; you could replace the images with `points`, in order to see where the wings would be placed).

```
png(paste0("Figures/empirical_wing_variation.png"),
    width = 4000, height = 4000, res = 300)

layout(matrix(1:2, ncol = 1))
par(mar = (c(3, 3.65, 0.5, 0.2)))

# First, we plot absolute areas

# Calculate range of x- and y-values
x_range <- range(timareta_data$yellow_cm2,
    na.rm = T)
y_range <- range(timareta_data$red_cm2,
```

```
    na.rm = T)
# Give some extra space in
# y-direction
y_range <- c(y_range[1] - 0.04 * diff(y_range),
    y_range[2] + 0.04 * diff(y_range))
# Open empty plot
plot(1, 1, xlim = x_range, ylim = y_range,
    type = "n", xaxt = "n", yaxt = "n",
    xlab = "", ylab = "")
# Axes and titles
axis(1, lwd = 0, line = -0.6)
axis(1, labels = F, tck = -0.01)
axis(2, lwd = 0, line = -0.6, las = 2)
axis(2, labels = F, tck = -0.01)
mtext(bquote("Area yellow band [" *
    cm^2 * "]"), 1, line = 1.72, cex = 1.1)
mtext(bquote("Area red band [" * cm^2 *
    "]"), 2, line = 2.32, cex = 1.1)
# Add images, ordered by magnitude
# of x-values
for (i in (1:nrow(timareta_data))[order(timareta_data$yellow_cm2)]) {
    if (!is.na(timareta_data$whole_wing_cm2[i])) {
        # Read in image
        ph1 <- readPNG(paste0("wing_photos/",
            timareta_data$female[i],
            ".png"))
        # We calculate a multiplier for the
        # image width, which will make it
        # relative to the actual size of the
        # wing. We calibrate to 200
        width_mult <- dim(ph1)[2]/200
        # Add wing image
        addImg(ph1, x = timareta_data$yellow_cm2[i],
            y = timareta_data$red_cm2[i],
            width = width_mult * 0.055 *
                diff(par("usr")[1:2]))
    }
}
# Make sure we have an intact
# plotting box
box(lwd = 1)

# Second, we plot areas relative to
# the whole wing size

# This is largely the same as above,
# with the exception that size of
# yellow and of red band get divided
# by whole wing area here

x_range <- range(timareta_data$yellow_rel,
    na.rm = T)
y_range <- range(timareta_data$red_rel,
```

```
    na.rm = T)
y_range <- c(y_range[1] - 0.04 * diff(y_range),
    y_range[2] + 0.04 * diff(y_range))
plot(1, 1, xlim = x_range, ylim = y_range,
    type = "n", xaxt = "n", yaxt = "n",
    xlab = "", ylab = "")

axis(1, lwd = 0, line = -0.6)
axis(1, labels = F, tck = -0.01)
axis(2, lwd = 0, line = -0.6, las = 2)
axis(2, labels = F, tck = -0.01)
mtext("Proportion yellow band", 1, line = 1.9,
    cex = 1.1)
mtext("Proportion red band", 2, line = 2.5,
    cex = 1.1)

for (i in (1:nrow(timareta_data))[order(timareta_data$yellow_rel)]) {
    if (!is.na(timareta_data$whole_wing_cm2[i])) {
        ph1 <- readPNG(paste0("wing_photos/",
            timareta_data$female[i],
            ".png"))
        width_mult <- dim(ph1)[2]/200
        addImg(ph1, x = timareta_data$yellow_rel[i],
            y = timareta_data$red_rel[i],
            width = width_mult * 0.055 *
                diff(par("usr")[1:2]))
    }
}

box(lwd = 1)

invisible(dev.off())
```

Show png we just made:

```
fig_disp <- readPNG("Figures/empirical_wing_variation.png")
grid.raster(fig_disp)
```

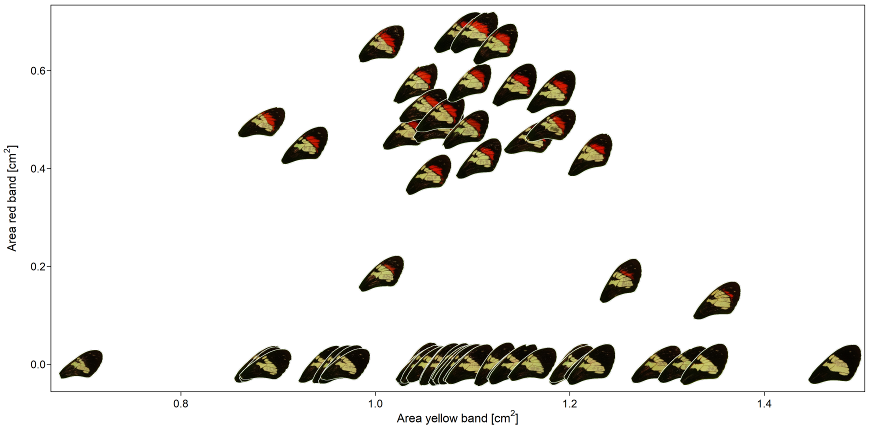


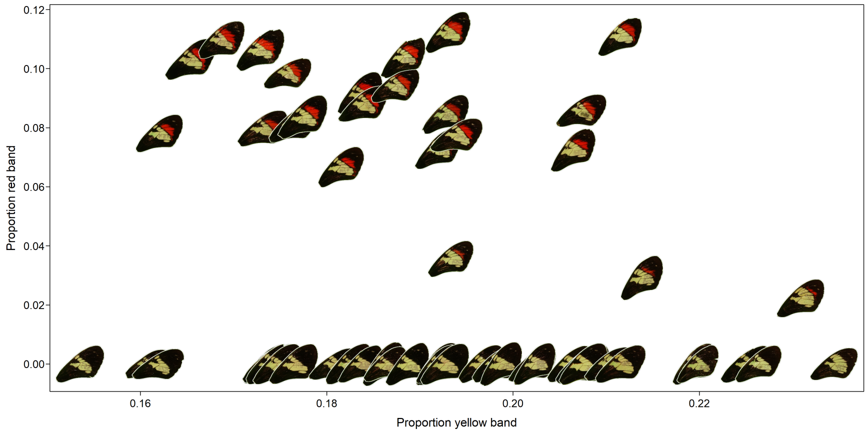


Size of the yellow band seems only weakly affected by presence/absence of the red band.

#### 4.4 Analyses of male harassment

##### 4.4.1 Experiment 1 (with *H. erato*)

Note that in this section, we repeat all the analyses for 2 different data sets: * A) Excluding females that didn't lay any eggs * B) Including females that didn't lay any eggs

Females that don't lay any eggs are likely unfit and it makes sense to exclude them. However, we want to show that excluding them does not qualitatively affect our results.

Add a column where all females that had a blacked out wing are labelled 'treatment' and all others (irrespective of treatment) are labelled 'control'

```
erato_data$contr_exp <- ifelse(erato_data$treatment ==
    "blacked_out", "treatment", "control")
```

Create a new dataset where we exclude those females that were completely untreated (these will appear again later in the egg-laying analyses)

```
erato_data1 <- erato_data[erato_data$treatment !=
    "completely_untreated", ]
```

Create yet another table, where each single observation period for a female gets an own row

```
col_names1 <- c("Eggs_with_males", "Eggs_without_males",
    "female", "observation_time_1",
    "court_time_1", "treatment", "contr_exp")
col_names2 <- c("Eggs_with_males", "Eggs_without_males",
```

```
    "female", "observation_time_2",
    "court_time_2", "treatment", "contr_exp")
col_names_new <- c("Eggs_with_males",
    "Eggs_without_males", "female",
    "obs_sec", "harass_sec", "treat1",
    "treat2")
erato_data_harass_glmm <- do.call(rbind,
    lapply(list(erato_data1[!is.na(erato_data1$observation_time_1),
        col_names1], erato_data1[!is.na(erato_data1$observation_time_2),
        col_names2]), function(x) {
        colnames(x) <- col_names_new
        x
    }))
```

Minimum, median and maximum for observation time per female:

```
# To make the Markdown prettier:
f_id <- erato_data_harass_glmm$female
print(quantile(sapply(unique(f_id),
    function(x) sum(erato_data_harass_glmm$obs_sec[f_id ==
        x])), c(0, 0.5, 1)))
```

```
##     0%    50%   100%
##  929.0 3607.5 7207.0
```

Add a new column where we calculate the number of seconds with harassment

```
erato_data_harass_glmm$no_harass_sec <- erato_data_harass_glmm$obs_sec -
    erato_data_harass_glmm$harass_sec
```

Test if females in the two control groups (transparent marker and only handled) differ. They do not:

```
joint_tests(glmer(cbind(harass_sec,
    no_harass_sec) ~ treat1 + (1 | female),
    family = "binomial", data = erato_data_harass_glmm[erato_data_harass_glmm$treat2 !=
        "treatment", ]))
```

Repeat the same, but excluding females that didn't lay

```
joint_tests(glmer(cbind(harass_sec,
    no_harass_sec) ~ treat1 + (1 | female),
    family = "binomial", data = erato_data_harass_glmm[erato_data_harass_glmm$treat2 !=
        "treatment" & (erato_data_harass_glmm$Eggs_with_males +
        erato_data_harass_glmm$Eggs_without_males) >
        0, ]))
```

Both tests reveal that the control groups do not differ, and hence, we will unite them in the following sections.

Now we calculate or each female the proportion of observation time that it was harassed

```
uni_female <- sort(unique(f_id))
pref_by_female_erato <- data.frame(female = uni_female,
    eggs_laid = sapply(uni_female, function(x) sum(erato_data_harass_glmm[f_id ==
        x, c("Eggs_with_males", "Eggs_without_males")][1,
        ])), treatment = sapply(uni_female,
        function(x) erato_data_harass_glmm$treat2[f_id ==
            x][1]), Prop_harass = sapply(uni_female,
        function(x) sum(erato_data_harass_glmm$harass_sec[f_id ==
```

```
            x])/sum(erato_data_harass_glmm$obs_sec[f_id ==
            x])))
```

We look at this with a boxplot

```
par(mar = c(2.5, 4, 0.2, 0.2))
boxplot(Prop_harass ~ treatment, data = pref_by_female_erato,
    xlab = "", ylab = "Proportion of observation time")
```

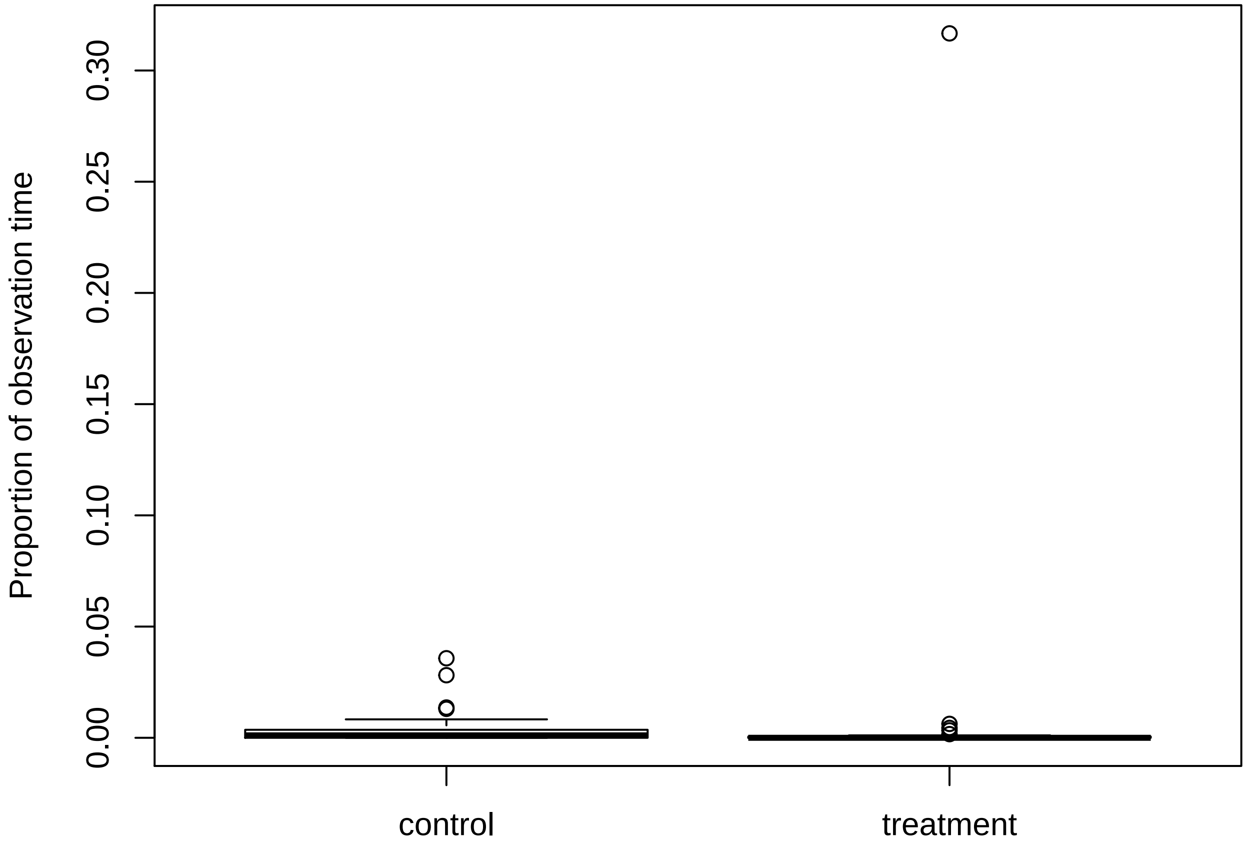


We see some quite extreme outliers! The most extreme point in the treatment group for example is 570x units of interquartile range (IQR) away from the 25-75% range. Usually, values >1.5 units away are considered outliers. These females will not allow a balanced data analysis. Now we search for these females, separately for the control and the treatment group! We extract all females that are more than 1.5 units of IQR away from the 25-75% range.

```
# For making the Markdown prettier,
# name some variables:
t_id <- pref_by_female_erato$treatment
e_laid <- pref_by_female_erato$eggs_laid
prop_h <- pref_by_female_erato$Prop_harass
outliers_by_group_erato <- sapply(c("control",
    "treatment"), function(x) boxplot.stats(prop_h[t_id ==
    x])$out)
# We repeat the same analysis
# excluding females that laid 0 eggs
outliers_by_group_erato_no0 <- sapply(c("control",
    "treatment"), function(x) boxplot.stats(prop_h[e_laid >
```

```
    0 & t_id == x])$out)
# We can see that both dataset yield
# the exact same outliers:
all.equal(outliers_by_group_erato, outliers_by_group_erato_no0)
```

```
## [1] TRUE
```

Give name of female to each of these outliers

```
for (renamer in 1:2) {
    names(outliers_by_group_erato[[renamer]]) <- sapply(outliers_by_group_erato[[renamer]],
        function(x) pref_by_female_erato$female[prop_h ==
            x])
}
```

Let's have a look at these outlier values

```
print(sapply(1:length(outliers_by_group_erato),
    function(x) round(outliers_by_group_erato[[x]],
        3)))
```

```
## [[1]]
##  M110   M66   M75   M78
## 0.014 0.028 0.036 0.013
##
## [[2]]
##  M103   M47   M53   M61   M99
## 0.317 0.006 0.003 0.002 0.004
```

We create a new dataset without these outliers:

```
erato_data_harass_glmm_no_outliers <- erato_data_harass_glmm[!f_id %in%
    unlist(sapply(1:length(outliers_by_group_erato),
        function(x) names(outliers_by_group_erato[[x]]))),
    ]
```

First, run models excluding females that laid no eggs. Create these datasets:

```
excl0 <- rowSums(erato_data_harass_glmm[,
    c("Eggs_with_males", "Eggs_without_males")]) >
    0
excl0_no_outl <- rowSums(erato_data_harass_glmm_no_outliers[,
    c("Eggs_with_males", "Eggs_without_males")]) >
    0
erato_data_harass_glmm_excl0 <- erato_data_harass_glmm[excl0,
    ]
erato_data_harass_glmm_no_outliers_excl0 <- erato_data_harass_glmm_no_outliers[excl0_no_outl,
    ]
```

We run a model with and without outliers

```
erato_harass_glmm <- glmer(cbind(harass_sec,
    no_harass_sec) ~ treat2 + (1 | female),
    family = "binomial", data = erato_data_harass_glmm_excl0)
erato_harass_glmm_2 <- glmer(cbind(harass_sec,
    no_harass_sec) ~ treat2 + (1 | female),
    family = "binomial", data = erato_data_harass_glmm_no_outliers_excl0)
```

Retrieve the EMMs with and without outliers

```r
(erato_harass_emm <- summary(emmeans(erato_harass_glmm,
    ~treat2, transform = "response")))
```

```r
(erato_harass_emm2 <- summary(emmeans(erato_harass_glmm_2,
    ~treat2, transform = "response")))
```

Check for significance with and without outliers

```r
(test_for_panelA_suppl <- joint_tests(erato_harass_glmm))
```

```r
(test_for_panelA <- joint_tests(erato_harass_glmm_2))
```

Confirm that rerunning these GLMMs and significance tests including females that laid no eggs gives similar results

Models with and without outliers

```r
erato_harass_glmm_incl0 <- glmer(cbind(harass_sec,
    no_harass_sec) ~ treat2 + (1 | female),
    family = "binomial", data = erato_data_harass_glmm)
erato_harass_glmm_2_incl0 <- glmer(cbind(harass_sec,
    no_harass_sec) ~ treat2 + (1 | female),
    family = "binomial", data = erato_data_harass_glmm_no_outliers)
```

Retrieve the EMMs with and without outliers

```r
emmeans(erato_harass_glmm_incl0, ~treat2,
    transform = "response")
```

```
##  treat2        prob       SE  df asymp.LCL asymp.UCL
##  control   3.70e-04 2.85e-04 Inf -1.88e-04  0.000928
##  treatment 3.55e-05 3.49e-05 Inf -3.29e-05  0.000104
##
## Confidence level used: 0.95
```

```r
emmeans(erato_harass_glmm_2_incl0, ~treat2,
    transform = "response")
```

```
##  treat2        prob       SE  df asymp.LCL asymp.UCL
##  control   3.65e-04 2.15e-04 Inf -5.72e-05  7.87e-04
##  treatment 2.27e-05 1.96e-05 Inf -1.58e-05  6.11e-05
##
## Confidence level used: 0.95
```

Check for significance with and without outliers

```r
joint_tests(erato_harass_glmm_incl0)
```

```r
joint_tests(erato_harass_glmm_2_incl0)
```

We get largely the same results.

###### 4.4.2 Experiment 2 (with *H. timareta*)

For the *H. timareta* dataset, no female ever laid 0 eggs during the complete experiment, so we don’t have to repeat analyses including and excluding females that didn’t lay eggs. Some females, however, have NA values in the eggs columns. These females were only tested for male harassment, but their egg count not systematically assessed (but all of them lay eggs).

```r
min(rowSums(timareta_data[, c("Eggs_with_males",
    "Eggs_without_males")]), na.rm = T)
```

```
## [1] 3
```

Again, we create a table, where each single observation period for a female gets an own row.

```r
col_names1 <- c("female", "observation_time_1",
    "court_time_1", "wing_colour", "yellow_rel",
    "red_rel")
col_names2 <- c("female", "observation_time_2",
    "court_time_2", "wing_colour", "yellow_rel",
    "red_rel")
col_names_new <- c("female", "obs_sec",
    "harass_sec", "wing_colour", "yellow_rel",
    "red_rel")
timareta_data_harass_glmm <- do.call(rbind,
    lapply(list(timareta_data[!is.na(timareta_data$observation_time_1),
        col_names1], timareta_data[!is.na(timareta_data$observation_time_2),
        col_names2]), function(x) {
        colnames(x) <- col_names_new
        x
    }))
```

Minimum, median and maximum for observation time per female:

```r
# To make the Markdown prettier:
f_id <- timareta_data_harass_glmm$female
print(quantile(sapply(unique(f_id),
    function(x) sum(timareta_data_harass_glmm$obs_sec[f_id ==
        x])), c(0, 0.5, 1)))
```

```
##   0%  50% 100%
## 1722 3547 3904
```

Again, add a new column where we calculate the number of seconds with harassment

```r
timareta_data_harass_glmm$no_harass_sec <- timareta_data_harass_glmm$obs_sec -
    timareta_data_harass_glmm$harass_sec
```

Again, we calculate or each female the proportion of observation time that it was harassed

```r
uni_female <- sort(unique(f_id))
pref_by_female_timareta <- data.frame(female = uni_female,
    treatment = sapply(uni_female, function(x) timareta_data_harass_glmm$wing_colour[f_id ==
        x][1]), Prop_harass = sapply(uni_female,
        function(x) sum(timareta_data_harass_glmm$harass_sec[f_id ==
            x])/sum(timareta_data_harass_glmm$obs_sec[f_id ==
            x])))
```

We look at this with a boxplot

```r
par(mar = c(2.5, 4, 0.2, 0.2))
boxplot(Prop_harass ~ treatment, data = pref_by_female_timareta,
    xlab = "", ylab = "Proportion of observation time")
```

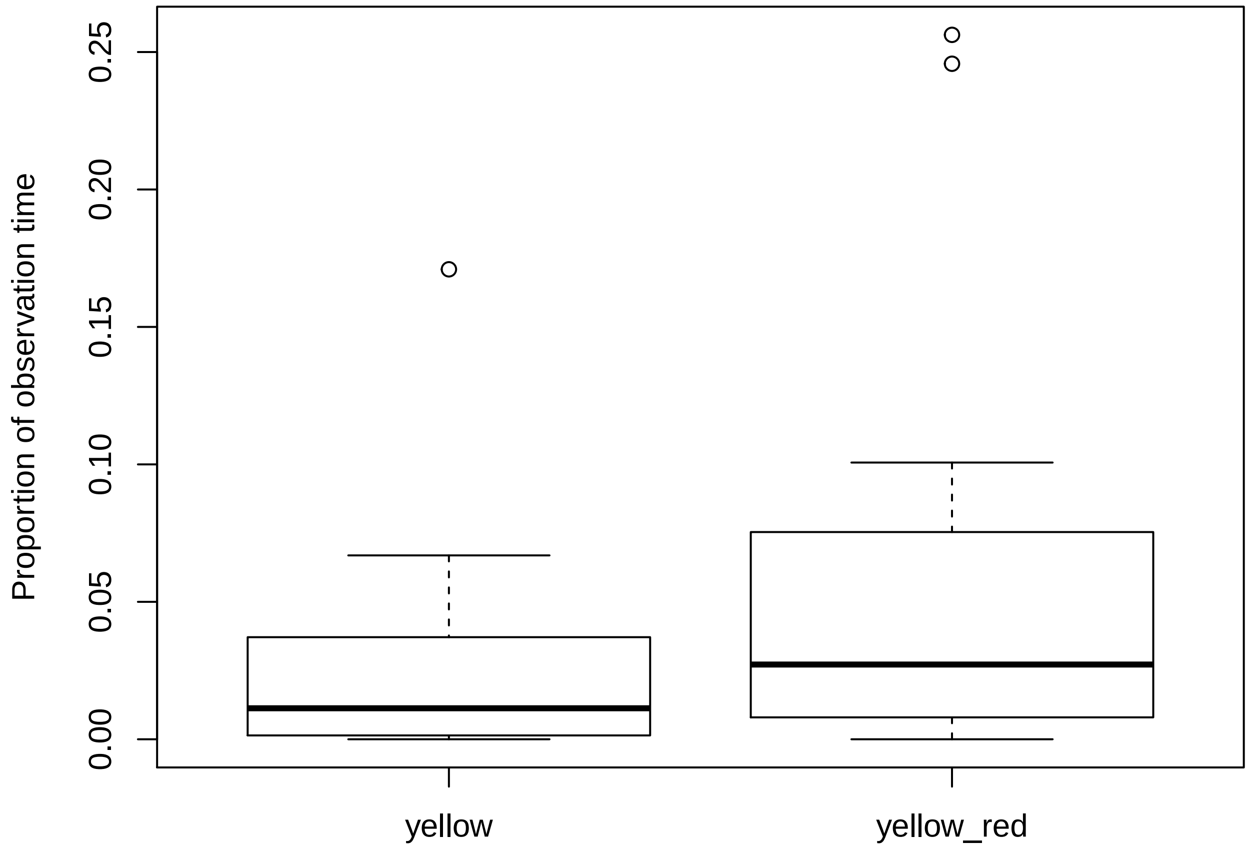


Again, we see some quite extreme outliers! They are not as extreme as before (the most extreme being 3.74 units of IQR away), but it makes still sense to exclude these. We apply the same rules as before:

```
# For making the Markdown prettier,
# name some variables:
t_id <- pref_by_female_timareta$treatment
prop_h <- pref_by_female_timareta$Prop_harass
outliers_by_group_timareta <- sapply(c("yellow",
    "yellow_red"), function(x) boxplot.stats(prop_h[t_id ==
    x])$out)
```

Give name of female to each of these outliers

```
for (renamer in 1:2) {
    names(outliers_by_group_timareta[[renamer]]) <- sapply(outliers_by_group_timareta[[renamer]],
        function(x) pref_by_female_timareta$female[prop_h ==
            x])
}
```

Let's have a look at these outlier values

```
print(sapply(1:length(outliers_by_group_timareta),
    function(x) round(outliers_by_group_timareta[[x]],
        3)))
```

```
## [[1]]
## A18_809
##   0.171
##
```

```
## [[2]]
## A18_800 A18_819
##   0.256   0.246
```

We create a new dataset without these outliers:

```
timareta_data_harass_glmm_no_outliers <- timareta_data_harass_glmm[!f_id %in%
    unlist(sapply(1:length(outliers_by_group_timareta),
        function(x) names(outliers_by_group_timareta[[x]]))),
    ]
```

We run a model with and without outliers

```
timareta_harass_glmm <- glmer(cbind(harass_sec,
    no_harass_sec) ~ wing_colour + (1 |
    female), family = "binomial", data = timareta_data_harass_glmm)
timareta_harass_glmm_2 <- glmer(cbind(harass_sec,
    no_harass_sec) ~ wing_colour + (1 |
    female), family = "binomial", data = timareta_data_harass_glmm_no_outliers)
```

Retrieve the EMMs with and without outliers

```
(timareta_harass_emm <- summary(emmeans(timareta_harass_glmm,
    ~wing_colour, transform = "response")))
```

```
(timareta_harass_emm2 <- summary(emmeans(timareta_harass_glmm_2,
    ~wing_colour, transform = "response")))
```

Check for significance with and without outliers

```
(test_for_panelB_suppl <- joint_tests(timareta_harass_glmm))
```

```
(test_for_panelB <- joint_tests(timareta_harass_glmm_2))
```

We can see here that excluding the outliers does seem to make a bit of a difference…

Finally, we fit the same type of model structure as above to the dataset without outliers, but this time considering size of the yellow patch on the wing (relative to the whole wing size, either including all individuals or only females with red on the wings), and/or size of the red patch (relative to the whole wing size, and only including females with red). Presence/absence of the red band might have an effect on how big the yellow band gets, and as we assume that males use this yellow band as a cue to detect females, bigger bands might be more attractive to males. Similarly, red bands might be unattractive, and the bigger the red band is, the more unattractive a female might be. We look at the corrected values (*i.e.* dividing by whole wing area) in order to correct for overall female size. While the absolute size of the bands might be a more important cue for males, both band sizes are correlated with total wing size, which might be a proxy for female fitness (*i.e.* bigger wings being more fit). Hence, to avoid any biases there, we look at relative size.

First, we have to make sure that among females with red on the wings, the two predictors are not strongly correlated (e.g., we can imagine that a bigger red band means a smaller red band). This plot is similar to the overview we created before, plotting photos of the actual wings, only that here, we exclude females without red on the wings. Also, a few individuals of the previous plot do not appear here, as no egg count could be performed on them.

For plotting, we create another dataset that has the values summarized by female ID (*i.e.* kind of reversing what we did above, where we split the dataset into single observations):

```
# The variable name
# timareta_data_harass_glmm_no_outliers
# is too long for the Markdown in
# this section. For this and the
```

```
# next sextions, we just rename it
# yy
yy <- timareta_data_harass_glmm_no_outliers
uniq_female <- unique(yy$female)
yy_plot <- data.frame(female = uniq_female,
    obs_sec = sapply(uniq_female, function(x) sum(yy$obs_sec[yy$female ==
        x])), harass_sec = sapply(uniq_female,
        function(x) sum(yy$harass_sec[yy$female ==
            x])), wing_colour = sapply(uniq_female,
        function(x) yy$wing_colour[yy$female ==
            x][1]), yellow_rel = sapply(uniq_female,
        function(x) yy$yellow_rel[yy$female ==
            x][1]), red_rel = sapply(uniq_female,
        function(x) yy$red_rel[yy$female ==
            x][1]), stringsAsFactors = F)
```

Now, look at possible collinearity. Note that no strong correlation can be seen, hence, both can be included as covariates in our model:

```
par(mar = c(3.5, 4, 0.2, 0.2))
plot(yy_plot$yellow_rel[yy_plot$wing_colour ==
    "yellow_red"], yy_plot$red_rel[yy_plot$wing_colour ==
    "yellow_red"], pch = 21, bg = adjustcolor("red",
    0.75), xaxt = "n", yaxt = "n", xlab = "",
    ylab = "")
axis(1, lwd = 0, line = -0.8, cex.axis = 0.8)
axis(1, labels = F, tck = -0.01)
axis(2, lwd = 0, line = -0.65, las = 2,
    cex.axis = 0.8)
axis(2, labels = F, tck = -0.01)
mtext("Proportion yellow band", 1, line = 1.8,
    cex = 1.15)
mtext("Proportion red band", 2, line = 2.3,
    cex = 1.15)
```

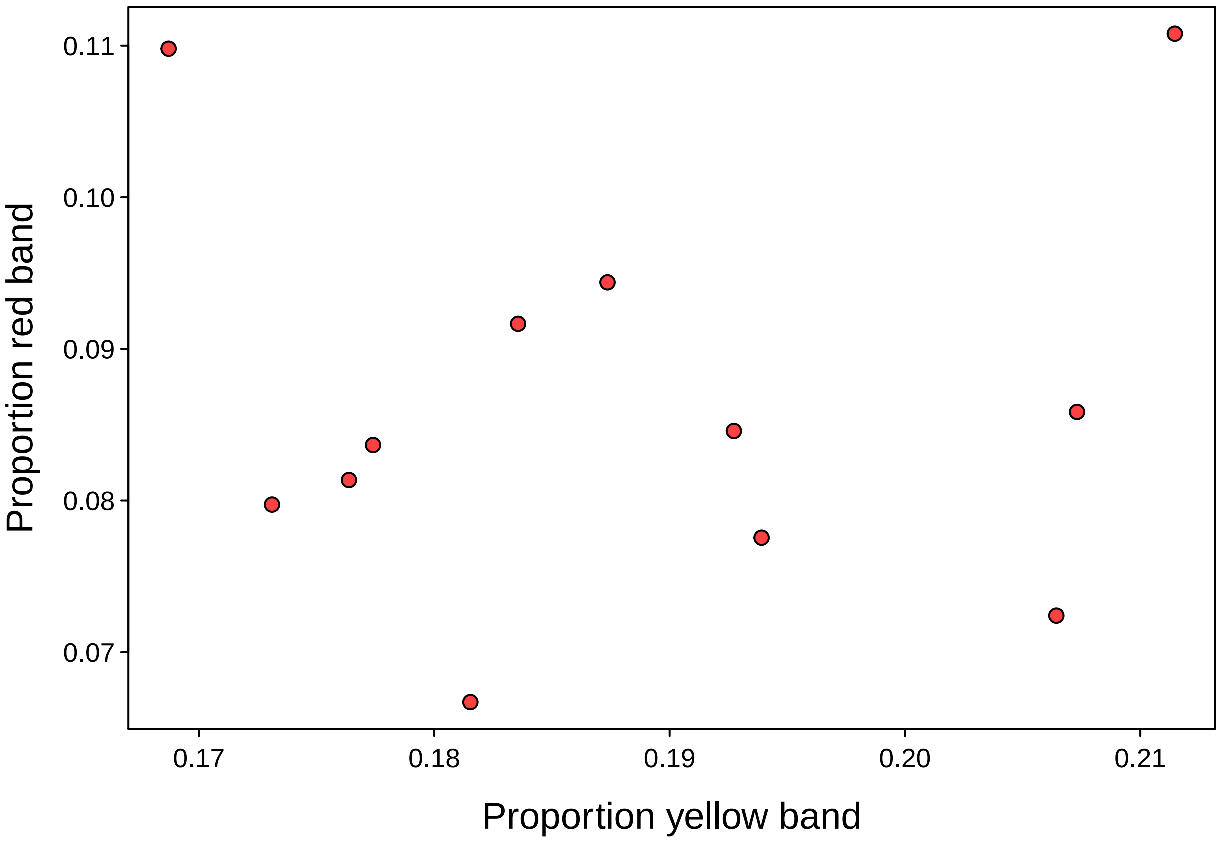


For modelling, create a dataset with only females with red on the wings:

```
yy_red <- yy[yy$wing_colour == "yellow_red",
    ]
```

We fit two models: * Model 1: testing for proportion of yellow band across all females * Model 2: testing for proportion of yellow band and proportion of red band as separate factors, only including females with red on the wing

```
timareta_harass_glmm_yellow <- glmer(cbind(harass_sec,
    no_harass_sec) ~ yellow_rel + (1 |
    female), family = "binomial", data = yy)
timareta_harass_glmm_red_and_yellow <- glmer(cbind(harass_sec,
    no_harass_sec) ~ red_rel + yellow_rel +
    (1 | female), family = "binomial",
    data = yy_red)
```

We test for significance of the slopes:

```
harass_trend_yellow <- emtrends(timareta_harass_glmm_yellow,
    ~yellow_rel, var = "yellow_rel")
print("Slope proportion yellow")
```

```
## [1] "Slope proportion yellow"
```

```
(harass_p_yellow <- summary(harass_trend_yellow,
    infer = c(TRUE, TRUE), null = 0,
    side = "two-sided"))
```

```r
harass_trend_red <- emtrends(timareta_harass_glmm_red_and_yellow,
    ~red_rel, var = "red_rel")
print("Slope proportion red (only individuals with red)")
```

```
## [1] "Slope proportion red (only individuals with red)"
```

```r
(harass_p_red <- summary(harass_trend_red,
    infer = c(TRUE, TRUE), null = 0,
    side = "two-sided"))
```

```r
harass_trend_yellow_red_indiv <- emtrends(timareta_harass_glmm_red_and_yellow,
    ~yellow_rel, var = "yellow_rel")
print("Slope proportion yellow (only individuals with red)")
```

```
## [1] "Slope proportion yellow (only individuals with red)"
```

```r
(harass_p_yellow_red_indiv <- summary(harass_trend_yellow_red_indiv,
    infer = c(TRUE, TRUE), null = 0,
    side = "two-sided"))
```

We get a significantly negative slope for the test on size of the red band. Let's plot the effect of yellow band and red band proportion on male harassment, for females that show red on the wings. First, we extract the model estimates:

```r
harass_pred_red <- emmip(timareta_harass_glmm_red_and_yellow,
    ~red_rel, transform = "response",
    CI = T, plotit = F, at = list(red_rel = seq(min(yy_red$red_rel,
        na.rm = T), max(yy_red$red_rel,
        na.rm = T), length.out = 100)))
harass_pred_yellow <- emmip(timareta_harass_glmm_red_and_yellow,
    ~yellow_rel, transform = "response",
    CI = T, plotit = F, at = list(yellow_rel = seq(min(yy_red$yellow_rel,
        na.rm = T), max(yy_red$yellow_rel,
        na.rm = T), length.out = 100)))
```

We plot this:

```r
png("Figures/empirical_proportion_red_harassment.png",
    width = 2700, height = 3600, res = 300)
layout(matrix(1:2, ncol = 1))
par(mar = c(3, 3.3, 1.8, 0.2))

# Range of y-values
yra <- range(c((yy_plot$harass_sec/yy_plot$obs_sec)[yy_plot$wing_colour ==
    "yellow_red"], harass_pred_red$LCL,
    harass_pred_red$UCL), na.rm = T)

# Effect of red

# Empty plot
plot(1, 1, type = "n", xaxt = "n", yaxt = "n",
    xlab = "", ylab = "", xlim = range(yy_plot$red_rel[yy_plot$wing_colour ==
        "yellow_red"], na.rm = T), ylim = yra)
# Add confidence interval of
# estimate
polygon(c(harass_pred_red$red_rel, rev(harass_pred_red$red_rel)),
    c(harass_pred_red$LCL, rev(harass_pred_red$UCL)),
```

```
    border = NA, col = adjustcolor("black",
        0.3))
# Add estimate
lines(harass_pred_red$red_rel, harass_pred_red$yvar,
    lwd = 2)
# Add data points
points(yy_plot$red_rel[yy_plot$wing_colour ==
    "yellow_red"], (yy_plot$harass_sec/yy_plot$obs_sec)[yy_plot$wing_colour ==
    "yellow_red"], pch = 21, bg = adjustcolor("red",
    0.75), cex = 0.02 * sqrt(yy_plot$obs_sec[yy_plot$wing_colour ==
    "yellow_red"]))
# Add axes, p-value, titles and
# panel ID
axis(1, lwd = 0, line = -0.8, cex.axis = 0.8)
axis(1, labels = F, tck = -0.01)
axis(2, lwd = 0, line = -0.65, las = 2,
    cex.axis = 0.8)
axis(2, labels = F, tck = -0.01)
mtext("Proportion red band", 1, line = 1.8,
    cex = 1.15)
mtext(bquote(italic(p) ~ .(ifelse(harass_p_red$p.value <
    0.001, "< 0.001", paste0("= ", round(harass_p_red$p.value,
    3))))), 3, line = 0, cex = 1.15)
text(par("usr")[1] + 0.97 * diff(par("usr")[1:2]),
    par("usr")[4] - 0.07 * diff(par("usr")[3:4]),
    "A", font = 2, cex = 1.7, pos = 2,
    offset = 0)


# Effect of yellow (basically the
# same)

plot(1, 1, type = "n", xaxt = "n", yaxt = "n",
    xlab = "", ylab = "", xlim = range(yy_plot$yellow_rel[yy_plot$wing_colour ==
        "yellow_red"], na.rm = T), ylim = yra)
polygon(c(harass_pred_yellow$yellow_rel,
    rev(harass_pred_yellow$yellow_rel)),
    c(harass_pred_yellow$LCL, rev(harass_pred_yellow$UCL)),
    border = NA, col = adjustcolor("black",
        0.3))
lines(harass_pred_yellow$yellow_rel,
    harass_pred_yellow$yvar, lwd = 2)
points(yy_plot$yellow_rel[yy_plot$wing_colour ==
    "yellow_red"], (yy_plot$harass_sec/yy_plot$obs_sec)[yy_plot$wing_colour ==
    "yellow_red"], pch = 21, bg = adjustcolor("red",
    0.75), cex = 0.02 * sqrt(yy_plot$obs_sec[yy_plot$wing_colour ==
    "yellow_red"]))
axis(1, lwd = 0, line = -0.8, cex.axis = 0.8)
axis(1, labels = F, tck = -0.01)
axis(2, lwd = 0, line = -0.65, las = 2,
    cex.axis = 0.8)
axis(2, labels = F, tck = -0.01)
mtext("Proportion yellow band", 1, line = 1.8,
```

```
    cex = 1.15)
mtext(bquote(italic(p) ~ .(ifelse(harass_p_yellow_red_indiv$p.value <
    0.001, "< 0.001", paste0("= ", round(harass_p_yellow_red_indiv$p.value,
    3))))), 3, line = 0, cex = 1.15)
text(par("usr")[1] + 0.97 * diff(par("usr")[1:2]),
    par("usr")[4] - 0.07 * diff(par("usr")[3:4]),
    "B", font = 2, cex = 1.7, pos = 2,
    offset = 0)

mtext("Proportion of observation time with male harassment",
    2, line = -1, cex = 1.15, outer = T)

invisible(dev.off())
```

Show png we just made:

```
fig_disp <- readPNG("Figures/empirical_proportion_red_harassment.png")
grid.raster(fig_disp)
```

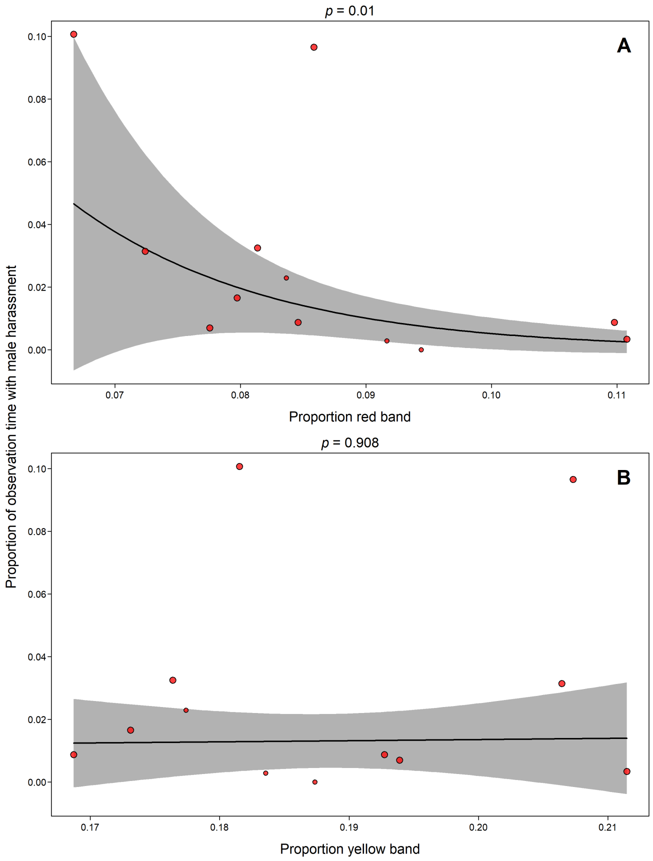


#### 4.5 Analyses of female fecundity

##### 4.5.1 Experiment 1 (with *H. erato*)

Add column indicating females that either have no egg count performed or laid in total zero eggs.

```
erato_data_eggs <- erato_data
erato_data_eggs$laid0 <- (ifelse(is.na(erato_data$Eggs_with_males),
```

```
    T, erato_data$Eggs_with_males ==
        0) + ifelse(is.na(erato_data$Eggs_without_males),
    T, erato_data$Eggs_without_males ==
        0)) == 2
```

Add a column calculating the difference between eggs with and without male

```
erato_data_eggs$egg_difference <- erato_data_eggs$Eggs_with_males -
    erato_data_eggs$Eggs_without_males
```

As before, we create a new table, where we bloat up the table, by giving each egg count from one of the phases an own row. We also add a column denoting whether males were present or not.

```
col_names1 <- c("female", "Eggs_with_males",
    "treatment", "contr_exp", "laid0")
col_names2 <- c("female", "Eggs_without_males",
    "treatment", "contr_exp", "laid0")
col_names_new <- c("female", "eggs",
    "treat1", "treat2", "laid0")
erato_data_eggs_glmm <- data.frame(do.call(rbind,
    lapply(list(erato_data_eggs[, col_names1],
        erato_data_eggs[, col_names2]),
        function(x) {
            colnames(x) <- col_names_new
            x
        })), Male_presence = c(rep("present",
    nrow(erato_data_eggs)), rep("not_present",
    nrow(erato_data_eggs))), stringsAsFactors = F)
```

Test if females in the two control groups (transparent marker and only handled) differ. We use a Poisson model, with egg count per 48h period as response variable. Exclude for now females from the 'completely_untreated' group.

```
joint_tests(glmer(eggs ~ treat1 * Male_presence +
    (1 | female), family = "poisson",
    data = erato_data_eggs_glmm[erato_data_eggs_glmm$treat2 !=
        "treatment" & erato_data_eggs_glmm$treat1 !=
        "completely_untreated", ]))
```

Repeat the same, but excluding females that didn't lay

```
joint_tests(glmer(eggs ~ treat1 * Male_presence +
    (1 | female), family = "poisson",
    data = erato_data_eggs_glmm[erato_data_eggs_glmm$treat2 !=
        "treatment" & erato_data_eggs_glmm$treat1 !=
        "completely_untreated" & !erato_data_eggs_glmm$laid0,
        ]))
```

The two controls do not differ (neither effect of treatment alone, nor interaction with male presence). We unite them for further analyses.

Now we test whether there is an interaction between warning pattern and male presence. We exclude completely "untreated" females. We exclude females that didn't lay from here, but below, we show that including them doesn't make a difference.

```
erato_eggs_glmm <- glmer(eggs ~ treat2 *
    Male_presence + (1 | female), family = "poisson",
    data = erato_data_eggs_glmm[erato_data_eggs_glmm$treat1 !=
```

```
        "completely_untreated" & !erato_data_eggs_glmm$laid0,
        ])
```

We look at the estimators

```
print(emmeans(erato_eggs_glmm, ~Male_presence |
    treat2, transform = "response"))
```

```
## treat2 = control:
##  Male_presence rate    SE  df asymp.LCL asymp.UCL
##  not_present   5.29 0.474 Inf      4.36      6.22
##  present       5.04 0.457 Inf      4.15      5.94
##
## treat2 = treatment:
##  Male_presence rate    SE  df asymp.LCL asymp.UCL
##  not_present   5.75 0.508 Inf      4.75      6.75
##  present       4.80 0.444 Inf      3.93      5.66
##
## Confidence level used: 0.95
```

We can see that both groups laid similar amounts of eggs. However, both had quite a drastic reduction in eggs when males were present.

We look at the effect sizes:

```
erato_eggs_eff_size <- summary(emmeans(erato_eggs_glmm,
    pairwise ~ Male_presence | treat2,
    transform = "response")$contrasts,
    infer = T)
# Inverse the numbers (as emmeans
# calculate without males - with
# males, but we want it the other
# way around)
erato_eggs_eff_size[, c("estimate",
    "asymp.LCL", "asymp.UCL")] <- erato_eggs_eff_size[,
    c("estimate", "asymp.LCL", "asymp.UCL")] *
    (-1)
# Aesthetical changes of the table
names(erato_eggs_eff_size)[6:7] <- names(erato_eggs_eff_size)[7:6]
erato_eggs_eff_size <- erato_eggs_eff_size[,
    c(1:5, 7, 6, 8:9)]
erato_eggs_eff_size$contrast <- as.character("present - not_present")
erato_eggs_eff_size[, -(1:2)] <- round(erato_eggs_eff_size[,
    -(1:2)], 3)
# Print
print(erato_eggs_eff_size)
```

```
##                contrast    treat2 estimate    SE  df asymp.LCL asymp.UCL
## 1 present - not_present   control   -0.248 0.458 Inf    -1.146     0.650
## 2 present - not_present treatment   -0.955 0.478 Inf    -1.891    -0.018
##   z.ratio p.value
## 1   0.541   0.588
## 2   1.997   0.046
```

The reduction in eggs can be seen in both groups, but surprisingly the stronger effect is in the treatment group.

Let's see which of the factors of the model are significant:

```
joint_tests(erato_eggs_glmm)
```

Male presence seems to be the only factor really playing a role.

We briefly confirm that we get the same result when including females that didn't lay:

```
joint_tests(glmer(eggs ~ treat2 * Male_presence +
    (1 | female), family = "poisson",
    data = erato_data_eggs_glmm[erato_data_eggs_glmm$treat1 !=
        "completely_untreated", ]))
```

We now test for the effect of male presence in isolation, this time including those females that had no warning pattern treatment (and that were excluded so far); also, again, excluding females that didn't lay (again, the effect of including them will be tested below):

```
erato_eggs_glmm_males <- glmer(eggs ~
    Male_presence + (1 | female), family = "poisson",
    data = erato_data_eggs_glmm[!erato_data_eggs_glmm$laid0,
        ])
```

We look at the estimators

```
(erato_estimates <- emmeans(erato_eggs_glmm_males,
    ~Male_presence, transform = "response"))
```

```
##  Male_presence rate    SE  df asymp.LCL asymp.UCL
##  not_present   5.45 0.322 Inf      4.82      6.08
##  present       4.72 0.289 Inf      4.16      5.29
##
## Confidence level used: 0.95
```

Again, we look at the effect sizes:

```
erato_eggs_eff_size_PLOT <- summary(emmeans(erato_eggs_glmm_males,
    pairwise ~ Male_presence, transform = "response")$contrasts,
    infer = T)
# Inverse again the numbers
erato_eggs_eff_size_PLOT[, c("estimate",
    "asymp.LCL", "asymp.UCL")] <- erato_eggs_eff_size_PLOT[,
    c("estimate", "asymp.LCL", "asymp.UCL")] *
    (-1)
# Aesthetical changes of the table
names(erato_eggs_eff_size_PLOT)[5:6] <- names(erato_eggs_eff_size_PLOT)[6:5]
erato_eggs_eff_size_PLOT <- erato_eggs_eff_size_PLOT[,
    c(1:5, 6, 5, 7:8)]
erato_eggs_eff_size_PLOT$contrast <- as.character("present - not_present")
erato_eggs_eff_size_PLOT[, -(1:2)] <- round(erato_eggs_eff_size_PLOT[,
    -(1:2)], 3)
# Print
print(erato_eggs_eff_size_PLOT)
```

```
##                  contrast   estimate    SE  df asymp.UCL asymp.LCL asymp.UCL.1
## 1 present - not_present -0.7293273 0.303 Inf    -0.135    -1.324      -0.135
##   z.ratio p.value
## 1   2.404   0.016
```

Testing for the overall effect of male presence will give the same p-value as the one printed above:

```
(test_for_panelA_eggs <- joint_tests(erato_eggs_glmm_males))
```

Again, briefly checking if including those that didn't lay makes a difference:

```
joint_tests(glmer(eggs ~ Male_presence +
    (1 | female), family = "poisson",
    data = erato_data_eggs_glmm))
```

We create ourselves a brief summary of the data, first with the actual raw numbers (again, we exclude those that never laid here):

```
# Text to paste
t1 <- "Mean with male = "
t2 <- "Mean without male = "
t3 <- "Mean difference = "
t4 <- "Without male x % more than with male = "
t5 <- "With male x % less than without male = "
wo_males <- erato_data_eggs$Eggs_without_males[!erato_data_eggs$laid0]
w_males <- erato_data_eggs$Eggs_with_males[!erato_data_eggs$laid0]
paste0(t1, round(mean(w_males), 2))
```

```
## [1] "Mean with male = 5.13"
```

```
paste0(t2, round(mean(wo_males), 2))
```

```
## [1] "Mean without male = 5.92"
```

```
paste0(t3, round(mean(erato_data_eggs$egg_difference[!erato_data_eggs$laid0]),
    2))
```

```
## [1] "Mean difference = -0.79"
```

```
paste0(t4, round(sum(wo_males)/sum(w_males) -
    1, 2) * 100, "%")
```

```
## [1] "Without male x % more than with male = 15%"
```

```
paste0(t5, round(sum(w_males)/sum(wo_males) -
    1, 2) * 100, "%")
```

```
## [1] "With male x % less than without male = -13%"
```

We repeat the same with model estimates:

```
wo_males <- summary(erato_estimates)$rate[summary(erato_estimates)$Male_presence ==
    "not_present"]
w_males <- summary(erato_estimates)$rate[summary(erato_estimates)$Male_presence ==
    "present"]
paste0(t1, round(w_males, 2))
```

```
## [1] "Mean with male = 4.72"
```

```
paste0(t2, round(wo_males, 2))
```

```
## [1] "Mean without male = 5.45"
```

```
paste0(t3, round(erato_eggs_eff_size_PLOT$estimate,
    2))
```

```
## [1] "Mean difference = -0.73"
```

```
paste0(t4, round(wo_males/w_males -
    1, 2) * 100, "%")
```

```
## [1] "Without male x % more than with male = 15%"
```

```
paste0(t5, round(w_males/wo_males -
    1, 2) * 100, "%")
```

```
## [1] "With male x % less than without male = -13%"
```

##### 4.5.2 Experiment 2 (with *H. timareta*)

For the *H. timareta* data, we again don't have to worry about females that never laid eggs, as all of them did. However, as previously mentioned, some females have an NA in the egg columns. We discard these right away:

```
timareta_data_eggs <- timareta_data[!is.na(timareta_data$Eggs_with_males) &
    !is.na(timareta_data$Eggs_without_males),
    ]
```

Add a column calculating the difference between eggs with and without male

```
timareta_data_eggs$egg_difference <- timareta_data_eggs$Eggs_with_males -
    timareta_data_eggs$Eggs_without_males
```

As before, we create a new table, where we bloat up the table, by giving each egg count from one of the phases an own row. We also add a column denoting whether males were present or not.

```
col_names1 <- c("female", "Eggs_with_males",
    "wing_colour", "yellow_rel", "red_rel")
col_names2 <- c("female", "Eggs_without_males",
    "wing_colour", "yellow_rel", "red_rel")
col_names_new <- c("female", "eggs",
    "wing_colour", "yellow_rel", "red_rel")
timareta_data_eggs_glmm <- data.frame(do.call(rbind,
    lapply(list(timareta_data_eggs[,
        col_names1], timareta_data_eggs[,
        col_names2]), function(x) {
        colnames(x) <- col_names_new
        x
    })), Male_presence = c(rep("present",
    nrow(timareta_data_eggs)), rep("not_present",
    nrow(timareta_data_eggs))), stringsAsFactors = F)
```

We test whether there is an interaction between warning pattern and male presence.

```
timareta_eggs_glmm <- glmer(eggs ~ wing_colour *
    Male_presence + (1 | female), family = "poisson",
    data = timareta_data_eggs_glmm)
```

We look at the estimators

```
print(emmeans(timareta_eggs_glmm, ~Male_presence |
    wing_colour, transform = "response"))
```

```
## wing_colour = yellow:
##  Male_presence rate    SE  df asymp.LCL asymp.UCL
##  not_present   8.91 0.815 Inf      7.32     10.51
##  present       7.17 0.684 Inf      5.83      8.51
##
```

```
## wing_colour = yellow_red:
##  Male_presence rate    SE  df asymp.LCL asymp.UCL
##  not_present   8.80 0.841 Inf      7.15     10.45
##  present       7.14 0.712 Inf      5.74      8.54
##
## Confidence level used: 0.95
```

Again, we can see that both groups laid similar amounts of eggs. However, both had quite a drastic reduction in eggs when males were present.

We look at the effect sizes:

```
timareta_eggs_eff_size <- summary(emmeans(timareta_eggs_glmm,
    pairwise ~ Male_presence | wing_colour,
    transform = "response")$contrasts,
    infer = T)
# Inverse the numbers (as emmeans
# calculate without males - with
# males, but we want it the other
# way around)
timareta_eggs_eff_size[, c("estimate",
    "asymp.LCL", "asymp.UCL")] <- timareta_eggs_eff_size[,
    c("estimate", "asymp.LCL", "asymp.UCL")] *
    (-1)
# Aesthetical changes of the table
names(timareta_eggs_eff_size)[6:7] <- names(timareta_eggs_eff_size)[7:6]
timareta_eggs_eff_size <- timareta_eggs_eff_size[,
    c(1:5, 7, 6, 8:9)]
timareta_eggs_eff_size$contrast <- as.character("present - not_present")
timareta_eggs_eff_size[, -(1:2)] <- round(timareta_eggs_eff_size[,
    -(1:2)], 3)
# Print
print(timareta_eggs_eff_size)
```

```
##                contrast wing_colour estimate    SE  df asymp.LCL asymp.UCL
## 1 present - not_present      yellow   -1.744 0.674 Inf    -3.065    -0.422
## 2 present - not_present  yellow_red   -1.660 0.712 Inf    -3.055    -0.264
##   z.ratio p.value
## 1   2.586    0.01
## 2   2.331    0.02
```

The reduction in eggs can be seen equally in both groups!

Let's see which of the factors of the model are significant:

```
joint_tests(timareta_eggs_glmm)
```

Again, male presence seems to be the only factor really playing a role!

We now test for the effect of male presence in isolation:

```
timareta_eggs_glmm_males <- glmer(eggs ~
    Male_presence + (1 | female), family = "poisson",
    data = timareta_data_eggs_glmm)
```

We look at the estimators

```
(timareta_estimates <- emmeans(timareta_eggs_glmm_males,
    ~Male_presence, transform = "response"))
```

```
##  Male_presence rate    SE  df asymp.LCL asymp.UCL
##  not_present   8.86 0.589 Inf      7.70     10.01
##  present       7.15 0.496 Inf      6.18      8.13
##
## Confidence level used: 0.95
```

Again, we look at the effect sizes:

```
timareta_eggs_eff_size_PLOT <- summary(emmeans(timareta_eggs_glmm_males,
    pairwise ~ Male_presence, transform = "response")$contrasts,
    infer = T)
# Inverse again the numbers
timareta_eggs_eff_size_PLOT[, c("estimate",
    "asymp.LCL", "asymp.UCL")] <- timareta_eggs_eff_size_PLOT[,
    c("estimate", "asymp.LCL", "asymp.UCL")] *
    (-1)
# Aesthetical changes of the table
names(timareta_eggs_eff_size_PLOT)[5:6] <- names(timareta_eggs_eff_size_PLOT)[6:5]
timareta_eggs_eff_size_PLOT <- timareta_eggs_eff_size_PLOT[,
    c(1:5, 6, 5, 7:8)]
timareta_eggs_eff_size_PLOT$contrast <- as.character("present - not_present")
timareta_eggs_eff_size_PLOT[, -(1:2)] <- round(timareta_eggs_eff_size_PLOT[,
    -(1:2)], 3)
# Print
print(timareta_eggs_eff_size_PLOT)
```

```
##               contrast  estimate   SE  df asymp.UCL asymp.LCL asymp.UCL.1
## 1 present - not_present -1.704124 0.49 Inf    -0.744    -2.664      -0.744
##   z.ratio p.value
## 1    3.48   0.001
```

Testing for the overall effect of male presence will give the same p-value as the one printed above:

```
(test_for_panelB_eggs <- joint_tests(timareta_eggs_glmm_males))
```

Again, we fit the same type of models as above, but considering size of the yellow patch on the wing (relative to the whole wing size, either including all individuals or only females with red on the wings), and/or size of the red patch (relative to the whole wing size, and only including females with red) as dependant variables. The reasoning behind this is the same as for the analyses of male harassment. * Model 1: testing for proportion of yellow band across all females * Model 2: testing for proportion of yellow band and proportion of red band as separate factor, only including females with red on the wing

```
timareta_eggs_glmm_yellow <- glmer(eggs ~
    Male_presence * yellow_rel + (1 |
        female), family = "poisson",
    data = timareta_data_eggs_glmm)
timareta_data_eggs_glmm_red <- timareta_data_eggs_glmm[timareta_data_eggs_glmm$wing_colour ==
    "yellow_red", ]
timareta_eggs_glmm_red_and_yellow <- glmer(eggs ~
    Male_presence * red_rel + Male_presence *
        yellow_rel + (1 | female), family = "poisson",
    data = timareta_data_eggs_glmm)
```

We test for significance of the slopes. Note, the slopes we check here are the effect size slopes, *i.e.* the difference between present minus not present. Due to alphabetical order, `emtrends` calculates the effect the other way around (not present minus present), but we don't bother for now to bring this into the right order, as there seems to be no effect whatsoever anyway:

```r
eggs_trend_yellow <- emtrends(timareta_eggs_glmm_yellow,
    pairwise ~ Male_presence * yellow_rel,
    var = "yellow_rel")$contrasts
print("Slope proportion yellow")
```

```
## [1] "Slope proportion yellow"
```

```r
(eggs_p_yellow <- summary(eggs_trend_yellow,
    infer = c(TRUE, TRUE), null = 0,
    side = "two-sided"))
```

```r
eggs_trend_red <- emtrends(timareta_eggs_glmm_red_and_yellow,
    pairwise ~ Male_presence * red_rel,
    var = "red_rel")$contrasts
print("Slope proportion red (only individuals with red)")
```

```
## [1] "Slope proportion red (only individuals with red)"
```

```r
(eggs_p_red <- summary(eggs_trend_red,
    infer = c(TRUE, TRUE), null = 0,
    side = "two-sided"))
```

```r
eggs_trend_yellow_red_indiv <- emtrends(timareta_eggs_glmm_red_and_yellow,
    pairwise ~ Male_presence * yellow_rel,
    var = "yellow_rel")$contrasts
print("Slope proportion yellow (only individuals with red)")
```

```
## [1] "Slope proportion yellow (only individuals with red)"
```

```r
(eggs_p_yellow_red_indiv <- summary(eggs_trend_yellow_red_indiv,
    infer = c(TRUE, TRUE), null = 0,
    side = "two-sided"))
```

This looks like a quite complex output, but it's in fact the exact same p-values we get from calling `summary` and looking at the interaction terms.

We create ourselves a brief summary of the data, first with the actual raw numbers:

```r
wo_males <- timareta_data_eggs$Eggs_without_males
w_males <- timareta_data_eggs$Eggs_with_males
paste0(t1, round(mean(w_males), 2))
```

```
## [1] "Mean with male = 7.73"
```

```r
paste0(t2, round(mean(wo_males), 2))
```

```
## [1] "Mean without male = 9.57"
```

```r
paste0(t3, round(mean(timareta_data_eggs$egg_difference),
    2))
```

```
## [1] "Mean difference = -1.84"
```

```r
paste0(t4, round(sum(wo_males)/sum(w_males) -
    1, 2) * 100, "%")
```

```
## [1] "Without male x % more than with male = 24%"
```

```r
paste0(t5, round(sum(w_males)/sum(wo_males) -
    1, 2) * 100, "%")
```

```
## [1] "With male x % less than without male = -19%"
```

We repeat the same with model estimates:

```
wo_males <- summary(timareta_estimates)$rate[summary(timareta_estimates)$Male_presence ==
    "not_present"]
w_males <- summary(timareta_estimates)$rate[summary(timareta_estimates)$Male_presence ==
    "present"]
paste0(t1, round(w_males, 2))
```

```
## [1] "Mean with male = 7.15"
```

```
paste0(t2, round(wo_males, 2))
```

```
## [1] "Mean without male = 8.86"
```

```
paste0(t3, round(timareta_eggs_eff_size_PLOT$estimate,
    2))
```

```
## [1] "Mean difference = -1.7"
```

```
paste0(t4, round(wo_males/w_males -
    1, 2) * 100, "%")
```

```
## [1] "Without male x % more than with male = 24%"
```

```
paste0(t5, round(w_males/wo_males -
    1, 2) * 100, "%")
```

```
## [1] "With male x % less than without male = -19%"
```

#### 4.6 Analyses of caterpillars hatched in experiment 2

In the following, we check for the correlation of eggs laid and caterpillars hatched, which was measured for a subset of females from experiment 2.

First, for easing later coding, we create 4 new vectors. For each "treatment" group (yellow and yellow-red females), we create a variable with the the eggs laid with and without male present, as well as the number of caterpillars hatched from these eggs.

```
eggs_laid_exp <- timareta_data$Eggs_with_males
eggs_laid_con <- timareta_data$Eggs_without_males
hatched_exp <- ifelse(timareta_data$Males_in_A_or_B ==
    "A", timareta_data$Day_3_4_hatch_A,
    timareta_data$Day_5_6_hatch_B)
hatched_con <- ifelse(timareta_data$Males_in_A_or_B ==
    "A", timareta_data$Day_5_6_hatch_B,
    timareta_data$Day_3_4_hatch_A)
```

There is one outlier datapoint, where a female laid much more than all others in a two-day interval. This datapoint lies also outside the 1.5*IQR range (IQR = interquartile range, left boxplot). It is also an extreme outlier for number of hatchlings (it comes from the same female; right boxplot)!

```
layout(matrix(1:2, ncol = 2))
par(mar = c(1, 4, 0.2, 0.5))
boxplot(c(eggs_laid_exp[!is.na(eggs_laid_exp)],
    eggs_laid_con[!is.na(eggs_laid_con)]),
    ylab = "Number of eggs laid")
boxplot(c(hatched_exp[!is.na(hatched_exp)],
```

```
    hatched_con[!is.na(hatched_con)]),
    ylab = "Number of caterpillars eclosed")
```

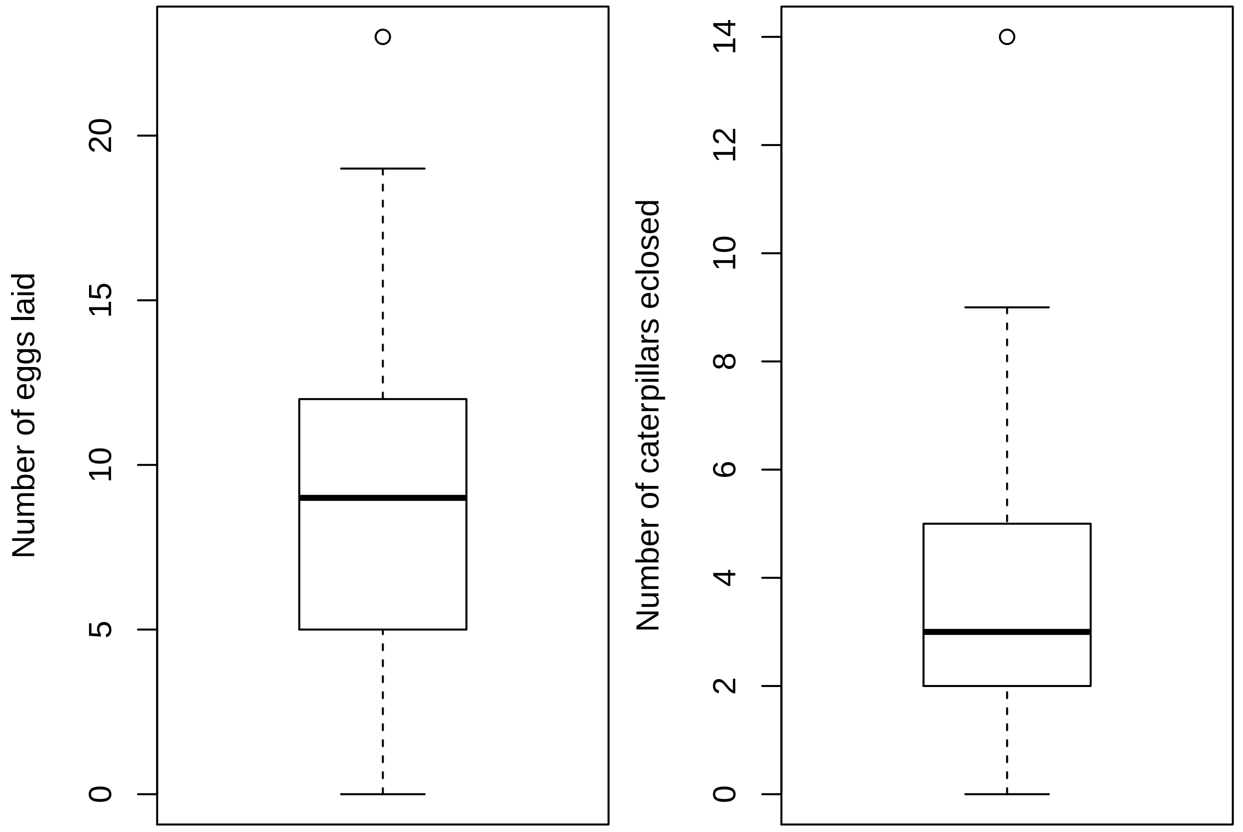


We exclude this data point, as it's otherwise making it harder to see the trends in the graph.

```
outlier_exp <- ifelse(is.na(eggs_laid_exp),
    T, eggs_laid_exp > 20)
outlier_con <- ifelse(is.na(eggs_laid_con),
    T, eggs_laid_con > 20)
eggs_laid_exp[outlier_exp] <- NA
eggs_laid_con[outlier_con] <- NA
hatched_exp[outlier_exp] <- NA
hatched_con[outlier_con] <- NA
```

We run a couple of `loess` regressions, in order to visualize the relationships. our prediction range goes from minimum to maximum number of eggs laid (when no informtation is available in the model, the prediction will simply be `NA`).

```
pred_range <- seq(min(c(eggs_laid_con,
    eggs_laid_exp), na.rm = T), max(c(eggs_laid_con,
    eggs_laid_exp), na.rm = T), length.out = 100)
# loess for all data (independent if
# male present or not), and
# predictions along range
lo <- loess(c(hatched_con, hatched_exp) ~
    c(eggs_laid_con, eggs_laid_exp),
    span = 0.9)
```

```
pred <- predict(lo, pred_range)
# loess for data when males were
# present
lom <- loess(hatched_exp ~ eggs_laid_exp,
    span = 0.9)
predm <- predict(lom, pred_range)
# loess for data when males were
# absent
lonm <- loess(hatched_con ~ eggs_laid_con,
    span = 0.9)
prednm <- predict(lonm, pred_range)
```

We plot this:

```
png(paste0("Figures/empirical_eggs_cats.png"),
    width = 1500, height = 1500, res = 300)
par(mar = c(2.2, 2, 0.2, 0.2))

# Open empty plot
plot(1, 1, xlim = range(c(eggs_laid_con,
    eggs_laid_exp), na.rm = T), ylim = range(c(hatched_con,
    hatched_exp), na.rm = T), pch = 21,
    type = "n", xlab = "", ylab = "",
    xaxt = "n", yaxt = "n")
# set seed for jittering
set.seed(42)
# Add points with slight jitter in x
# and y. Add triangles for eggs laid
# with male and circles for eggs
# laid without
points(jitter(c(eggs_laid_con, eggs_laid_exp),
    amount = 0.015 * diff(par("usr")[1:2])),
    jitter(c(hatched_con, hatched_exp),
        amount = 0.015 * diff(par("usr")[3:4])),
    pch = c(rep(21, length(eggs_laid_con)),
        rep(24, length(eggs_laid_exp))),
    bg = adjustcolor(rep(ifelse(timareta_data$wing_colour ==
        "yellow", "yellow", "red")),
        0.5), cex = 0.9)
# Add predictions for females with
# male
lines(pred_range, predm, col = "dodgerblue")
# Add predictions for females
# without male
lines(pred_range, prednm, col = "green")
# Add predictions for all females
lines(pred_range, pred)

# Axes and titles
axis(1, line = -0.9, lwd = 0, cex.axis = 0.8)
axis(1, labels = F, tck = -0.01)
axis(2, line = -0.7, lwd = 0, las = 1,
    cex.axis = 0.8)
axis(2, labels = F, tck = -0.01)
```

```
mtext("Eggs laid", 1, line = 1.1, cex = 0.9)
mtext("Caterpillars hatched", 2, line = 1.1,
    cex = 0.9)

invisible(dev.off())
```

Show png we just made:

```
fig_disp <- readPNG("Figures/empirical_eggs_cats.png")
grid.raster(fig_disp)
```

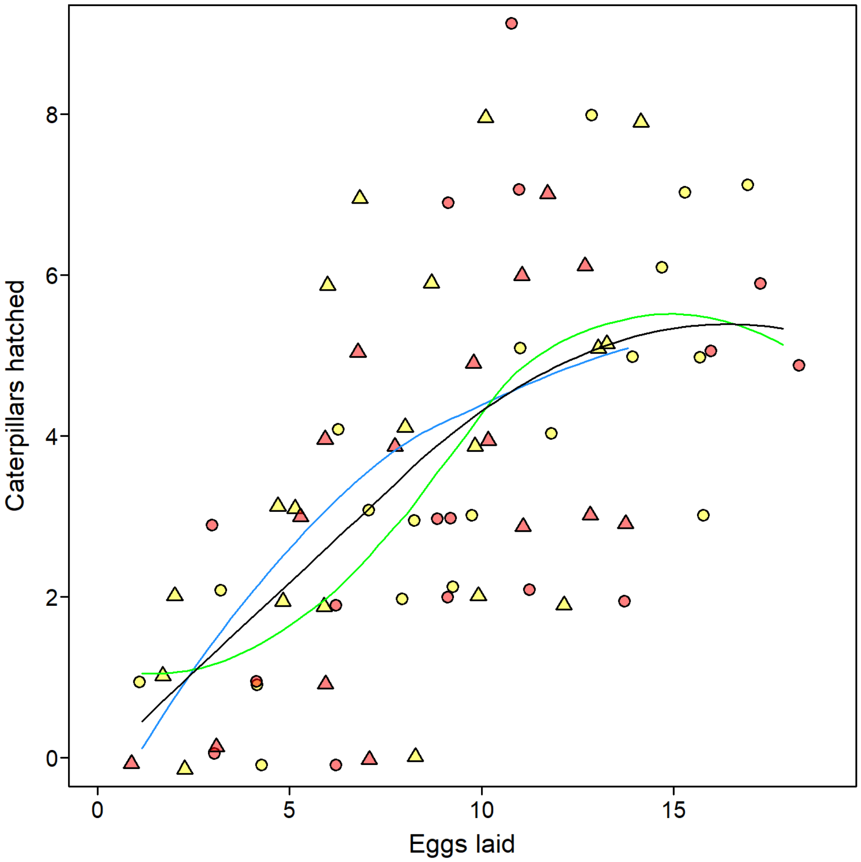


##### 4.7 Combined plot for male harassment and female fecundity

Read in photos from different butterfly pictures. Note: These will **NOT** be provided in the online supplement. If you rerun this, comment out the code where these variables appear.

```
erato_red <- readPNG("wing_photos/erato_red.png")
erato_black <- readPNG("wing_photos/erato_black.png")
BC_yell <- readPNG("wing_photos/BC_no_red.png")
BC_yellred <- readPNG("wing_photos/BC_red.png")
```

Plot harassment results including outliers.

```
png("Figures/empirical_male_harassment_with_outliers.png",width=3765,height=3600,res=600)
layout(matrix(1:2,nrow=2))
par(oma=c(0,0.6,0,0))

#Experiment 1. Use plot_proportion_stats code.
```

```
#The extreme outlier
all_extr<-outliers_by_group_erato[[2]]
extr<-all_extr[all_extr==max(all_extr)]

outputto<-invisible(plot_proportion_stats(
  input_data=erato_data_harass_glmm[erato_data_harass_glmm$female!=names(extr),],
                                         # The data set
                                         sum_by_col_plot=c("female"),
                                         # Sum by ID for plot
                                         response_col=list(c("harass_sec"),
                                                           c("no_harass_sec")),
                                         # Response variable
                                         sub1_col="treat2",
                                         # Fixed effect in model
                                         sub1_states=c("treatment","control"),
                                         # Panels from left to right
                                         stats_for_1sub=F,
                                         # Don't show automatically created GLMM results
                                         #(although they would be identical)
                                         #Aesthetics (not so important):
                                         par_mar=c(1,3.2,1.5,7.7),
                                         return_plot_table = T,
                                         ylims = "0_to_max",
                                         sub1_labels=c("",""),
                                         show_N=F,
                                         italics_sub1 = F,
                                         yaxs_label="",
                                         y_tck=-0.015,
                                         horiz_0.5=T,
                                         panel_dividers=F,
                                         dot_size=0.0145,
                                         dot_colours=list("black","red"),
                                         transparency = 0,
                                         line_colours = adjustcolor("white",0),
                                         dot_lwd = 1.8,
                                         sub_box_yn=F,
                                         numb_iterations=1000,
                                         legend_dots=c(),
                                         border_space=0.49,
                                         seedi=123,
                                         verbose=F
))
#This is a purely aesthetical aspect. We add the dots calculated from above function,
#but with varying point characters
pch_type<-sapply(outputto[[2]]$female,function(x)
  erato_data_harass_glmm$treat1[erato_data_harass_glmm$female==x][1])
pch_type<-ifelse(pch_type!="only_handled",21,24)
points(outputto[[3]],outputto[[2]]$preference_DUMMY,
       bg=adjustcolor(outputto[[2]]$dot_coloursss,0.75),
       cex=0.0145*sqrt(outputto[[2]]$appearance_DUMMY),
       pch=pch_type)

#Mark outliers
```

```r
points(c(0.5,1.5,1.5,1.5),
       c(0.005,0.0134,0.0281,0.0345),
       pch=1,cex=4.5)
arrows(0.25,0.007,
       0.48,0.0019,
       length=0.03)
text(0.22,0.0083,"Also an outlier",cex=0.6)

#Add info about the one excluded outlier
arrows(0.5,par("usr")[3]+0.91*diff(par("usr")[3:4]),
       0.5,par("usr")[3]+0.96*diff(par("usr")[3:4]),
       length=0.03)
text(0.5,par("usr")[3]+0.89*diff(par("usr")[3:4]),
     paste0("one more at ",
            format(round(extr,3),nsmall=3)),cex=0.6)

#Add sample sizes:
axis(1,at=c(0.5,1.5),
     paste0("(",c(sum(outputto[[2]]$positioning==0.5),
                  sum(outputto[[2]]$positioning==1.5)),")"),
     lwd=0,line=-1,cex.axis=0.8)

#Add estimators
add_CI(EMM_obj = erato_harass_emm,at=1:2,match_col="treat2",
       match_vec = c("treatment","control"))

#Add p-value
p_value_add<-ifelse(test_for_panelA_suppl$p.value<0.001,"< 0.001",
                    paste0("= ",format(round(test_for_panelA_suppl$p.value,3),nsmall=3)))
axis(3,at=c(0.5,1.5),line=0.55,tck=0.02,labels=F)
mtext(bquote(italic(p)~.(p_value_add)),3,line=0.5,cex=0.73)

#We add a letter to the figure for this panel
text(par("usr")[1]+(0.3/par("pin")[1])*diff(par("usr")[1:2]),
     par("usr")[4]-(0.23/par("pin")[2])*diff(par("usr")[3:4]),
     "A",font=2,cex=1.1,
     pos=2,offset=0)

#Legend
par(xpd=NA)
#Images
addImg(erato_black,x=2.64,y=par("usr")[3]+0.6*diff(par("usr")[3:4]),
       width = 0.18)
addImg(erato_red,x=2.64,y=par("usr")[3]+0.4*diff(par("usr")[3:4]),
       width = 0.18)

#Lines indicating where each picture belongs to
segments(2.525,par("usr")[3]+0.535*diff(par("usr")[3:4]),
         2.525,par("usr")[3]+0.265*diff(par("usr")[3:4]),
         col="black",lwd=2.2)
segments(2.525,par("usr")[3]+0.635*diff(par("usr")[3:4]),
         2.525,par("usr")[3]+0.565*diff(par("usr")[3:4]),lwd=2.2)
segments(2.525,par("usr")[3]+0.535*diff(par("usr")[3:4]),
```

```
         2.525,par("usr")[3]+0.265*diff(par("usr")[3:4]),
         col="red",lwd=1.5)
#Further legend info
rect(2.05,par("usr")[3]+0.25*diff(par("usr")[3:4]),
     2.75,par("usr")[3]+0.75*diff(par("usr")[3:4]),
     col=NA,border="black",lwd=0.5)
text(2.4,par("usr")[3]+0.7*diff(par("usr")[3:4]),
     font=2,cex=0.8,"Experiment 1")
text(2.18,par("usr")[3]+0.60*diff(par("usr")[3:4]),
     cex=0.6,"manipulated",pos=4,offset=0)
text(2.18,par("usr")[3]+0.516*diff(par("usr")[3:4]),
     cex=0.6,"control",pos=4,offset=0)
text(2.18,par("usr")[3]+0.484*diff(par("usr")[3:4]),
     cex=0.47,"transparent pen",pos=4,offset=0)
text(2.18,par("usr")[3]+0.416*diff(par("usr")[3:4]),
     cex=0.6,"control",pos=4,offset=0)
text(2.18,par("usr")[3]+0.384*diff(par("usr")[3:4]),
     cex=0.47,"only handled",pos=4,offset=0)
text(2.18,par("usr")[3]+0.3*diff(par("usr")[3:4]),
     cex=0.6,"untreated",pos=4,offset=0)
points(rep(2.115,4),
       par("usr")[3]+c(0.6,0.5,0.4,0.3)*diff(par("usr")[3:4]),
       cex=1.3,pch=c(21,21,24,22),bg=adjustcolor(c("black",rep("red",3)),0.75))
par(xpd=F)


#Experiment 2 (same as above)
outputto<-invisible(plot_proportion_stats(input_data=timareta_data_harass_glmm,
                                          # The data set
                                          sum_by_col_plot=c("female"),
                                          # Sum by ID for plot
                                          response_col=list(c("harass_sec"),
                                                            c("no_harass_sec")),
                                          # Response variable
                                          sub1_col="wing_colour",
                                          # Fixed effect in model
                                          sub1_states=c("yellow_red","yellow"),
                                          # Panels from left to right
                                          stats_for_1sub=F,
                                          # Don't show automatically created GLMM results
                                          # (although they would be identical)
                                          #Aesthetics (not so important):
                                          par_mar=c(1,3.2,1.5,7.7),
                                          return_plot_table = T,
                                          ylims = "0_to_max",
                                          sub1_labels=c("",""),
                                          show_N=F,
                                          italics_sub1 = F,
                                          yaxs_label="",
                                          y_tck=-0.015,
                                          horiz_0.5=T,
                                          panel_dividers=F,
                                          dot_size=0.0145,
```

```
                                              dot_colours=list("red","yellow"),
                                              transparency = 0,
                                              line_colours = adjustcolor("white",0),
                                              dot_lwd = 1.8,
                                              sub_box_yn=F,
                                              numb_iterations=1000,
                                              legend_dots=c(),
                                              border_space=0.49,
                                              seedi=123,
                                              verbose=F
))

#Similar as above, we add the dots separately
points(outputto[[3]],outputto[[2]]$preference_DUMMY,pch=21,
       bg=adjustcolor(outputto[[2]]$dot_coloursss,0.75),
       cex=0.0145*sqrt(outputto[[2]]$appearance_DUMMY))

#Mark outliers
points(c(0.5,1.5),
       c(0.247,0.171),
       pch=1,cex=4.5)

#Add sample sizes:
axis(1,at=c(0.5,1.5),
     paste0("(",c(sum(outputto[[2]]$positioning==0.5),
                  sum(outputto[[2]]$positioning==1.5)),")"),
     lwd=0,line=-1,cex.axis=0.8)


#Add estimators
add_CI(EMM_obj = timareta_harass_emm,at=1:2,match_col="wing_colour",
       match_vec = c("yellow_red","yellow"))

#Add p-value
p_value_add<-ifelse(test_for_panelB_suppl$p.value<0.001,"< 0.001",
                    paste0("= ",format(round(test_for_panelB_suppl$p.value,3),nsmall=3)))
axis(3,at=c(0.5,1.5),line=0.55,tck=0.02,labels=F)
mtext(bquote(italic(p)~.(p_value_add)),3,line=0.5,cex=0.73)

#We add a letter to the figure for this panel
text(par("usr")[1]+(0.3/par("pin")[1])*diff(par("usr")[1:2]),
     par("usr")[4]-(0.23/par("pin")[2])*diff(par("usr")[3:4]),
     "B",font=2,cex=1.1,
     pos=2,offset=0)

#Legend (same as before)
par(xpd=NA)

addImg(BC_yellred,x=2.64,y=par("usr")[3]+0.5375*diff(par("usr")[3:4]),
       width = 0.16)
addImg(BC_yell,x=2.64,y=par("usr")[3]+0.3975*diff(par("usr")[3:4]),
       width = 0.155)
segments(2.525,par("usr")[3]+0.5725*diff(par("usr")[3:4]),
```

```
         2.525,par("usr")[3]+0.5025*diff(par("usr")[3:4]),lwd=2.2,"black")
segments(2.525,par("usr")[3]+0.4325*diff(par("usr")[3:4]),
         2.525,par("usr")[3]+0.3625*diff(par("usr")[3:4]),
         col="black",lwd=2.2)
segments(2.525,par("usr")[3]+0.5725*diff(par("usr")[3:4]),
         2.525,par("usr")[3]+0.5025*diff(par("usr")[3:4]),lwd=1.5,"red")
segments(2.525,par("usr")[3]+0.4325*diff(par("usr")[3:4]),
         2.525,par("usr")[3]+0.3625*diff(par("usr")[3:4]),
         col="yellow",lwd=1.5)
rect(2.05,par("usr")[3]+0.3125*diff(par("usr")[3:4]),
     2.75,par("usr")[3]+0.6875*diff(par("usr")[3:4]),
     col=NA,border="black",lwd=0.5)
text(2.4,par("usr")[3]+0.6375*diff(par("usr")[3:4]),
     font=2,cex=0.8,"Experiment 2")
text(2.18,par("usr")[3]+0.5375*diff(par("usr")[3:4]),
     cex=0.6,"manipulated",pos=4,offset=0)
text(2.18,par("usr")[3]+0.3975*diff(par("usr")[3:4]),
     cex=0.6,"control",pos=4,offset=0)
points(rep(2.115,2),
       par("usr")[3]+c(0.5375,0.3975)*diff(par("usr")[3:4]),
       cex=1.3,pch=21,bg=adjustcolor(c("red","yellow"),0.75))
par(xpd=F)

mtext("Proportion of observation time with male harassment",2,line=-0.4,outer=T)

invisible(dev.off())
```

Show png we just made:

```
fig_disp <- readPNG("Figures/empirical_male_harassment_with_outliers.png")
grid.raster(fig_disp)
```

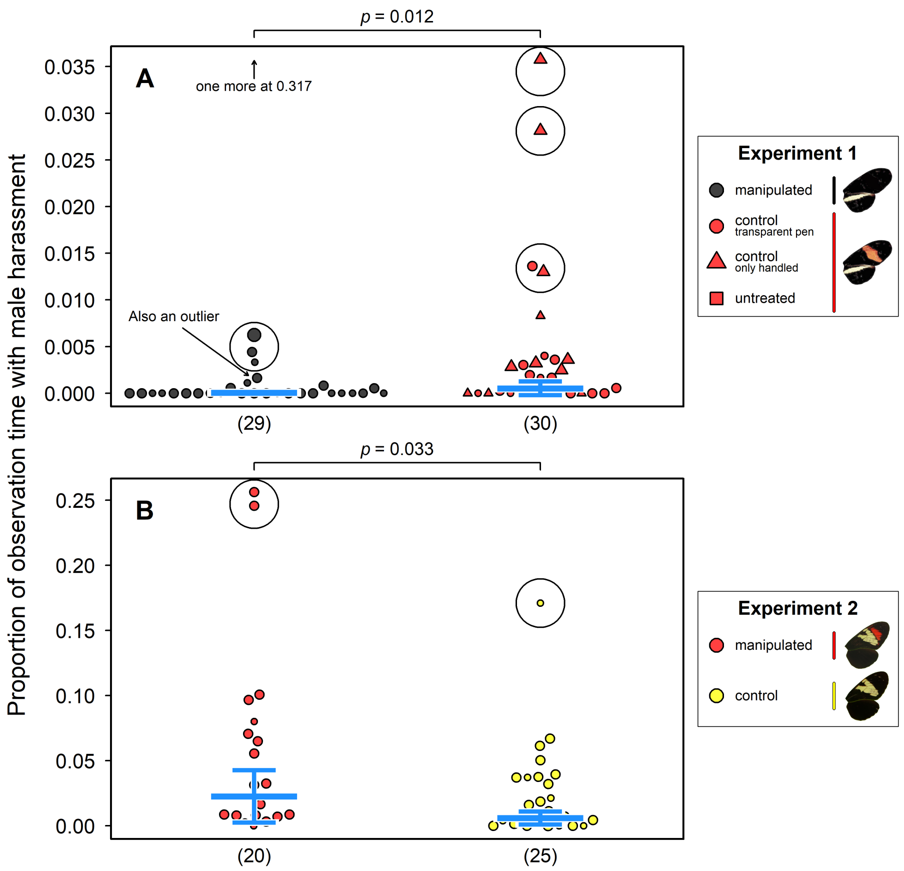


Plot harassment results excluding outliers, as well as fecundity data.

```
png("Figures/empirical_male_harassment_and_eggs_without_outliers.png",width=5800,
    height=3600,res=600)
layout(matrix(c(1:4),nrow=2),widths = c(1.85,1))
par(oma=c(0,0.6,0,0))

#Experiment 1 - harassment (as before)
outputto<-invisible(plot_proportion_stats(input_data=erato_data_harass_glmm_no_outliers,
                                          # The data set
                                          sum_by_col_plot=c("female"),
                                          # Sum by ID for plot
                                          response_col=list(c("harass_sec"),
                                                            c("no_harass_sec")),
                                          # Response variable
                                          sub1_col="treat2",
                                          # Fixed effect in model
                                          sub1_states=c("treatment","control"),
                                          # Panels from left to right
                                          stats_for_1sub=F,
                                          # Don't show automatically created GLMM results
                                          # (although they would be identical)
                                          #Aesthetics (not so important):
                                          par_mar=c(1,3.2,1.5,9.7),
                                          return_plot_table = T,
                                          ylims = "0_to_max",
                                          sub1_labels=c("",""),
```

```
                                                  show_N=F,
                                                  italics_sub1 = F,
                                                  yaxs_label="",
                                                  y_tck=-0.015,
                                                  horiz_0.5=T,
                                                  panel_dividers=F,
                                                  dot_size=0.02,
                                                  dot_colours=list("black","red"),
                                                  transparency = 0,
                                                  line_colours = adjustcolor("white",0),
                                                  dot_lwd = 1.8,
                                                  sub_box_yn=F,
                                                  numb_iterations=1000,
                                                  legend_dots=c(),
                                                  border_space=0.49,
                                                  seedi=123,
                                                  verbose=F
))
#Again, adding the dots separately
pch_type<-sapply(outputto[[2]]$female,function(x)
  erato_data_harass_glmm_no_outliers$treat1[erato_data_harass_glmm_no_outliers$female==x][1])
pch_type<-ifelse(pch_type!="only_handled",21,24)
points(outputto[[3]],outputto[[2]]$preference_DUMMY,
       bg=adjustcolor(outputto[[2]]$dot_coloursss,0.75),
       cex=0.02*sqrt(outputto[[2]]$appearance_DUMMY),
       pch=pch_type)

#Add sample sizes:
axis(1,at=c(0.5,1.5),
     paste0("(",c(sum(outputto[[2]]$positioning==0.5),
                  sum(outputto[[2]]$positioning==1.5)),")"),
     lwd=0,line=-1,cex.axis=0.8)

#Add estimators
add_CI(EMM_obj = erato_harass_emm2,at=1:2,match_col="treat2",
       match_vec = c("treatment","control"))

#Add p-value
p_value_add<-ifelse(test_for_panelA$p.value<0.001,"< 0.001",
                    paste0("= ",format(round(test_for_panelA$p.value,3),nsmall=3)))
axis(3,at=c(0.5,1.5),line=0.55,tck=0.02,labels=F)
mtext(bquote(italic(p)~.(p_value_add)),3,line=0.5,cex=0.73)

#We add a letter to the figure for this panel
text(par("usr")[1]+(0.3/par("pin")[1])*diff(par("usr")[1:2]),
     par("usr")[4]-(0.23/par("pin")[2])*diff(par("usr")[3:4]),
     "A",font=2,cex=1.5,
     pos=2,offset=0)

#Legend (as before)
par(xpd=NA)
addImg(erato_black,x=2.64,y=par("usr")[3]+0.6*diff(par("usr")[3:4]),
       width = 0.18)
```

```
addImg(erato_red,x=2.64,y=par("usr")[3]+0.4*diff(par("usr")[3:4]),
       width = 0.18)
segments(2.525,par("usr")[3]+0.535*diff(par("usr")[3:4]),
         2.525,par("usr")[3]+0.265*diff(par("usr")[3:4]),
         col="black",lwd=2.2)
segments(2.525,par("usr")[3]+0.635*diff(par("usr")[3:4]),
         2.525,par("usr")[3]+0.565*diff(par("usr")[3:4]),lwd=2.2)
segments(2.525,par("usr")[3]+0.535*diff(par("usr")[3:4]),
         2.525,par("usr")[3]+0.265*diff(par("usr")[3:4]),
         col="red",lwd=1.5)
rect(2.05,par("usr")[3]+0.25*diff(par("usr")[3:4]),
     2.75,par("usr")[3]+0.75*diff(par("usr")[3:4]),
     col=NA,border="black",lwd=0.5)
text(2.4,par("usr")[3]+0.7*diff(par("usr")[3:4]),
     font=2,cex=1,"Experiment 1")
text(2.18,par("usr")[3]+0.60*diff(par("usr")[3:4]),
     cex=0.8,"manipulated",pos=4,offset=0)
text(2.18,par("usr")[3]+0.516*diff(par("usr")[3:4]),
     cex=0.8,"control",pos=4,offset=0)
text(2.18,par("usr")[3]+0.484*diff(par("usr")[3:4]),
     cex=0.67,"transparent pen",pos=4,offset=0)
text(2.18,par("usr")[3]+0.416*diff(par("usr")[3:4]),
     cex=0.8,"control",pos=4,offset=0)
text(2.18,par("usr")[3]+0.384*diff(par("usr")[3:4]),
     cex=0.67,"only handled",pos=4,offset=0)
text(2.18,par("usr")[3]+0.3*diff(par("usr")[3:4]),
     cex=0.8,"untreated",pos=4,offset=0)
points(rep(2.115,4),
       par("usr")[3]+c(0.6,0.5,0.4,0.3)*diff(par("usr")[3:4]),
       cex=1.3,pch=c(21,21,24,22),bg=adjustcolor(c("black",rep("red",3)),0.75))
par(xpd=F)

#Experiment 2 (also same as above)
outputto<-invisible(plot_proportion_stats(input_data=timareta_data_harass_glmm_no_outliers,
                                          # The data set
                                          sum_by_col_plot=c("female"),
                                          # Sum by ID for plot
                                          response_col=list(c("harass_sec"),
                                                            c("no_harass_sec")),
                                          # Response variable
                                          sub1_col="wing_colour",
                                          # Fixed effect in model
                                          sub1_states=c("yellow_red","yellow"),
                                          # Panels from left to right
                                          stats_for_1sub=F,
                                          # Don't show automatically created GLMM results
                                          # (although they would be identical)
                                          #Aesthetics (not so important):
                                          par_mar=c(1,3.2,1.5,9.7),
                                          return_plot_table = T,
                                          ylims = "0_to_max",
                                          sub1_labels=c("",""),
                                          show_N=F,
```

```
                                                italics_sub1 = F,
                                                yaxs_label="",
                                                y_tck=-0.015,
                                                horiz_0.5=T,
                                                panel_dividers=F,
                                                dot_size=0.02,
                                                dot_colours=list("red","yellow"),
                                                transparency = 0,
                                                line_colours = adjustcolor("white",0),
                                                dot_lwd = 1.8,
                                                sub_box_yn=F,
                                                numb_iterations=1000,
                                                legend_dots=c(),
                                                border_space=0.49,
                                                seedi=123,
                                                verbose=F
))
#Again, add dots separately
points(outputto[[3]],outputto[[2]]$preference_DUMMY,pch=21,
       bg=adjustcolor(outputto[[2]]$dot_coloursss,0.75),
       cex=0.02*sqrt(outputto[[2]]$appearance_DUMMY))

#Add sample sizes:
axis(1,at=c(0.5,1.5),
     paste0("(",c(sum(outputto[[2]]$positioning==0.5),
                  sum(outputto[[2]]$positioning==1.5)),")"),
     lwd=0,line=-1,cex.axis=0.8)

#Add estimators
add_CI(EMM_obj = timareta_harass_emm2,at=1:2,match_col="wing_colour",
       match_vec = c("yellow_red","yellow"))

#Add p-value
p_value_add<-ifelse(test_for_panelB$p.value<0.001,"< 0.001",
                    paste0("= ",format(round(test_for_panelB$p.value,3),nsmall=3)))
axis(3,at=c(0.5,1.5),line=0.55,tck=0.02,labels=F)
mtext(bquote(italic(p)~.(p_value_add)),3,line=0.5,cex=0.73)

#We add a letter to the figure for this panel
text(par("usr")[1]+(0.3/par("pin")[1])*diff(par("usr")[1:2]),
     par("usr")[4]-(0.23/par("pin")[2])*diff(par("usr")[3:4]),
     "C",font=2,cex=1.5,
     pos=2,offset=0)

#Legend (as before)
par(xpd=NA)
addImg(BC_yellred,x=2.64,y=par("usr")[3]+0.5375*diff(par("usr")[3:4]),
       width = 0.16)
addImg(BC_yell,x=2.64,y=par("usr")[3]+0.3975*diff(par("usr")[3:4]),
       width = 0.155)
segments(2.525,par("usr")[3]+0.5725*diff(par("usr")[3:4]),
         2.525,par("usr")[3]+0.5025*diff(par("usr")[3:4]),lwd=2.2,"black")
segments(2.525,par("usr")[3]+0.4325*diff(par("usr")[3:4]),
```

```r
         2.525,par("usr")[3]+0.3625*diff(par("usr")[3:4]),
         col="black",lwd=2.2)
segments(2.525,par("usr")[3]+0.5725*diff(par("usr")[3:4]),
         2.525,par("usr")[3]+0.5025*diff(par("usr")[3:4]),lwd=1.5,"red")
segments(2.525,par("usr")[3]+0.4325*diff(par("usr")[3:4]),
         2.525,par("usr")[3]+0.3625*diff(par("usr")[3:4]),
         col="yellow",lwd=1.5)
rect(2.05,par("usr")[3]+0.3125*diff(par("usr")[3:4]),
     2.75,par("usr")[3]+0.6875*diff(par("usr")[3:4]),
     col=NA,border="black",lwd=0.5)
text(2.4,par("usr")[3]+0.6375*diff(par("usr")[3:4]),
     font=2,cex=1,"Experiment 2")
text(2.18,par("usr")[3]+0.5375*diff(par("usr")[3:4]),
     cex=0.8,"manipulated",pos=4,offset=0)
text(2.18,par("usr")[3]+0.3975*diff(par("usr")[3:4]),
     cex=0.8,"control",pos=4,offset=0)
points(rep(2.115,2),
       par("usr")[3]+c(0.5375,0.3975)*diff(par("usr")[3:4]),
       cex=1.3,pch=21,bg=adjustcolor(c("red","yellow"),0.75))
par(xpd=F)

mtext("Proportion of observation time with male harassment",2,line=-0.4,outer=T)

#set seed for same randomization of datasets
set.seed(42)

randomize_erato<-sample(1:nrow(erato_data_eggs))
randomize_timareta<-sample(1:nrow(timareta_data_eggs))

par(mar=c(1,3.2,1.5,0.3))

#Empty plot
plot(1,1,type="n",xaxt="n",yaxt="n",xlab="",ylab="",
     xlim=c(0.5,1.35),xaxs="i",
     ylim=c(-1,1)*max(c(abs(erato_data_eggs[!erato_data_eggs$laid0,]$egg_difference),
                                                  abs(timareta_data_eggs$egg_difference))))
#Add a dashed line for the 'null hypothesis'
abline(h=0,lty="dashed")

#Add beeswarms
treat <- erato_data_eggs[!erato_data_eggs$laid0,]$treatment[randomize_erato]
beeswarm(erato_data_eggs[!erato_data_eggs$laid0,]$egg_difference[randomize_erato],
         at=1,add=T,pch=21,cex=1.3,spacing=1.15,
         pwbg=adjustcolor(ifelse(treat=="blacked_out","black","red"),0.75),
         pwpch=ifelse(treat=="only_handled",24,
                      ifelse(treat=="completely_untreated",22,21)))

#Add estimators
add_CI(EMM_obj = erato_eggs_eff_size_PLOT,at=c(1.5),match_col="",response="estimate")

#p-value
pv<-ifelse(test_for_panelA_eggs$p.value<0.001,
           paste0("'\u003C'~","0.001"),
```

```
          paste0("'\u003D'~",
                 format(round(test_for_panelA_eggs$p.value,3),
                        nsmall=3)))
p_value_add<-paste0("bquote(italic(p)~",pv,")")
cmd1<-",cex=0.83)*(diff(par('usr')[3:4])/diff(par('usr')[1:2]))*(par('pin')[1]/par('pin')[2])"
cmd2<-",cex=0.83)*(diff(par('usr')[1:2])/diff(par('usr')[3:4]))*(par('pin')[2]/par('pin')[1])"
panelA_eggs_width<-eval(parse(text=paste0("strwidth(",p_value_add,cmd1)))
panelA_eggs_height<-eval(parse(text=paste0("strheight(",p_value_add,cmd2)))

#Add effect size (with white in the background)
segments(0.675,0,0.675,erato_eggs_eff_size_PLOT$estimate,
         lend=2,lwd=4,col="white")
segments(0.675,0,0.69,0,
         lend=2,lwd=4,col="white")
segments(0.675,erato_eggs_eff_size_PLOT$estimate,
         0.69,erato_eggs_eff_size_PLOT$estimate,
         lend=2,lwd=4,col="white")
segments(0.675,0,0.675,erato_eggs_eff_size_PLOT$estimate,
         lend=2,lwd=3)
segments(0.675,0,0.69,0,
         lend=2,lwd=3)
segments(0.675,erato_eggs_eff_size_PLOT$estimate,
         0.69,erato_eggs_eff_size_PLOT$estimate,
         lend=2,lwd=3)
#Add a little black bracket showing the effect size
segments(0.675,0.5*erato_eggs_eff_size_PLOT$estimate,
         0.625,0.5*erato_eggs_eff_size_PLOT$estimate)
segments(0.625,0.5*erato_eggs_eff_size_PLOT$estimate,
         0.625,0.5*erato_eggs_eff_size_PLOT$estimate-0.09*diff(par("usr")[3:4]))
segments(0.61,0.5*erato_eggs_eff_size_PLOT$estimate-0.09*diff(par("usr")[3:4]),
         0.625,0.5*erato_eggs_eff_size_PLOT$estimate-0.09*diff(par("usr")[3:4]))
#Add p-value
rect(0.61-1.5*panelA_eggs_height,
     0.5*erato_eggs_eff_size_PLOT$estimate-0.12*diff(par("usr")[3:4])-0.55*panelA_eggs_width,
     0.61,
     0.5*erato_eggs_eff_size_PLOT$estimate-0.12*diff(par("usr")[3:4])+0.55*panelA_eggs_width,
     col="white",border="black",lwd=0.3)
cmd1.1<-"text(0.61-0.75*panelA_eggs_height,0.5*erato_eggs_eff_size_PLOT$estimate"
cmd1.2<-"-0.12*diff(par('usr')[3:4]),"
cmd2<-",cex=0.85,srt=90)"
eval(parse(text=paste0(cmd1.1,cmd1.2,p_value_add,cmd2)))

#We add a letter to the figure for this panel
text(par("usr")[1]+(0.3/par("pin")[1])*diff(par("usr")[1:2]),
     par("usr")[4]-(0.23/par("pin")[2])*diff(par("usr")[3:4]),
     "B",font=2,cex=1.5,
     pos=2,offset=0)

#Axes
axis(2,labels=F,tck=-0.015,lwd=1)
axis(2,las=1,cex.axis=0.85,line=-0.45,lwd=0)
axis(1,at=1,
     paste0("(",sum(!erato_data_eggs$laid0),")"),
```

```
     lwd=0,line=-1,cex.axis=0.8)

#Empty plot
plot(1,1,type="n",xaxt="n",yaxt="n",xlab="",ylab="",
     xlim=c(0.5,1.35),xaxs="i",
     ylim=c(-1,1)*max(c(abs(erato_data_eggs[!erato_data_eggs$laid0,]$egg_difference),
                                                  abs(timareta_data_eggs$egg_difference))))
#Null hypothesis
abline(h=0,lty="dashed")

#Data as beeswarm
beeswarm(timareta_data_eggs$egg_difference[randomize_timareta],
         at=1,add=T,pch=21,cex=1.3,spacing=1.15,
         pwbg=adjustcolor(ifelse(timareta_data_eggs$wing_colour[randomize_timareta]=="yellow",
                                 "yellow","red"),0.75))

#Add estimators
add_CI(EMM_obj = timareta_eggs_eff_size_PLOT,at=c(1.5),match_col="",response="estimate")

#p-value
pv<-ifelse(test_for_panelB_eggs$p.value<0.001,
           paste0("'\u003C'~","0.001"),
           paste0("'\u003D'~",
                  format(round(test_for_panelB_eggs$p.value,3),
                         nsmall=3)))
p_value_add<-paste0("bquote(italic(p)~",pv,")")
cmd1<-",cex=0.83)*(diff(par('usr')[3:4])/diff(par('usr')[1:2]))*(par('pin')[1]/par('pin')[2])"
cmd2<-",cex=0.83)*(diff(par('usr')[1:2])/diff(par('usr')[3:4]))*(par('pin')[2]/par('pin')[1])"
panelB_eggs_width<-eval(parse(text=paste0("strwidth(",p_value_add,cmd1)))
panelB_eggs_height<-eval(parse(text=paste0("strheight(",p_value_add,cmd2)))

#Show effect size
segments(0.675,0,0.675,timareta_eggs_eff_size_PLOT$estimate,
         lend=2,lwd=4,col="white")
segments(0.675,0,0.69,0,
         lend=2,lwd=4,col="white")
segments(0.675,timareta_eggs_eff_size_PLOT$estimate,
         0.69,timareta_eggs_eff_size_PLOT$estimate,
         lend=2,lwd=4,col="white")
segments(0.675,0,0.675,timareta_eggs_eff_size_PLOT$estimate,
         lend=2,lwd=3)
segments(0.675,0,0.69,0,
         lend=2,lwd=3)
segments(0.675,timareta_eggs_eff_size_PLOT$estimate,
         0.69,timareta_eggs_eff_size_PLOT$estimate,
         lend=2,lwd=3)
#Add again bracket for effect size
segments(0.675,0.5*timareta_eggs_eff_size_PLOT$estimate,
         0.625,0.5*timareta_eggs_eff_size_PLOT$estimate)
segments(0.625,0.5*timareta_eggs_eff_size_PLOT$estimate,
         0.625,0.5*timareta_eggs_eff_size_PLOT$estimate-0.09*diff(par("usr")[3:4]))
segments(0.61,0.5*timareta_eggs_eff_size_PLOT$estimate-0.09*diff(par("usr")[3:4]),
         0.625,0.5*timareta_eggs_eff_size_PLOT$estimate-0.09*diff(par("usr")[3:4]))
```

```r
#Add p-value
rect(0.61-1.5*panelB_eggs_height,
     0.5*timareta_eggs_eff_size_PLOT$estimate-0.12*diff(par("usr")[3:4])-0.55*panelB_eggs_width,
     0.61,
     0.5*timareta_eggs_eff_size_PLOT$estimate-0.12*diff(par("usr")[3:4])+0.55*panelB_eggs_width,
     col="white",border="black",lwd=0.3)
cmd1.1<-"text(0.61-0.75*panelB_eggs_height,0.5*timareta_eggs_eff_size_PLOT$estimate"
cmd1.2<-"-0.12*diff(par('usr')[3:4]),"
cmd2<-",cex=0.85,srt=90)"
eval(parse(text=paste0(cmd1.1,cmd1.2,p_value_add,cmd2)))

#We add a letter to the figure for this panel
text(par("usr")[1]+(0.3/par("pin")[1])*diff(par("usr")[1:2]),
     par("usr")[4]-(0.23/par("pin")[2])*diff(par("usr")[3:4]),
     "D",font=2,cex=1.5,
     pos=2,offset=0)

#Axes
axis(2,labels=F,tck=-0.015,lwd=1)
axis(2,las=1,cex.axis=0.85,line=-0.45,lwd=0)
axis(1,at=1,
     paste0("(",nrow(timareta_data_eggs),")"),
     lwd=0,line=-1,cex.axis=0.8)

mtext("Eggs with male presence - without male presence",2,line=-38.7,outer=T)

invisible(dev.off())
```

Show png we just made:

```r
fig_disp <- readPNG("Figures/empirical_male_harassment_and_eggs_without_outliers.png")
grid.raster(fig_disp)
```

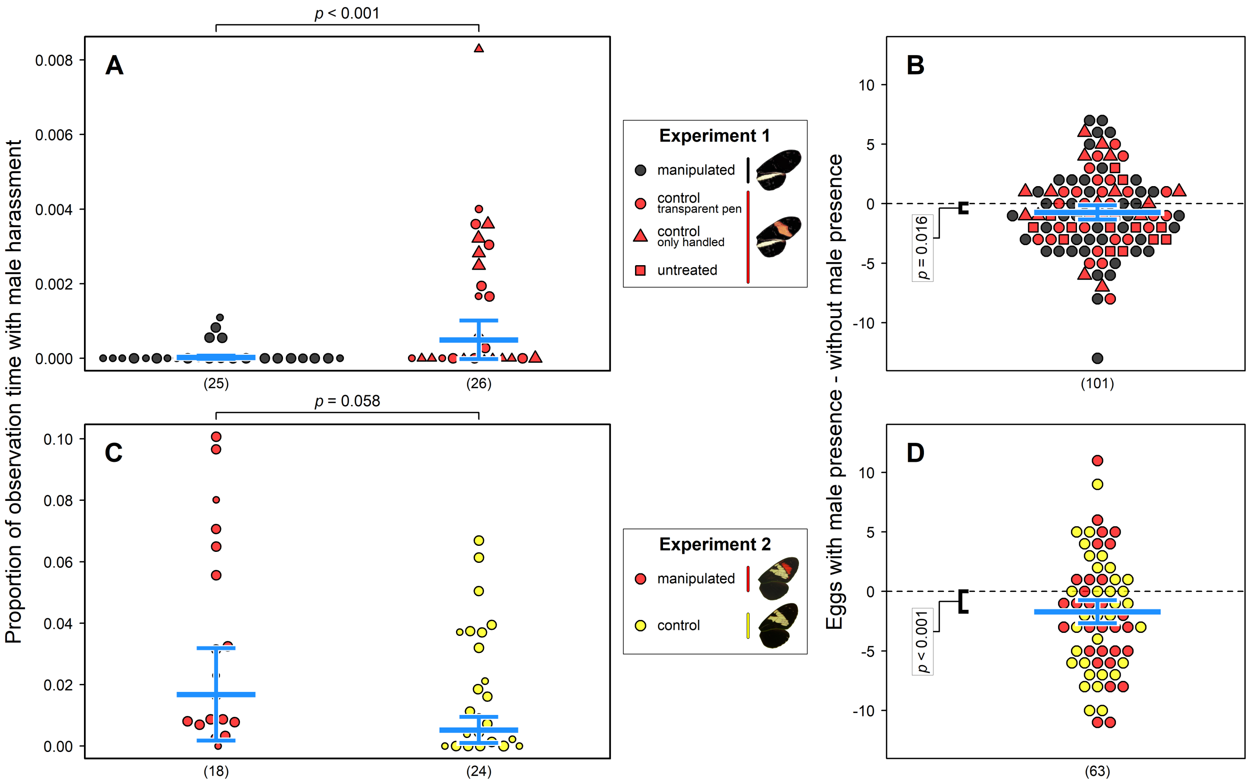


#### 4.8 Testing for effect of harassment on female fecundity

We can now attempt to combine these two analyses and check if females that were harassed more during our observation periods were also more affected in their fecundity by male presence. This analyses is very preliminary and has some caveats, mainly the fact that we observed male behaviours usually only for about 1 hour of the 48 hours of male presence (~24 of which had daylight). Hence, it is unlikely that our resolution is high enough to uncover such trends. Also, harassment data has to be summarized by individual and we didn't find a solid way to weight by observation time in our linear models. Lastly, one further confounding factor is that females might be harassed more if they are especially active during a day (as a moving female will be noticed more by males). However, when they're more active, they might also lay more eggs. We might therefore see an opposite trend, where laying rates with high harassment by males are actually high. Overall, this is analysis can only give a rough idea.

For the *H. erato* data, we create a new dataset and remove previously identified outliers, as well completely untreated females. Also, we add a column showing difference in eggs and one with the 'harassment proportion'

```
erato_eggs_harass <- erato_data[(!erato_data$female %in%
    c(names(outliers_by_group_erato[[1]]),
        names(outliers_by_group_erato[[2]]))) &
    erato_data$treatment != "completely_untreated",
    ]
erato_eggs_harass$eggs_diff <- erato_eggs_harass$Eggs_with_males -
    erato_eggs_harass$Eggs_without_males
eh <- erato_eggs_harass$total_court_time/erato_eggs_harass$total_observation_time
erato_eggs_harass$prop_harass <- eh
```

The difference in eggs laid is roughly normally distributed

```
layout(1)
par(mar = c(4.1, 4, 0.5, 0.5))
par(oma = c(0, 0, 0, 0))
hist(erato_eggs_harass$eggs_diff, main = "",
    xlab = "Eggs with male - eggs without male")
```

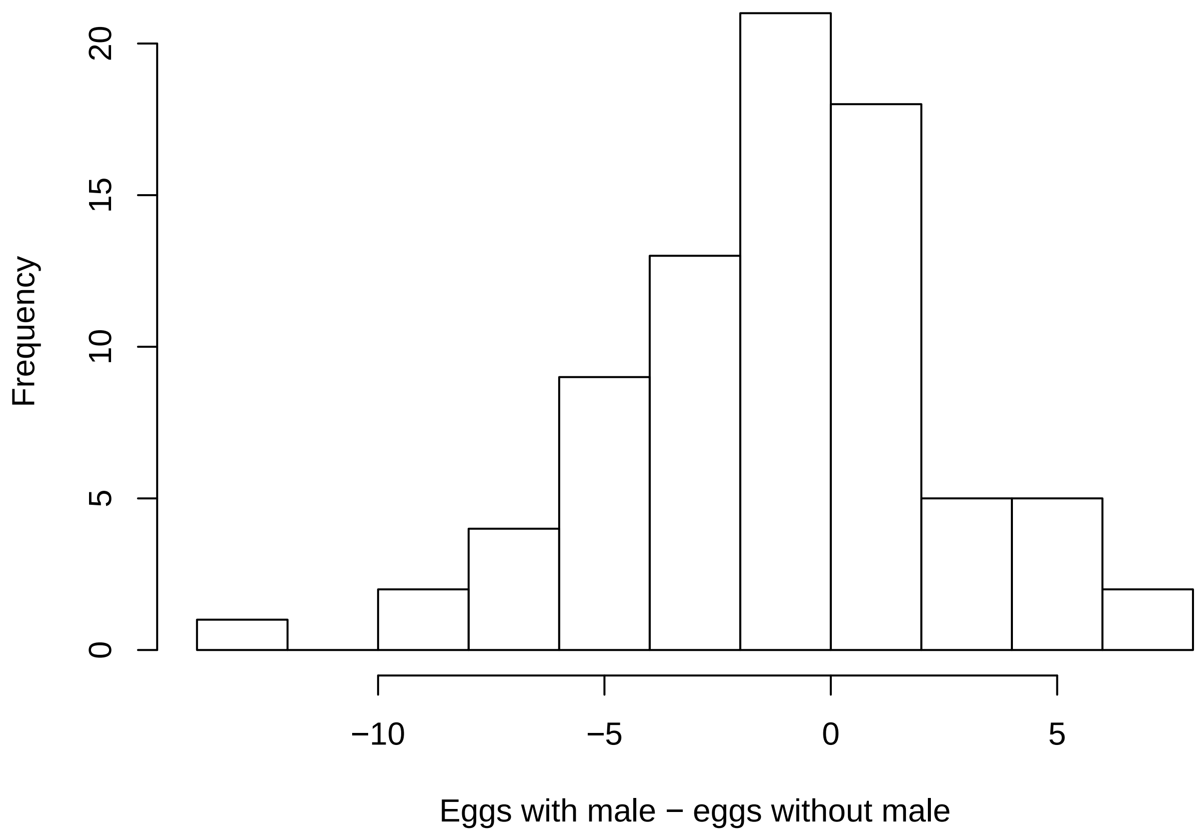


We fit a linear model explaining difference in eggs laid by total harassment proportion, in interaction with female treatment group.

```
erato_eggs_harass_glmm <- lm(eggs_diff ~
    prop_harass * contr_exp, data = erato_eggs_harass)
```

We test which of the slopes is significantly different from 0. First for the two treatment groups separately, then combined. No slope differs significantly from 0.

```
eggs_harass_trend_bygr_erato <- emtrends(erato_eggs_harass_glmm,
    ~prop_harass * contr_exp, var = "prop_harass")
eggs_harass_trend_all_erato <- emtrends(erato_eggs_harass_glmm,
    ~prop_harass, var = "prop_harass")
```

```
## NOTE: Results may be misleading due to involvement in interactions
```

```
summary(eggs_harass_trend_bygr_erato,
    infer = c(TRUE, TRUE), null = 0)
```

```
summary(eggs_harass_trend_all_erato,
    infer = c(TRUE, TRUE), null = 0)
```

We extract model estimators for the control and for the treatment group

```
eggs_harass_pred_erato_contr <- emmip(erato_eggs_harass_glmm,
    ~prop_harass, CIs = T, plotit = F,
    at = list(contr_exp = "control",
        prop_harass = seq(min(erato_eggs_harass$prop_harass[erato_eggs_harass$contr_exp ==
            "control"], na.rm = T),
            max(erato_eggs_harass$prop_harass[erato_eggs_harass$contr_exp ==
                "control"], na.rm = T),
            length.out = 100)))
```

```
## NOTE: Results may be misleading due to involvement in interactions
```

```
eggs_harass_pred_erato_treat <- emmip(erato_eggs_harass_glmm,
    ~prop_harass, CIs = T, plotit = F,
    at = list(contr_exp = "treatment",
        prop_harass = seq(min(erato_eggs_harass$prop_harass[erato_eggs_harass$contr_exp ==
            "treatment"], na.rm = T),
            max(erato_eggs_harass$prop_harass[erato_eggs_harass$contr_exp ==
                "treatment"], na.rm = T),
            length.out = 100)))
```

```
## NOTE: Results may be misleading due to involvement in interactions
```

We produce a new table for the *H. timareta* data, using almost the same logic.

```
timareta_eggs_harass <- timareta_data[!timareta_data$female %in%
    c(names(outliers_by_group_timareta[[1]]),
        names(outliers_by_group_timareta[[2]])),
    ]
timareta_eggs_harass$eggs_diff <- timareta_eggs_harass$Eggs_with_males -
    timareta_eggs_harass$Eggs_without_males
th <- timareta_eggs_harass$total_court_time/timareta_eggs_harass$total_observation_time
timareta_eggs_harass$prop_harass <- th
```

The difference in eggs laid is again roughly normally distributed

```
layout(1)
par(mar = c(4.1, 4, 0.5, 0.5))
par(oma = c(0, 0, 0, 0))
hist(timareta_eggs_harass$eggs_diff,
    main = "", xlab = "Eggs with male - eggs without male")
```

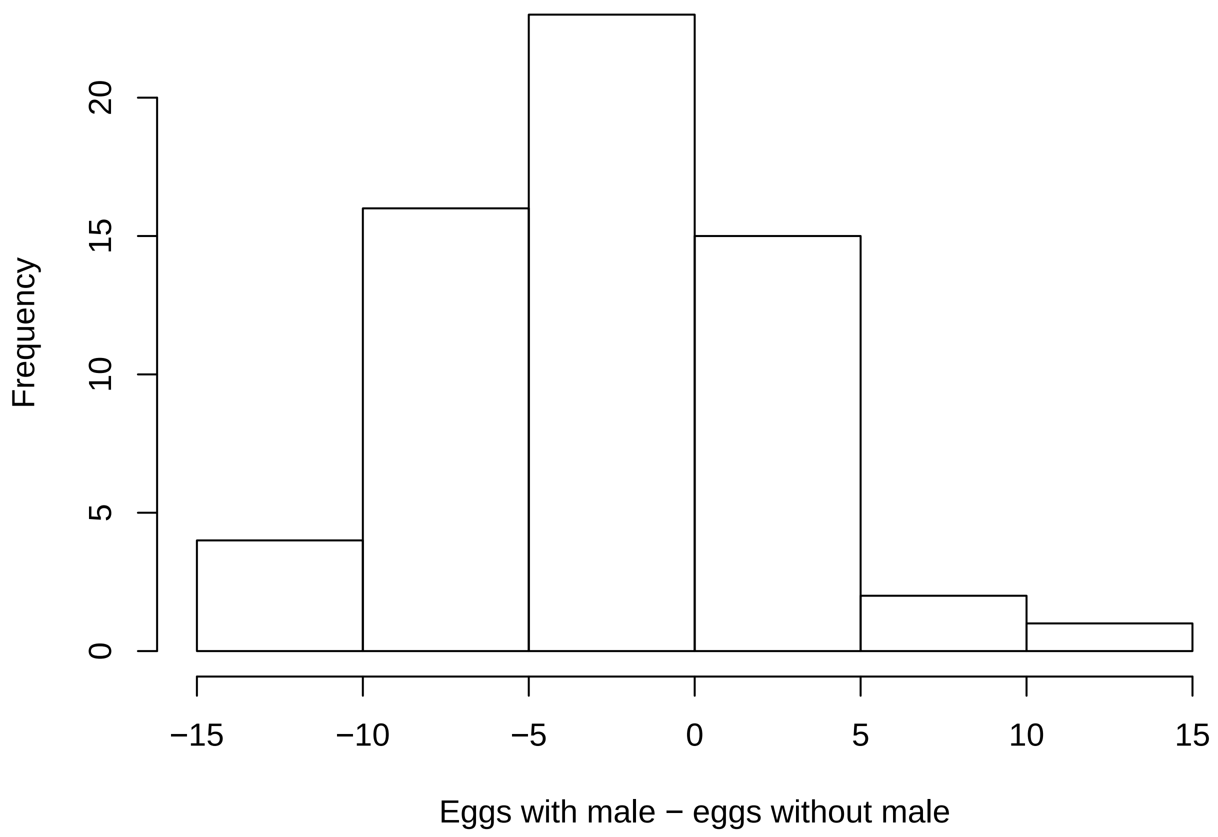


We fit again a linear model

```
timareta_eggs_harass_glmm <- lm(eggs_diff ~
    prop_harass * wing_colour, data = timareta_eggs_harass)
```

We test again which of the slopes is significant. Again, none is signficantly different from 0.

```
eggs_harass_trend_bygr_timareta <- emtrends(timareta_eggs_harass_glmm,
    ~prop_harass * wing_colour, var = "prop_harass")
eggs_harass_trend_all_timareta <- emtrends(timareta_eggs_harass_glmm,
    ~prop_harass, var = "prop_harass")
```

```
## NOTE: Results may be misleading due to involvement in interactions
```

```
summary(eggs_harass_trend_bygr_timareta,
    infer = c(TRUE, TRUE), null = 0)
```

```
summary(eggs_harass_trend_all_timareta,
    infer = c(TRUE, TRUE), null = 0)
```

We extract model estimators for the control and for the treatment group

```
min1 <- min(timareta_eggs_harass$prop_harass[timareta_eggs_harass$wing_colour ==
    "yellow"], na.rm = T)
max1 <- max(timareta_eggs_harass$prop_harass[timareta_eggs_harass$wing_colour ==
    "yellow"], na.rm = T)
min2 <- min(timareta_eggs_harass$prop_harass[timareta_eggs_harass$wing_colour ==
    "yellow_red"], na.rm = T)
max2 <- max(timareta_eggs_harass$prop_harass[timareta_eggs_harass$wing_colour ==
```

```
    "yellow_red"], na.rm = T)
eggs_harass_pred_timareta_contr <- emmip(timareta_eggs_harass_glmm,
    ~prop_harass, CIs = T, plotit = F,
    at = list(wing_colour = "yellow",
        prop_harass = seq(min1, max1,
            length.out = 100)))
```

```
## NOTE: Results may be misleading due to involvement in interactions
```

```
eggs_harass_pred_timareta_treat <- emmip(timareta_eggs_harass_glmm,
    ~prop_harass, CIs = T, plotit = F,
    at = list(contr_exp = "treatment",
        prop_harass = seq(min2, max2,
            length.out = 100)))
```

```
## NOTE: Results may be misleading due to involvement in interactions
```

We plot the slopes for the two different treatment groups out, separated by experiment. We don't include a slope for the overall trend, as the error margin is quite huge in those.

```
png("Figures/empirical_proportion_vs_egg_diff.png",
    width = 1500, height = 2500, res = 300)

layout(matrix(1:2, ncol = 1))
par(mar = c(1.7, 2.5, 1.5, 0.2))
par(oma = c(1.3, 1, 0, 0))

# Set seed to make jitter
# reproducible
set.seed(42)

# Add points, with a slight jitter
# in both directions to avoid
# overlap.
plot(jitter(erato_eggs_harass$prop_harass,
    amount = diff(range(erato_eggs_harass$prop_harass,
        na.rm = T))/50), jitter(erato_eggs_harass$eggs_diff,
    amount = diff(range(erato_eggs_harass$eggs_diff,
        na.rm = T))/50), pch = ifelse(erato_eggs_harass$treatment ==
    "only_handled", 24, 21), bg = adjustcolor(ifelse(erato_eggs_harass$contr_exp ==
    "treatment", "black", "red"), 0.75),
    cex = 0.02 * sqrt(erato_eggs_harass$total_observation_time),
    ylab = "", xlab = "", xaxt = "n",
    yaxt = "n", ylim = range(c(erato_eggs_harass$eggs_diff,
        eggs_harass_pred_erato_contr[,
            c("LCL", "UCL")], eggs_harass_pred_erato_treat[,
            c("LCL", "UCL")]), na.rm = T))
# Add axes
axis(1, lwd = 0, line = -0.5)
axis(1, labels = F, tck = -0.02)
axis(2, las = 2, lwd = 0, line = -0.5)
axis(2, labels = F, tck = -0.02)
mtext(substitute(italic("H. erato demophoon")),
    3, line = 0.2)
```

```r
# Add model estimates with
# confidence intervals
polygon(c(eggs_harass_pred_erato_contr$prop_harass,
    rev(eggs_harass_pred_erato_contr$prop_harass)),
    c(eggs_harass_pred_erato_contr$LCL,
        rev(eggs_harass_pred_erato_contr$UCL)),
    border = NA, col = adjustcolor("red",
        0.2), lwd = 2)
lines(eggs_harass_pred_erato_contr$prop_harass,
    eggs_harass_pred_erato_contr$yvar,
    col = "red", lwd = 3)
polygon(c(eggs_harass_pred_erato_treat$prop_harass,
    rev(eggs_harass_pred_erato_treat$prop_harass)),
    c(eggs_harass_pred_erato_treat$LCL,
        rev(eggs_harass_pred_erato_treat$UCL)),
    border = NA, col = adjustcolor("black",
        0.2), lwd = 2)
lines(eggs_harass_pred_erato_treat$prop_harass,
    eggs_harass_pred_erato_treat$yvar,
    col = "black", lwd = 3)

# Repeat the same for H. timareta
plot(jitter(timareta_eggs_harass$prop_harass,
    amount = diff(range(timareta_eggs_harass$prop_harass,
        na.rm = T))/50), jitter(timareta_eggs_harass$eggs_diff,
    amount = diff(range(timareta_eggs_harass$eggs_diff,
        na.rm = T))/50), pch = 21, bg = adjustcolor(ifelse(timareta_eggs_harass$wing_colour ==
    "yellow", "yellow", "red"), 0.75),
    cex = 0.02 * sqrt(timareta_eggs_harass$total_observation_time),
    ylab = "", xlab = "", xaxt = "n",
    yaxt = "n", ylim = range(c(timareta_eggs_harass$eggs_diff,
        eggs_harass_pred_timareta_contr[,
            c("LCL", "UCL")], eggs_harass_pred_timareta_treat[,
            c("LCL", "UCL")]), na.rm = T))
axis(1, lwd = 0, line = -0.5)
axis(1, labels = F, tck = -0.02)
axis(2, las = 2, lwd = 0, line = -0.5)
axis(2, labels = F, tck = -0.02)
mtext(substitute(italic("H. timareta linaresi")),
    3, line = 0.2)

polygon(c(eggs_harass_pred_timareta_contr$prop_harass,
    rev(eggs_harass_pred_timareta_contr$prop_harass)),
    c(eggs_harass_pred_timareta_contr$LCL,
        rev(eggs_harass_pred_timareta_contr$UCL)),
    border = NA, col = adjustcolor("goldenrod",
        0.2), lwd = 2)
lines(eggs_harass_pred_timareta_contr$prop_harass,
    eggs_harass_pred_timareta_contr$yvar,
    col = "goldenrod", lwd = 3)
polygon(c(eggs_harass_pred_timareta_treat$prop_harass,
    rev(eggs_harass_pred_timareta_treat$prop_harass)),
    c(eggs_harass_pred_timareta_treat$LCL,
```

```
        rev(eggs_harass_pred_timareta_treat$UCL)),
    border = NA, col = adjustcolor("red",
        0.2), lwd = 2)
lines(eggs_harass_pred_timareta_treat$prop_harass,
    eggs_harass_pred_timareta_treat$yvar,
    col = "red", lwd = 3)

# Add axes titles
mtext("Eggs with male presence - without male presence",
    2, line = 0, outer = T)
mtext("Proportion of observation time with male harassment",
    1, line = 1.9)

invisible(dev.off())
```

Show png we just made:

```
fig_disp <- readPNG("Figures/empirical_proportion_vs_egg_diff.png")
grid.raster(fig_disp)
```

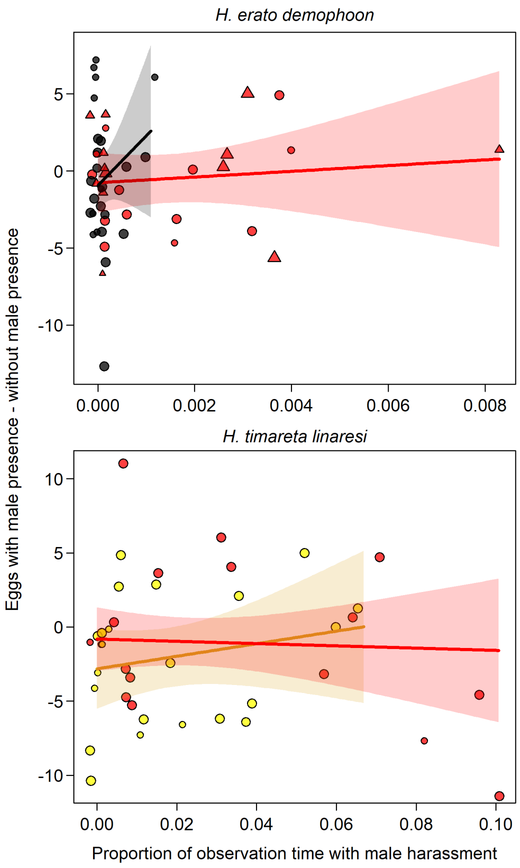


### 5 Session Info

```
si = sessionInfo()
# Remove 'locale' for double-blind
# review
```

```r
si$locale = c(";")
utils:::print.sessionInfo(si)
```

```
## R version 3.6.1 (2019-07-05)
## Platform: x86_64-w64-mingw32/x64 (64-bit)
## Running under: Windows 10 x64 (build 19042)
##
## Matrix products: default
##
## locale:
## [1]
##
## attached base packages:
## [1] grid      stats     graphics  grDevices utils     datasets  methods
## [8] base
##
## other attached packages:
##  [1] truncnorm_1.0-8         brms_2.13.0             Rcpp_1.0.6
##  [4] arm_1.10-1              ENmisc_1.2-7            RColorBrewer_1.1-2
##  [7] vcd_1.4-7               Hmisc_4.4-0             ggplot2_3.3.0
## [10] Formula_1.2-3           survival_3.1-12         lattice_0.20-38
## [13] beeswarm_0.2.3          lme4_1.1-23             Matrix_1.2-18
## [16] png_0.1-7               dominanceanalysis_2.0.0 MASS_7.3-51.5
## [19] emmeans_1.4.6           binom_1.1-1             rmdformats_0.3.7
## [22] knitr_1.28
##
## loaded via a namespace (and not attached):
##   [1] TH.data_1.0-10       minqa_1.2.4          colorspace_1.4-1
##   [4] ellipsis_0.3.0       ggridges_0.5.2       rsconnect_0.8.16
##   [7] estimability_1.3     htmlTable_1.13.3     markdown_1.1
##  [10] base64enc_0.1-3      rstudioapi_0.11      rstan_2.19.3
##  [13] DT_0.13              mvtnorm_1.1-0        bridgesampling_1.0-0
##  [16] codetools_0.2-16     splines_3.6.1        shinythemes_1.1.2
##  [19] bayesplot_1.7.2      jsonlite_1.6.1       nloptr_1.2.2.1
##  [22] cluster_2.1.0        shiny_1.4.0.2        compiler_3.6.1
##  [25] backports_1.1.4      assertthat_0.2.1     fastmap_1.0.1
##  [28] cli_2.4.0            later_1.0.0          formatR_1.7
##  [31] prettyunits_1.1.1    acepack_1.4.1        htmltools_0.4.0
##  [34] tools_3.6.1          igraph_1.2.5         coda_0.19-3
##  [37] gtable_0.3.0         glue_1.4.2           reshape2_1.4.4
##  [40] dplyr_0.8.3          vctrs_0.2.4          nlme_3.1-140
##  [43] crosstalk_1.1.0.1    lmtest_0.9-37        xfun_0.13
##  [46] stringr_1.4.0        ps_1.6.0             mime_0.9
##  [49] miniUI_0.1.1.1       lifecycle_0.2.0      gtools_3.8.2
##  [52] statmod_1.4.34       zoo_1.8-7            scales_1.0.0
##  [55] colourpicker_1.0     promises_1.1.0       Brobdingnag_1.2-6
##  [58] parallel_3.6.1       sandwich_2.5-1       inline_0.3.15
##  [61] shinystan_2.5.0      yaml_2.2.1           gridExtra_2.3
##  [64] StanHeaders_2.21.0-1 loo_2.2.0            rpart_4.1-15
##  [67] latticeExtra_0.6-29  stringi_1.5.3        dygraphs_1.1.1.6
##  [70] checkmate_2.0.0      pkgbuild_1.2.0       boot_1.3-24
##  [73] rlang_0.4.10         pkgconfig_2.0.3      matrixStats_0.56.0
##  [76] evaluate_0.14        purrr_0.3.4          rstantools_2.0.0
##  [79] htmlwidgets_1.5.1    labeling_0.3         processx_3.5.1
```

```
##  [82] tidyselect_1.1.0     plyr_1.8.6          magrittr_2.0.1
##  [85] bookdown_0.18        R6_2.5.0            multcomp_1.4-13
##  [88] pillar_1.4.3         foreign_0.8-76      withr_2.4.1
##  [91] xts_0.12-0           abind_1.4-5         nnet_7.3-13
##  [94] tibble_3.0.1         crayon_1.4.1        rmarkdown_2.1
##  [97] jpeg_0.1-8.1         data.table_1.12.8   callr_3.6.0
## [100] threejs_0.3.3        digest_0.6.25       xtable_1.8-4
## [103] httpuv_1.5.2         stats4_3.6.1        munsell_0.5.0
## [106] shinyjs_1.1
```
