## Supplementary methods and results for "Does sexual conflict contribute to the evolution of novel warning patterns?"

[2] Present address: Department of Evolutionary Neuroethology, Max Planck Institute for Chemical Ecology, Hans-Knöll-Straße 8, 07745 Jena, Germany

#### Supplementary Methods

##### Visual modelling

For experiment 1, to assess visual equivalence of the two control treatments as well as equivalence of a blacked-out red band and black wing color, we applied black and colorless markers (Copic Ciao 100 and 0, respectively) on the red forewing bands of 6 pinned *H. erato demophoon* females. We randomly assigned females into two groups: One group received the blacked-out treatment on the right forewing and the colorless treatment on the left forewing, whereas female in the other group received the reverse. We applied the markers on a rectangular area (~ 0.5 x 1 cm) at the mid-section of the red bands. We then took two photographs of each butterfly with a Nikon Nikkor D7000 camera (Nikon, Melville NY, USA) using a visible light (400 – 680 nm) or a UV (320 – 380 nm) filter (Baader, Mammendorf, Germany). A 40% gray standard (Labsphere, North Sutton NH, USA) was included in each photograph for color calibration. The visible light and UV photos of each wing were combined to generate a multispectral image, which the software (Troscianko and Stevens 2015) then converted to cone catch models based on the visual sensitivity and relative abundance of cone receptors for *H. erato* males (McCulloch et al. 2017; Parnell et al. 2018). Based on the cone catch model results, we calculated pairwise just noticeable differences (JND) between 1) the blacked-out area on the red band and an adjacent black area of similar size, and 2) the area of the red band painted with colorless marker and an adjacent area of similar size where the red band was untreated, using a Weber fraction of 0.05.

**K-means clustering**

For experiment 2, to assess the sizes of the yellow and the red bands in backcross to *H. timareta* individuals, we took photos of the right forewing of 55 females used in the experiments (some females could not be sampled, or had too damaged wings for proper analysis). Photos were taken with a Sony™ Cyber-shot DSC-HX90V camera attached to a tripod (1/6 shutter speed, 5.6 aperture, 58mm zoom, ISO 100, system setting for flash-light white balance, 4896 x 3672 resolution, self-timer), with a conventional bulb and the built-in flash of the camera as light source. The wings were positioned at a fixed distance from the camera on a green cardboard background. An area of 2396 x 1797 pixels was cropped from each photo. PhotoScissors™ software was used to remove the green background.

Photos were read into R (R Core Team 2019) and then compressed in width and height by 4.1-fold (down to 584 x 438 pixels) to speed up the following analyses, using the *imager* package (Barthelme 2020). We then performed k-means clustering on the RGB dimension of the pictures (all transparent background pixels were considered as black). For wings with only a yellow band present, k = 4, and for wings with yellow and red patches, k = 5 clusters were used. These clusters summarized, respectively: the purely black pixels of the background, the (nearly) black scales of the wing, miscellaneous pixels (e.g. damaged parts of the wing), as well as the pixels of the yellow and (if present) of the red band. For photos of 2 yellow-red and of 1 yellow individual, one more cluster had to be applied as a miscellaneous category, *i.e.* k = 5 for yellow and k = 6 for yellow-red (scales in parts of these wings were damaged).

We then visualized the original next to the k-means-clustered image and selected by eye which category matched the yellow and (if present) the red band and used the pixel count of these categories as a proxy for band size. To assess the size of the complete wing, number of transparent (*i.e.* background) pixels was subtracted from total pixel count. We then took a photo of a measuring tape in the spot where wings were positioned, using the same camera settings and manipulating the image with the same pipeline we used for the wing photos. In ImageJ™, we counted how many pixels covered 1 cm of the measuring tape. Considering that the aspect ratio of the pixels on the sensor of the camera was 1:1, this could be used as a scale to translate number of pixels to $cm^2$.

**Removal of outliers from male harassment data**

Counts of seconds with courtship are likely to result in extreme outliers (e.g. if certain males are particularly active). For each experiment and treatment group, we calculated the proportion of time each female was courted, and then removed all females from subsequent analyses with values more than 1.5 units of IQR (interquartile range) above the 75th or below the 25th percentile (highlighted in Fig. S8). However, we also performed the same analyses on the dataset including the outliers (Fig. S8).

### Supplementary Tables

**Table S1: Probability of survival of the novel allele under different parameter combinations.** Probabilities are shown with 95% binomial confidence interval, and red shading is strongest at highest values. Presented parameters include: probability of encounter $e$, strength of attraction $\alpha$, sex of the first mutant, number of loci $N$ encoding for each preference phenotype, presence/absence of bottleneck and the predator learning threshold $Q$. The resulting 288 parameter combinations only include cases where sexual conflict was present and predators returned after 100 generations. Without sexual conflict and/or an earlier return of the predators, the novel allele could never go to fixation. Additionally to these parameters, for each combination of $e$ and $\alpha$ and assuming a patch containing males typical for the first generation with minimum attraction to novel and maximum attraction to ancestral pattern, the expected relative difference in number of offspring sired by a novel patterned female versus an ancestrally patterned female is shown. During predator absence, the novel allele is effectively neutral in males (in terms of direct fitness).

| | | | | $e$ = 0.001 | $e$ = 0.001 | $e$ = 0.0025 | $e$ = 0.0025 | $e$ = 0.005 | $e$ = 0.005 | $e$ = 0.01 | $e$ = 0.01 |
|---|---|---|---|---|---|---|---|---|---|---|---|
| | | | | $\alpha$ = 1 | $\alpha$ = 3 | $\alpha$ = 1 | $\alpha$ = 3 | $\alpha$ = 1 | $\alpha$ = 3 | $\alpha$ = 1 | $\alpha$ = 3 |
| | | | | $\Delta$ = 0.029 | $\Delta$ = 0.043 | $\Delta$ = 0.072 | $\Delta$ = 0.104 | $\Delta$ = 0.141 | $\Delta$ = 0.198 | $\Delta$ = 0.272 | $\Delta$ = 0.36 |
| First Mutant = Female | Loci = 1 | Bottleneck | Q = 1 | - | - | - | - | - | - | 2.4% [1.5-3.5%] | 3.8% [2.7-5.2%] |
| | | | Q = 2 | - | - | - | - | - | - | 0.9% [0.4-1.7%] | 3.7% [2.6-5%] |
| | | | Q = 4 | - | - | - | - | - | - | - | 2.4% [1.5-3.6%] |
| | | No Bottleneck | Q = 1 | - | - | - | - | - | - | 2% [1.2-3.1%] | 23.9% [21.3-26.7%] |
| | | | Q = 2 | - | - | - | - | - | - | - | 16.2% [14-18.6%] |
| | | | Q = 4 | - | - | - | - | - | - | - | 6% [4.6-7.6%] |
| | Loci = 5 | Bottleneck | Q = 1 | - | - | - | - | - | - | 2.3% [1.5-3.4%] | 3.5% [2.5-4.9%] |
| | | | Q = 2 | - | - | - | - | - | - | 1.2% [0.6-2.1%] | 3.9% [2.8-5.3%] |
| | | | Q = 4 | - | - | - | - | - | - | 0.2% [0-0.7%] | 2.3% [1.5-3.4%] |
| | | No Bottleneck | Q = 1 | - | - | - | - | - | - | 1.2% [0.6-2.1%] | 23.9% [21.3-26.7%] |
| | | | Q = 2 | - | - | - | - | - | - | - | 18.9% [16.5-21.4%] |
| | | | Q = 4 | - | - | - | - | - | - | - | 5.1% [3.8-6.6%] |
| | Loci = 10 | Bottleneck | Q = 1 | - | - | - | - | - | - | 1.9% [1.1-2.9%] | 4.2% [3.1-5.7%] |
| | | | Q = 2 | - | - | - | - | - | - | 0.4% [0.1-1%] | 3.8% [2.7-5.2%] |
| | | | Q = 4 | - | - | - | - | - | - | 0.2% [0-0.7%] | 2.7% [1.8-3.9%] |
| | | No Bottleneck | Q = 1 | - | - | - | - | - | - | 2.3% [1.5-3.4%] | 26% [23.3-28.9%] |
| | | | Q = 2 | - | - | - | - | - | - | - | 15.8% [13.6-18.2%] |
| | | | Q = 4 | - | - | - | - | - | - | - | 4.3% [3.1-5.7%] |
| First Mutant = Male | Loci = 1 | Bottleneck | Q = 1 | - | - | - | - | - | 0.1% [0-0.6%] | 2.9% [2-4.2%] | 4.6% [3.4-6%] |
| | | | Q = 2 | - | - | - | - | - | - | 1.2% [0.6-2.1%] | 3.4% [2.4-4.7%] |
| | | | Q = 4 | - | - | - | - | - | - | 0.2% [0-0.7%] | 3% [2-4.2%] |
| | | No Bottleneck | Q = 1 | - | - | - | - | - | - | 0.9% [0.4-1.7%] | 9.8% [8-11.8%] |
| | | | Q = 2 | - | - | - | - | - | - | - | 4.9% [3.7-6.5%] |
| | | | Q = 4 | - | - | - | - | - | - | - | 1.8% [1.1-2.9%] |
| | Loci = 5 | Bottleneck | Q = 1 | - | - | - | - | - | 0.1% [0-0.6%] | 2.7% [1.8-3.9%] | 4.3% [3.1-5.7%] |
| | | | Q = 2 | - | - | - | - | - | - | 0.4% [0.1-1%] | 4.9% [3.7-6.4%] |
| | | | Q = 4 | - | - | - | - | - | - | 0.4% [0.1-1%] | 2.1% [1.3-3.2%] |
| | | No Bottleneck | Q = 1 | - | - | - | - | - | - | 0.4% [0.1-1%] | 9.8% [8-11.8%] |
| | | | Q = 2 | - | - | - | - | - | - | - | 5.9% [4.5-7.6%] |
| | | | Q = 4 | - | - | - | - | - | - | - | 2.3% [1.4-3.4%] |
| | Loci = 10 | Bottleneck | Q = 1 | - | - | - | - | - | - | 3.5% [2.5-4.9%] | 4.5% [3.3-5.9%] |
| | | | Q = 2 | - | - | - | - | - | - | 0.7% [0.3-1.4%] | 4.8% [3.6-6.3%] |
| | | | Q = 4 | - | - | - | - | - | - | 0.1% [0-0.6%] | 3.2% [2.2-4.5%] |
| | | No Bottleneck | Q = 1 | - | - | - | - | - | - | 0.5% [0.2-1.2%] | 10.6% [8.8-12.6%] |
| | | | Q = 2 | - | - | - | - | - | - | - | 6.1% [4.7-7.8%] |
| | | | Q = 4 | - | - | - | - | - | - | - | 1.4% [0.8-2.4%] |

**Table S2: Type III Anova results for the model explaining survival of the novel pattern allele.** p-values were corrected for multiple testing using the Bonferroni correction. Parameter annotation as in Tables 1 and S1. Presence/absence of sexual conflict as well as the time for the return of the predictors were not included in the model, to avoid complete separation. However, both parameters had the biggest effect on simulation outcome.

| **Model Term** | ***d.f.*** | **F ratio** | ***p*-value** | $R^2$ |
|---|---|---|---|---|
| *e* | 3 | 36.57 | < 0.001 | 0.634 |
| $\alpha$ : bottleneck | 1 | 304.385 | < 0.001 | 0.089 |
| $\alpha$ : *Q* | 2 | 56.039 | < 0.001 | 0.084 |
| $\alpha$ | 1 | 569.593 | < 0.001 | 0.056 |
| bottleneck : sex_first_mutant | 1 | 220.306 | < 0.001 | 0.035 |
| *Q* : bottleneck | 2 | 58.521 | < 0.001 | 0.034 |
| *Q* | 2 | 160.863 | < 0.001 | 0.020 |
| sex_first_mutant | 1 | 118.329 | < 0.001 | 0.010 |
| bottleneck | 1 | 31.493 | < 0.001 | 0.009 |

### Supplementary Figures

**Figure S1: Phenotypic effect of male attraction traits dependent on genomic architecture and male strength of attraction to colors.** y-axis is log-scaled and shows the x-fold increase in probability (compared to minimum attraction) that male interest is elicited by a female pattern, dependent on number of alleles conferring attraction in the male. This relationship is shown for different genetic architectures of male preference, and different strengths of attraction to colors in males.

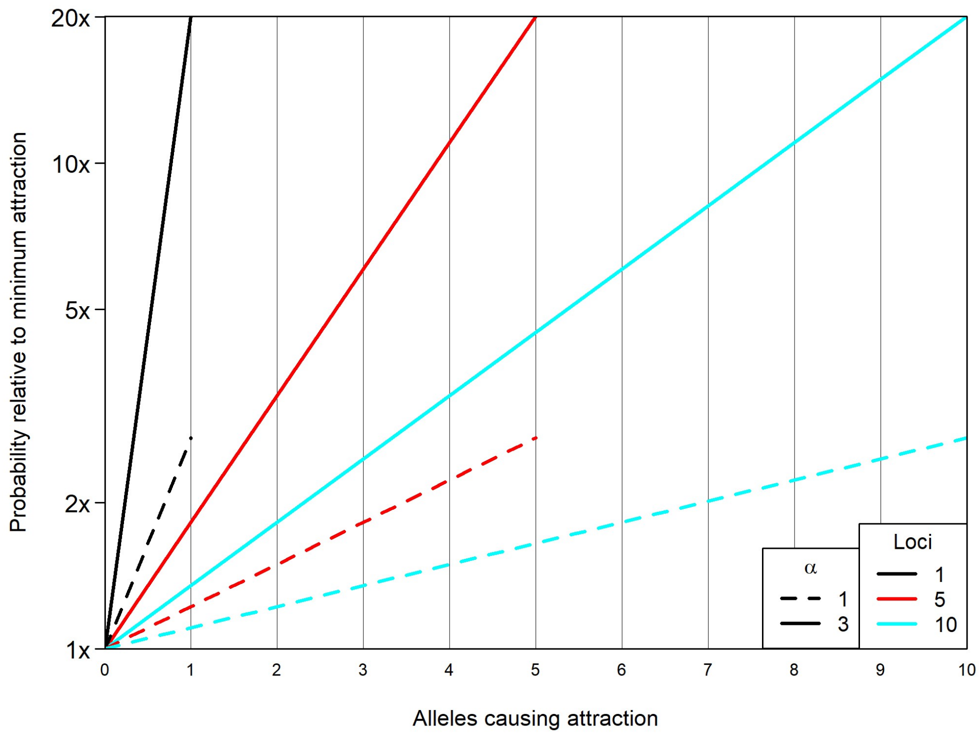


**Figure S2: Positioning of first mutant in the arena affects survival of the novel mutation.** Position of first mutants in the central (initially predation-free) zone of the arena from simulation runs where the novel pattern survived. The furthest away the novel mutation was placed from the area where predation is not relaxed in the beginning, the more likely it is that it survives.

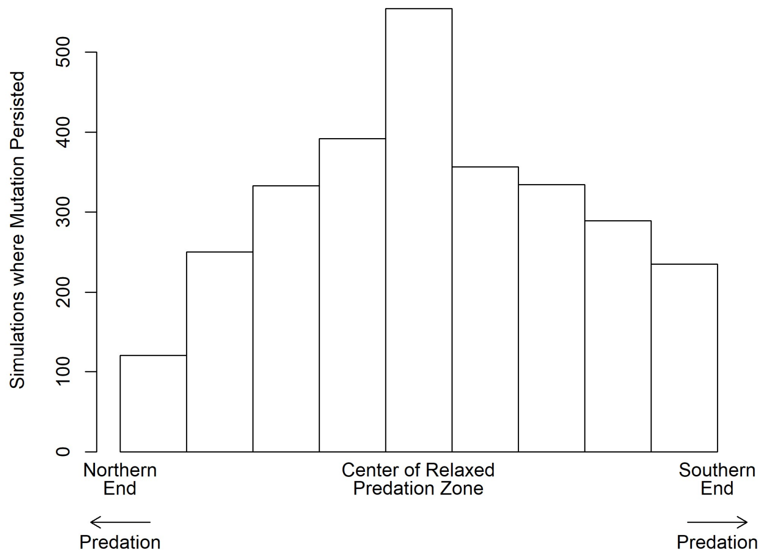

**Figure S3: Frequency of novel pattern allele in central patches at generation 100 has to be high before predators return.** Only data from simulations where predators return after generation 100 are shown, across different other simulation scenarios. A: Allele frequency of novel pattern. B: Phenotype frequency of novel pattern. Cases where the novel pattern allele survived until generation 2500 are shown as red dots; cases where it did not survive are either from scenarios involving sexual conflict (= black dots) or not involving sexual conflict (= gray dots).

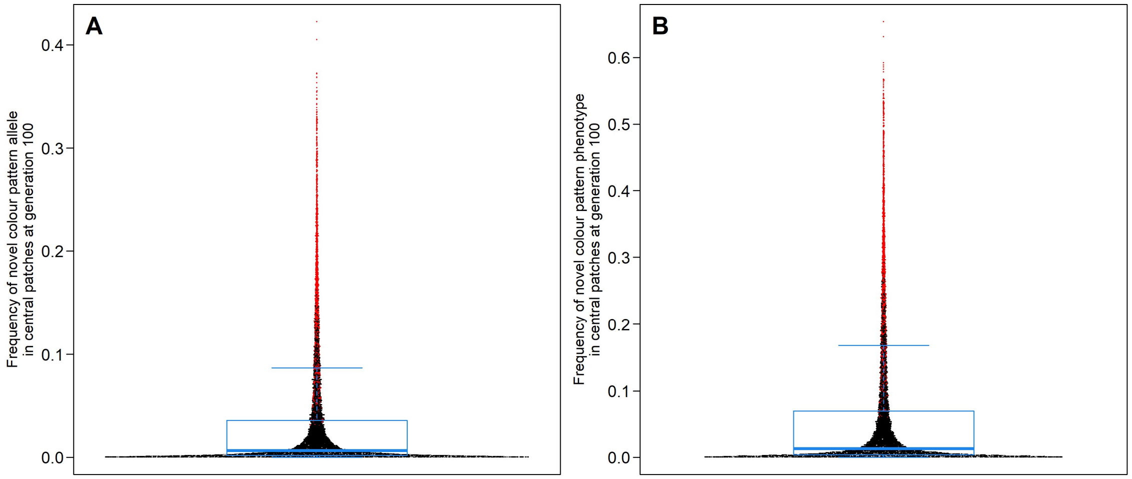


**Figure S4: Genetic architecture of male attraction affects 'lag' between (mean) allele frequencies for novel pattern and for attraction to this pattern across simulation scenarios.** Allele frequencies of novel alleles over generation time. All 'scenarios' that ever resulted in the novel pattern allele surviving until generation 2500 are displayed. Parameter settings (except number of loci encoding attraction traits) are displayed on the left of each row; number of loci encoding attraction traits are displayed on top of each row. Allele frequencies either correspond to novel alleles at the warning pattern locus (gray), at loci encoding male attraction to novel pattern (orange; novel alleles increase attraction) or at loci encoding for male attraction to ancestral pattern (blue; novel alleles reduce attraction). For the latter two, mean allele frequencies were calculated, whenever multiple loci were involved. x-axis is log-scaled. Vertical line at 100 generations shows where period of relaxed predation ends.

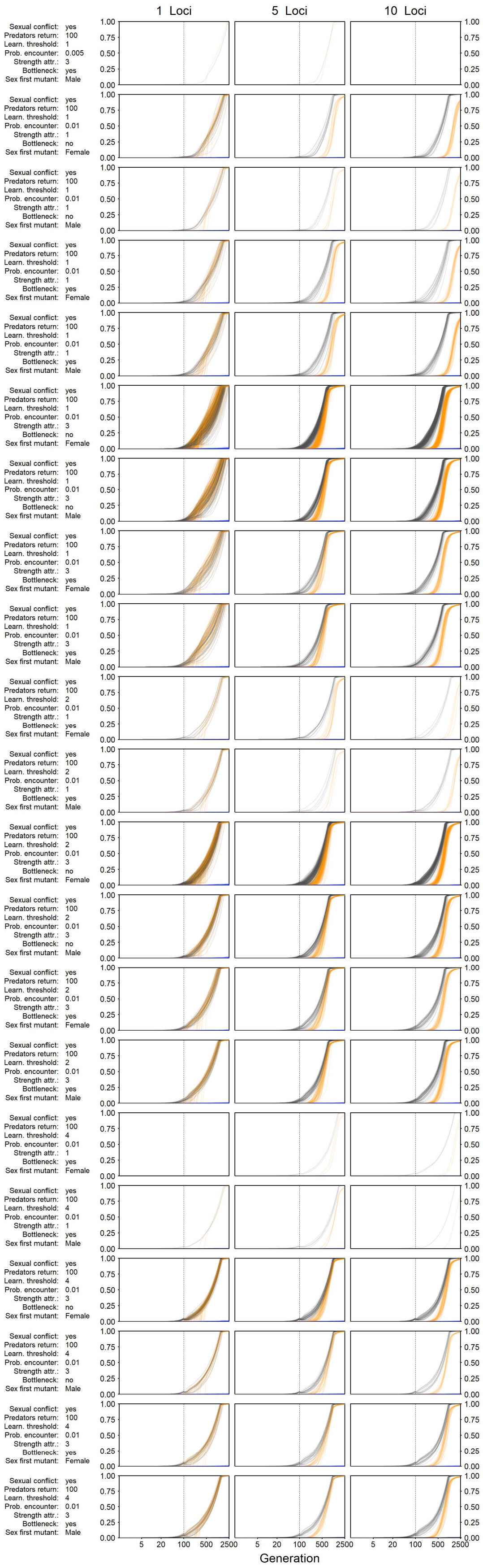
1 Loci
5 Loci
10 Loci
Sexual conflict: yes
Predators return: 100
Learn. threshold: 1
Prob. encounter: 0.005
Strength attr.: 3
Bottleneck: yes
Sex first mutant: Male
Sexual conflict: yes
Predators return: 100
Learn. threshold: 1
Prob. encounter: 0.01
Strength attr.: 1
Bottleneck: no
Sex first mutant: Female
Sexual conflict: yes
Predators return: 100
Learn. threshold: 1
Prob. encounter: 0.01
Strength attr.: 1
Bottleneck: no
Sex first mutant: Male
Sexual conflict: yes
Predators return: 100
Learn. threshold: 1
Prob. encounter: 0.01
Strength attr.: 1
Bottleneck: yes
Sex first mutant: Female
Sexual conflict: yes
Predators return: 100
Learn. threshold: 1
Prob. encounter: 0.01
Strength attr.: 1
Bottleneck: yes
Sex first mutant: Male
Sexual conflict: yes
Predators return: 100
Learn. threshold: 1
Prob. encounter: 0.01
Strength attr.: 3
Bottleneck: no
Sex first mutant: Female
Sexual conflict: yes
Predators return: 100
Learn. threshold: 1
Prob. encounter: 0.01
Strength attr.: 3
Bottleneck: no
Sex first mutant: Male
Sexual conflict: yes
Predators return: 100
Learn. threshold: 1
Prob. encounter: 0.01
Strength attr.: 3
Bottleneck: yes
Sex first mutant: Female
Sexual conflict: yes
Predators return: 100
Learn. threshold: 1
Prob. encounter: 0.01
Strength attr.: 3
Bottleneck: yes
Sex first mutant: Male
Sexual conflict: yes
Predators return: 100
Learn. threshold: 2
Prob. encounter: 0.01
Strength attr.: 1
Bottleneck: yes
Sex first mutant: Female
Sexual conflict: yes
Predators return: 100
Learn. threshold: 2
Prob. encounter: 0.01
Strength attr.: 1
Bottleneck: yes
Sex first mutant: Male
Sexual conflict: yes
Predators return: 100
Learn. threshold: 2
Prob. encounter: 0.01
Strength attr.: 3
Bottleneck: no
Sex first mutant: Female
Sexual conflict: yes
Predators return: 100
Learn. threshold: 2
Prob. encounter: 0.01
Strength attr.: 3
Bottleneck: no
Sex first mutant: Male
Sexual conflict: yes
Predators return: 100
Learn. threshold: 2
Prob. encounter: 0.01
Strength attr.: 3
Bottleneck: yes
Sex first mutant: Female
Sexual conflict: yes
Predators return: 100
Learn. threshold: 2
Prob. encounter: 0.01
Strength attr.: 3
Bottleneck: yes
Sex first mutant: Male
Sexual conflict: yes
Predators return: 100
Learn. threshold: 4
Prob. encounter: 0.01
Strength attr.: 1
Bottleneck: yes
Sex first mutant: Female
Sexual conflict: yes
Predators return: 100
Learn. threshold: 4
Prob. encounter: 0.01
Strength attr.: 1
Bottleneck: yes
Sex first mutant: Male
Sexual conflict: yes
Predators return: 100
Learn. threshold: 4
Prob. encounter: 0.01
Strength attr.: 3
Bottleneck: no
Sex first mutant: Female
Sexual conflict: yes
Predators return: 100
Learn. threshold: 4
Prob. encounter: 0.01
Strength attr.: 3
Bottleneck: no
Sex first mutant: Male
Sexual conflict: yes
Predators return: 100
Learn. threshold: 4
Prob. encounter: 0.01
Strength attr.: 3
Bottleneck: yes
Sex first mutant: Female
Sexual conflict: yes
Predators return: 100
Learn. threshold: 4
Prob. encounter: 0.01
Strength attr.: 3
Bottleneck: yes
Sex first mutant: Male
0.00
0.25
0.50
0.75
1.00
5
20
100
500
2500
Generation

**Figure S5: Males never harass the novel pattern more than the ancestral pattern.** The ‘phenotype ratios’ resulting from Fig S4, *i.e.* number individuals with novel pattern divided by ancestral pattern (gray) and mean male attraction to novel pattern divided by mean attraction to ancestral pattern (turquoise). Other annotations are as in Fig. S4. Four different phases are displayed with different background color intensity, corresponding to the different forces of selection acting: 1) no natural selection in central patches and selection from male harassment favors novel pattern; 2) natural selection favors ancestral pattern and selection from male harassment favors novel pattern; 3) both forces of selection favor the novel pattern; 4) natural selection favors ancestral pattern and selection from male harassment is quasi-absent. x- and y-axis are log-scaled. Note that two different y-axis scales were applied (color of labels corresponds to the respective ratio).

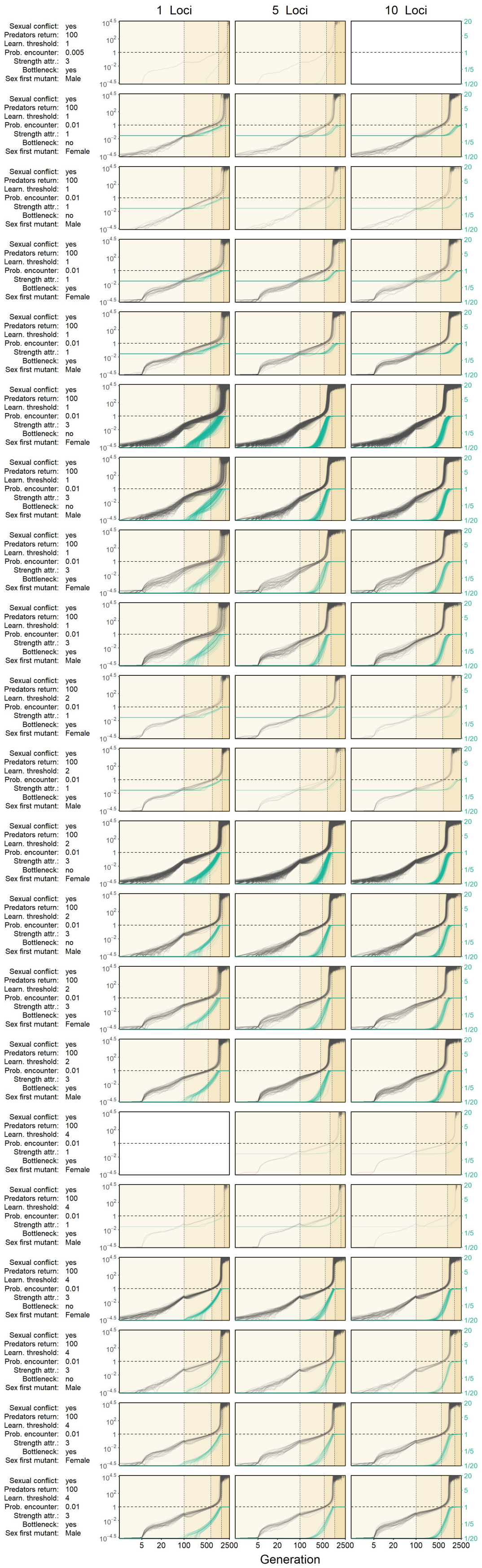
1 Loci
5 Loci
10 Loci
Sexual conflict: yes
Predators return: 100
Learn. threshold: 1
Prob. encounter: 0.005
Strength attr.: 3
Bottleneck: yes
Sex first mutant: Male
Sexual conflict: yes
Predators return: 100
Learn. threshold: 1
Prob. encounter: 0.01
Strength attr.: 1
Bottleneck: no
Sex first mutant: Female
Sexual conflict: yes
Predators return: 100
Learn. threshold: 1
Prob. encounter: 0.01
Strength attr.: 1
Bottleneck: no
Sex first mutant: Male
Sexual conflict: yes
Predators return: 100
Learn. threshold: 1
Prob. encounter: 0.01
Strength attr.: 1
Bottleneck: yes
Sex first mutant: Female
Sexual conflict: yes
Predators return: 100
Learn. threshold: 1
Prob. encounter: 0.01
Strength attr.: 1
Bottleneck: yes
Sex first mutant: Male
Sexual conflict: yes
Predators return: 100
Learn. threshold: 1
Prob. encounter: 0.01
Strength attr.: 3
Bottleneck: no
Sex first mutant: Female
Sexual conflict: yes
Predators return: 100
Learn. threshold: 1
Prob. encounter: 0.01
Strength attr.: 3
Bottleneck: no
Sex first mutant: Male
Sexual conflict: yes
Predators return: 100
Learn. threshold: 1
Prob. encounter: 0.01
Strength attr.: 3
Bottleneck: yes
Sex first mutant: Female
Sexual conflict: yes
Predators return: 100
Learn. threshold: 1
Prob. encounter: 0.01
Strength attr.: 3
Bottleneck: yes
Sex first mutant: Male
Sexual conflict: yes
Predators return: 100
Learn. threshold: 2
Prob. encounter: 0.01
Strength attr.: 1
Bottleneck: yes
Sex first mutant: Female
Sexual conflict: yes
Predators return: 100
Learn. threshold: 2
Prob. encounter: 0.01
Strength attr.: 1
Bottleneck: yes
Sex first mutant: Male
Sexual conflict: yes
Predators return: 100
Learn. threshold: 2
Prob. encounter: 0.01
Strength attr.: 3
Bottleneck: no
Sex first mutant: Female
Sexual conflict: yes
Predators return: 100
Learn. threshold: 2
Prob. encounter: 0.01
Strength attr.: 3
Bottleneck: no
Sex first mutant: Male
Sexual conflict: yes
Predators return: 100
Learn. threshold: 2
Prob. encounter: 0.01
Strength attr.: 3
Bottleneck: yes
Sex first mutant: Female
Sexual conflict: yes
Predators return: 100
Learn. threshold: 2
Prob. encounter: 0.01
Strength attr.: 3
Bottleneck: yes
Sex first mutant: Male
Sexual conflict: yes
Predators return: 100
Learn. threshold: 4
Prob. encounter: 0.01
Strength attr.: 1
Bottleneck: yes
Sex first mutant: Female
Sexual conflict: yes
Predators return: 100
Learn. threshold: 4
Prob. encounter: 0.01
Strength attr.: 1
Bottleneck: yes
Sex first mutant: Male
Sexual conflict: yes
Predators return: 100
Learn. threshold: 4
Prob. encounter: 0.01
Strength attr.: 3
Bottleneck: no
Sex first mutant: Female
Sexual conflict: yes
Predators return: 100
Learn. threshold: 4
Prob. encounter: 0.01
Strength attr.: 3
Bottleneck: no
Sex first mutant: Male
Sexual conflict: yes
Predators return: 100
Learn. threshold: 4
Prob. encounter: 0.01
Strength attr.: 3
Bottleneck: yes
Sex first mutant: Female
Sexual conflict: yes
Predators return: 100
Learn. threshold: 4
Prob. encounter: 0.01
Strength attr.: 3
Bottleneck: yes
Sex first mutant: Male
$10^{4.5}$
$10^{2}$
1
$10^{-2}$
$10^{-4.5}$
20
5
1
1/5
1/20
5 20 100 500 2500
Generation

**Figure S6: Effect of genetic architecture of preference phenotypes on (mean) allele frequencies at patterning or attraction loci at the moment of maximum allele frequency of the novel allele.** Cases where mutation survived until the last generation are highlighted in red (and their proportion is indicated above the graph). y-axis is log-scaled. Frequency of novel color pattern allele at scenarios of 1, 5 or 10 loci per preference phenotype (first row). In scenarios with one preference locus (*i.e.* male adaptations to female novelty is easiest), it seems as if the novel mutation has a slightly harder time reaching fixation, and may died out more often after having already reached a respectable frequency. Male preference phenotypes can evolve more easily to strongest attraction to novel pattern when there is fewer loci involved (second row). The preference phenotype for the ancestral pattern seems to not change much, independent of the scenario (third row).

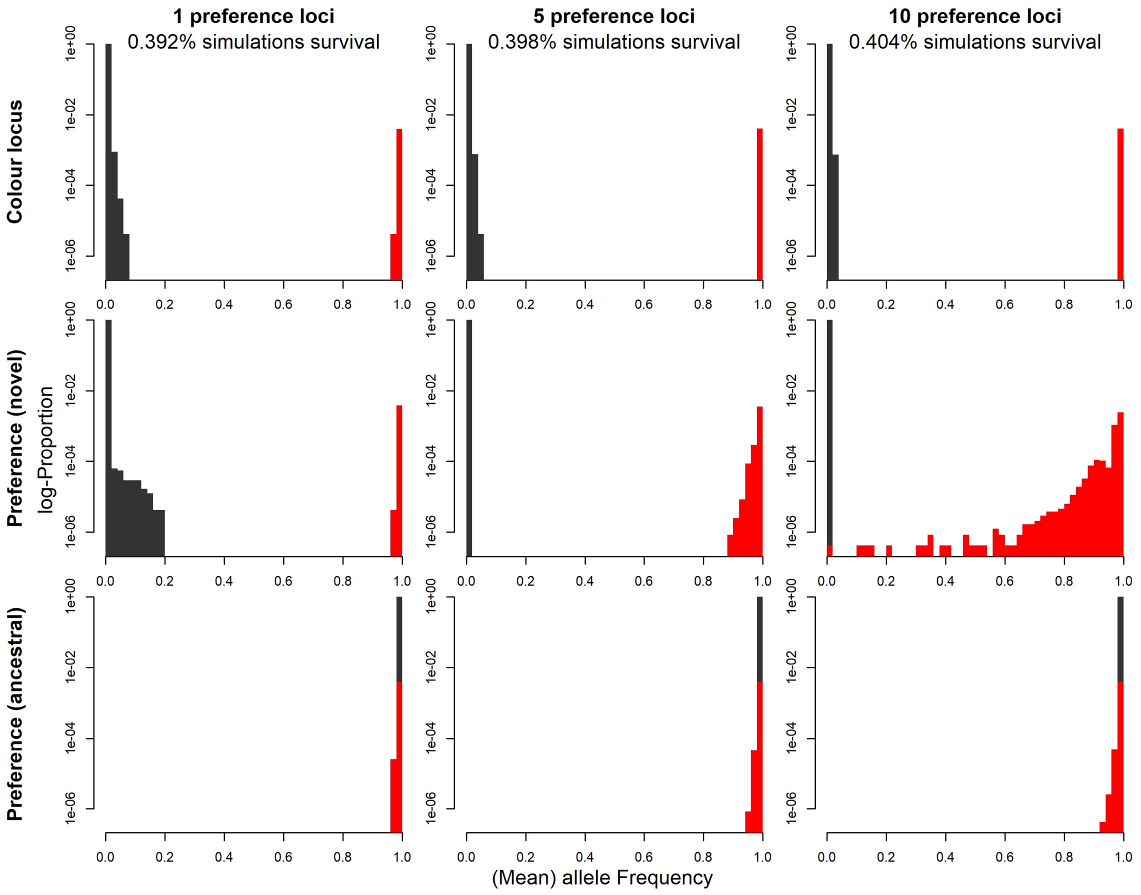

**Figure S7: Transparently-painted red appears as red wing color (and blacked-out red pattern appears as black wing color) to the *H. erato* visual system.** Just noticeable differences (JND) as predicted by a model of the *H. erato* visual system. All values lie well within the range that is commonly assumed to be non-distinguishable.

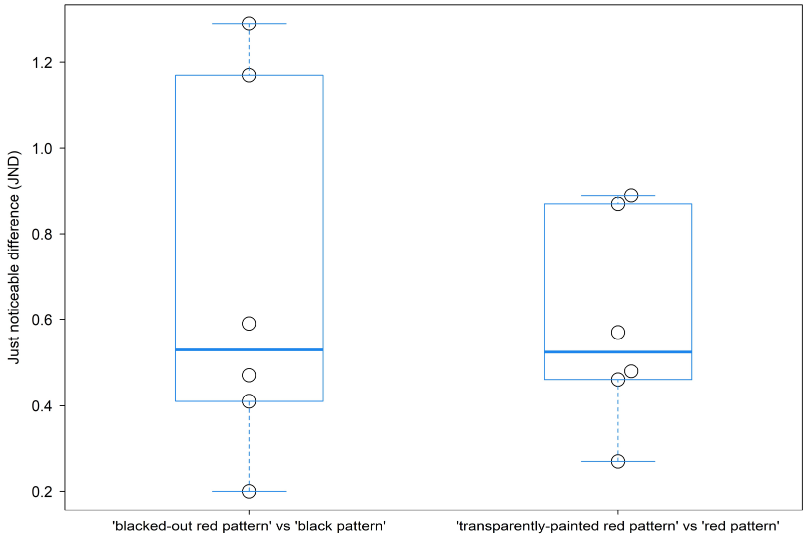


**Figure S8: Size of yellow and of red band on backcross to *H. timareta* wings.** Absolute area of yellow and red band (top) and area as relative to whole forewing (bottom). Wings are displayed at the respective x- and y-coordinates in the plots. Sizes of the displayed wings are scaled to the actual sizes of the images. Each wing received a white outline, which helps identifying where a wing covers a wing below it.

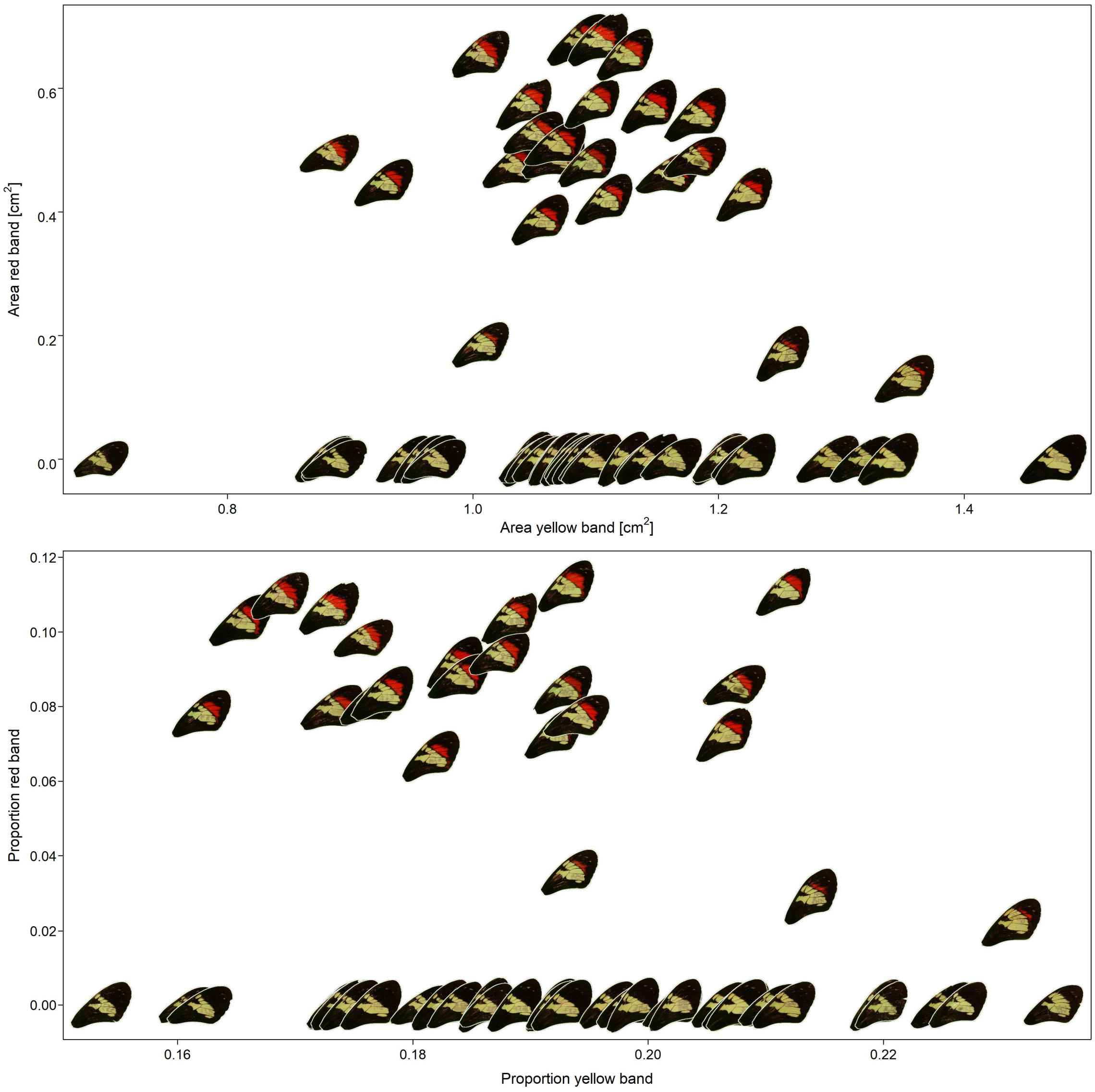


**Figure S9: Including outliers in male harassment analyses qualitatively affects results from experiment 2.** Same as Fig. 3A+C, but including data from 'outlier' females. 'Outlier females' are highlighted with circles/arrows. One female is not shown as its y-value was much larger than of any other female in experiment 1. Estimated marginal means and p-values are based on a model fitted to the data including the outliers.

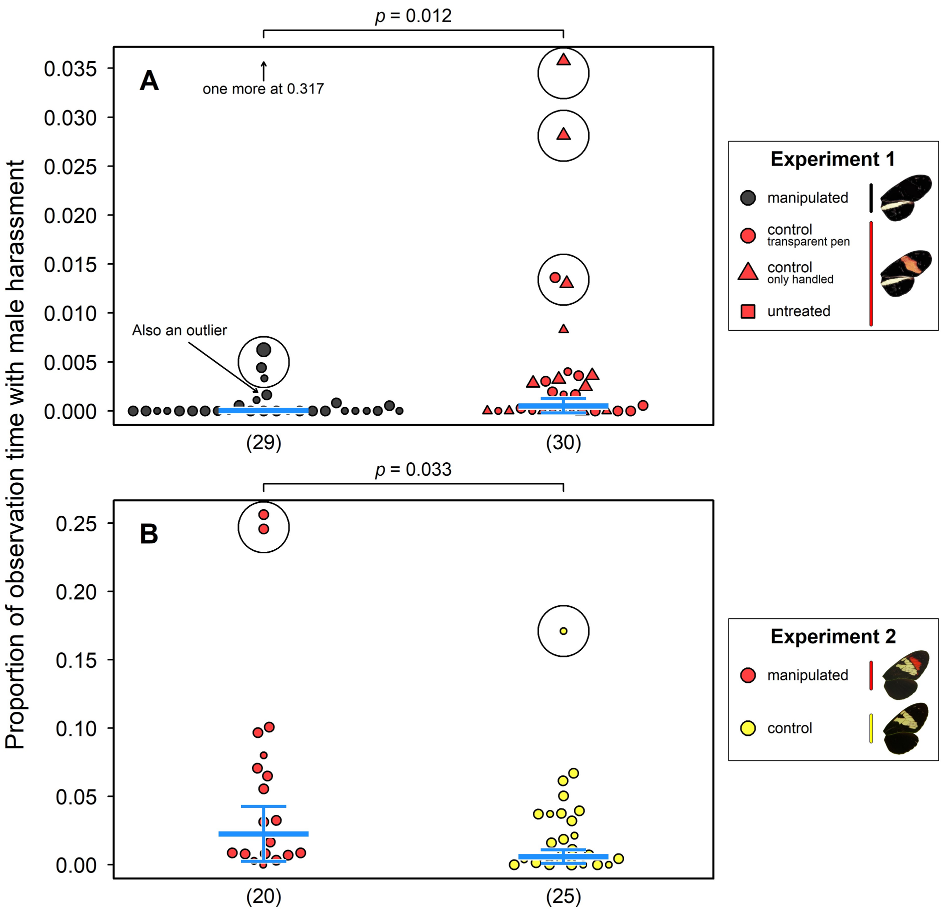


**Fig. S10: Size of the red band (proportional to whole wing) in yellow-red backcross to *H. timareta* hybrids affects male harassment.** Proportion of total observation time males spent harassing yellow-red backcross to *H. timareta* females as a function of red band size (A) and yellow band size (B). Size of bands is considered as relative to total wing area. Area of dots is relative to total number of observed seconds. Estimated marginal means are shown as a line and their confidence intervals (CIs) in gray. Note that the same outliers indicated in Fig. S9B were excluded from this analysis. Band size measurements from 6 of the 18 females from this treatment group were not available.

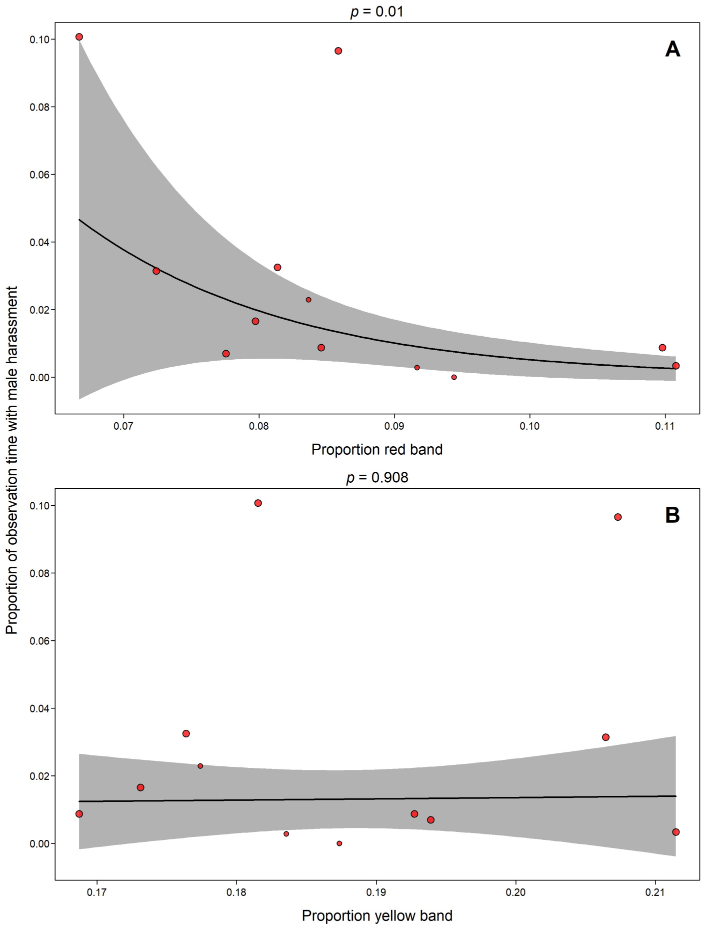


**Fig. S11: Hatching success is independent of male presence.** Number of eggs laid by differently colored females of experiment 2 and corresponding number of hatchlings in the presence (triangles) or absence (circles) of males. Dots were minimally jittered in x- and y-direction. For clarity, one data point was not included, specifically a female that laid 23 eggs in a two-day period (the only data point lying outside 1.5x the interquartile range for number of eggs). Yellow-red ('novel-colored') females have red symbols, yellow ('control') females have yellow symbols. Lines were determined by local polynomial regression fitting. Black line = all data, green line = male absence, blue line = male presence. Number of eggs and number of hatchlings is correlated. However, above a certain number of eggs laid, it seems that number of hatchlings stagnates. This could either mean that females that lay 'too much' invest less in the quality of eggs, or it could be an artefact of data collection, as we stored collected eggs in small plastic containers, where siblings hatched together. Young hatchlings can be cannibalistic, eating their siblings or eggs containing their siblings, which might influence our hatchling counts. The proportion of this might grow with number of hatchlings in a container.

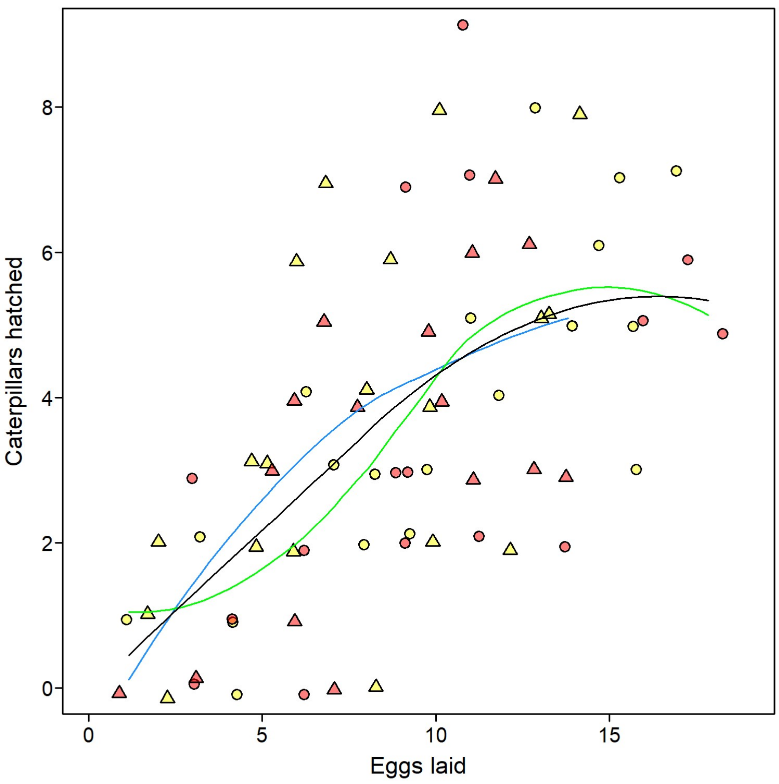

Caterpillars hatched
Eggs laid
0
2
4
6
8
0
5
10
15
